# Synaptic decline precedes dopaminergic neuronal loss in human midbrain organoids harboring a triplication of the *SNCA* gene

**DOI:** 10.1101/2021.07.15.452499

**Authors:** Jennifer Modamio, Claudia Saraiva, Gemma Gomez Giro, Sarah Louise Nickels, Javier Jarazo, Paul Antony, Peter Barbuti, Rashi Hadler, Christian Jäger, Rejko Krüger, Enrico Glaab, Jens Christian Schwamborn

## Abstract

Increased levels of the protein alpha-synuclein (α-syn) are associated with the development of neurodegenerative diseases like Parkinson’s disease (PD). In physiological conditions, α-syn modulates synaptic plasticity, neurogenesis and neuronal survival. However, its pathogenic accumulation and aggregation results in toxicity and neurodegeneration.

Here, we used a PD patient specific midbrain organoid model derived from induced pluripotent stem cells harboring a triplication in the *SNCA* gene to study PD-associated phenotypes. The model recapitulates the two main hallmarks of PD, which are α-syn aggregation and loss of dopaminergic neurons. Additionally, impairments in astrocyte differentiation were detected. Transcriptomics data indicate that synaptic function is impaired in PD specific midbrain organoids. This is further confirmed by alterations in synapse number and electrophysiological activity. We found that synaptic decline precedes neurodegeneration. Finally, this study substantiates that patient specific midbrain organoids allow a personalized phenotyping, which make them an interesting tool for precision medicine and drug discovery.

## 2. Introduction

Parkinson’s disease (PD), the second most common neurodegenerative disorder, is characterized by the loss of dopaminergic neurons in the *substantia nigra* (SN) *pars compacta* (Foix and Nicolesco, 1925; Hassler, 1938) and by the presence of proteinaceous aggregates known as Lewy bodies and Lewy neurites (Lewy, 1912). PD is thought to be caused mostly by an interplay of genetic-, environmental- and age-related factors. The *SNCA* gene encodes for the alpha-synuclein protein (α-syn), which was identified as the main component of Lewy bodies and Lewy neurites (Goedert et al., 2013). Multiplications and point mutations of *SNCA* are causative of familial PD (Appel-Cresswell et al., 2013; Ghosh et al., 2013; Krüger, 1998; Lesage et al., 2013; Polymeropoulos et al., 1997; Zarranz et al., 2004). Individuals carrying a *SNCA* triplication present full PD penetrance, likely caused by higher α-syn levels, which increases the propensity of α-syn to form oligomers, fibrils, and aggregates with cytotoxic effects (Bousset et al., 2013; Lashuel, 2013; Tsigelny et al., 2012).

Under physiological conditions, α-syn is mostly present in the pre-synaptic terminal contributing to its essential functions in synaptic vesicle trafficking, neurotransmitter release and chaperon activity (Burré, 2015; Jensen et al., 1998). In addition, several reports suggested α-syn implication in neuronal differentiation and regulation of the dopaminergic system (Butler et al., 2017; Pavlou et al., 2017). In fact, lack of α-syn leads to a reduced number of dopaminergic neurons during embryonic development (Garcia-Reitboeck et al., 2013). A-syn also regulates the maintenance of adult neural stem cells in the subventricular zone, and dopamine levels in the SN (Perez-Villalba et al., 2018). However, the structural flexibility of α-syn allows the adoption of several conformations, which favors oligomerization and aggregation that may result in neurotoxicity and degeneration (Ghiglieri et al., 2018). To date, despite a large number of studies on α-syn functionality in the synaptic terminal, the exact role of α-syn in neurogenesis and neurodegeneration remains poorly understood. Therefore, it is key to better understand the interplay between α-syn and neurogenesis.

One of the main limitations in the study of PD is the lack of models that properly recapitulate the key hallmarks of the disease. We and others have previously developed human midbrain organoid (MO) models that mirror key characteristics of PD (Kim et al., 2019; Monzel et al., 2017; Nickels et al., 2020; Smits et al., 2019). These MO models contain high numbers of dopaminergic neurons, the most vulnerable cell population in PD. This enrichment of DA neurons allows the study of neurogenesis, neurodegeneration, and neuron associated mechanism (e.g. synapse formation and synaptic activity) in a context resembling the cellular composition and structural organization of the human midbrain. In addition, these MOs also enable the study of other cell types of interest, including astrocytes and oligodendrocytes.

Here, we describe the generation of PD patient specific MOs harboring a triplication in the *SNCA* gene. These MOs showed increased intracellular and extracellular levels of total α-syn, as well as increased intracellular phosphorylated (pS129) α-syn. Additionally, MOs mimic the subcellular translocation of α-syn from the cytosol to the synaptic terminal observed in full polarized neurons, where it co-localizes with the presynaptic marker synaptophysin. Our results indicate that increased α-syn levels influence synapse number and function. Electrophysiological activity analysis revealed an early boost in synaptic function followed by a decline over time, which coincides with a reduction synapse number. Accordingly, MOs harboring a triplication of the *SNCA* gene show an accelerated dopaminergic neuron differentiation, followed by the characteristic dopaminergic neuron loss observed in PD patients.

Overall, we showed that human iPSC-derived midbrain organoids can recapitulate key molecular features of PD pathogenesis (i.e. α-syn accumulation and dopaminergic neurodegeneration). Moreover, using patient specific lines we showed that α-syn triplication influences dopaminergic neurogenesis, namely by showing faster differentiation and maturation, which is followed by a quicker decline. Dopaminergic neuron decline is preceded by alterations in the gene expression signature, electrophysiological activity and synapse number. These data shall contribute to explain the earlier onset age seen on these PD patients, but also to the development of novel and personalized models and treatments for PD.

## 3. Results

### 3.1 MOs recapitulate midbrain main features

In this study, we generated MOs from PD patient specific induced pluripotent stem cells (iPSCs) with a triplication in the *SNCA* gene. We included two independent clones from a PD patient harboring a triplication in the *SNCA* gene (3xSNCA), age- and gender-matched control iPSCs from four healthy donors, and one engineered *SNCA* Knock out (SNCA-KO) line (Table. S1). iPSCs were characterized for common stemness markers via immunocytochemistry and reverse transcription-quantitative polymerase chain reaction (RT-qPCR). iPSCs were positive for the stemness markers SRY-Box Transcription Factor 2 (SOX2), TRA-1-81 (Fig. S1), the octamer-binding transcription factor 4 (OCT4), TRA-1-60 (Fig. S2), NANOG, and SSEA-4 (Fig. S3). Expression of *SOX2* and *OCT4* was further confirmed via RT-qPCR (Fig. S4A, B) (Table.S2). Fluorescence in-situ hybridization (FISH) analysis was performed confirming the presence of additional copies of the *SNCA* gene in both 3xSNCA lines (DB317 and DB336) (Fig. S4C).

iPSCs were used to derive midbrain floor plate neural progenitor cells (mfNPCs) as described previously (Smits et al., 2019). Characterization via immunostaining showed positive signal for the stemness marker SOX2 and the neuroectodermal stem cell marker (NESTIN) (Fig. S5A). mfNPCs were cultured until homogeneous and defined colonies were present (Fig. S5B). mfNPCs were subsequently used for MOs generation following the recently published optimized protocol, which ensures reduced variability and absence of a necrotic core (Nickels et al., 2020).

MOs generated from mfNCPs show a regional organization with a central stem cell niche enriched in progenitor cells, and mature differentiated cells in the periphery (Fig. S6A, B). Progenitor cells at the MOs core expressed classic midbrain floor plate markers such as FOXA2, the LIM Homeobox Transcription Factor 1 Alpha (LMX1A), and the Engrailed Homeobox 1 (EN1) (Fig. 6C).

Increased vulnerability of dopaminergic neurons in PD is one of the main hallmarks of the disease. Therefore, the cellular composition of MOs, with a specific focus on the number of dopaminergic neurons, was characterized at different time points. We started the analysis at an early immature state (day 15 of culture) until a more mature state (day 90 of culture).

Dopaminergic neurons were identified by tyrosine hydroxylase (TH) staining (Fig. S6A, B, D). Importantly, we identified dopaminergic neurons specific for the A9 region of the midbrain which are double positive for TH and the G-protein-regulated inwardly rectifying potassium channel 2 (GIRK2) expression, as well as those specific for the A10 region, double positive for TH and the marker Calbindin (Fig. S6D). In addition, positive immunocytochemistry signals for DOPA decarboxylase (DDC) and dopamine neurotransmitter confirmed dopamine synthesis and presence in MOs (Fig. S6D).

### 3.2 3xSNCA MOs show increased levels of total and pS129 alpha-synuclein

Higher levels and aggregation of α-syn are a hallmark of PD. Hence, the first phenotype that we analyzed in the 3xSNCA MOs was α-syn protein expression and pathology. Here and for all following analysis we show the results for the two 3xSNCA lines and the four healthy control lines averaged. The SNCA-KO line is used to verify anti-α-syn antibody specificity.

During development, α-syn translocates from the cytosolic compartment in immature cells to a synaptic location in mature polarized neurons (Chandra et al., 2004). MOs recapitulated this phenomenon. In 15- day-old MOs α-syn showed a diffuse cytoplasmic distribution, which became confined in a dot like pattern from 30 days of differentiation onwards (Fig. S7A, B).

Via immunofluorescence staining we were able to detect α-syn and pS129 α-syn (α-syn with a phosphorylation in the Serine residue 129) in MOs (Fig. 1A). Co-localization of both α-syn antibodies via immunofluorescence and lack of signal in SNCA-KO MOs supported the specificity of the α-syn detection (Fig. 1A). The immunostaining analysis revealed significantly higher levels of α-syn in 3xSNCA MOs in comparison to healthy controls at different time points of differentiation (*p < 0.05, ***p < 0.001) (Fig. 1B). This result was further confirmed via Western blotting (Fig. 1C, D) (**p < 0.01, ****p < 0.0001). Furthermore, increased extracellular α-syn in 3xSNCA MOs cultures was also observed via Elisa and dot blotting (Fig. 1E, F, G) (***p < 0.001, ****p < 0.0001).

**Figure 1:**
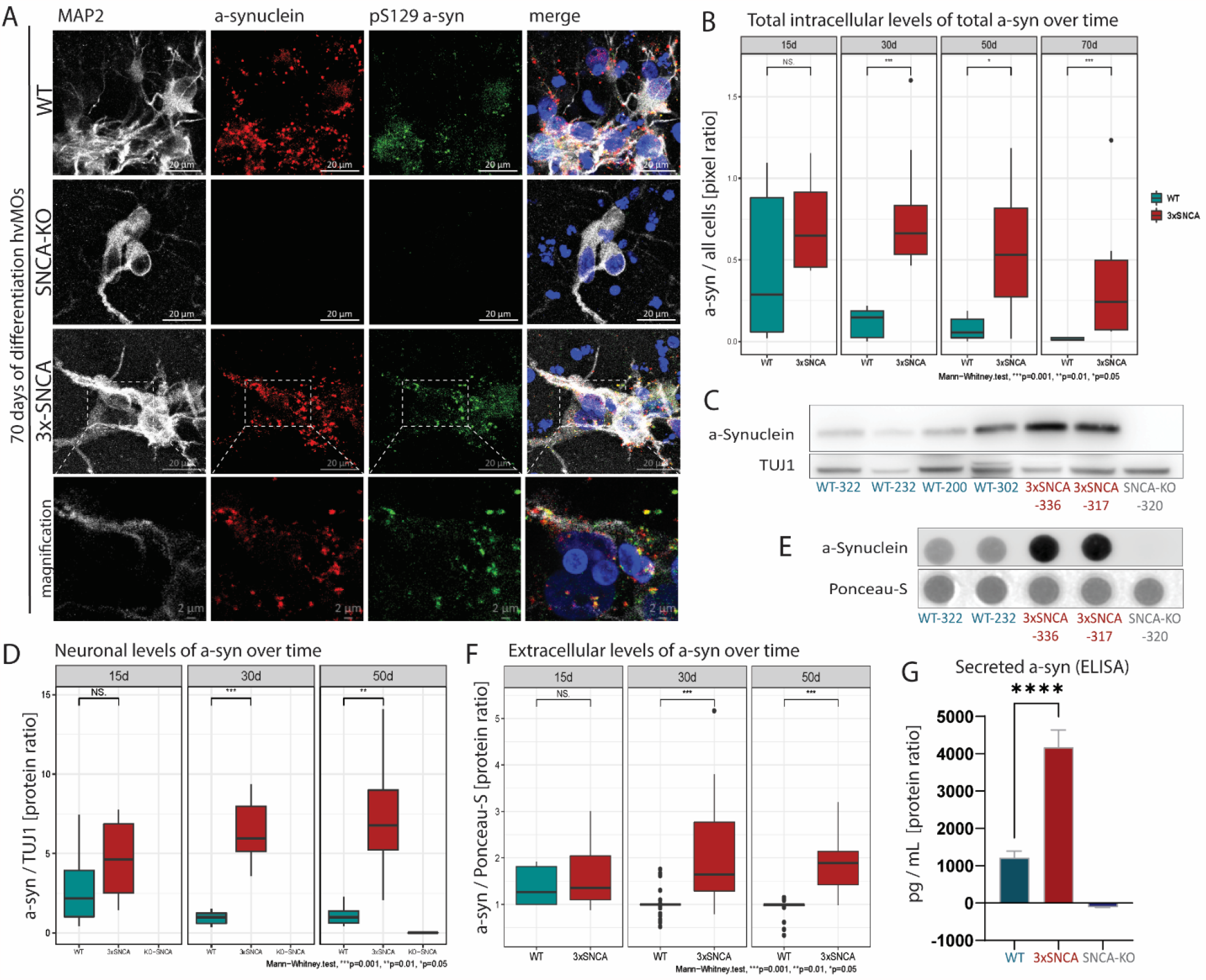
3xSNCA MOs show increased extra-and intra-cellular total α-syn. (A) Representative confocal images (60X) for total α-syn and pS129 α-syn from 70-day-old WT, 3xSNCA, and SNCA-KO MOs. (B) Inmunofluorescence quantification of intracellular total α-syn over time in WT and 3xSNCA hMOS in all cells. (C) Representative Western blot membranes for total α-syn and TUJ1 from 50-day-old MOs. (D) Western blot quantification for total α-syn and TUJ1 from 15, 30, and 50-day-old MOs (3 MOs pooled per line and batch, 3 independent batches. n=3 per line, n>=6 per condition (condition = WT, 3xSNCA, SNCA-KO)). (E) Representative dot blot membranes for total α-syn and Ponceau-S from 50-day-old MOs. (F) Dot blot quantification for α-syn in media normalized to Ponceau-S from 15, 30, and 50-day-old MOs (media pooled from 3 MOs per line and batch of MOs (>=6 independent batches), n=16 per condition, n=2 for SNCA-KO). (G) ELISA quantification for total α-syn in media from 50 days-old MOs (3 MOs pooled per line and batch (n=6 independent batches). (B, D-G) Statistical significance by Mann–Whitney U test *p < 0.05, **p < 0.01 and ***p < 0.001.

Since pS129 α-syn is associated with aggregation and consequent toxicity (Oueslati, 2016), we further investigate its presence in the MOs. Firstly, we tested the specificity of the antibody towards the phosphorylation via the usage of lambda phosphatase. Phosphatase treated MOs showed a significant reduction of pS129 α-syn signal compared to untreated MOs (Fig. 2A, B) (**p < 0.01).

**Figure 2:**
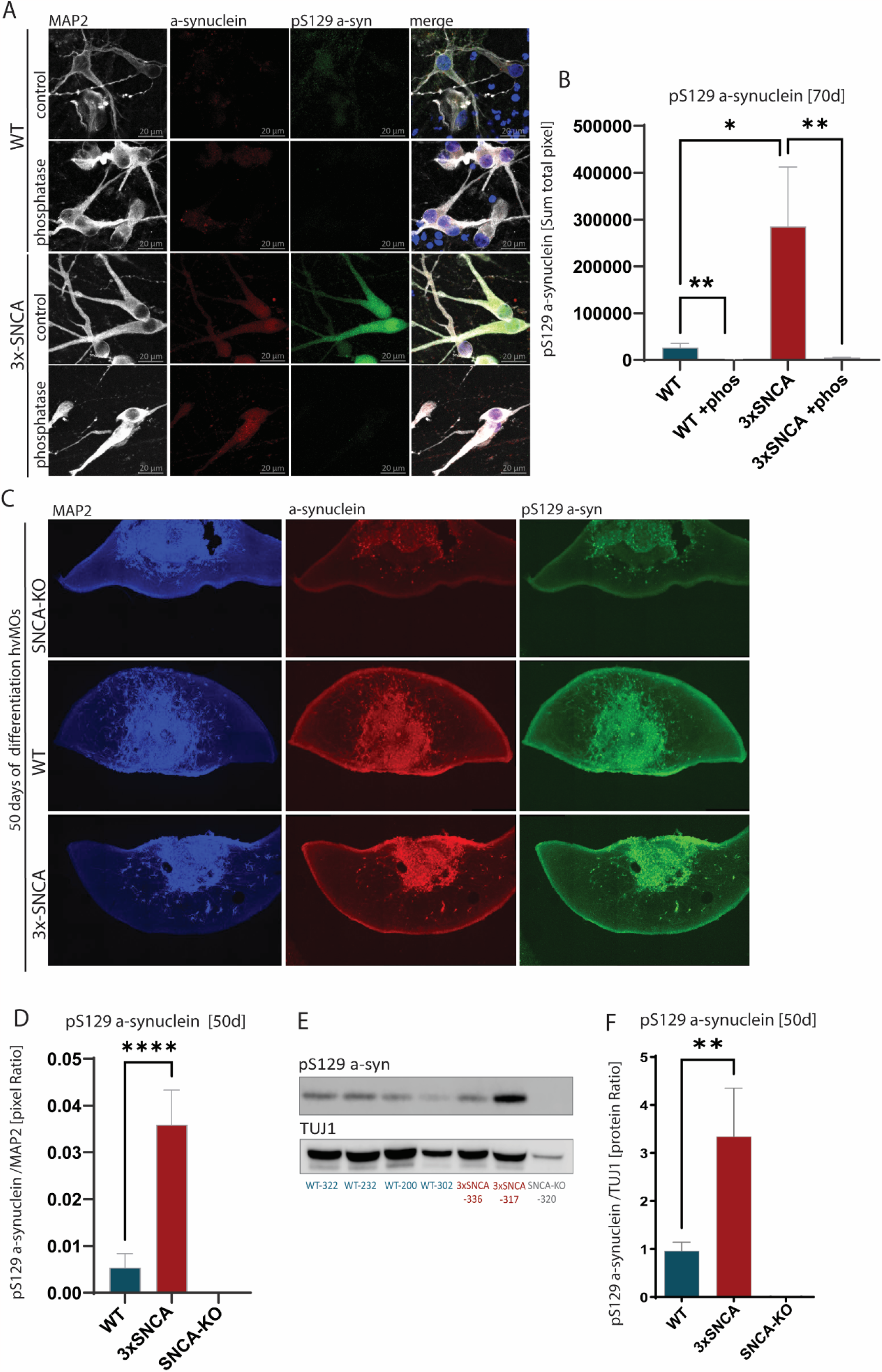
3xSNCA MOs show increased levels of phosphorylated (pS129) α-syn. (A) Representative confocal images (60X) for total α-syn and pS129 α-syn from WT and 3xSNCA MOs, untreated and treated with Lambda phosphatase. (B) Inmunofluorescence quantification of intracellular pS129 α-syn of 70-day-old MOs in WT and 3xSNCA hMOS, untreated and treated with Lambda phosphatase. (C) Representative immunofluorescence images from a confocal microscope, 20X high numerical aperture, scale bar 200um (D) Quantification of pS129 α-syn from 50-day-old SNCA-KO, 3xSNCA, and WT MOs corresponding to figure C. (E) Representative western blot membrane for pS129 α-syn and TUJ1 from 50-day-old MOs. (F) Western blot quantification for pS129 α-syn and TUJ1 from 50-day-old MOs (n=3 per line, n=6 per condition). (B, D, F) Statistical significance by Mann–Whitney U test *p < 0.05, **p < 0.01 and ***p < 0.001.

The immunostaining analysis revealed increased pS129 α-syn levels in 3xSNCA MOs (*p < 0.05, ****p < 0.0001) (Fig. 2B, C, D, S8A). These increased pS129 α-syn levels in 3xSNCA MOs were further confirmed via Western blotting (*p < 0.05) (Fig. 2E, F).

In summary, the patient specific MOs recapitulate higher levels and phosphorylation of α-syn as seen during pathology in patients’ brains.

### 3.3 3xSNCA MOs show progressive dopaminergic neuron loss

The second hallmark of PD, alongside with α-syn aggregation, is the loss of dopaminergic neurons during the progression of the disease. To analyze whether patient specific MOs recapitulate this process of neurodegeneration, we analyzed the number of the dopaminergic neuron population over time.

High content imaging was used to analyze the general neuronal population (TUJ1 positive) as well as the dopaminergic neuron population (TH positive) (Fig. 3A). Quantification of the amount of dopaminergic neurons normalized to the total neuronal population showed an increased amount of dopaminergic neurons at early time points of the differentiation process in 3xSNCA MOs (15 day-old MOs) (Fig 3B). Importantly, overtime the amount of dopaminergic neurons in the patient specific MOs progressively declined under the level seen in healthy control organoids (90 day-old MOs) (Fig. 3B) (*p < 0.05, ***p < 0.001). Hence, the patient specific MOs mirror the characteristic progressive decline described in PD pathology.

**Figure 3:**
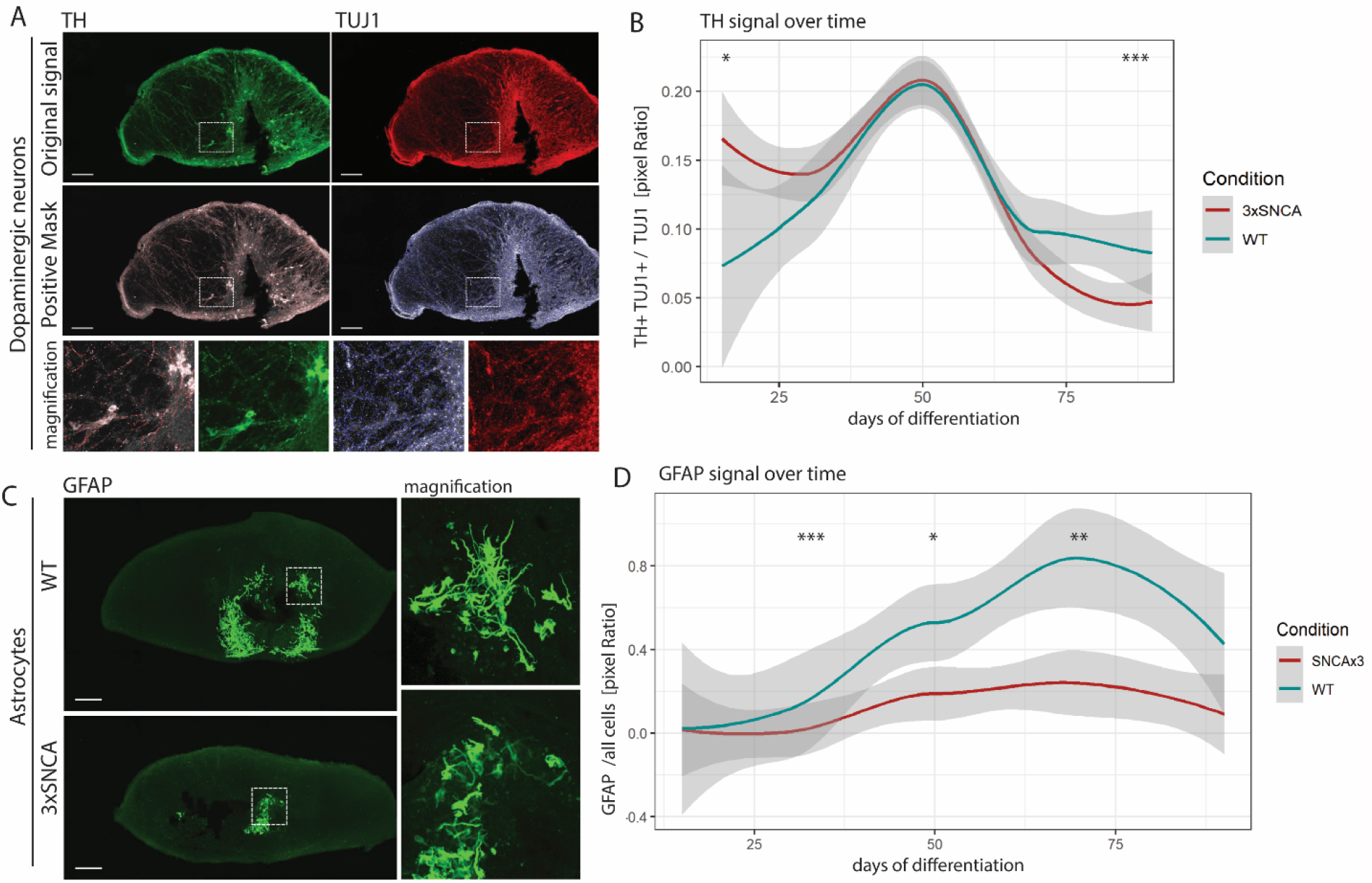
3xSNCA MOs present dopaminergic neuronal loss and deficient astrogenesis. (A) Representative images were obtained with the High Content Operetta microscope 20x for TH and TUJ1 signal, and masks were generated for the positive signal. (B) Time-line representation of the TH+ and TUJ1+ population (double-positive) and normalized to total TUJ1+ signal based on images from A. (C) Representative images obtained with the High Content Operetta microscope 20x for GFAP. (D) Time-line representation of the GFAP+ population and normalized to total population (Hoechst signal excluding pyknotic nuclei) based on images from C and mask from (Fig. S8C). (B, D) Statistical significance by Mann–Whitney U test *p < 0.05, **p < 0.01 and ***p < 0.001 (n>=3. Sections from n>=3 independent organoids per generation. Total of 3-6 independent MOs generation).

In addition, since astrocytes have gained substantial importance in PD research and MOs allow to study other cellular populations from the CNS, we investigated the astrocytic population in the patient specific MOs. High content imaging analysis of astrocytes, being positive for the Glial fibrillary acidic protein (GFAP), revealed a delayed and reduced astrocytic differentiation in the 3xSNCA MOs (30, 50 & 70 day-old MOs) (Fig. 3C, D, S8C) (*p < 0.05, **p < 0.01, ***p < 0.001).

Altogether, our data suggest that patient specific MOs represent a robust model that recapitulates key molecular features of PD and might shed light into the mechanism of neurodegeneration in 3xSNCA PD.

### 3.4 Gene expression is largely altered in patient specific MOs

To detect possible causes for the observed phenotypes, we performed a whole-transcriptome analysis via RNA-sequencing of 30-day-old MOs.

Hierarchical clusters clearly separated healthy control (wild type, WT) and patient specific midbrain organoid (3xSNCA) populations (n=3) (Fig. 4A). A total of 3200 genes were found significantly dysregulated (FDR < 0.05) including the PD-associated genes *NURR1* (nuclear receptor related 1 protein), *PODXL* (Podocalyxin Like), and *CHCHD2* (Coiled-Coil-Helix-Coiled-Coil-Helix Domain Containing 2); and the *SNCA*-associated genes *MAOB (*monoamine Oxidase B), *KLK6* (Kallikrein Related Peptidase 6), and *PPP2R5D* (Protein Phosphatase 2 Regulatory Subunit B’Delta) (FDR < 0.05) (Table S3A, B, C).

**Figure 4:**
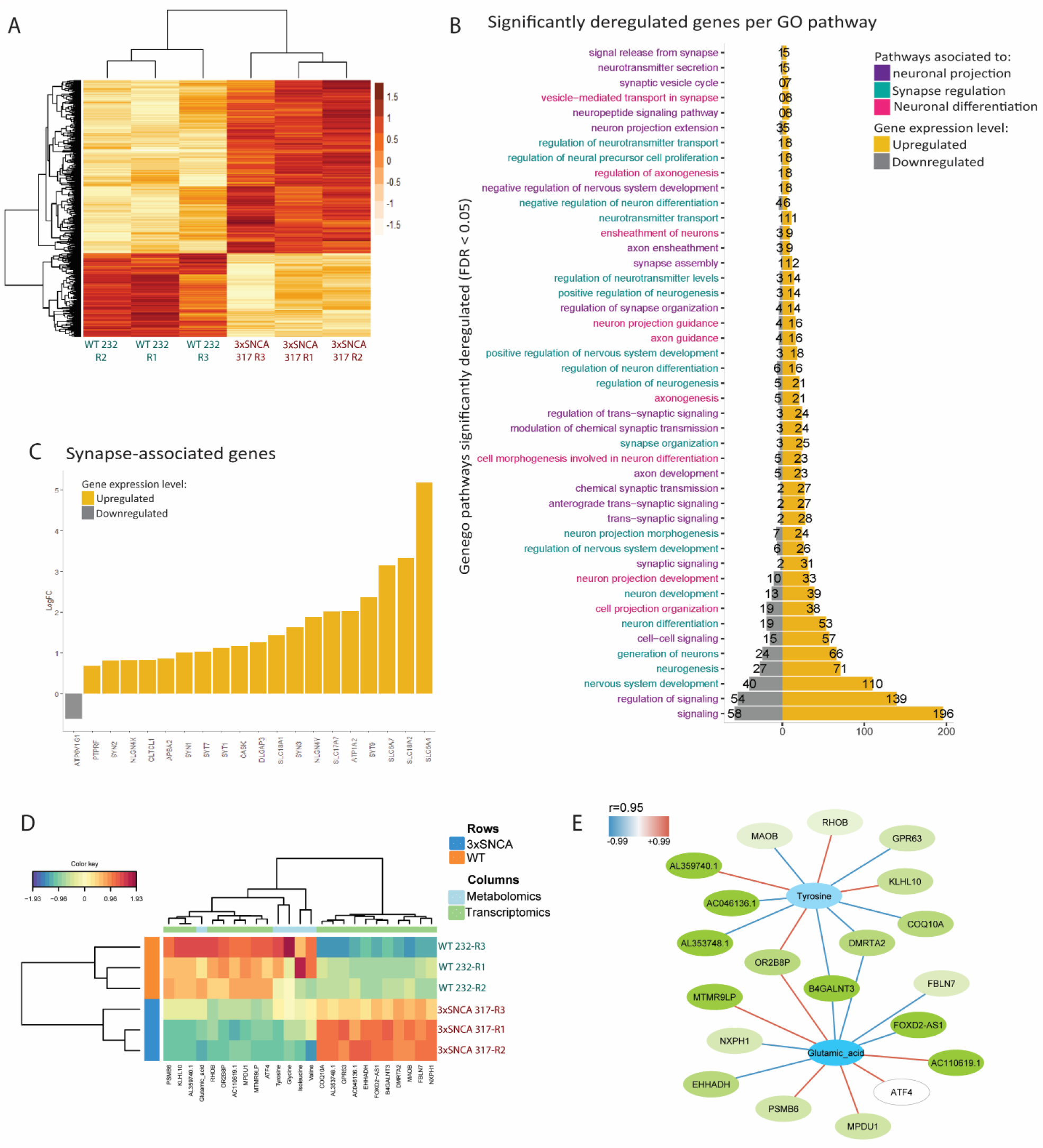
Transcriptomic data reveals dysregulation of neuronal and synapse-associated pathways in 3xSNCA MOs. (A) Hierarchical cluster of z-score values for dysregulated genes in 30-day-old 3xSNCA and WT MOs (FDR < 0.05) (n=3, 9 pooled MOs from 3 independent batches). (B) Selected dysregulated GO processes network in 3xSNCA MOs. Representation of the number of significantly deregulated genes (FDR < 0.05) for each pathway. (C) Bar plot representing synapse associated genes logFC (FDR < 0.05). Synapse-associated genes were obtained from PathCards database (https://pathcards.genecards.org). (D) Clustered Image Map representing the multiomics signature in relation with the samples using principal component 1 from the discriminative PCA. (E) Metabolite-gene association network abs(r) >= 0.95, Positive correlations in red, negative correlations in blue. Transparency of nodes depicts importance of each variable in PCA1.

In addition, gene ontology (GO) pathway analysis revealed 601 processes significantly dysregulated between WT and 3xSNCA MOs (FDR < 0.05) (Table. S3D). Interestingly, when looking at the gene expression within pathways of interest, we could observe that most genes associated with neuronal differentiation, axonal projection, and synapse showed increased expression in 3xSNCA MOs (Fig. 4B). These results are in accordance with the early dopaminergic neuron specification that we described here (Fig. 3B) and further supported by increased expression of most genes associated with tyrosine metabolism, dopamine metabolism, and dopamine neurotransmitter such as *DDC*, monoamine Oxidase A (*MAOA*), and *MAOB* (Fig. S9A, Table. S3E).

A-syn is a presynaptic protein previously described to influence synapses (Burré, 2015). Hence, the increased expression of most genes associated with the synapse and the synaptic vesicle cycle might be caused by high α-syn expression or high α-syn intracellular levels (Fig. 4C). Accordingly, we also observed upregulation of genes associated with neurotransmitter release (Table. S3F).

Interestingly, the transcriptomic analysis revealed also decreased expression of genes associated with astrocytic differentiation such as *GFAP, S100b* (S100 Calcium Binding Protein B), *AGER* (Advanced Glycosylation End-Product Specific Receptor), *TSPAN2* (Tetraspanin 2), *VIM* (Vimentin), *GPR183* (G Protein-Coupled Receptor 183), and *NKX2-2* (NK2 Homeobox 2) (FDR < 0.05). This is consistent with the reduced amount of astrocytes observed in the immunofluorescence staining based analysis (Fig. 3D).

In addition, targeted metabolomic analysis of the spent culture medium from 30-day-old healthy and 3xSNCA MOs was performed. A significant increase in lactate levels as well as a decrease in glutamate levels was observed in 3xSNCA MOs (Fig S9B, C) – both considered biomarkers for CNS related diseases (Donatti et al., 2020). In PD patients’ blood lactate levels are increased (Nakagawa-Hattori et al., 1992), while glutamate levels are decreased in the brain (O’Gorman Tuura et al., 2018).

Moreover, we use Data Integration Analysis for Biomarker discovery using Latent cOmponents (DIABLO) of combined metabolomic and transcriptomic data to separate healthy from PD patient MOs. The separation between conditions was obtained using the correlation of the top 20 genes with the top 5 metabolites from the discriminative PCA (component 1) (Fig. S9D) (Singh et al., 2018). Heatmap visualization lists the genetic and metabolic expression profile responsible for cluster separation between conditions (Fig. 4D). Contribution plots rank the importance of each gene or metabolite for the separation of both conditions (Fig. S9E, F). Pearson correlations (abs(r) >= 0.7) between genes and metabolites as well as their expression levels are further visualized in a circus plot (Fig. S9G). From these correlations the top ones (abs(r) >= 0.95) are represented within a network (Fig. 4E). Interestingly, the amino acid tyrosine and the gene MAOB are negatively correlated. Their combined deregulation goes in accordance with the aforementioned dopaminergic alterations observed in 3xSNCA MOs, and matches the possible neurotoxic role of increased MAOB previously described in PD patients (Tong et al., 2017).

### 3.4 Synapse number and electrophysiological activity is progressively reduced in patient specific MOs

Synaptic dysfunction has been previously linked to PD (Bridi and Hirth, 2018; Ghiglieri et al., 2018). However, whether it is a cause or a consequence of neurodegeneration is still under debate. Since the transcriptomic results point to an increased expression of synapse related genes in 30-day-old 3xSNCA MOs (Fig. 4C), we performed Microelectrode array (MEA) measurements to investigate synaptic activity (Fig. 5A, B, C, D). Electrophysiological activity of MOs was recorded from 15 to 90 days of differentiation.

**Figure 5:**
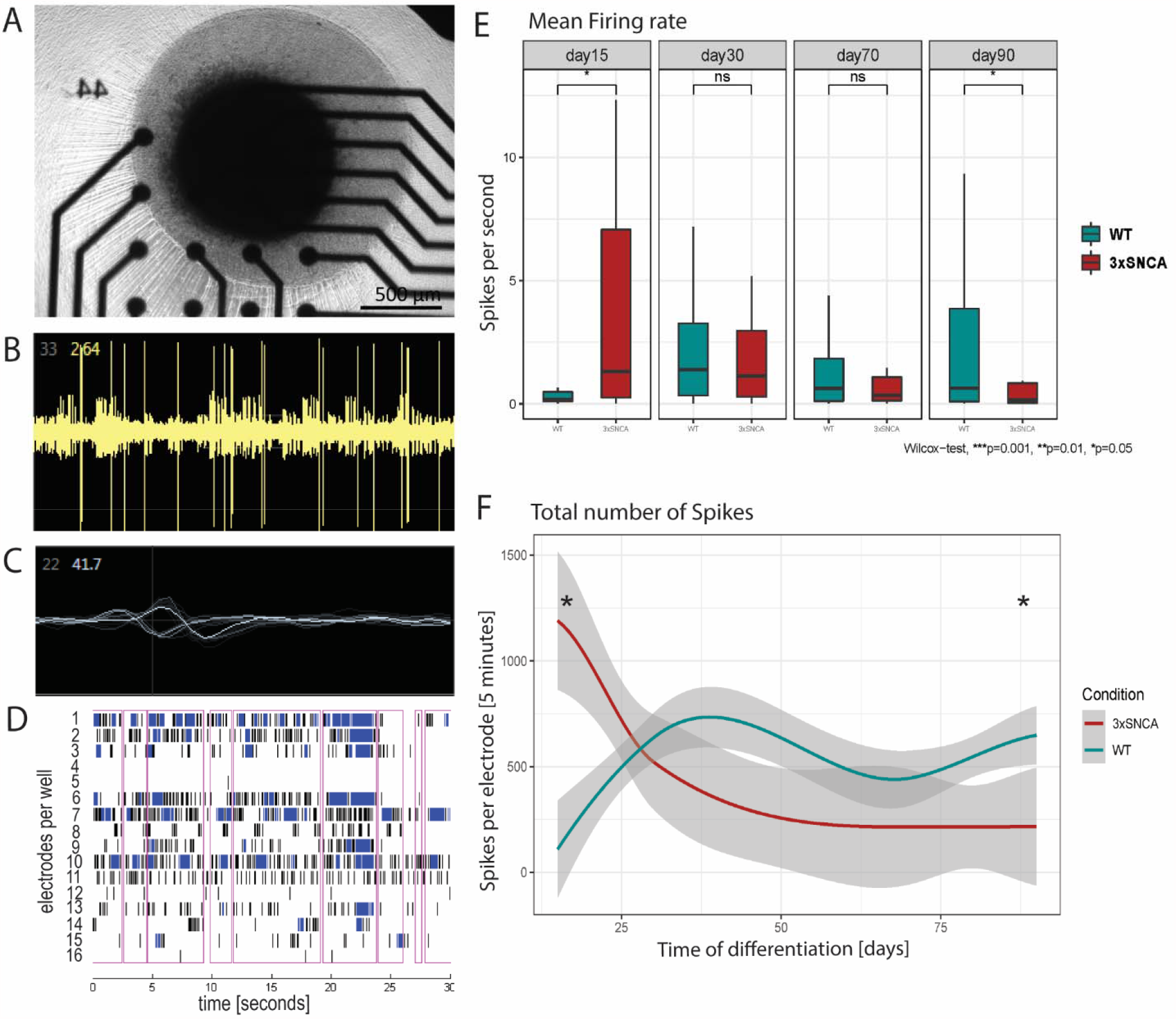
3xSNCA MOs present a decrease in electrophysiological activity over time. (A) Representative image of a MO in a MEA well consisting of 16 electrodes. (B) Representative image of the spike recording from an electrode. (C) Representative image of the spike waves from an electrode. (D) Representative image of the network burst from a MEA well. Each row represents an electrode within the well. Each spike is represented with a vertical line in black, each burst in blue, and connectivity between electrodes is marked by a pink rectangle. (E) Box-plot representation of the mean firing rate (spikes per second). (E) Time-line representation of the total number of spikes per electrode (threshold >5 Spikes/minute/electrode). (E, F) Statistical significance by Mann–Whitney U test *p < 0.05.

Patient specific MOs with the 3xSNCA genotype showed an increase in the number of spikes, mean firing rate, inter-spike interval and bursting events in comparison to healthy controls at early stages (15-day-old MOs). However, these values progressively decline below the values for the healthy control MOs at later time points (Fig. 5E, F, S10A, B) (*p < 0.05).

In order to delineate the molecular reason for the progressive decline in the electrophysiological activity, we analyzed the amount of presynaptic (positive for Synaptophysin) and postsynaptic (positive for PSD95) signals via immunofluorescence (Fig. 6A).

**Figure 6:**
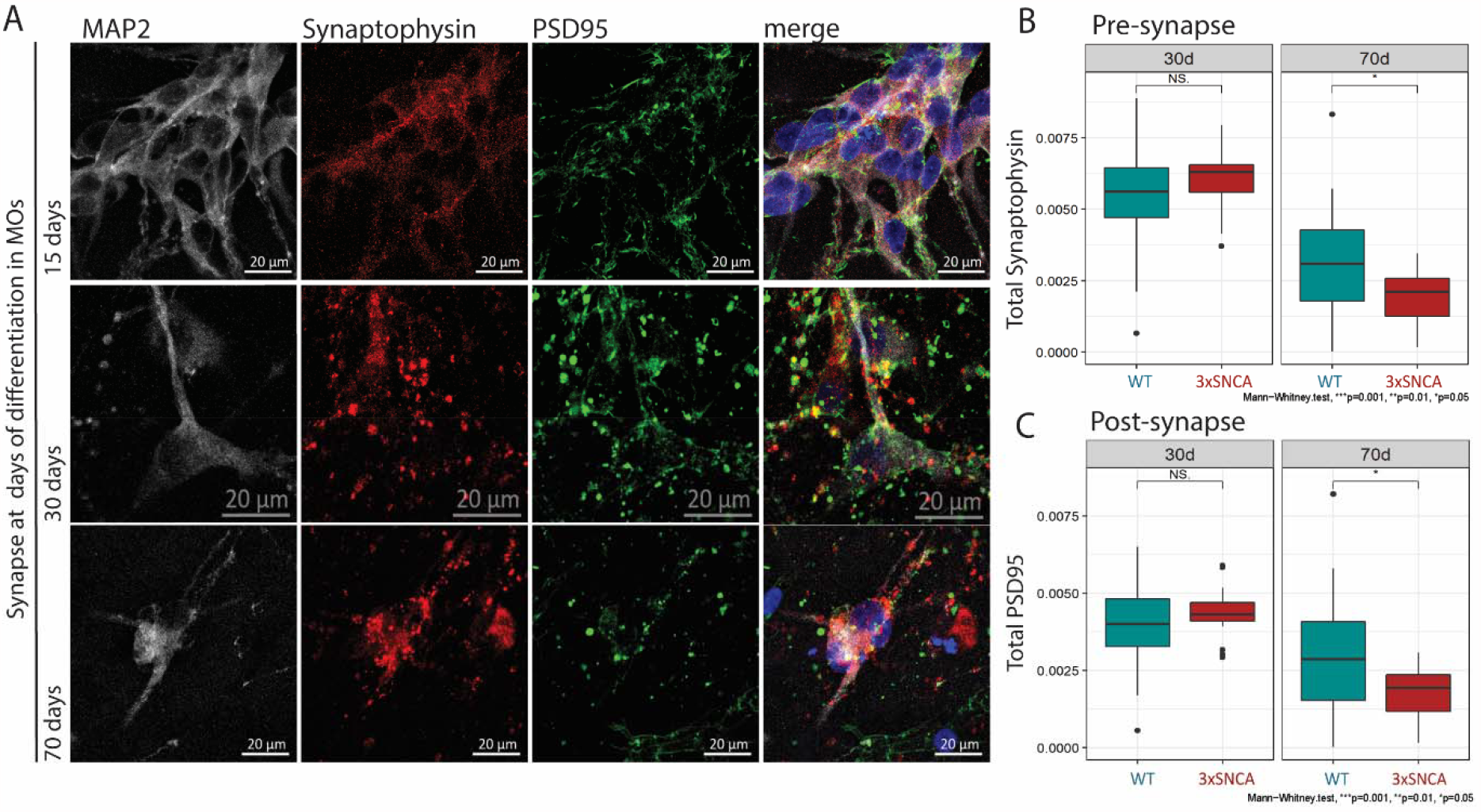
3xSNCA MOs present reduced synapse. (A) Representative confocal images (60X) for the neuronal marker MAP2, the presynaptic marker synaptophysin, and the postsynaptic marker PSD-95. Images from 15, 30, and 70-day-old MOs. (B) Box-plot synaptophysin counts in the synapse (pre-/post-synaptic marker in proximity) from 30 and 70-day-old MOs. (C) Box-plot of PSD-95 counts in the synapse (pre-/post-synaptic marker in proximity) from 30 and 70-day-old MOs. (B, C) (n=9 per line, n=18 per condition, 3 biological replicates, 3 area scan, 10 Z-planes, 0.46 um per sample). (B, C) Statistical significance by Mann–Whitney U test *p < 0.05.

Synaptic markers showed a diffuse cytoplasmic distribution in 15-day-old MOs, which became confined in a dot-like pattern from 30 days of differentiation onwards (Fig. 6A). Furthermore, immunofluorescence analysis indicated colocalization of α-syn and the presynaptic markers synaptophysin (Fig. S10C). Synaptic events were only considered when both pre-/post-synaptic marker were in proximity (Fig. S10D). Interestingly, although no differences in synapse number were encountered at early time points of differentiation, 70-day-old 3xSNCA MOs showed a significant reduction in synapse count compared to WT MOs (*p < 0.05) (Fig. 6B, C). These results indicate that loss of synapses in the patient specific organoids accompanied with reduced electrophysiological activity, precedes degeneration of dopaminergic neurons.

## 4. Discussion

Loss of dopaminergic neurons and α-syn aggregation are main hallmarks of PD. Nevertheless, there is still no clear association between α-syn, neurogenesis and neurodegeneration. In addition, the lack of translatable models has result in contradictory data in the field. Here, we show a robust and reproducible model which recapitulates major PD molecular hallmarks as well as human midbrain cellular complexity and structure. MOs harboring a triplication in the *SNCA* gene show a 2-fold increase of total and pS129 α-syn and mimic α-syn synaptic location. Using this model we aimed to contribute to a more detailed understanding of the role of α-syn in PD. For this purpose, we assessed the 3xSNCA MOs cellular phenotype followed by an investigation of the possible causes behind it.

The here described 3xSNCA MOs mimic the loss of dopaminergic neurons observed in PD. Interestingly, this progressive loss of dopaminergic neurons is preceded by an overproduction of the dopaminergic neuron population at earlier stages of differentiation. Consistently, 3xSNCA MOs also presented an increased expression of genes associated to neurogenesis and the dopaminergic system, supporting the aforementioned phenotype. The positive effect of α-syn on dopaminergic neuron differentiation is in accordance with previous reports (Garcia-Reitboeck et al., 2013; Tani et al., 2010). Although lower levels of dopaminergic neuron differentiation have also been linked previously to *SNCA* triplication models (Oliveira et al., 2015), these results were based on 2D neuronal cultures and a single time point of differentiation. The here presented developmental model allow the study of α-syn role overtime. Our observations raise the intriguing question whether also in patients’ brain the amount of dopaminergic neurons is initially higher than in healthy individuals. This hypothesis would be in line with a recent publication that actually implied a toxic function of excess levels of dopamine, leading to posterior neurodegeneration (Sackner-Bernstein, 2021). In fact, the here presented combinatorial analysis of metabolomic and transcriptomic data from 3xSNCA MOs indicated a correlation between tyrosine metabolism and the MAOB gene. Increased MAOB expression in 3xSNCA MOs matches with the previously described toxic effects of the dopamine metabolism in PD (Alborghetti and Nicoletti, 2018).

Additionally, the findings presented here support a role of the presynaptic protein α-syn in early synapse development. According to the literature, α-syn contributes to changes in synaptic protein levels, synapse structure, vesicle turnover, and regulation of dopamine transmission in dopaminergic neurons (Bellani et al., 2010; Bendor et al., 2013; Burré, 2015; Burré et al., 2014, 2010; Mosharov et al., 2010). The functionality of α-syn, its co-localization with the presynaptic marker synaptophysin, and the parallelism of both proteins in the confinement to a dotted pattern support α-syn synaptic functionality in the MO system used here. Early in the differentiation process (15-day-old), 3xSNCA MOs showed higher electrophysiological activity than healthy MOs. In line with this result, we observed an increased expression of synapse-associated genes in patient specific MOs. However, mature 3xSNCA MOs (90-day-old) showed a decline in electrophysiological activity below healthy MOs values. Accordingly, synaptic events were significantly reduced at later time points of differentiation in 3xSNCA MOs, likely causing the observed decreased in electrophysiological activity. These results further support the previously described synaptic decline in PD (Ghiglieri et al., 2018). We speculate that potentially, an early overactivation, caused by higher α-syn levels might explain a later decline (Bridi and Hirth, 2018).

Interestingly, patient-derived MOs showed a ∼50% reduction in astrocyte number compared to healthy MOs. This result matches with the decreased expression of astrocyte-specific genes in 3xSNCA MOs and may suggest that the early increased neuronal differentiation is at expense of astrogenesis. Indeed, previous literature describe a negative effect of α-syn on the maturation capacity of glial cells (Ettle et al., 2014). Furthermore, astrocytes are necessary for neuronal homeostasis and synaptic functionality (Allen and Eroglu, 2017; Araque et al., 2014; Bélanger and Magistretti, 2009). In fact, the incorporation of astrocytes to neuronal culture increased the number and strength of synapse (Pfrieger and Barres, 1997; Ullian et al., 2001). Therefore, the reduced astrocytic population might additionally contribute to the synaptic decline and neurodegeneration observed in 3xSNCA MOs.

Parkinson’s disease is generally associated with toxic forms of α-syn. However, the results presented here suggest a higher complexity, thus pointing to a role of α-syn in the composition and functionality of the developing midbrain. On the one hand, the pre-synaptic protein α-syn enhances early dopaminergic differentiation (Garcia-Reitboeck et al., 2013). On the other hand, increased dopamine metabolism caused by upregulation of MAOB leads to oxidative stress contributing to toxicity and neurodegeneration as previously described for PD (Nagatsu and Sawada, 2006). Accumulation and/or aggregation of toxic α-syn forms also lead to synaptic decline and neurodegeneration (Bridi and Hirth, 2018). Here, we observed that increased levels of α-syn lead to increased early synaptic activity followed by synaptic decline and neurodegeneration, which might be exacerbated by the lack of neuronal supporter cells such as astrocytes.

Altogether, our results support a key involvement of α-syn in the dopaminergic system, synaptic activity and astrogenesis, revealing some of its more complex and multifaceted roles in the physiological and pathological midbrain. Importantly, here we show that synaptic decline precedes neurodegeneration. Furthermore, this study underpins that patient specific midbrain organoids enable a detailed personalized phenotyping in PD, since MOs endogenously recapitulate the key molecular hallmarks on PD with unprecedented fidelity. Patient specific MOs may therefore serve as a new versatile tool for personalized medicine and drug discovery.

## 5. Material and methods

### iPSCs, mfNPCs and organoid culture

Induced pluripotent stem cells were obtained from (Reinhardt et al., 2013), purchased from (https://nindsgenetics.org/), acquired from (https://www.coriell.org, cell no. ND27760) and genetically modified (Barbuti et al. *in preparation*). More information is detailed in (Table. S1). iPSCs were culture in Essential 8 medium (Thermo Fisher, cat no. A1517001) with 1% Penicillin/Streptomycin (Invitrogen, cat no. 15140122) in Matrigel-coated plates (Corning, cat no. 354277). Splitting was perform using Accutase (Sigma, cat no. A6964) and including ROCK inhibitor Y-27632 (Merck Milipore, cat no. 688000) at 10uM for 6-24□h after seeding.

Fluorescence in situ hybridization analysis was performed by Cell Line Genetics (https://clgenetics.com/) in mfNPCs cells containing a triplication of the *SNCA* gene (lines DB317 and DB336). Midbrain floor plate neural progenitor cells were derived from iPSCs as previously described by us (Smits et al., 2019). mfNPCs were maintained in 2D condition and culture in freshly supplemented N2B27 as described in (Nickels et al., 2020). To ensure a pure population of mfNPCs, the cells were passed at least 9 times before hvMO generation.

Midbrain organoids were generated as detailed in our previous study from (Nickels et al., 2020), using a reduced seeding density of 6000 cells. MOs were embedded in geltrex (Invitrogen, cat no. A1413302) at day 8 of differentiation and kept at 37C, 5% CO2 under dynamic conditions (80 rpm) for up to 180 days (Nickels et al., 2020).

### Immunocytochemistry

Immunofluorsecent stainings for iPSCs and mfNPCs were perform as described in (Gomez-Giro et al., 2019) and (Smits et al., 2019), respectively.

MOs were collected at several time points (15d, 30d, 50d, 70d, 90d, 110d, 130d, 150d, 180d, 200d). MOs were fix in 4% PFA (overnight at 4°C), washed in PBS1x and sectioned into 70 μm thick slices (Nickels et al., 2020). Immunostaining of MOs were performed as indicated in (Nickels et al., 2020) with minor modifications. Briefly, sections were blocked and permeabilized in 2% BSA, 5% Donkey serum and 0.5% Triton X-100 for 120 minutes. Primary antibodies were diluted in 2% BSA, 5% Donkey serum and0.1% Triton X-100 for 72h at 4 °C in dynamic conditions. Secondary antibodies incubation and mounting was then performed as in (Nickels et al., 2020). The antibodies used in this manuscript are summarized in (Table. S5).

#### Lambda phosphatase treatment

Prefixed (4% PFA) MOs sections (70um) were treated with 400 Units of lambda phosphatase reaction buffer (Santa Cruz, cat no. sc-200312A) for 30 minutes at 37C. Immunostaining was performed as usual. Lambda phosphatase reaction buffer was prepared by mixing, 400 unites of lambda phosphatase in 1X lambda phosphatase buffer, and 1X MnCl2 2 mM solution.

### Image acquisition

Bright field images were taken using the AxioVert.A1 from Zeiss. Confocal images for iPSCs, mfNPCs and MOs were taken with Microscope ZEISS confocal LSM 710. For synapse and dotted α-syn images, 12 z-planes with separation of 0.46um were acquired with a 60X oil objective. For overview of MOs 20X objective was used with tile-scan and online stitching. Operetta CLS High-content Imaging System (Perkin-Elmer) was used for acquisition and quantification of MOs sections. 20X objective with high numerical aperture was used to acquired high resolution images of the entire MOs, using a 15% overlapping between regions and 30-35 planes per section. The distance between sections was the recommended by the manufacture for the specified objective (0.8 um).

### Image quantification and data analysis

Confocal image (Microscope ZEISS confocal LSM 710) analysis was done in MatLab (Version 2019a, Mathworks). Script for data analysis is included in the corresponding Gitlab repository. MOs section image analysis (Operetta CLS High-content Imaging System) was done with Matlab (Version 2017b, Mathworks) using in-house developed image analysis algorithm for automated marker quantification (Bolognin et al., 2019; Monzel et al., 2019, 2017; Nickels et al., 2020; Smits et al., 2019). Script for data analysis is included in the corresponding Gitlab repository. Further data analysis and representation was done with R (R-3.6.3) and R studio (Version 1.2.5033). HMOs area were automatically determined using Microplate Cytation5M Cell imaging Multi Mode Reader (Biotek).

### Western Blotting

Pool of 2-3 MOs were lysed in RIPA lysis buffer (Abcam, cat no. ab156034) containing 1X protease inhibitor cocktail (PIC)(Sigma). Samples were sonicated in Bioruptor (Diagenode) for 10 cycles 15 sec ON 15 sec OFF. Protein samples were resolved by precast polyacrylamide gels (Thermofisher, cat no. NW04120BOX). Proteins were transferred from the gel to PVDF membranes in an iBlot2 device (Thermo Fisher, cat no. IB24001). Depending on the protein, membranes were fix in 0.04% PFA for 30 minutes at room temperature (RT). Further washed 5 minutes in PBS. Membranes were blocked in 0.2% Tween-20, 5% Milk in 1xPBS for 60 min at RT. Primary antibodies were incubated in 0.02% Tween-20, 5% Milk in 1xPBS at 4°C overnight. Secondary antibodies were incubated for 60 min at RT in the same buffer as the one described for primary antibodies. Membranes were revealed using the SuperSignal West Pico Chemiluminescent Substrate (Thermo, cat no. #34580). Enhanced chemiluminescent signal was detected in a STELLA imaging system. The antibodies used in this manuscript are summarized in (Table. S5).

For phosphorylated proteins RIPA lysis buffer was further supplemented with PIC and 1X phosphatase inhibitor cocktail (Merck Millipore, cat no. 524629-1ML). A cocktail of phosphatase inhibitors (B-Glycerolphosphat (25mM), NaF (5mM) and Na3VO4 (1mM)) was also included during the incubation of primary and secondary antibodies.

### Dot Blot

Media from 2-3 MOs was pooled, then snap frozen or directly used. Media was thaw on ice, and spinned down at 300g and 4°C for 5 minutes before usage. Nitrocellulose membrane (Sigma-Aldrich, cat no. GE10600001) was hydrated before usage in PBS. Dot Blot Minifold I (Whatman; 10447900) was used as manufacturer recommended using 300 µl of media, and 1:2 and 1:4 dilutions in PBS were loaded. Membrane was re-hydrated twice with 300ul of PBS per well before sample loading. After sample run (vacuum ON), the membrane was washed with 300ul of PBS per well. Membrane was retrieved and directly fix in 0.04% PFA. Afterwards, the membrane was washed in PBS 1 min for removal of PFA. Then, the membrane was incubated for 1 min in Ponceau S solution (Sigma-Aldrich, cat no. P7170-1L) and imaged. Ponceau signal was used for sample normalization. Blocking and antibody incubation were done as indicated in the western blotting area. Images were acquired with ODYSSEY LI-COR or STELLA imaging system. Images were analyzed with Image Studio Lite Ver 5.2. The antibodies used in this manuscript are summarized in (Table. S5).

### qPCR

RNeasy Mini Kit (Qiagen, cat no. 74106) was used following manufacturer instructions for total RNA isolation. RNase-Free DNase Set (Qiagen, cat no. 79254) was used for further DNase digestion. RNA-to-cDNA Kit (Invitrogen, cat no. 4387406) was used following manufacturer’s instructions to synthesize complementary DNA. GoTaq G2 Hot Start Green Master Mix (Promega, cat no. M7423) was used together with primers list in table 1 to perform reverse transcription PCR reaction. AriaMx Real-Time PCR system Aligent machine was used to perform qPCR and AriaMx PC software for the extraction and analysis of the data.

### Microelectrode array

The Maestro microelectrode array (MEA, Axion BioSystems) platform was used to record spontaneous activity of MOs along the maturation process. 48-well MEA plates consistent of 16 electrodes per well were precoated with 0.1-mg/ml poly-D-lysinehydrobromide (PDL SIGMA #P7886-50MG) overnight (ON) at 37C, 48 hours prior experiment. Carefully, PDL was removed and 10μg/ml laminin (SIGMA #L2020-1MG) was applied ON at 37C, 24 hours prior experiment. On the day of the experiment, 15-day-old MOs were carefully placed onto the array and covered by a droplet of geltrex (Invitrogen, cat no. A1413302). After 5 minutes of incubation at 37C (geltrex polymerization), neuronal maturation media was added. Organoids electrical activity was measured at several time points. 24 hours before each recording media was changed to supplemented BrainPhys^™^ Neuronal Medium and SM1 Kit (STEMCELL Technologies SARL, cat no. #05792). Axion Integrated Studio (AxIS 2.1) was used to perform analysis and data extraction. False positive and missed detections were minimized by using a Butterworth band pass filter with 200–3000 Hz cutoff frequency and a threshold of 6× SD. Spontaneous activity was recorded for 5 minutes at 37C and a sampling rate of 12.5 kHz. Electrodes were considered active when presenting an average of ≥ 5 spikes/min. Neural stat compiler files were used for the data analysis in R. Further information regarding The Maestro microelectrode array is included (Bardy et al.2015). Script for data analysis is included in the corresponding Gitlab repository.

### ELISA

Human Alpha-synuclein ELISA Kit (ab260052) was performed for the quantitative determination of alpha-synuclein secreted by 50-day-old MOs. Supernatant of 3 MOs per condition was pooled, snap frozen and store at −80C until needed. The ELISA was performed according to the manufacturer’s instructions with 50 μl of sample volume.

### Statistical analysis and graphical representation

Statistical analyses were performed in GraphPad Prism (Version 6.01) or R software (package: ggsignif). Significance asterisks represent *P*□<□0.05 *, *P*□<□0.01 **, *P*□<□0.001 ***, *P*□<□0.0001 ****; ns stands for not significant.

### Fluorescence in situ hybridization

FISH analysis was perform by Cell Line Genetics (https://clgenetics.com/) in mfNPCs cells containing a triplication of the *SNCA* gene (lines DB317 and DB336).

### Transcriptomics

Total RNA was isolated from 5 pooled organoids using Trizol reagent (cat. no. 15596026, ThermoFischer) according to the manufacturers protocol. 1 µg of total RNA was used for library preparation using TruSeq stranded mRNA library preparation kit (cat. no. 20020594, Illumina). Briefly, the mRNA pull-down was done using the magnetic beads with oligodT primer. To preserve the strand information the second strand synthesis was done such that during PCR amplification only first strand was amplified. The libraries were quantified using Qubit dsDNA HS assay kit (Thermofisher) and the size distribution was determined using Agilent 2100 Bioanalyzer. Pooled libraries were sequenced on NextSeq500 using manufacturer’s instructions.

### Transcriptomic data analysis

The raw RNA-seq fastq files were quality-checked using the FastQC software (“Babraham Institute Bioinformatics Group. FastQC, version 0.11.9,” 2010), and the data was pre-processed using Kallisto (version 0.46.0) (Bray et al., 2016) for alignment-free transcriptome quantification and human transcriptome annotations from the Ensembl 96 release. Gene-level differential expression analysis was conducted in R (version 3.5.1) (R Core Team, 2019) using the Bioconductor package edgeR (Robinson et al., 2009), and filtering out genes with low expression counts using the filterByExpr-function with default parameters. Normalization factors to scale the raw library size were determined using the calcNormFactors-function with default settings, and posterior dispersion estimates were obtained by applying the estimateDisp-function with the robust-parameter set to true in order to robustify the estimation against outliers.

Pathway enrichment analyses were conducted with the GeneGo MetaCore^™^ software (https://portal.genego.com) using the gene-level differential expression analysis results obtained with edgeR as input for the standard Enrichment Analysis workflow. The pathway over-representation analysis statistics, including false-discovery rate (FDR) scores according to the method by Benjamini and Hochberg (Benjamini and Hochberg, 1995), were determined for Gene Ontology biological processes and the GeneGo collections of cellular pathway maps and process networks.

Script for data analysis is included in the corresponding Gitlab repository.

### Polar metabolite extraction, derivatization, and GC-MS measurement

MOs were culture individually in 500ul culture media before collection. For each condition and biological replicate, media from 2-3 MOs was pooled, snap frozen and store at −80C until usage.

Extracellular metabolites from media samples were extracted using a methanolic extraction fluid (5:1, methanol/water mixture, v/v). The water fraction contained two internal standards Pentanedioic acid-D6 (*c* = 10 µg/mL; C/D/N Isotopes Inc.) and [UL-^13^C_5_]-Ribitol (*c* = 20 µg/mL; Omicron Biochemicals). 40 µL of medium was added to 240 µL ice-cold extraction fluid. After adding 100 µl ice-cold chloroform, the mixture was shaken for 5 min at 4 °C. For phase separation, 100 µl chloroform and 100 µl water were added and vortexed for 1 min. Then, the mixture was centrifuged at 21,000 xg for 5 min at 4 °C. 250 µL of the polar (upper) phase was transferred to GC glass vial with micro insert (5-250 µL) and evaporated to dry under vacuum at −4 °C.

Metabolite derivatization was performed by using a multi-purpose sample preparation robot (Gerstel). Dried medium extracts were dissolved in 30 µl pyridine, containing 20 mg/mL methoxyamine hydrochloride (Sigma-Aldrich), for 120 min at 45 °C under shaking. After adding 30 µl N-methyl-N-trimethylsilyl-trifluoroacetamide (Macherey-Nagel), samples were incubated for 30 min at 45 °C under continuous shaking.

GC-MS analysis was performed by using an Agilent 7890A GC coupled to an Agilent 5975C inert XL Mass Selective Detector (Agilent Technologies). A sample volume of 1 µl was injected into a Split/Splitless inlet, operating in split mode (10:1) at 270 °C. The gas chromatograph was equipped with a 30 m (I.D. 0.25 mm, film 0.25 µm) DB-5ms capillary column (Agilent J&W GC Column) with 5 m guard column in front of the analytical column. Helium was used as carrier gas with a constant flow rate of 1.2 ml/min. The GC oven temperature was held at 90 °C for 1 min and increased to 220 °C at 10 °C/min. Then, the temperature was increased to 280 °C at 20 °C/min followed by 5 min post run time at 325 °C. The total run time was 22 min. The transfer line temperature was set to 280 °C. The MSD was operating under electron ionization at 70 eV. The MS source was held at 230 °C and the quadrupole at 150 °C. Mass spectra were acquired in selected ion monitoring (SIM) mode for precise semi-quantification of medium components. The following masses were used for quantification and qualification of the derivatized target analytes (dwell times between 25 and 70 ms):

| Analyte Name | Quantification Ion (m/z) | Qualification Ion I (m/z) | Qualification Ion II (m/z) |
| --- | --- | --- | --- |
| Pyruvic acid 1MEOX 1TMS | 174 | 158.1 | 189.1 |
| Lactic acid 2TMS | 219.1 | 190.1 | 117.1 |
| Alanine 2TMS | 218.1 | 190.1 | 116.1 |
| Valine 2TMS | 144.1 | 218.1 | 246.2 |
| Leucine 2TMS | 158.1 | 218.1 | 232.2 |
| Isoleucine 2TMS | 158.1 | 218.1 | 232.2 |
| Glycine 3TMS | 276.1 | 248.1 | 174 |
| Serine 3TMS | 306.1 | 218.1 | 204.1 |
| Threonine 3TMS | 218.1 | 291.2 | 320.2 |
| IS Pentanedioic acid-D6 2TMS | 267.1 | 163.1 | 239.1 |
| Methionine 2TMS | 176.1 | 250.1 | 293.1 |
| Glutamic acid 3TMS | 363.2 | 246.1 | 348.2 |
| Phenylalanine 2TMS | 192.1 | 218.1 | 266.1 |
| Asparagine 3TMS | 231.1 | 132.1 | 348.1 |
| IS [UL-13C5]-Ribitol 5TMS | 220.1 | 310.2 | 323.2 |
| Glutamine 3TMS | 347.2 | 245.1 | 156.1 |
| Glucose 1MEOX 5TMS | 319.2 | 217.1 | 220.1 |
| Lysine 4TMS | 174.1 | 317.2 | 434.3 |
| Tyrosine 3TMS | 218.1 | 179.1 | 280.2 |
| Inositol 6TMS | 305.1 | 318.1 | 507.3 |
| Tryptophan 2TMS | 218.1 | 130 | 348.2 |
| Cystine 4TMS | 218.1 | 297.1 | 411.2 |

### Data processing and normalization

All GC-MS chromatograms were processed using MetaboliteDetector, v3.2.20190704 (Hiller et al., 2009). Compounds were annotated by retention time and mass spectrum using an in-house mass spectral (SIM) library (overall similarity: >0.85). The following deconvolution settings were applied: Peak threshold: 2; Minimum peak height: 2; Bins per scan: 10; Deconvolution width: 8 scans; No baseline adjustment; Minimum 1 peaks per spectrum; No minimum required base peak intensity.

The internal standards were added at the same concentration to every medium sample to correct for uncontrolled sample losses and analyte degradation during metabolite extraction. The data set was normalized by using the response ratio of the integrated peak area (QI)-analyte and the integrated peak area(QI)-internal standard.

### DIABLO and MINT analysis

Data was generated from three WT and three 3xSNCA MOs replicate samples where simultaneously RNA was extracted and media was collected. Metabolite and transcriptomic data was integrated using DIABLO (Singh et al., 2018) vignette: http://mixomics.org/mixdiablo/case-study-tcga/ and https://mixomicsteam.github.io/Bookdown/diablo.html#principle-of-diablo. Script for data analysis is included in the corresponding Gitlab repository.

## Ethics Statement

Written informed consent was obtained from all individuals who donated samples to this study and all work with human stem cells was done after approval of the national ethics board, Comité National d’Ethique de Recherche (CNER), under the approval numbers 201305/04 and 201901/01.

## Code Availability

The codes used for image analysis are included in: https://github.com/LCSB-DVB/Modamio_2021

## Data Availability

All data can be found under: https://doi.org/10.17881/1yzp-qv41

## Acknowledgements

The authors thank Prof. Dr. Hans R. Schöler from the Max-Planck-Gesellschaft, Prof. Dr. Thomas Gasser and Christine Klein from the Universitats klinikum Tuebingen, Dr. Jared Sterneckert from the CRTD, Prof. Dr. Rejko Krüger, and Gabriela Novak from the University of Luxembourg for providing us with cell lines. Moreover, we thank Isabel Rosety for the derivation of two cell lines. We thank the LCSB Metabolomics Platform, and specially Christian Jäger and Xiangyi Dong for their contribution to this manuscript.

J.J. is supported by a Pelican award from the Fondation du Pelican de Mie et Pierre Hippert-Faber. The JCS lab is supported by the Fonds National de la Recherche (FNR) Luxembourg (BRIDGES18/BM/12719664_MOTASYN; INTER/FWF/19/14117540/PDage). This project has received funding from the European Union’s Horizon 2020 research and innovation programme H2020-FETPROACT-2018-01 under grant agreement No 824070. This work was supported by the U.S. Army Medical Research Materiel Command endorsed by the U.S. Army through the Parkinson’s Research Program Investigator-Initiated Research Award under Award No. W81XWH-17-PRP-IIRA. Opinions, interpretations, conclusions and recommendations are those of the author and are not necessarily endorsed by the U.S. Army. EG and JCS would like to acknowledge support from the Fondation Gustave et Simone Prevot. We also would like to thank the private donors who support our work at the Luxembourg Centre for Systems Biomedicine.

## Rights retention Statement

“*This research was funded in whole or in part by [Institute and Grant number]. For the purpose of Open Access, the author has applied a CC BY public copyright license to any Author Accepted Manuscript (AAM) version arising from this submission*.”

## Author statement

Jennifer Modamio: Formal analysis, Investigation, Methodology, Data curation, Writing-Original draft preparation. Claudia Saraiva: Investigation, Writing -Review & Editing. Gemma Gomez Giro: Investigation, Writing -Review & Editing. Sarah Louise Nickels: Formal analysis, Writing -Review & Editing. Javier Jarazo: Software. Paul Antony: Software. Silvia Bolognin: Software. Peter Barbuti: Resources. Rashi Hadler: Data Curation. Christian Jäger: Data Curation. Rejko Krüger: Resources. Enrico Glaab: Formal analysis. Jens Christian Schwamborn: Supervision, Conceptualization, Resources, Writing -Review & Editing

