## Supplementary Information for "Synaptic decline precedes dopaminergic neuronal loss in human midbrain organoids harboring a triplication of the *SNCA* gene"

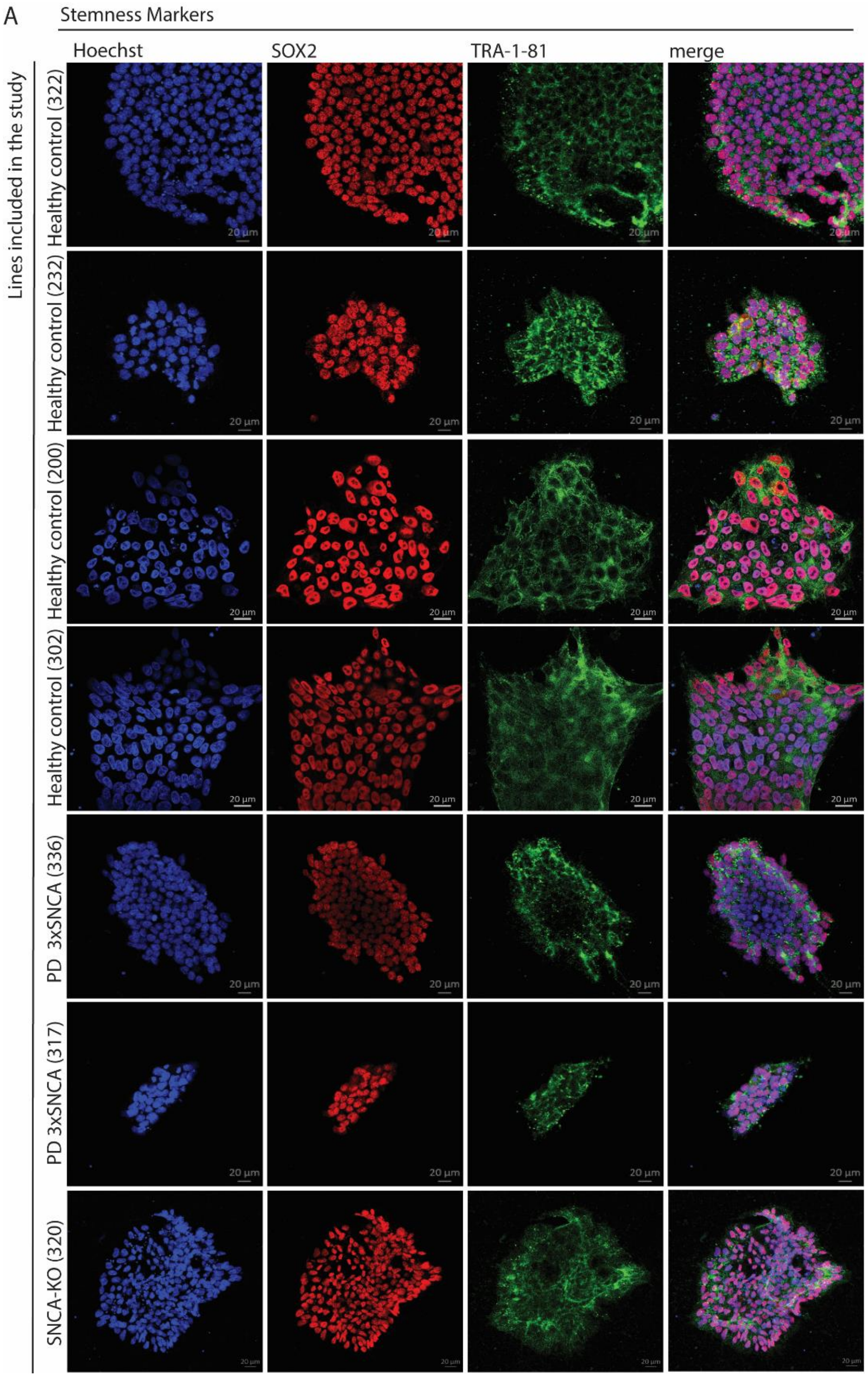


**Supplementary figure 1: HiPSCs showed positive immunostaining for the stemness markers SOX2 and TRA-1-81.** (A) Representative images from confocal microscope (60X) of iPSCs used in the current study showed positive signal for the nuclear stemness marker SOX2 (red) and the surface stemness marker TRA-1-81 (green). DNA stained by hoechst (blue).

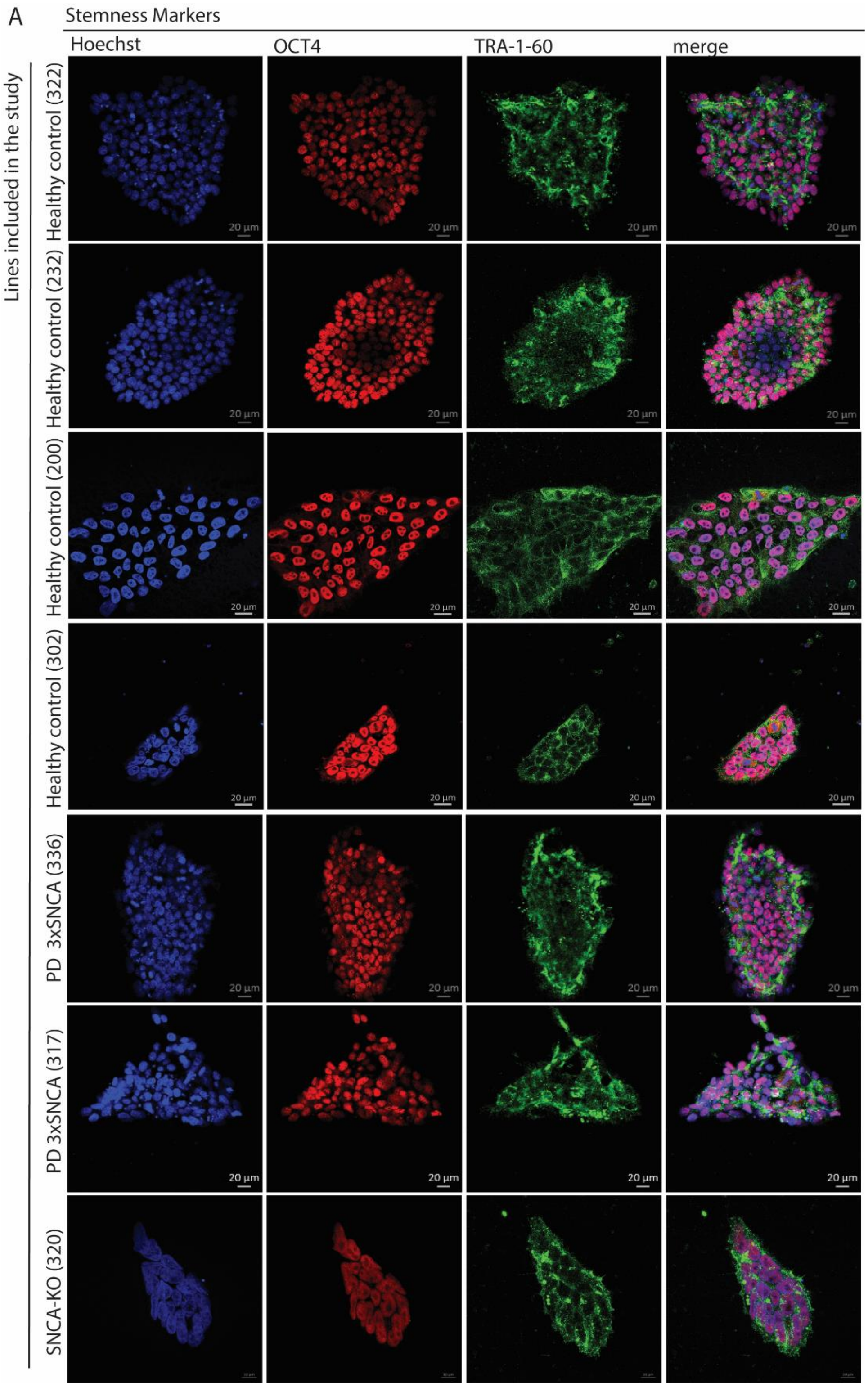


**Supplementary figure 2: HiPSCs showed positive immunostaining for the stemness markers OCT4 and TRA-1-60.** (A) Representative images from confocal microscope (60X) of iPSCs used in the current study showed positive signal for the nuclear stemness marker OCT4 (red) and the surface stemness marker TRA-1-60 (green). DNA stained by hoechst (blue).

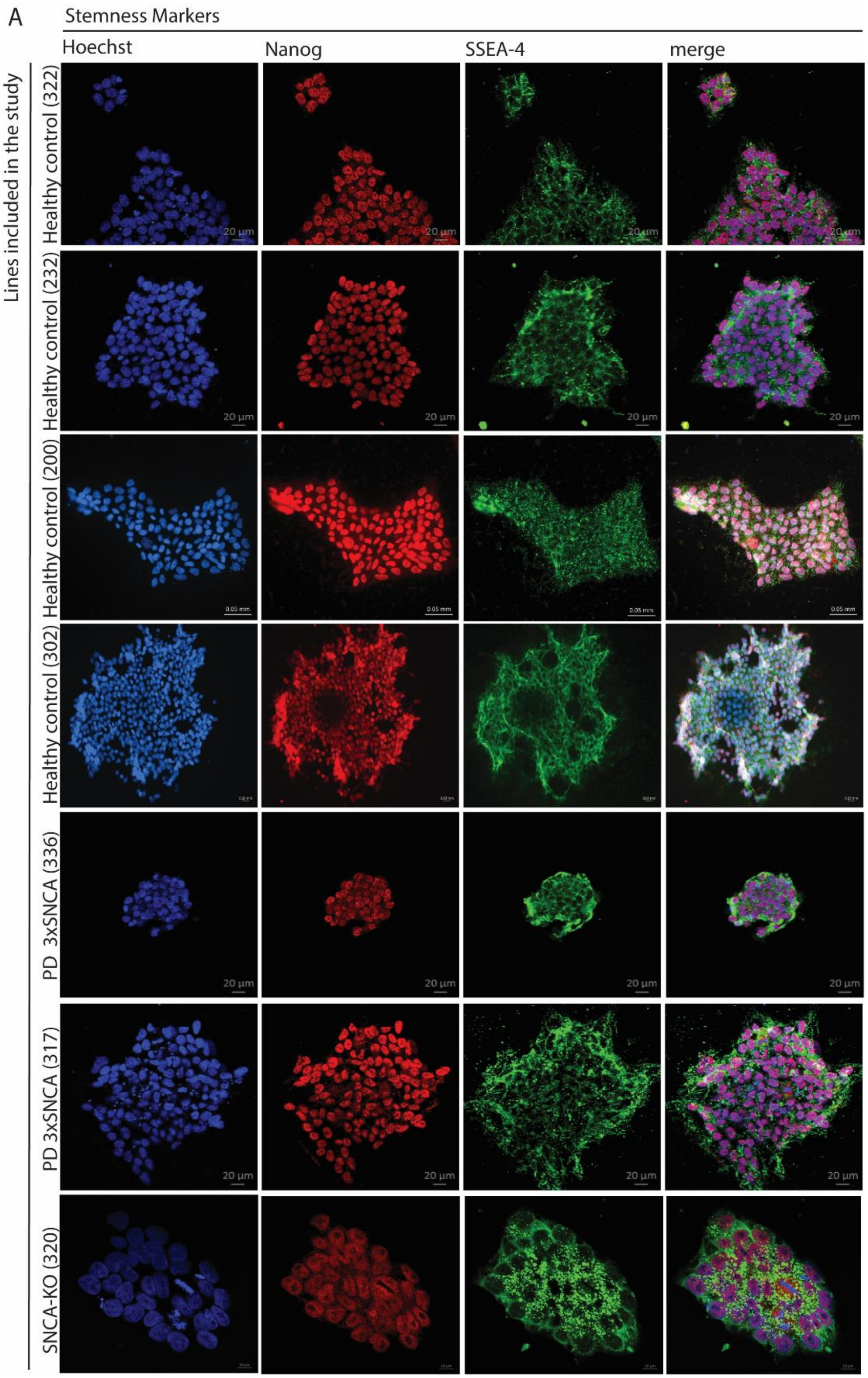


**Supplementary figure 3: HiPSCs showed positive immunostaining for the stemness markers NANOG and SSEA4.** (A) Representative images from confocal microscope (60X) of iPSCs used in the current study showed positive signal for the nuclear stemness marker NANOG (red) and the surface stemness marker SSEA4 (green). DNA stained by hoechst (blue).

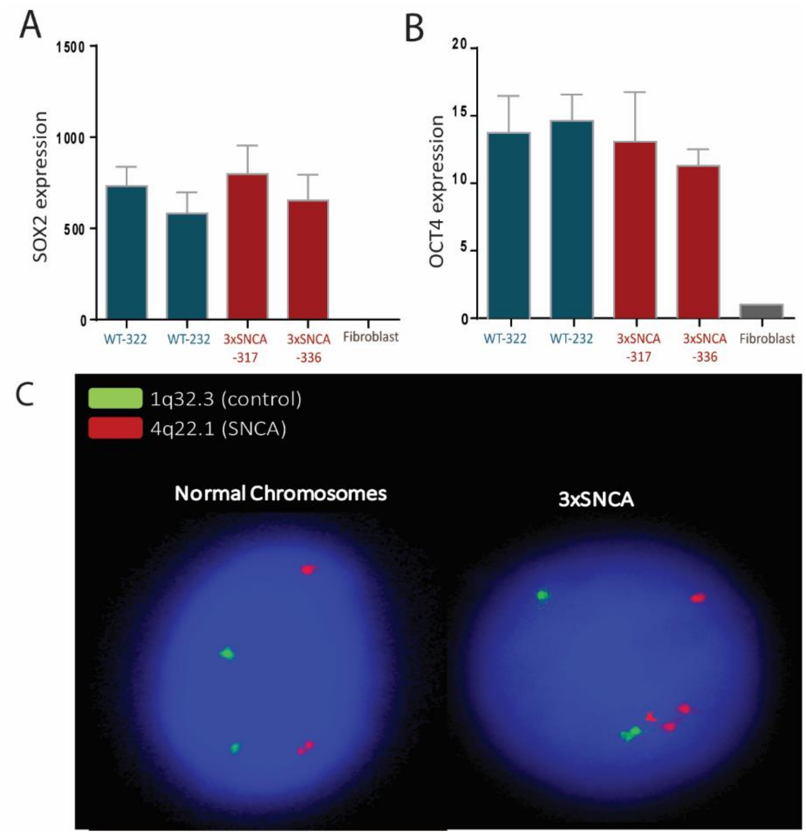


**Supplementary figure 4: RT-qPCR confirmed positive expression for stemness markers and FISH analysis confirms the presence of *SNCA* multiplication.** (A-B) RT-qPCR analysis for the stemness markers *SOX2* (A) and *OCT4* (B) in iPSCs. Fibroblast were used as negative control, ± SEM, ΔΔ Ct method (3 technical, 1 biological replicate (n=1)). (C) Representative image of Fluorescence *in situ* hybridation (FISH) of a normal condition (ratio 2G/2R) and a *SNCA* triplication condition (ratio 2G/4R) (2G = 2 green – control gene, 2R = 2 red – *SNCA* gene).

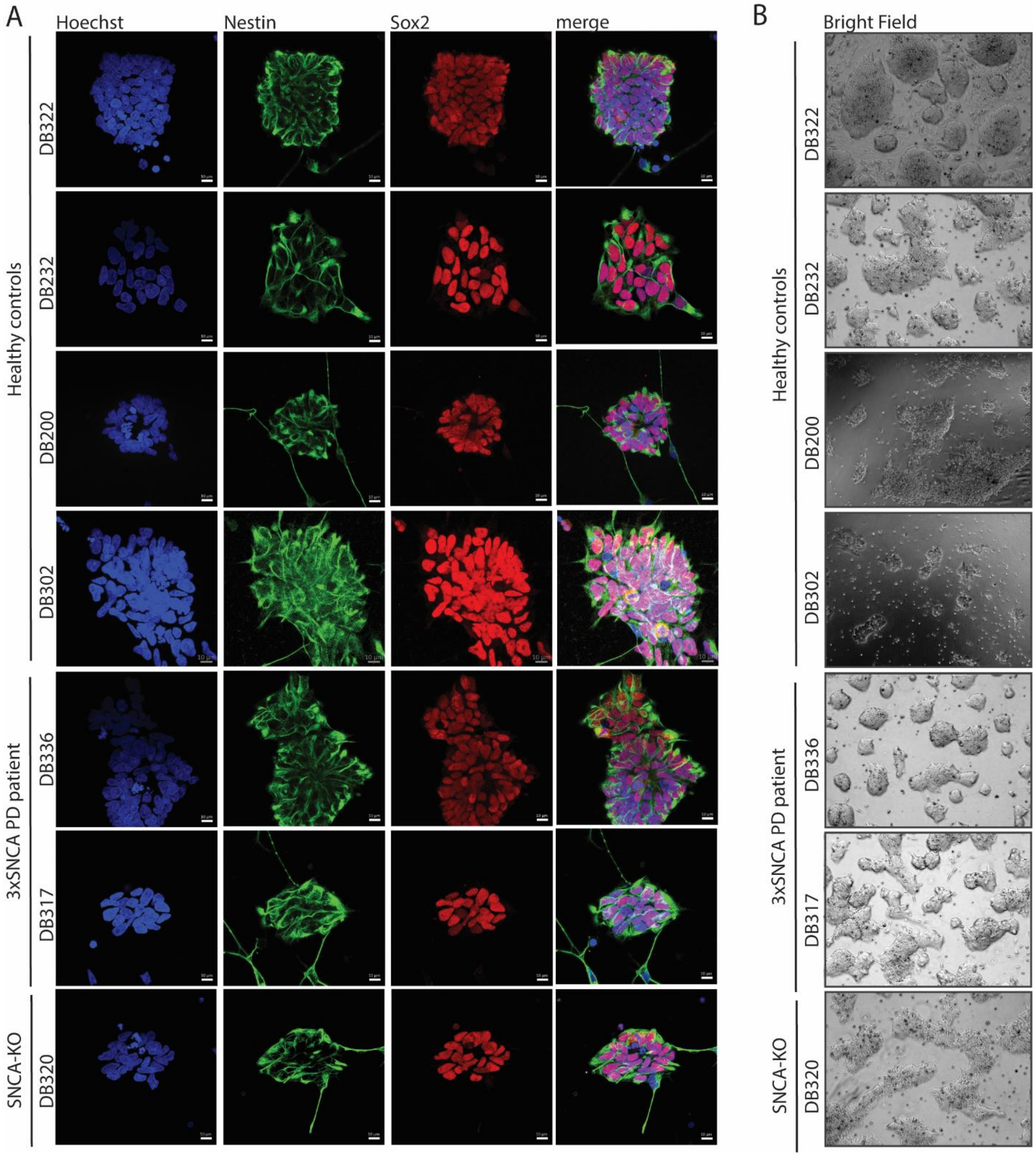


**Supplementary figure 5: mfNPCs presented midbrain ventral identity.** (A) Panel with representative images from confocal microscope (60X). mfNPCs presented positive signal for the stemness markers SOX2 and NESTIN. (B) Representative bright-field image (5X) from mfNPCs colonies.

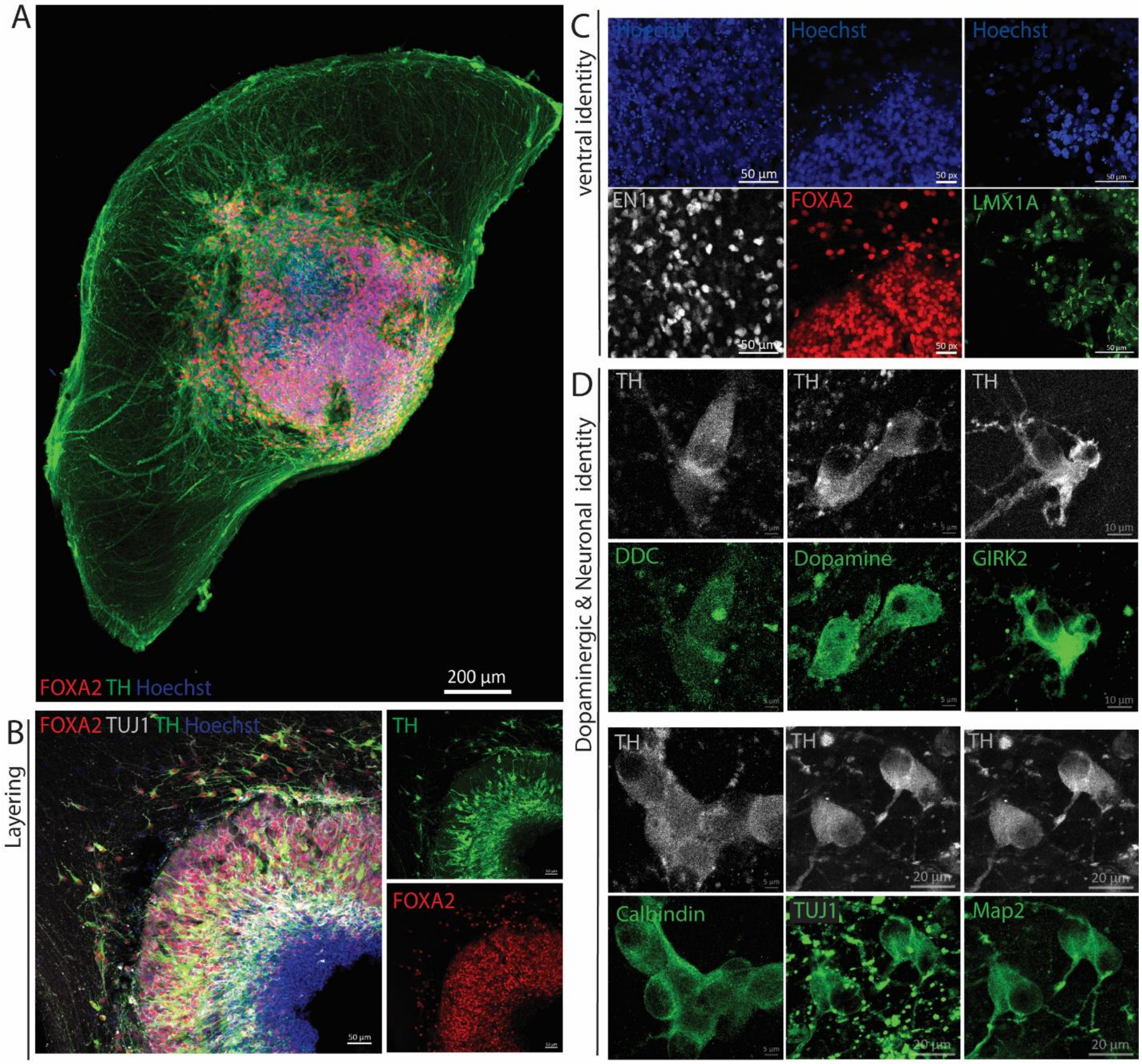


**Figure S6: Characterization of the MO model.** (A) Representative confocal image of a 30 days-old MO with positive signal for FOXA2, TUJ1 and TH. (B) Representative confocal image (20X) of a MO layering, with a central stem cell niche and peripheral differentiated cells. (C) Representative confocal images for the ventral markers FOXA2, LMX1A and EN1. (D) Representative confocal images for early (TUJ1) and mature (MAP2) neuronal markers, as well as dopaminergic neuronal markers such as TH, DDC, Dopamine, GIRK2 and Calbindin.

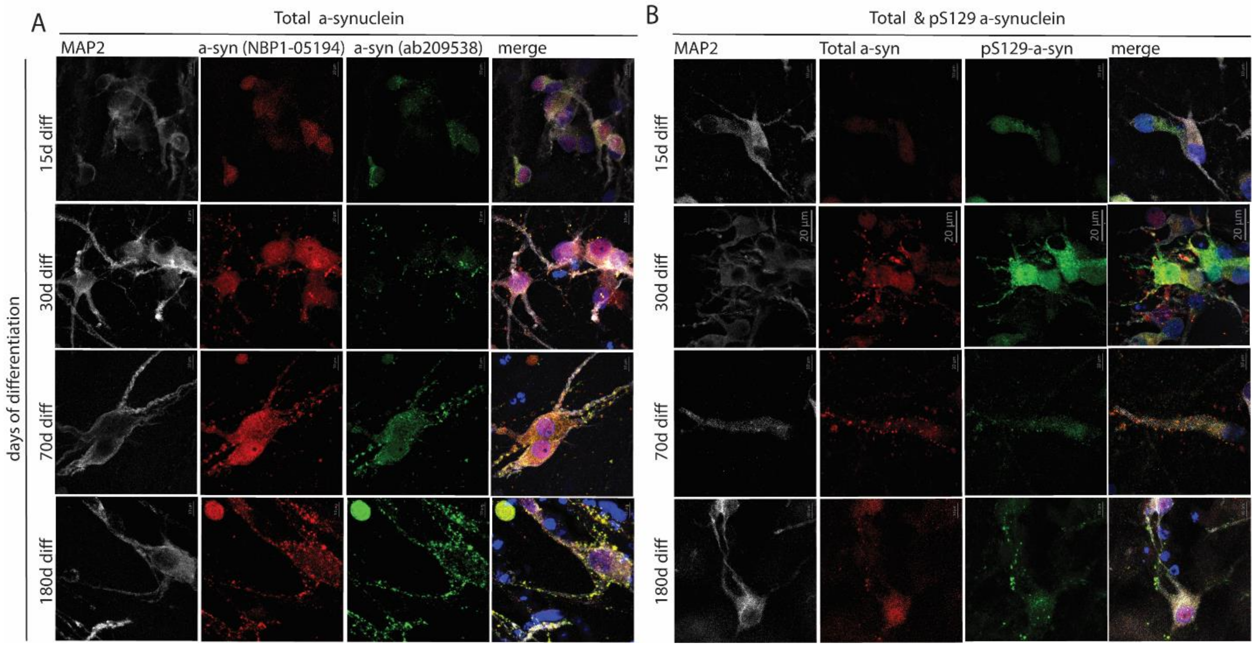


**Supplementary figure 7: Subcellular location of α-syn during the maturation process and vesicle-like dotted α-syn.** Representative confocal images (60X) of α-syn subcellular location over time. (A) Total α-syn (ab: NBP1-05194) and conformation-specific α-syn (ab: ab209538 - signal loss in denaturalized α-syn). Colocalization of both markers at several time points. (B) Total α-syn and pS129 α-syn. Colocalization of both markers at several time points.

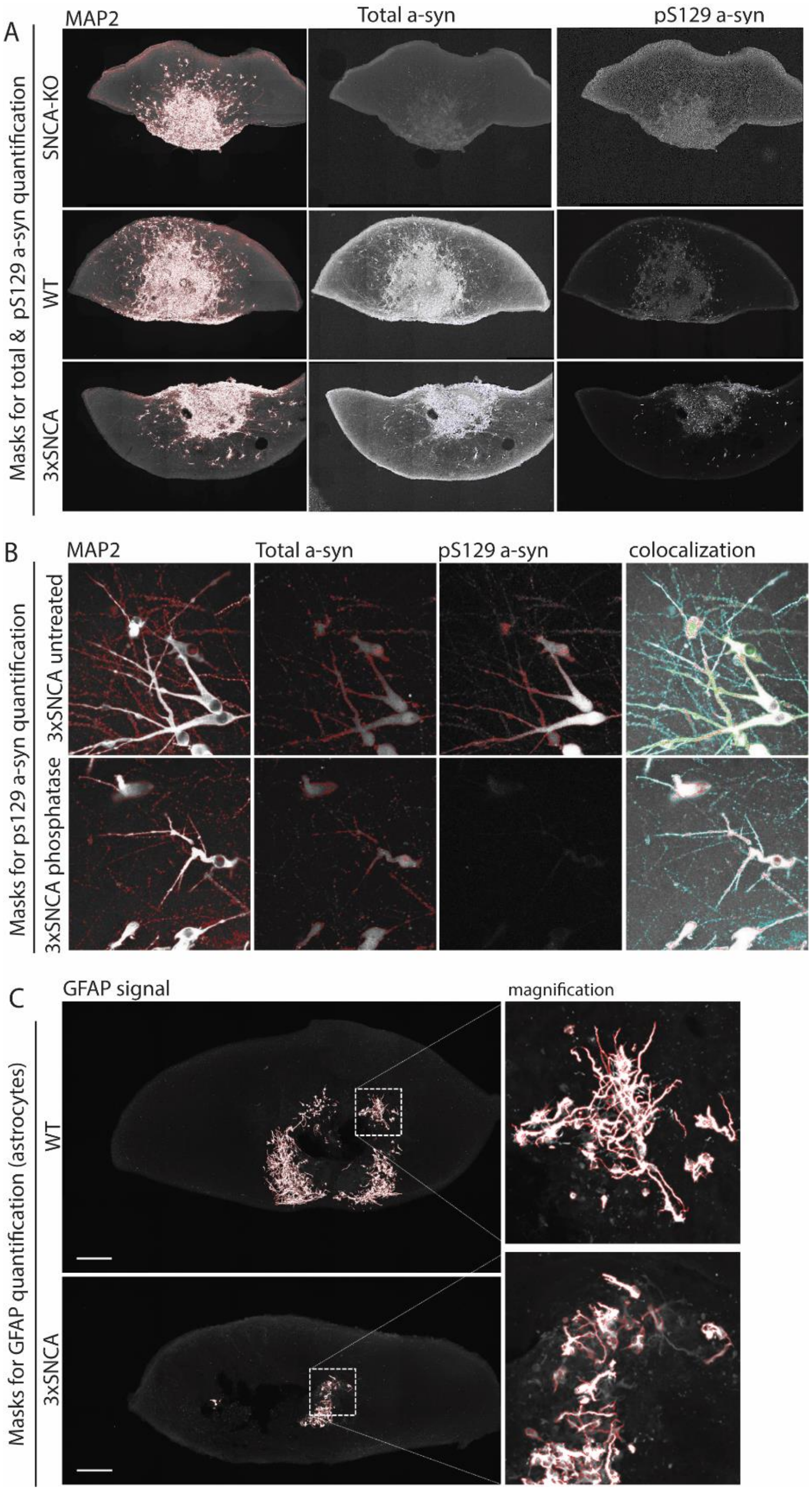


**Supplementary figure 8: Positive masks for high content image analysis. (A)** Representative masks used for quantification of MAP2, total α-syn and pS129 α-syn corresponding to the representative images from (Fig. 2C). (B) Representative masks used for the quantification of MAP2, total α-syn and pS129 α-syn corresponding to the representative images from (Fig. 2A). (C) Representative masks used for quantification of GFAP+ corresponding to the representative images from (Fig. 3C). Zoomed images show the mask consider as positive signal for the quantification.

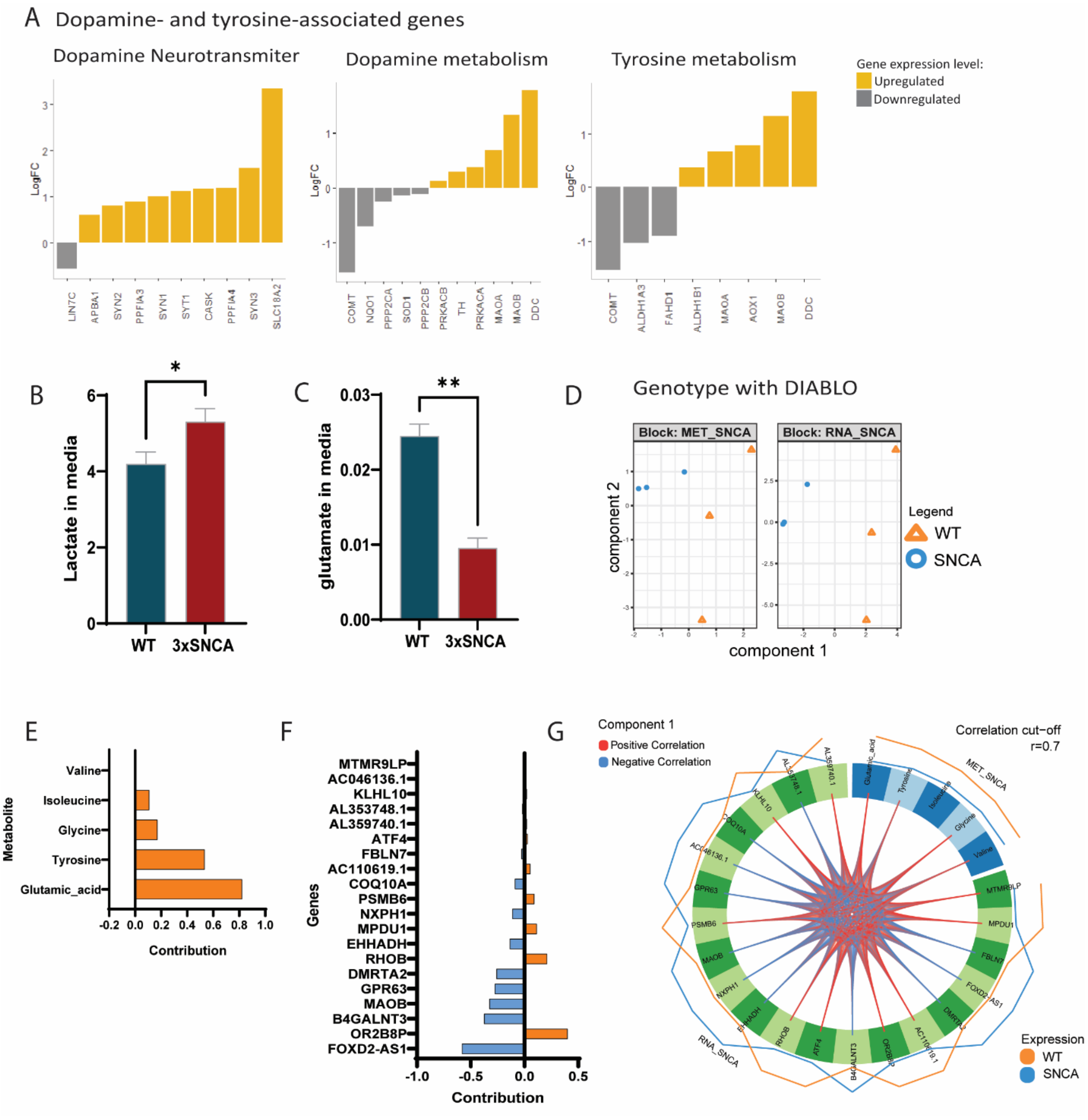


**Supplementary figure 9: Combinatorial analysis of transcriptomic and metabolomic data.** (A) Significantly deregulated genes associated to the dopamine neurotransmitter, dopamine metabolism and tyrosine metabolism (FDR < 0.05 or pValue < 0.05). (B) Box-plot for Lactate in media (*p < 0.05). (C) Box-plot for glutamate in media (****p < 0.0001). (D) Genotype with DIABLO analysis. Discriminative PCA (component 1 & 2) for transcriptomic and metabolomic data from top selected genes and metabolites. (E) Contribution plot from top extracellular metabolites selected in DIABLO analysis (5 metabolites selected in the discriminative PCA – component 1). (F) Contribution plot from top genes selected in DIABLO analysis (20 genes selected in the discriminative PCA – component 1). (E) Correlation network between top metabolites and genes from component 1 (abs(r) >= 0.7).

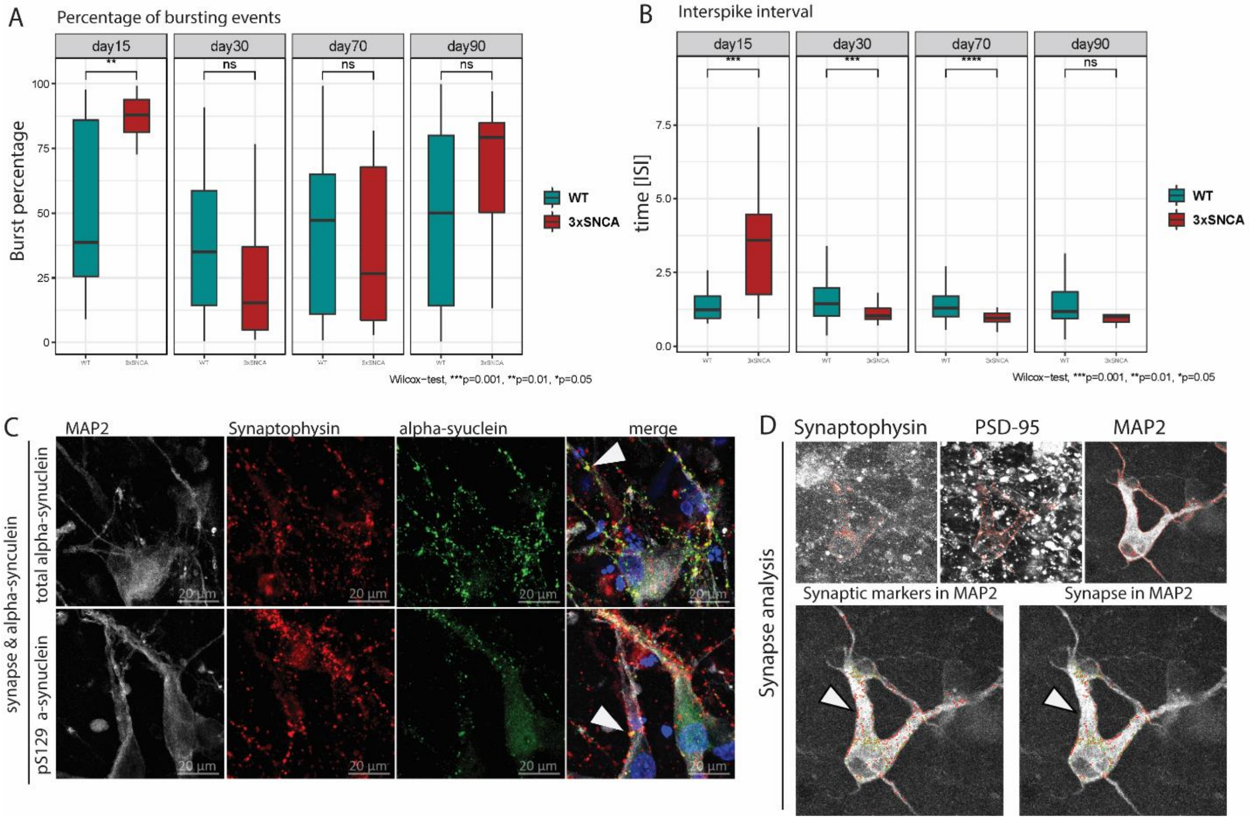


**Supplementary figure 10: Reduced synaptic activity and synapse count in 3xSNCA MOs.** (A) Percentage or bursting event over time in 3xSNCA and WT MOs. (B) Interspike interval over time in 3xSNCA and WT MOs. (C) Co-localization of the presynaptic marker synaptophysin with total α-syn and pS129 α-syn. (D) Masks used for the quantification of synaptic events from (Fig. 6A).

**Table S1: List of cell lines used in the present study.**

| Biological samples | | |
|---|---|---|
| **Database number** | **Origen** | **Gender, Age, condition** |
| DB 322 | Grossmann et al, 2019 | Female, 55 years, WT healthy |
| DB 232 | Reinhardt et al, 2013 | Female, 53 years, WT healthy |
| DB 200 | Reinhardt et al, 2013 | Female, 81 years, WT healthy |
| DB 302 | Universitats klinikum Tuebingen | Female, 68 years, WT healthy |
| DB 336* | EBISC Edi001_A01 | Female, 55 years, PD 3xSNCA |
| DB 317* | Coriell ND27760 | Female, 55 years, PD 3xSNCA |
| DB 320 | Barbuti et al. *in preparation* | Male, 67 years, WT-GE SNCA-KO |

WT, Wild type; PD, Parkinson's Disease; GE, genetically engineered; (*), same donor, independent clone

**Table S1:** Information for each iPSC line used in this project. Internal database number, cell line origen, gender, age and condition.

**Table S2: Primers for qPCR used in the iPSCs characterization.**

| Primers | | | |
|---|---|---|---|
| **Gene** | **Forward/Reverse** | **sequence** | **Company** |
| OCT-4 | h-OCT4-F | GTGGAGGAAGCTGACAACAA | Sigma |
| OCT-4 | h-OCT4-R | ATTCTCCAGGTTGCCTCTCA | Sigma |
| SOX2 | hSOX2 endo F | TGGCGAACCATCTCTGTGGT | Eurogentec |
| SOX2 | hSOX2 endo R | CCAACGGTGTCAACCTGCAT | Eurogentec |

**Table S2:** Primer sequence used for the analysis of the stemness markers OCT-4 and SOX2.

**Table. S3A: LogFC significantly deregulated genes.**

| Gene symbol | logFC | logCPM | F | PValue | FDR |
|---|---|---|---|---|---|
| CCNYL2 | 7.243903 | 2.151256 | 436.0476 | 2.59E-09 | 2.69E-05 |
| CTSF | -9.74738 | 5.184203 | 448.8375 | 2.62E-09 | 2.69E-05 |
| ZNF558 | -12.5826 | 3.929767 | 805.97 | 6.19E-09 | 2.98E-05 |
| ZNF736 | 7.656001 | 3.096631 | 344.3107 | 8.98E-09 | 2.98E-05 |
| ZXDA | 5.380893 | 2.451911 | 332.6284 | 9.41E-09 | 2.98E-05 |
| PCDHGB4 | 5.183086 | 2.95387 | 334.2692 | 1.03E-08 | 2.98E-05 |
| CUTALP | -3.9586 | 4.070692 | 332.8783 | 1.05E-08 | 2.98E-05 |
| C5orf17 | -5.26645 | 2.361398 | 316.8681 | 1.16E-08 | 2.98E-05 |
| PCDHB5 | 3.13969 | 4.185071 | 304.4568 | 1.39E-08 | 3.07E-05 |
| LINC00839 | 12.1785 | 3.526411 | 648.1414 | 1.52E-08 | 3.07E-05 |
| AC079949.2 | 8.611137 | 1.849903 | 301.8007 | 1.65E-08 | 3.07E-05 |
| ZNF229 | 3.959082 | 4.034348 | 294.4481 | 1.85E-08 | 3.15E-05 |
| GABRG3 | 4.212878 | 4.9109 | 270.4563 | 2.73E-08 | 4.31E-05 |
| TCEAL5 | 3.683467 | 5.647359 | 261.2193 | 3.21E-08 | 4.69E-05 |
| HTR3A | 6.073423 | 3.814823 | 244.2865 | 4.36E-08 | 5.95E-05 |
| A2M-AS1 | -5.84998 | 1.55674 | 239.2705 | 4.72E-08 | 6.04E-05 |
| MTND4P12 | 2.5108 | 6.677763 | 235.1633 | 5.19E-08 | 6.07E-05 |
| TCERG1L | 11.25155 | 2.608606 | 461.7234 | 5.34E-08 | 6.07E-05 |
| TMEM132B | 3.247346 | 5.385478 | 230.6875 | 5.67E-08 | 6.11E-05 |
| STUM | 3.111856 | 3.606777 | 219.5963 | 6.65E-08 | 6.81E-05 |
| AJAP1 | 2.218388 | 5.397745 | 191.6945 | 1.19E-07 | 0.000112 |
| TMEM132C | 10.68531 | 2.048897 | 369.4276 | 1.21E-07 | 0.000112 |
| PCDHGA11 | 5.534924 | 0.517499 | 186.2163 | 1.35E-07 | 0.000112 |
| AC069277.1 | 5.643679 | 1.140904 | 188.671 | 1.42E-07 | 0.000112 |
| LINC00654 | 5.172987 | 2.018105 | 188.4558 | 1.42E-07 | 0.000112 |
| NPY | 4.252355 | 3.337429 | 185.917 | 1.51E-07 | 0.000112 |
| POU6F2 | 3.005282 | 3.803736 | 184.3668 | 1.57E-07 | 0.000112 |
| PCDHGA7 | 7.943698 | 1.969963 | 184.0963 | 1.58E-07 | 0.000112 |
| PTPRT | 10.94133 | 2.303042 | 342.7327 | 1.6E-07 | 0.000112 |
| DPP6 | 2.172069 | 5.226497 | 178.4406 | 1.64E-07 | 0.000112 |
| TMEM255A | 1.860437 | 5.663434 | 175.2866 | 1.78E-07 | 0.000118 |
| NNMT | -2.2462 | 4.727873 | 171.9281 | 1.94E-07 | 0.000121 |
| EEF1GP1 | 2.533966 | 3.924972 | 172.4861 | 1.95E-07 | 0.000121 |
| AC079062.1 | 7.738777 | 1.769787 | 174.2112 | 2.03E-07 | 0.000122 |
| POU3F4 | 4.560591 | 3.300145 | 169.3956 | 2.31E-07 | 0.000135 |
| PCDHA5 | 4.613308 | 2.795469 | 165.0293 | 2.59E-07 | 0.000144 |
| TMC3 | 3.993454 | 1.351988 | 161.5517 | 2.61E-07 | 0.000144 |
| TEKT4P2 | -9.8107 | 1.201427 | 286.432 | 2.69E-07 | 0.000145 |
| NNAT | 9.314708 | 9.312468 | 162.7207 | 2.76E-07 | 0.000145 |
| AL132780.4 | 5.127891 | 0.651619 | 158.374 | 2.95E-07 | 0.000151 |
| UGT2B4 | 4.347293 | 1.401453 | 158.4039 | 3.12E-07 | 0.000156 |
| PCDHGA3 | 7.381796 | 0.664869 | 154.2172 | 3.52E-07 | 0.000171 |
| FEV | 3.774647 | 4.927502 | 153.4627 | 3.6E-07 | 0.000171 |
| LINC01060 | 6.210102 | 0.792153 | 151.0214 | 3.86E-07 | 0.000177 |

| | | | | | |
|---|---|---|---|---|---|
| TP53TG3F | 9.591516 | 0.985638 | 259.0776 | 3.89E-07 | 0.000177 |
| FAM19A5 | 3.213454 | 6.281302 | 149.3642 | 4.06E-07 | 0.000181 |
| ZP3 | -2.54341 | 3.181243 | 144.6416 | 4.26E-07 | 0.000186 |
| PSPHP1 | -2.75096 | 2.475571 | 140.1354 | 4.92E-07 | 0.00021 |
| TP53TG3B | 9.473585 | 0.871936 | 239.7733 | 5.18E-07 | 0.000213 |
| FMO3 | -5.3009 | 1.645342 | 141.321 | 5.2E-07 | 0.000213 |
| MIR4458HG | 2.136422 | 4.273787 | 135.7053 | 5.84E-07 | 0.000225 |
| FMOD | 2.46061 | 3.13278 | 134.8306 | 5.85E-07 | 0.000225 |
| CXCR4 | 2.99487 | 5.712683 | 136.9719 | 5.98E-07 | 0.000225 |
| FAR2P1 | -2.74565 | 5.398995 | 136.6225 | 6.05E-07 | 0.000225 |
| HLA-A | -1.83652 | 5.733763 | 135.0291 | 6.11E-07 | 0.000225 |
| AC109439.2 | 2.834616 | 2.338089 | 133.7302 | 6.16E-07 | 0.000225 |
| FIRRE | 2.267656 | 2.912134 | 131.3268 | 6.58E-07 | 0.000236 |
| CSRP1 | 3.269589 | 5.99451 | 132.1484 | 7.02E-07 | 0.000248 |
| DAZAP2P1 | 4.401472 | 0.189539 | 128.4151 | 7.28E-07 | 0.000253 |
| LINC00865 | 6.63669 | 0.707278 | 129.4778 | 7.69E-07 | 0.000262 |
| PAX5 | 2.995007 | 3.057225 | 128.6228 | 7.92E-07 | 0.000266 |
| POMZP3 | -2.05743 | 3.14119 | 119.4277 | 1.01E-06 | 0.000333 |
| TENM1 | 3.019146 | 5.138445 | 120.5719 | 1.05E-06 | 0.000343 |
| NSDHL | 1.54721 | 5.620728 | 115.4333 | 1.17E-06 | 0.000375 |
| AC139491.7 | -4.90095 | 0.664249 | 116.851 | 1.21E-06 | 0.000381 |
| GFAP | -3.4144 | 5.214701 | 116.218 | 1.24E-06 | 0.000385 |
| CYP4F24P | 8.954809 | 0.376627 | 186.7456 | 1.3E-06 | 0.000397 |
| TLR4 | 3.230765 | 4.532381 | 112.4609 | 1.43E-06 | 0.000425 |
| BRS3 | -2.11371 | 4.898632 | 112.2034 | 1.45E-06 | 0.000425 |
| CBLN4 | 5.966097 | 0.562677 | 112.1107 | 1.45E-06 | 0.000425 |
| TRIM58 | 3.705562 | 0.738562 | 111.1043 | 1.48E-06 | 0.000426 |
| GATA2 | 2.060355 | 4.34492 | 111.1522 | 1.51E-06 | 0.000429 |
| PCDHGA10 | 4.797073 | 3.21937 | 106.4993 | 1.82E-06 | 0.000511 |
| TMOD1 | 1.930148 | 3.88943 | 104.4205 | 1.86E-06 | 0.000515 |
| RHOXF1-AS1 | 5.159383 | 0.453768 | 104.8754 | 1.95E-06 | 0.00053 |
| CU633967.1 | -1.82594 | 5.911788 | 104.6306 | 1.97E-06 | 0.00053 |
| ZNF619 | 3.061976 | 1.50833 | 103.8697 | 2.03E-06 | 0.000541 |
| OTOF | 3.473943 | 2.04997 | 103.3378 | 2.08E-06 | 0.000546 |
| AFF2 | 1.743253 | 7.091104 | 101.6043 | 2.24E-06 | 0.00058 |
| ZNF572 | 6.651652 | 0.730678 | 101.1195 | 2.29E-06 | 0.000585 |
| HEPACAM2 | 4.88538 | 0.202344 | 100.8014 | 2.32E-06 | 0.000585 |
| LINC00682 | 2.583869 | 2.025833 | 99.65542 | 2.34E-06 | 0.000585 |
| KDR | 3.798564 | 1.459465 | 99.52887 | 2.45E-06 | 0.000597 |
| CU633904.1 | -1.93358 | 4.587379 | 99.52384 | 2.45E-06 | 0.000597 |
| URB1-AS1 | 2.777056 | 2.000639 | 98.89908 | 2.52E-06 | 0.000607 |
| OR2L13 | 3.153483 | 2.907541 | 98.61304 | 2.55E-06 | 0.000607 |
| AC011477.2 | 4.190662 | -0.41246 | 96.50003 | 2.59E-06 | 0.000609 |
| PHOX2B | 3.000231 | 4.11338 | 98.04169 | 2.62E-06 | 0.000609 |
| ANKRD1 | 3.42946 | 0.242048 | 95.87131 | 2.67E-06 | 0.00061 |
| FSTL4 | 1.99774 | 3.381108 | 96.31759 | 2.68E-06 | 0.00061 |
| TRIM60P18 | 6.65197 | -0.01289 | 96.97606 | 2.74E-06 | 0.000617 |

| | | | | | |
|---|---|---|---|---|---|
| ADGRG5 | 3.518829 | 0.72831 | 96.44109 | 2.81E-06 | 0.000626 |
| CU634019.1 | -1.9727 | 4.339878 | 94.89726 | 3.02E-06 | 0.000661 |
| PSMD5 | -2.52287 | 4.821313 | 94.75426 | 3.04E-06 | 0.000661 |
| IRF6 | 4.033492 | 1.382643 | 94.29698 | 3.1E-06 | 0.000661 |
| AC138907.8 | 5.127322 | -0.20865 | 93.97972 | 3.15E-06 | 0.000661 |
| TRDN | 3.084607 | 0.202099 | 92.29257 | 3.15E-06 | 0.000661 |
| TOGARAM2 | -2.76243 | 3.310557 | 93.7071 | 3.19E-06 | 0.000661 |
| STC1 | 1.825411 | 6.218487 | 93.50547 | 3.22E-06 | 0.000661 |
| CBLN1 | 2.603886 | 2.380889 | 93.32017 | 3.24E-06 | 0.000661 |
| TP53TG3 | 8.525437 | -0.02271 | 146.9176 | 3.26E-06 | 0.000661 |
| AC079160.1 | 3.992385 | -0.16947 | 89.87966 | 3.54E-06 | 0.000701 |
| AC016582.3 | 2.557726 | 2.967054 | 91.38985 | 3.55E-06 | 0.000701 |
| MAGED1 | 1.494697 | 9.668881 | 89.90832 | 3.56E-06 | 0.000701 |
| ILDR2 | 1.939942 | 5.100948 | 90.92171 | 3.63E-06 | 0.000708 |
| CHCHD2 | -1.36634 | 6.379506 | 88.80828 | 3.73E-06 | 0.000715 |
| NBPF2P | 6.324775 | -0.31909 | 90.30507 | 3.74E-06 | 0.000715 |
| KLHL13 | 1.787135 | 6.065684 | 89.85291 | 3.82E-06 | 0.000715 |
| LINC02511 | -3.10708 | 0.96397 | 89.433 | 3.9E-06 | 0.000715 |
| LINC01266 | 4.229003 | -0.37691 | 87.86803 | 3.91E-06 | 0.000715 |
| DUSP9 | 2.532355 | 2.109236 | 89.38468 | 3.91E-06 | 0.000715 |
| LEFTY2 | -2.2989 | 6.775775 | 89.37411 | 3.91E-06 | 0.000715 |
| TP53TG3E | 5.896864 | 0.020468 | 88.88277 | 4.01E-06 | 0.000726 |
| SMIM31 | -2.8215 | 1.791259 | 88.36936 | 4.11E-06 | 0.000738 |
| MMP24OS | 1.87102 | 3.029395 | 86.38661 | 4.21E-06 | 0.000749 |
| ZNF506 | 2.740179 | 2.922313 | 86.60826 | 4.48E-06 | 0.000787 |
| MYOM2 | 1.85241 | 3.257975 | 85.0793 | 4.5E-06 | 0.000787 |
| MECP2 | 1.227249 | 7.118486 | 84.7551 | 4.57E-06 | 0.000787 |
| FRMPD1 | 2.519367 | 1.884788 | 86.17055 | 4.58E-06 | 0.000787 |
| ZNF603P | -2.45922 | 2.090084 | 86.01903 | 4.61E-06 | 0.000787 |
| TUBB8P7 | 2.342169 | 1.416899 | 84.03496 | 4.74E-06 | 0.000802 |
| NLGN4Y | 1.870007 | 3.539672 | 85.04092 | 4.78E-06 | 0.000802 |
| CPXM2 | 2.093145 | 2.227621 | 83.05533 | 4.99E-06 | 0.000831 |
| COLEC12 | -3.36919 | 4.117786 | 84.26987 | 5.04E-06 | 0.000832 |
| MLXIPL | 2.32934 | 2.062036 | 83.53336 | 5.11E-06 | 0.000837 |
| MAF | -1.72043 | 6.670163 | 83.56851 | 5.23E-06 | 0.000844 |
| TPH2 | 4.866023 | 5.423391 | 83.29833 | 5.3E-06 | 0.000844 |
| LYPD6 | 1.687133 | 2.992504 | 81.83955 | 5.32E-06 | 0.000844 |
| OTP | 4.516377 | 1.394157 | 83.21667 | 5.32E-06 | 0.000844 |
| MAP10 | 2.71958 | 1.992811 | 82.71784 | 5.46E-06 | 0.000854 |
| AC139491.5 | -6.06344 | -0.5225 | 82.61043 | 5.49E-06 | 0.000854 |
| USP11 | 1.504072 | 8.649508 | 82.5636 | 5.5E-06 | 0.000854 |
| NAP1L5 | 1.811205 | 8.973951 | 82.0416 | 5.66E-06 | 0.000871 |
| IRX2 | 1.881754 | 3.143878 | 80.92091 | 5.72E-06 | 0.000873 |
| ZFP3 | 2.358823 | 2.298345 | 81.52243 | 5.81E-06 | 0.000881 |
| AC068756.1 | 8.644815 | 0.085279 | 126.7002 | 5.88E-06 | 0.000884 |
| BEX5 | 2.439274 | 4.93557 | 80.76766 | 6.05E-06 | 0.000904 |
| TWIST1 | -2.28631 | 3.465029 | 80.59242 | 6.11E-06 | 0.000906 |

| | | | | | |
|---|---|---|---|---|---|
| PRSS12 | 2.504263 | 5.165708 | 80.43072 | 6.16E-06 | 0.000907 |
| LHX5 | 2.744528 | 0.322715 | 78.3265 | 6.43E-06 | 0.000939 |
| SLCO1A2 | 2.465537 | 2.597564 | 79.52125 | 6.47E-06 | 0.000939 |
| RAB39B | 1.332441 | 6.529975 | 78.67606 | 6.52E-06 | 0.000939 |
| KIF3B | 1.454893 | 7.998154 | 78.74817 | 6.74E-06 | 0.00096 |
| GABRA1 | 5.596172 | 1.098557 | 78.61537 | 6.79E-06 | 0.00096 |
| R3HDML | -2.4258 | 4.305801 | 78.59558 | 6.8E-06 | 0.00096 |
| AC073263.1 | -3.2805 | 0.129277 | 77.41897 | 6.86E-06 | 0.000961 |
| AP002852.1 | 3.583274 | -0.7024 | 76.65214 | 7.06E-06 | 0.000981 |
| KCNJ16 | 3.012056 | 3.058705 | 77.82684 | 7.09E-06 | 0.000981 |
| IQSEC2 | 1.634104 | 5.345973 | 77.67076 | 7.15E-06 | 0.000983 |
| SYT6 | 1.630712 | 4.539732 | 77.40914 | 7.25E-06 | 0.000984 |
| MAB21L1 | 4.723465 | 3.054641 | 77.3632 | 7.27E-06 | 0.000984 |
| LANCL3 | 1.647371 | 4.355524 | 77.2827 | 7.31E-06 | 0.000984 |
| FAM182B | 3.188219 | 0.691103 | 77.01051 | 7.42E-06 | 0.000992 |
| CHRNA3 | 2.05105 | 3.028058 | 76.37384 | 7.68E-06 | 0.001018 |
| TGFB3 | -1.63706 | 3.584285 | 75.10048 | 7.71E-06 | 0.001018 |
| IAPP | 4.593614 | 0.965723 | 74.80885 | 8.39E-06 | 0.001096 |
| BMP6 | -2.07397 | 5.096853 | 74.68658 | 8.45E-06 | 0.001096 |
| GDI2P1 | 2.804555 | 0.685219 | 74.06177 | 8.49E-06 | 0.001096 |
| AC133551.1 | 7.867737 | -0.61601 | 110.9544 | 8.55E-06 | 0.001096 |
| TMEM42 | -1.4658 | 3.735473 | 73.18574 | 8.61E-06 | 0.001096 |
| ZNF208 | 4.621178 | -0.65106 | 74.22523 | 8.62E-06 | 0.001096 |
| PLXDC2 | -1.9586 | 6.863074 | 74.06088 | 8.76E-06 | 0.001105 |
| PLTP | -1.6222 | 6.935772 | 73.98923 | 8.79E-06 | 0.001105 |
| SLC6A4 | 5.184564 | 5.502426 | 73.48943 | 9.05E-06 | 0.001124 |
| ZFP92 | 4.190583 | -0.1888 | 73.39641 | 9.1E-06 | 0.001124 |
| CLCN5 | 1.089539 | 7.025807 | 72.22268 | 9.11E-06 | 0.001124 |
| UTS2B | 1.406348 | 4.891963 | 72.14193 | 9.22E-06 | 0.00113 |
| SMIM10L2A | 1.320626 | 7.626235 | 73.04726 | 9.29E-06 | 0.001132 |
| ZNF528-AS1 | 2.779466 | 2.704055 | 72.85694 | 9.39E-06 | 0.001137 |
| AC092490.1 | 3.26372 | 0.86632 | 72.75762 | 9.44E-06 | 0.001137 |
| HERC5 | 2.404789 | 0.650837 | 71.25857 | 9.65E-06 | 0.001156 |
| OR2L3 | 4.108766 | -0.07307 | 72.26839 | 9.72E-06 | 0.001157 |
| BX890604.2 | 1.09397 | 6.846204 | 70.89988 | 9.86E-06 | 0.001166 |
| AL138899.1 | 8.126276 | -0.39084 | 109.3037 | 9.91E-06 | 0.001166 |
| SLC18A2 | 3.321849 | 2.117558 | 71.60916 | 1.01E-05 | 0.001176 |
| MFAP4 | -2.57911 | 5.388332 | 71.59152 | 1.01E-05 | 0.001176 |
| SLC7A8 | -1.50115 | 4.368 | 70.74243 | 1.02E-05 | 0.001181 |
| ZNF667-AS1 | 1.20065 | 5.474867 | 70.23696 | 1.03E-05 | 0.001181 |
| AC136612.1 | 8.056312 | -0.44928 | 108.0226 | 1.03E-05 | 0.001182 |
| NOP56P1 | 3.742189 | -0.78641 | 69.78127 | 1.06E-05 | 0.0012 |
| TSPAN7 | 1.422557 | 7.948338 | 70.11609 | 1.1E-05 | 0.001249 |
| HOXA10 | -2.88091 | 0.514812 | 69.86808 | 1.12E-05 | 0.001256 |
| AL138899.2 | 6.350132 | 0.435694 | 69.82057 | 1.12E-05 | 0.001258 |
| CNTN2 | 1.337904 | 6.331429 | 69.73107 | 1.13E-05 | 0.001258 |
| TP53TG3D | 7.621711 | -0.83322 | 102.1437 | 1.15E-05 | 0.00127 |

| | | | | | |
|---|---|---|---|---|---|
| OTOR | 4.422387 | 0.809316 | 68.5598 | 1.21E-05 | 0.001336 |
| PCDHGA6 | 4.570108 | -0.35237 | 68.1741 | 1.24E-05 | 0.001361 |
| LINC01630 | 2.112205 | 2.750287 | 67.96056 | 1.26E-05 | 0.001372 |
| PRPS2 | 1.315826 | 4.366304 | 66.20893 | 1.32E-05 | 0.001425 |
| NPTXR | 1.484817 | 6.759839 | 67.17092 | 1.32E-05 | 0.001425 |
| MYH14 | 1.808498 | 3.732501 | 66.87891 | 1.35E-05 | 0.001425 |
| PRSS3 | 4.66779 | 0.424332 | 66.7388 | 1.36E-05 | 0.001425 |
| SPINT1 | 3.218354 | 1.288533 | 66.73696 | 1.36E-05 | 0.001425 |
| TCEAL6 | 2.240896 | 1.535773 | 66.43669 | 1.36E-05 | 0.001425 |
| BMP8B | 2.182547 | 2.613317 | 66.69511 | 1.36E-05 | 0.001425 |
| GRIN3A | 3.633272 | 1.504652 | 66.69166 | 1.36E-05 | 0.001425 |
| SLC6A7 | 3.139086 | -0.09991 | 65.55116 | 1.41E-05 | 0.001462 |
| LIPH | 2.198187 | 0.85473 | 65.04507 | 1.42E-05 | 0.001462 |
| AC018804.1 | -1.67419 | 2.742732 | 65.01129 | 1.43E-05 | 0.001462 |
| GAD1 | 1.596769 | 6.242948 | 65.96434 | 1.43E-05 | 0.001462 |
| GRIPAP1 | 1.056667 | 6.761841 | 64.36764 | 1.49E-05 | 0.001512 |
| FZD10-DT | 3.646386 | 0.219476 | 65.2836 | 1.49E-05 | 0.001512 |
| NRP2 | 1.202826 | 6.806833 | 64.676 | 1.51E-05 | 0.001524 |
| TMEM185A | 1.463656 | 3.909479 | 63.79709 | 1.54E-05 | 0.001546 |
| INSM1 | 2.260052 | 2.487076 | 64.70753 | 1.55E-05 | 0.001546 |
| GNG12 | -1.55501 | 5.339978 | 64.50766 | 1.57E-05 | 0.001559 |
| GABRQ | 1.129139 | 7.35923 | 63.03064 | 1.62E-05 | 0.001597 |
| GATA3 | 2.506363 | 4.098911 | 63.94983 | 1.63E-05 | 0.001597 |
| AL117382.1 | -2.71245 | 4.357496 | 63.81342 | 1.64E-05 | 0.001597 |
| PTPRR | 1.650371 | 3.767133 | 63.7945 | 1.64E-05 | 0.001597 |
| GPR173 | 1.21303 | 7.439197 | 63.66607 | 1.65E-05 | 0.001597 |
| LINC01235 | 5.59791 | 0.567655 | 63.69692 | 1.65E-05 | 0.001597 |
| STK32A | -1.27487 | 6.927842 | 63.51588 | 1.67E-05 | 0.001608 |
| MAGEA4 | 7.493837 | -0.9413 | 91.41177 | 1.7E-05 | 0.001627 |
| AC073264.3 | -1.44842 | 3.344669 | 62.0787 | 1.73E-05 | 0.001649 |
| FAM90A11P | 8.160221 | -0.35315 | 93.03552 | 1.75E-05 | 0.001654 |
| MAMLD1 | 1.544531 | 5.027887 | 62.80463 | 1.75E-05 | 0.001655 |
| SYT9 | 2.353371 | 3.662039 | 62.53578 | 1.79E-05 | 0.001666 |
| TMEM185AP1 | 1.772588 | 2.197823 | 61.58705 | 1.79E-05 | 0.001666 |
| LHX5-AS1 | 3.225733 | -0.1307 | 62.45664 | 1.8E-05 | 0.001666 |
| PNMA5 | 2.27673 | 4.073058 | 62.36771 | 1.81E-05 | 0.001666 |
| TMEM132D | 1.972147 | 4.560668 | 62.3098 | 1.81E-05 | 0.001666 |
| ZCCHC12 | 1.290478 | 7.755464 | 62.26467 | 1.82E-05 | 0.001666 |
| MOSPD2 | 1.198214 | 5.377666 | 61.32319 | 1.82E-05 | 0.001666 |
| CACNA1I | 1.809038 | 3.464366 | 61.98143 | 1.85E-05 | 0.001679 |
| TBX2 | 3.021665 | 1.915753 | 61.97737 | 1.85E-05 | 0.001679 |
| PCDHB18P | 1.957115 | 1.565467 | 61.01865 | 1.86E-05 | 0.001679 |
| GPC3 | 2.688948 | 3.178673 | 61.653 | 1.89E-05 | 0.001701 |
| SIK1 | 1.421814 | 3.601228 | 60.60661 | 1.92E-05 | 0.001707 |
| IDH3G | 1.1108 | 6.07288 | 60.58936 | 1.92E-05 | 0.001707 |
| C1RL | -1.29965 | 3.755879 | 60.48605 | 1.93E-05 | 0.001712 |
| JADE3 | 1.24786 | 4.349999 | 60.2158 | 1.97E-05 | 0.001737 |

| | | | | | |
|---|---|---|---|---|---|
| SHE | 3.572975 | -0.52359 | 60.42436 | 2.05E-05 | 0.001799 |
| AC133561.1 | 2.734537 | 2.548824 | 60.22979 | 2.09E-05 | 0.001818 |
| RBPMS2 | 1.641634 | 3.35562 | 60.18309 | 2.09E-05 | 0.001818 |
| ZNF275 | 1.054172 | 6.265136 | 59.2661 | 2.1E-05 | 0.001818 |
| AC092070.2 | 1.927132 | 1.541165 | 59.2596 | 2.1E-05 | 0.001818 |
| ZNF135 | 1.721913 | 2.760114 | 59.55224 | 2.13E-05 | 0.001833 |
| C12orf56 | 2.517211 | -0.04489 | 59.02361 | 2.14E-05 | 0.001833 |
| KCNC3 | 1.59812 | 4.663249 | 59.79768 | 2.15E-05 | 0.001835 |
| DIRAS3 | 2.977126 | 4.184937 | 59.30088 | 2.23E-05 | 0.001892 |
| PAMR1 | -1.73586 | 3.079662 | 59.21756 | 2.24E-05 | 0.001895 |
| PCDHB10 | 1.177947 | 5.40382 | 58.29229 | 2.26E-05 | 0.0019 |
| GJB7 | 2.243735 | 1.407061 | 58.97925 | 2.28E-05 | 0.001911 |
| IFITM2 | -1.27105 | 3.947246 | 57.87991 | 2.32E-05 | 0.00194 |
| XIST | -4.03487 | -0.51857 | 58.64113 | 2.33E-05 | 0.001941 |
| HTR3B | 4.726723 | -0.99806 | 58.5392 | 2.35E-05 | 0.001947 |
| MPP1 | 1.895363 | 5.070487 | 58.39106 | 2.37E-05 | 0.00196 |
| ALX1 | -3.76767 | -0.17656 | 57.94304 | 2.45E-05 | 0.002015 |
| DAB2 | -2.39608 | 2.114924 | 57.81311 | 2.47E-05 | 0.002026 |
| PLAGL1 | -1.29429 | 4.28719 | 56.67754 | 2.53E-05 | 0.002067 |
| S100A11 | -1.59297 | 4.040098 | 57.27296 | 2.57E-05 | 0.002074 |
| AL356274.2 | 2.648416 | 0.355409 | 57.26461 | 2.57E-05 | 0.002074 |
| FGL1 | 2.544305 | 0.322734 | 56.78425 | 2.58E-05 | 0.002074 |
| NACA3P | 2.101794 | 2.538613 | 57.21048 | 2.58E-05 | 0.002074 |
| SLC6A8 | 1.39496 | 6.825917 | 57.13955 | 2.6E-05 | 0.002076 |
| AC060834.2 | 4.254908 | 0.38825 | 56.9685 | 2.63E-05 | 0.002093 |
| GPR176 | 1.428098 | 4.052431 | 56.91972 | 2.64E-05 | 0.002093 |
| PRKX | 1.102318 | 5.859756 | 56.03201 | 2.66E-05 | 0.002097 |
| SMIM32 | 4.371358 | -0.52817 | 56.78495 | 2.66E-05 | 0.002097 |
| UVSSA | -1.32054 | 5.35603 | 56.53596 | 2.71E-05 | 0.002128 |
| CD1D | 4.220235 | -0.39156 | 56.43925 | 2.73E-05 | 0.002134 |
| HERC3 | 1.45759 | 5.264406 | 56.26422 | 2.77E-05 | 0.002153 |
| NKX2-5 | 5.187803 | 2.327009 | 56.21653 | 2.78E-05 | 0.002153 |
| FLRT1 | 1.755994 | 2.877693 | 56.05992 | 2.81E-05 | 0.00217 |
| SLC6A10P | 3.950946 | -0.21369 | 55.93845 | 2.83E-05 | 0.002181 |
| ZNF571-AS1 | 1.610269 | 2.38516 | 55.03361 | 2.86E-05 | 0.002189 |
| SYTL5 | 1.270876 | 4.882191 | 55.59483 | 2.87E-05 | 0.002189 |
| C4A | -1.44686 | 5.242795 | 55.73686 | 2.88E-05 | 0.002189 |
| LINC02302 | 7.402665 | -1.01768 | 79.89278 | 2.9E-05 | 0.002196 |
| HTR2C | 3.10274 | 0.977619 | 55.46969 | 2.93E-05 | 0.00221 |
| RWDD2B | -1.20908 | 4.496299 | 54.66947 | 2.94E-05 | 0.00221 |
| CTGF | -1.15242 | 5.84967 | 55.2517 | 2.95E-05 | 0.00221 |
| H2AFJ | 1.13017 | 4.809386 | 54.60422 | 2.96E-05 | 0.00221 |
| RTL8B | 1.091322 | 5.387828 | 54.3847 | 3.01E-05 | 0.002239 |
| TPM3P9 | -1.40488 | 3.148375 | 54.32634 | 3.02E-05 | 0.002241 |
| LINC02303 | -3.60935 | 0.82457 | 55.02493 | 3.03E-05 | 0.002241 |
| NPAP1 | 2.626723 | -0.32661 | 54.1482 | 3.06E-05 | 0.002255 |
| CRYBA2 | 2.780326 | 1.954385 | 54.76549 | 3.09E-05 | 0.002263 |

| | | | | | |
|---|---|---|---|---|---|
| MVP | -1.41677 | 3.943999 | 54.68135 | 3.11E-05 | 0.002263 |
| TEX41 | 7.529246 | -0.90851 | 78.80365 | 3.11E-05 | 0.002263 |
| CASK | 1.166581 | 8.312961 | 54.60319 | 3.13E-05 | 0.002264 |
| RASSF6 | 3.384116 | -0.6825 | 54.17762 | 3.13E-05 | 0.002264 |
| CBLN2 | 2.4059 | 4.727446 | 54.5535 | 3.14E-05 | 0.002264 |
| ARHGDIB | 1.907249 | 2.526004 | 54.29087 | 3.2E-05 | 0.002297 |
| AC022148.1 | 3.607148 | -0.17643 | 54.21564 | 3.22E-05 | 0.002297 |
| IGSF10 | 1.587706 | 2.465265 | 53.48409 | 3.22E-05 | 0.002297 |
| MGAM | 2.43224 | 0.185412 | 53.44878 | 3.23E-05 | 0.002297 |
| IRX6 | -2.93142 | -0.15255 | 54.04029 | 3.27E-05 | 0.002313 |
| MIXL1 | 2.755903 | 0.072391 | 53.91333 | 3.3E-05 | 0.002327 |
| CHRM5 | 1.875497 | 1.595748 | 53.06732 | 3.33E-05 | 0.002341 |
| RNF113A | 1.21332 | 4.356568 | 53.02107 | 3.34E-05 | 0.002341 |
| PLXNA2 | 1.210363 | 5.744301 | 53.4031 | 3.43E-05 | 0.00239 |
| PRLHR | 3.740568 | 1.455212 | 53.38228 | 3.43E-05 | 0.00239 |
| IFITM3 | 1.467677 | 3.250604 | 52.74204 | 3.45E-05 | 0.00239 |
| IDS | 1.179239 | 10.24332 | 52.84307 | 3.46E-05 | 0.00239 |
| APOOL | 1.053955 | 5.694807 | 52.18835 | 3.56E-05 | 0.002457 |
| CPA6 | -2.30604 | 1.826279 | 52.66122 | 3.63E-05 | 0.002493 |
| AC079949.1 | 7.234783 | -1.15869 | 74.10222 | 3.64E-05 | 0.002493 |
| CRH | 2.995699 | 1.188248 | 52.45549 | 3.69E-05 | 0.002515 |
| ARMCX6 | 1.390106 | 3.987218 | 52.36025 | 3.71E-05 | 0.002515 |
| TNXA | -2.00118 | 3.02186 | 52.3204 | 3.73E-05 | 0.002515 |
| OTOL1 | 2.721454 | -0.34123 | 51.60173 | 3.73E-05 | 0.002515 |
| C1orf216 | 1.557361 | 5.770334 | 52.28973 | 3.74E-05 | 0.002515 |
| CLUL1 | 1.679063 | 3.137343 | 52.232 | 3.75E-05 | 0.002516 |
| COLEC11 | -2.9327 | 0.514349 | 52.20099 | 3.76E-05 | 0.002516 |
| C4B | -1.67086 | 3.396266 | 52.15443 | 3.77E-05 | 0.002517 |
| ST20 | -1.67116 | 3.220317 | 51.99466 | 3.82E-05 | 0.002541 |
| AL390334.1 | 2.335521 | 0.835412 | 51.94528 | 3.84E-05 | 0.002542 |
| CDKL5 | 1.105877 | 5.84531 | 51.59484 | 3.88E-05 | 0.002563 |
| ARHGAP4 | 1.552602 | 3.09006 | 51.7432 | 3.9E-05 | 0.002566 |
| KLF2P1 | -2.27833 | 0.084293 | 51.00364 | 3.92E-05 | 0.00257 |
| L1CAM | 1.726374 | 9.363325 | 51.52502 | 3.97E-05 | 0.00259 |
| ZXDB | 1.048186 | 5.653144 | 50.82728 | 3.97E-05 | 0.00259 |
| AC093912.1 | -5.02437 | -0.27987 | 51.44034 | 3.99E-05 | 0.00259 |
| FAM19A4 | 1.886 | 3.478939 | 51.38647 | 4.01E-05 | 0.00259 |
| BX842568.2 | 1.866904 | 0.884389 | 50.70791 | 4.01E-05 | 0.00259 |
| KIT | 1.760948 | 3.474289 | 51.16794 | 4.08E-05 | 0.002627 |
| RPL10 | 1.0111 | 8.147836 | 50.28733 | 4.15E-05 | 0.002656 |
| BX322639.1 | 1.999928 | 3.847437 | 50.9506 | 4.15E-05 | 0.002656 |
| SLC39A4 | 3.033075 | 0.680051 | 50.88113 | 4.17E-05 | 0.002662 |
| SOWAHD | 2.71232 | -0.35652 | 50.03108 | 4.24E-05 | 0.002694 |
| TP53TG3C | 7.021129 | -1.33623 | 70.19382 | 4.28E-05 | 0.002715 |
| PDZD4 | 1.315743 | 8.659678 | 50.3787 | 4.35E-05 | 0.002746 |
| CNGA3 | 1.372025 | 4.226838 | 50.31982 | 4.37E-05 | 0.00275 |
| SMIM10L2B | 1.292032 | 6.043969 | 50.2264 | 4.4E-05 | 0.002756 |

| | | | | | |
|---|---|---|---|---|---|
| BTBD17 | 2.323433 | 1.892549 | 50.20379 | 4.41E-05 | 0.002756 |
| SNRPN | 1.258323 | 7.636329 | 50.1822 | 4.42E-05 | 0.002756 |
| FAR2P4 | -2.10308 | 1.78731 | 50.02556 | 4.47E-05 | 0.002782 |
| AJ009632.2 | -3.22994 | 0.491422 | 49.89922 | 4.52E-05 | 0.002802 |
| PRRG1 | 1.322693 | 4.799931 | 49.74168 | 4.58E-05 | 0.002825 |
| LINC00648 | 1.772752 | 2.562046 | 49.7254 | 4.58E-05 | 0.002825 |
| CYP27A1 | 1.708671 | 2.523357 | 49.63199 | 4.62E-05 | 0.002832 |
| MTMR1 | 0.980557 | 5.656706 | 48.98316 | 4.62E-05 | 0.002832 |
| AC124312.2 | 2.104724 | 2.383845 | 49.56138 | 4.64E-05 | 0.002838 |
| CSAG3 | 7.472094 | -0.95973 | 69.72941 | 4.73E-05 | 0.00285 |
| NIPAL2 | -1.09592 | 5.945572 | 49.30929 | 4.74E-05 | 0.00285 |
| AC120036.4 | 1.631064 | 3.222042 | 49.26369 | 4.76E-05 | 0.00285 |
| EFNB1 | 1.530755 | 4.599001 | 49.2141 | 4.78E-05 | 0.00285 |
| C11orf88 | -7.57198 | -0.86889 | 69.50581 | 4.78E-05 | 0.00285 |
| MAGEL2 | 1.117412 | 7.270787 | 49.16005 | 4.8E-05 | 0.00285 |
| HSPA2 | 2.26995 | 0.684397 | 49.15083 | 4.8E-05 | 0.00285 |
| OR2W3 | 4.175152 | -0.69313 | 49.14476 | 4.8E-05 | 0.00285 |
| PLXNB3 | 2.002143 | 2.353035 | 49.14133 | 4.81E-05 | 0.00285 |
| FOXD2 | 3.120301 | 1.097003 | 49.12108 | 4.81E-05 | 0.00285 |
| Z68871.1 | 1.97173 | 0.731364 | 48.48616 | 4.82E-05 | 0.00285 |
| RADX | 1.865231 | 5.821627 | 49.04132 | 4.85E-05 | 0.002851 |
| AC108865.2 | 4.267204 | -0.93966 | 49.03799 | 4.85E-05 | 0.002851 |
| RAB11FIP1 | 1.386543 | 3.151718 | 48.27234 | 4.9E-05 | 0.002876 |
| HERC2P5 | 2.903536 | 0.326483 | 48.7813 | 4.95E-05 | 0.002896 |
| GFY | 2.51525 | -0.50156 | 48.10625 | 4.97E-05 | 0.0029 |
| GABRB2 | 3.894396 | 2.138549 | 48.65877 | 5E-05 | 0.002909 |
| TPH1 | 2.438425 | 2.99361 | 48.56107 | 5.04E-05 | 0.002924 |
| SYNDIG1L | 2.591358 | 0.551717 | 48.365 | 5.12E-05 | 0.002964 |
| MUC5AC | -1.22609 | 9.328902 | 48.32585 | 5.14E-05 | 0.002965 |
| EHHADH | 2.014442 | 1.876263 | 48.27794 | 5.16E-05 | 0.002969 |
| AF228730.5 | 1.657383 | 2.134163 | 47.7572 | 5.22E-05 | 0.002993 |
| TAC3 | 2.031492 | 3.647162 | 48.02432 | 5.27E-05 | 0.003004 |
| HTR1D | 1.79801 | 2.122021 | 47.97669 | 5.29E-05 | 0.003004 |
| PNPLA4 | 1.246081 | 5.031547 | 47.97669 | 5.29E-05 | 0.003004 |
| PNCK | 1.499853 | 6.318871 | 47.96943 | 5.3E-05 | 0.003004 |
| RHOJ | -1.96874 | 5.292479 | 47.7894 | 5.38E-05 | 0.003041 |
| PKP1 | 4.12975 | -0.47197 | 47.70797 | 5.41E-05 | 0.003054 |
| USP27X | 0.997227 | 5.559376 | 46.92347 | 5.5E-05 | 0.003094 |
| DDX11 | 1.548609 | 3.473803 | 47.42869 | 5.54E-05 | 0.00311 |
| CDH23 | 1.478542 | 2.864923 | 47.02001 | 5.58E-05 | 0.00312 |
| GPR149 | 3.406583 | -0.08887 | 47.27293 | 5.62E-05 | 0.003134 |
| AMHR2 | 3.4058 | -1.05607 | 46.63824 | 5.64E-05 | 0.003138 |
| GPC4 | 1.613619 | 4.017394 | 47.10254 | 5.7E-05 | 0.003155 |
| PCDHGB7 | 2.374983 | 2.521882 | 47.09965 | 5.7E-05 | 0.003155 |
| CENPF | 1.629079 | 1.572457 | 46.32044 | 5.8E-05 | 0.003198 |
| SMC1A | 0.889954 | 6.667016 | 46.25965 | 5.83E-05 | 0.003207 |
| CASC8 | 3.049414 | -0.08743 | 46.69576 | 5.9E-05 | 0.003239 |

| | | | | | |
|---|---|---|---|---|---|
| KIRREL3 | 1.58289 | 3.347378 | 46.6404 | 5.93E-05 | 0.00324 |
| PWAR5 | 1.412265 | 2.821817 | 46.01262 | 5.95E-05 | 0.00324 |
| TXNRD2 | -1.37759 | 2.485344 | 45.91267 | 6.01E-05 | 0.00324 |
| FAM90A12P | 7.047033 | -1.31392 | 64.70188 | 6.02E-05 | 0.00324 |
| SLC9A3 | 1.872633 | 0.738475 | 45.87853 | 6.02E-05 | 0.00324 |
| CLCN4 | 1.22228 | 8.051785 | 46.45153 | 6.03E-05 | 0.00324 |
| AC003973.3 | 7.594805 | -0.8612 | 64.93229 | 6.03E-05 | 0.00324 |
| MKRN3 | 1.325434 | 3.970578 | 46.44738 | 6.03E-05 | 0.00324 |
| SLC9A7 | 1.088444 | 7.729249 | 46.3098 | 6.1E-05 | 0.00327 |
| SLCO1B1 | 5.753044 | -0.80939 | 46.18722 | 6.17E-05 | 0.003293 |
| RLN2 | 2.430546 | -0.39769 | 45.59498 | 6.18E-05 | 0.003293 |
| GRB10 | 0.938881 | 6.902814 | 45.54145 | 6.21E-05 | 0.0033 |
| HRK | 1.423441 | 4.693581 | 45.95132 | 6.29E-05 | 0.003335 |
| FBLN5 | -1.52526 | 3.934742 | 45.93234 | 6.3E-05 | 0.003335 |
| GUCA1B | 2.628113 | 0.434666 | 45.82082 | 6.37E-05 | 0.003359 |
| AIF1L | 2.244732 | 0.836224 | 45.57847 | 6.5E-05 | 0.003422 |
| DCAF12L2 | 1.36545 | 3.353216 | 45.31632 | 6.53E-05 | 0.00343 |
| FAM161B | 1.065047 | 4.358963 | 44.92212 | 6.56E-05 | 0.003434 |
| PAX2 | 3.09282 | -0.60665 | 45.35278 | 6.63E-05 | 0.003459 |
| ALLC | -2.53209 | -0.22631 | 45.00526 | 6.64E-05 | 0.003459 |
| ATP7B | 1.163225 | 4.014099 | 44.62717 | 6.73E-05 | 0.003499 |
| C9orf64 | -3.36364 | -0.3601 | 45.119 | 6.77E-05 | 0.003503 |
| PAPPA | 1.742472 | 3.549495 | 45.085 | 6.79E-05 | 0.003503 |
| SFRP1 | 1.293606 | 5.720475 | 45.07166 | 6.8E-05 | 0.003503 |
| EOMES | 2.713383 | -0.76722 | 44.50252 | 6.81E-05 | 0.003503 |
| SPATA18 | 1.894111 | 1.807222 | 45.00835 | 6.84E-05 | 0.003508 |
| HOXD9 | 7.278743 | -1.12906 | 62.44221 | 6.88E-05 | 0.003523 |
| CYP21A2 | -2.05995 | 1.579404 | 44.82041 | 6.95E-05 | 0.003546 |
| NHSL2 | 1.200917 | 6.182024 | 44.79357 | 6.97E-05 | 0.003546 |
| SLC5A7 | 0.908425 | 6.258985 | 44.22808 | 6.98E-05 | 0.003546 |
| AL021395.1 | 7.77509 | -0.69485 | 62.09394 | 7.01E-05 | 0.003555 |
| LMCD1 | -1.3263 | 5.13277 | 44.67504 | 7.04E-05 | 0.00356 |
| LINC00649 | 1.868684 | 3.428488 | 44.63257 | 7.07E-05 | 0.00356 |
| SEC14L5 | 1.836766 | 1.039381 | 44.07608 | 7.08E-05 | 0.00356 |
| Z93403.1 | 3.103442 | -0.73584 | 44.5398 | 7.13E-05 | 0.003577 |
| AC025588.1 | 2.209723 | 0.495097 | 44.45672 | 7.18E-05 | 0.003595 |
| SEMA5A | 1.23415 | 5.086952 | 44.41832 | 7.21E-05 | 0.003598 |
| PKIB | 1.470049 | 6.503243 | 44.38275 | 7.23E-05 | 0.003601 |
| DOCK11 | 1.090445 | 4.569695 | 43.80467 | 7.26E-05 | 0.003605 |
| EPHB6 | 2.097258 | 3.306846 | 44.25761 | 7.31E-05 | 0.003624 |
| AGTR1 | 3.636816 | 1.294531 | 44.17223 | 7.37E-05 | 0.003643 |
| APOL4 | -1.62808 | 3.909277 | 44.13626 | 7.39E-05 | 0.003644 |
| CD44 | -1.38985 | 4.794055 | 44.11587 | 7.41E-05 | 0.003644 |
| MRC2 | -1.27105 | 7.095644 | 43.89884 | 7.55E-05 | 0.003707 |
| Z82214.2 | -3.97434 | 1.008307 | 43.86655 | 7.57E-05 | 0.003709 |
| IL17RA | 2.342489 | 4.100743 | 43.76972 | 7.64E-05 | 0.003729 |
| AC007368.1 | 6.789595 | -1.51795 | 59.28382 | 7.65E-05 | 0.003729 |

| | | | | | |
|---|---|---|---|---|---|
| MGAM2 | 3.14444 | -0.86982 | 43.57082 | 7.68E-05 | 0.003732 |
| CDH6 | -1.15059 | 6.765555 | 43.54807 | 7.8E-05 | 0.003782 |
| AC009495.3 | -3.98559 | 1.588003 | 43.42529 | 7.88E-05 | 0.003807 |
| METTL11B | 2.36835 | -0.61334 | 42.90253 | 7.88E-05 | 0.003807 |
| AC097639.1 | -2.89346 | 5.415734 | 43.35678 | 7.93E-05 | 0.003821 |
| KDM5C | 1.255664 | 7.119368 | 43.30365 | 7.97E-05 | 0.003825 |
| TM9SF4 | 0.999395 | 6.657467 | 43.26535 | 7.98E-05 | 0.003825 |
| CCKAR | 3.540007 | -0.95317 | 43.22647 | 8.03E-05 | 0.00384 |
| PRLR | 1.774101 | 2.907084 | 43.19493 | 8.05E-05 | 0.003842 |
| FAM156B | 0.840382 | 6.457752 | 42.6368 | 8.08E-05 | 0.003844 |
| ASXL1 | 0.930031 | 7.234632 | 42.62451 | 8.09E-05 | 0.003844 |
| TSPYL2 | 1.081987 | 7.146534 | 43.02093 | 8.18E-05 | 0.003872 |
| MKI67 | 2.055969 | 0.222064 | 42.48682 | 8.2E-05 | 0.003872 |
| AC004233.2 | -5.70936 | 3.738165 | 42.98421 | 8.21E-05 | 0.003872 |
| VSTM2A | 1.097618 | 6.094696 | 42.93821 | 8.24E-05 | 0.003875 |
| TSPEAR-AS1 | 1.603928 | 1.268808 | 42.41345 | 8.25E-05 | 0.003875 |
| HLA-DQA2 | -3.64669 | 1.354807 | 42.73688 | 8.4E-05 | 0.003934 |
| XIAP | 1.039362 | 6.991437 | 42.67057 | 8.45E-05 | 0.00395 |
| LRP2 | 1.550886 | 5.294394 | 42.60359 | 8.5E-05 | 0.003961 |
| CDK16 | 1.250006 | 8.242649 | 42.5898 | 8.51E-05 | 0.003961 |
| CTHRC1 | 1.70917 | 1.247579 | 41.99228 | 8.59E-05 | 0.003987 |
| ATP1A2 | 2.021218 | 3.460505 | 42.38335 | 8.68E-05 | 0.00402 |
| TMSB15A | 1.160428 | 5.750082 | 42.27296 | 8.77E-05 | 0.00405 |
| CHMP4C | 2.412059 | -0.19271 | 41.95407 | 8.8E-05 | 0.00405 |
| ARHGAP36 | 1.290936 | 9.056353 | 42.22788 | 8.81E-05 | 0.00405 |
| GALNT3 | -3.01765 | 5.520411 | 42.18555 | 8.84E-05 | 0.00405 |
| MICB | 2.33292 | -0.12583 | 41.75952 | 8.88E-05 | 0.00405 |
| ITGA2 | 1.50803 | 2.882834 | 42.13458 | 8.88E-05 | 0.00405 |
| HLA-G | 1.563523 | 1.527402 | 41.61053 | 8.91E-05 | 0.00405 |
| CXorf40A | 1.028291 | 5.29049 | 41.80574 | 8.92E-05 | 0.00405 |
| IL18 | -2.62578 | 1.22323 | 42.0867 | 8.92E-05 | 0.00405 |
| FAHD1 | -0.90894 | 5.481232 | 41.54503 | 8.96E-05 | 0.004059 |
| C14orf39 | 4.331644 | -0.06244 | 41.98278 | 9.01E-05 | 0.004072 |
| TCTEX1D1 | -2.04291 | 3.03186 | 41.92831 | 9.06E-05 | 0.004084 |
| HEPACAM | 2.816415 | -1.11463 | 41.36673 | 9.12E-05 | 0.004101 |
| XDH | 3.743831 | 0.318229 | 41.83071 | 9.14E-05 | 0.004103 |
| GAB3 | 3.243102 | -0.35743 | 41.79942 | 9.17E-05 | 0.004106 |
| CCDC169 | 1.425604 | 2.433854 | 41.25539 | 9.21E-05 | 0.004118 |
| CCNQ | 1.047683 | 5.112793 | 41.34439 | 9.33E-05 | 0.004153 |
| PTH1R | 1.967324 | 2.911191 | 41.5964 | 9.34E-05 | 0.004153 |
| GDI1 | 0.98358 | 10.20546 | 41.10087 | 9.35E-05 | 0.004153 |
| AL031595.2 | 1.851331 | 1.016699 | 41.29274 | 9.38E-05 | 0.004154 |
| RNASEL | 1.085531 | 4.377059 | 40.96476 | 9.47E-05 | 0.004189 |
| MET | 1.687808 | 4.729252 | 41.39311 | 9.53E-05 | 0.004203 |
| CHST7 | 1.862129 | 1.818596 | 41.23321 | 9.67E-05 | 0.004258 |
| MAGEA2B | 6.969465 | -1.38125 | 56.38061 | 9.71E-05 | 0.004261 |
| IGSF11 | 1.510544 | 1.320818 | 40.65538 | 9.76E-05 | 0.004261 |

| | | | | | |
|---|---|---|---|---|---|
| XKR7 | 1.359394 | 5.847619 | 41.12681 | 9.77E-05 | 0.004261 |
| BCL6 | -1.09173 | 5.942236 | 41.11093 | 9.79E-05 | 0.004261 |
| AL136226.1 | -3.71985 | -1.03768 | 41.09507 | 9.8E-05 | 0.004261 |
| ANO2 | 1.905049 | 1.253356 | 41.08513 | 9.81E-05 | 0.004261 |
| NMRAL1 | 1.227802 | 2.910259 | 40.59141 | 9.82E-05 | 0.004261 |
| PWAR6 | 1.211957 | 8.13859 | 41.03795 | 9.85E-05 | 0.004265 |
| QPRT | -0.95155 | 5.925385 | 40.58262 | 9.92E-05 | 0.004285 |
| DSG2 | -1.16713 | 4.821732 | 40.91769 | 9.97E-05 | 0.004296 |
| ST18 | 2.066098 | 2.677833 | 40.86983 | 0.0001 | 0.0043 |
| QPCT | 1.246048 | 5.721161 | 40.86557 | 0.0001 | 0.0043 |
| POU4F2 | 7.277664 | -1.12021 | 55.80256 | 0.0001 | 0.004303 |
| ATP6AP1 | 1.117131 | 8.514341 | 40.71295 | 0.000102 | 0.004342 |
| NLGN4X | 0.820864 | 7.213622 | 40.22056 | 0.000102 | 0.004342 |
| USP9X | 0.857924 | 8.175388 | 40.20687 | 0.000102 | 0.004342 |
| SLC30A8 | 2.514292 | 0.95883 | 40.58939 | 0.000103 | 0.00437 |
| S100A6 | 1.097715 | 4.797826 | 40.40251 | 0.000104 | 0.00442 |
| PLXNC1 | 1.397667 | 5.026361 | 40.34039 | 0.000105 | 0.004458 |
| ANKS1B | 1.222862 | 5.1229 | 40.12053 | 0.000108 | 0.004545 |
| COL23A1 | 1.652233 | 2.889286 | 40.08462 | 0.000108 | 0.004552 |
| ARHGAP6 | 1.460903 | 3.166886 | 40.03813 | 0.000109 | 0.004563 |
| GLRA2 | 1.163412 | 7.675048 | 39.99499 | 0.000109 | 0.004565 |
| CXorf40B | 1.101757 | 4.077809 | 39.53129 | 0.000109 | 0.004565 |
| KANTR | 1.034793 | 6.52591 | 39.93381 | 0.00011 | 0.004574 |
| ATP2B3 | 1.490422 | 4.880539 | 39.92949 | 0.00011 | 0.004574 |
| PQBP1 | 0.855711 | 5.867499 | 39.32093 | 0.000111 | 0.004634 |
| AC022150.3 | 6.525089 | -1.7213 | 52.96469 | 0.000112 | 0.004653 |
| RENBP | 1.20765 | 4.202844 | 39.52571 | 0.000114 | 0.00473 |
| TRAF1 | -1.645 | 2.606764 | 39.48533 | 0.000115 | 0.00474 |
| ZNF503-AS2 | -1.2398 | 2.361182 | 38.98974 | 0.000115 | 0.004743 |
| EYA1 | 1.853627 | 3.414907 | 39.43827 | 0.000115 | 0.004743 |
| CCDC34 | -1.0518 | 4.616159 | 38.89535 | 0.000116 | 0.00476 |
| EFCAB2 | -1.67901 | 4.167579 | 39.32369 | 0.000116 | 0.00476 |
| PIEZO2 | 1.310879 | 3.570882 | 39.31662 | 0.000117 | 0.00476 |
| JADE1 | -1.33025 | 5.937882 | 39.29972 | 0.000117 | 0.00476 |
| SLC7A14 | 1.306171 | 5.406958 | 39.29318 | 0.000117 | 0.00476 |
| SLC9A6 | 1.174598 | 8.555544 | 39.27143 | 0.000117 | 0.00476 |
| FHL1 | 1.112121 | 9.089745 | 39.26094 | 0.000117 | 0.00476 |
| FCRLA | 3.887257 | 0.423683 | 39.19458 | 0.000118 | 0.004773 |
| COL9A3 | -2.47132 | 3.996332 | 39.19218 | 0.000118 | 0.004773 |
| GRP | 2.404718 | 0.615979 | 39.12614 | 0.000119 | 0.004786 |
| KCNQ1 | 2.260133 | 1.212057 | 39.11707 | 0.000119 | 0.004786 |
| CLEC3B | -2.48557 | 0.614741 | 39.10527 | 0.000119 | 0.004786 |
| DUSP15 | 1.356483 | 4.447495 | 39.01608 | 0.00012 | 0.00482 |
| AL591848.4 | 1.31514 | 2.662233 | 38.46596 | 0.000121 | 0.004861 |
| COMT | -1.53814 | 5.126131 | 38.89146 | 0.000122 | 0.004861 |
| ZNF578 | 3.89333 | -0.25139 | 38.74552 | 0.000123 | 0.004923 |
| CSMD1 | 1.935471 | 2.923833 | 38.72318 | 0.000124 | 0.004924 |

| | | | | | |
|---|---|---|---|---|---|
| DNM3OS | -3.85632 | 3.18679 | 38.69134 | 0.000124 | 0.004926 |
| LRIG3 | -1.12005 | 5.715444 | 38.68112 | 0.000124 | 0.004926 |
| NIPAL1 | 1.705164 | 1.157484 | 38.30965 | 0.000124 | 0.004927 |
| CFAP58-DT | 2.533928 | 0.678676 | 38.57746 | 0.000125 | 0.004959 |
| OSBPL10 | -1.61155 | 5.816732 | 38.55649 | 0.000126 | 0.004959 |
| MED15P9 | -2.37305 | 0.607522 | 38.46894 | 0.000127 | 0.004994 |
| TRHDE | 1.444874 | 3.968585 | 38.42082 | 0.000127 | 0.005009 |
| IFI35 | -1.98606 | 1.098024 | 38.39617 | 0.000128 | 0.005011 |
| AC018688.1 | 2.14221 | -0.04168 | 38.00007 | 0.000128 | 0.005017 |
| LINC02145 | -2.25854 | -0.35098 | 37.91691 | 0.000128 | 0.005017 |
| AC104692.1 | -4.70893 | -1.63121 | 38.32447 | 0.000129 | 0.005019 |
| BCAP31 | 0.947124 | 7.558746 | 38.27852 | 0.000129 | 0.005033 |
| NDRG1 | 0.975134 | 6.117217 | 38.25581 | 0.00013 | 0.005035 |
| SMN2 | -0.92953 | 6.700504 | 38.14453 | 0.00013 | 0.005052 |
| PRKY | 1.557745 | 1.567212 | 37.69725 | 0.000131 | 0.005084 |
| NLRP2 | 1.792674 | 3.191664 | 38.08407 | 0.000132 | 0.005095 |
| HTR1E | 2.857702 | -0.33192 | 38.0378 | 0.000133 | 0.005109 |
| SH3GL3 | 1.09805 | 4.810621 | 38.00936 | 0.000133 | 0.005109 |
| UBQLN2 | 0.864863 | 7.76008 | 37.55818 | 0.000133 | 0.005109 |
| ELN | -1.16479 | 6.569053 | 37.97195 | 0.000133 | 0.005109 |
| ITGB4 | -1.97449 | 2.045026 | 37.96396 | 0.000134 | 0.005109 |
| AC023034.2 | 1.983378 | 0.284561 | 37.55928 | 0.000134 | 0.005127 |
| MKRN7P | -1.65304 | 1.955119 | 37.8924 | 0.000134 | 0.005127 |
| FAM78B | 1.279639 | 2.943702 | 37.52393 | 0.000135 | 0.005128 |
| TRPM2 | 2.961121 | -0.45751 | 37.85402 | 0.000135 | 0.005128 |
| SYNPO2 | 1.144176 | 3.274801 | 37.39196 | 0.000136 | 0.005132 |
| RPS2P32 | 2.129249 | 0.440482 | 37.81019 | 0.000136 | 0.005132 |
| NHLRC1 | -2.6451 | 0.013082 | 37.73905 | 0.000137 | 0.00516 |
| PNMA6F | 1.533794 | 2.972342 | 37.68566 | 0.000137 | 0.005179 |
| ABCB1 | 1.989781 | 3.397058 | 37.60459 | 0.000139 | 0.005213 |
| SPAG6 | -2.63253 | 3.418057 | 37.52437 | 0.00014 | 0.005247 |
| SRPX | 1.437792 | 3.417384 | 37.341 | 0.000142 | 0.005337 |
| NOV | 1.290197 | 3.876493 | 37.29217 | 0.000143 | 0.00535 |
| FAM66E | 3.045284 | 0.626554 | 37.28261 | 0.000143 | 0.00535 |
| KCNJ5 | 1.84985 | 1.591204 | 37.10386 | 0.000146 | 0.00544 |
| ZBTB16 | 2.238838 | 1.635529 | 37.06732 | 0.000146 | 0.005442 |
| AC016582.1 | 3.25976 | -1.41109 | 36.65755 | 0.000146 | 0.005442 |
| ZNF737 | 1.309939 | 3.036747 | 36.92232 | 0.000149 | 0.005514 |
| PEX5L | 1.753668 | 3.123589 | 36.85649 | 0.00015 | 0.005542 |
| KANSL1-AS1 | 1.99437 | 0.916983 | 36.82799 | 0.00015 | 0.005548 |
| C1QL3 | 2.15794 | 0.63143 | 36.75251 | 0.000151 | 0.005582 |
| ZDHHC11 | -1.44975 | 5.027741 | 36.6956 | 0.000152 | 0.005606 |
| CHGB | 1.36856 | 8.950367 | 36.61943 | 0.000153 | 0.00564 |
| TMEM273 | 2.83783 | -0.56333 | 36.57979 | 0.000154 | 0.005652 |
| FAM218A | 1.765975 | 2.758017 | 36.56617 | 0.000154 | 0.005652 |
| MYH7 | 1.399304 | 2.309818 | 36.26923 | 0.000156 | 0.005685 |
| APOL2 | -0.95797 | 5.180979 | 36.08231 | 0.000157 | 0.00573 |

| | | | | | |
|---|---|---|---|---|---|
| ADRA2A | 1.586325 | 2.477249 | 36.34344 | 0.000158 | 0.005747 |
| GCOM1 | 1.716431 | 2.161588 | 36.34032 | 0.000158 | 0.005747 |
| PRKCB | 1.351781 | 3.832264 | 36.28722 | 0.000159 | 0.005769 |
| ZNF502 | 1.327113 | 3.314468 | 36.23289 | 0.00016 | 0.00579 |
| AC009403.2 | 2.822411 | 0.838112 | 36.21926 | 0.00016 | 0.00579 |
| AC074281.2 | 7.417321 | -1.0024 | 48.3707 | 0.000161 | 0.00583 |
| ENTPD5 | 1.063184 | 3.705627 | 35.64669 | 0.000163 | 0.005883 |
| AC108865.1 | 3.473418 | -0.03236 | 35.9866 | 0.000164 | 0.005904 |
| AC132807.2 | 2.905984 | -0.51119 | 35.95948 | 0.000165 | 0.005907 |
| ARHGAP25 | 1.398315 | 1.700112 | 35.55983 | 0.000165 | 0.005907 |
| NOL4L | 0.836791 | 7.740265 | 35.52623 | 0.000165 | 0.005918 |
| AC009163.6 | 1.800888 | 1.000246 | 35.82127 | 0.000167 | 0.005967 |
| PID1 | 1.401657 | 4.626376 | 35.80512 | 0.000167 | 0.005967 |
| MAP3K13 | 1.104683 | 5.315013 | 35.78693 | 0.000168 | 0.005968 |
| SLC18A1 | 1.429469 | 2.099633 | 35.54435 | 0.000168 | 0.005973 |
| LDOC1 | 1.013021 | 7.201496 | 35.73105 | 0.000169 | 0.005983 |
| SNHG14 | 0.90534 | 8.738894 | 35.44572 | 0.000169 | 0.005985 |
| AC135050.3 | -1.98545 | 0.411176 | 35.68236 | 0.00017 | 0.005985 |
| PIRT | 1.472467 | 2.202456 | 35.6798 | 0.00017 | 0.005985 |
| IRX1 | 1.676713 | 1.856188 | 35.61754 | 0.000171 | 0.006015 |
| FGL2 | -3.36678 | 0.461471 | 35.51131 | 0.000173 | 0.006067 |
| CCL25 | -2.1567 | -0.66345 | 35.12419 | 0.000173 | 0.006067 |
| BLCAP | -0.92314 | 8.42254 | 35.44082 | 0.000173 | 0.006074 |
| UBE3A | -0.89168 | 7.346533 | 35.39675 | 0.000174 | 0.006077 |
| HOXA9 | -2.93657 | 4.026369 | 35.44372 | 0.000174 | 0.006077 |
| AF165147.1 | 2.790593 | -0.70521 | 35.40518 | 0.000175 | 0.006077 |
| HS6ST3 | 1.097931 | 6.5058 | 35.40261 | 0.000175 | 0.006077 |
| NOTUM | 2.432252 | 0.707997 | 35.39545 | 0.000175 | 0.006077 |
| OTUD5 | 0.848159 | 6.95748 | 34.88697 | 0.000177 | 0.006155 |
| LINC02082 | -1.79874 | 0.342097 | 34.83963 | 0.000178 | 0.006175 |
| TBC1D10C | 2.347484 | -0.92415 | 34.73714 | 0.00018 | 0.006226 |
| WARS2 | -1.20974 | 3.815507 | 35.10989 | 0.00018 | 0.006226 |
| SATB2 | -1.9708 | 5.20076 | 35.04018 | 0.000182 | 0.006263 |
| CILP | -1.24247 | 6.940581 | 35.00046 | 0.000183 | 0.006279 |
| DGKG | 2.511408 | 2.383486 | 34.93534 | 0.000184 | 0.006314 |
| FGF13 | 0.920369 | 8.659895 | 34.84429 | 0.000185 | 0.006329 |
| TBX3 | 4.420994 | -0.21261 | 34.87424 | 0.000185 | 0.006329 |
| VAX2 | 3.031156 | 2.976496 | 34.8668 | 0.000185 | 0.006329 |
| MAPRE1 | 0.845476 | 8.263319 | 34.44235 | 0.000186 | 0.006353 |
| EMD | 0.926774 | 6.430289 | 34.79876 | 0.000187 | 0.006353 |
| PAPSS2 | -1.36745 | 5.841462 | 34.77574 | 0.000187 | 0.006353 |
| AC093809.1 | -2.22865 | -0.00223 | 34.7721 | 0.000187 | 0.006353 |
| PCDHA7 | 1.954093 | 0.335223 | 34.67987 | 0.000189 | 0.006398 |
| LINC01505 | 2.820728 | -0.09408 | 34.61969 | 0.00019 | 0.006439 |
| C5orf38 | 1.843976 | 2.250731 | 34.54961 | 0.000192 | 0.006478 |
| AC136616.1 | 2.708353 | -0.36105 | 34.4883 | 0.000193 | 0.006499 |
| KCNC4 | 1.228338 | 6.02231 | 34.4816 | 0.000193 | 0.006499 |

| | | | | | |
|---|---|---|---|---|---|
| MIR22HG | -0.89101 | 4.622812 | 34.11125 | 0.000193 | 0.006499 |
| CAV2 | 2.024112 | 2.686383 | 34.45687 | 0.000194 | 0.006501 |
| GEMIN8 | 1.005189 | 4.385016 | 34.04157 | 0.000195 | 0.00653 |
| ATP11C | 1.082303 | 5.630074 | 34.36214 | 0.000196 | 0.006548 |
| CDHR3 | -1.38976 | 6.552069 | 34.31601 | 0.000197 | 0.006563 |
| AL121757.1 | 4.299724 | -1.34007 | 34.29458 | 0.000197 | 0.006563 |
| ASB2 | 1.510405 | 3.011989 | 34.29246 | 0.000197 | 0.006563 |
| WDR66 | 1.104295 | 2.808107 | 33.91786 | 0.000198 | 0.006563 |
| CPVL | -1.3528 | 5.065886 | 34.23995 | 0.000198 | 0.006563 |
| RAB11FIP1P1 | 2.647939 | -0.01527 | 34.23942 | 0.000198 | 0.006563 |
| SPANXN3 | 2.917553 | -1.0383 | 34.23915 | 0.000198 | 0.006563 |
| AC006042.3 | 2.748534 | 0.148142 | 34.15981 | 0.0002 | 0.006611 |
| ESYT3 | 1.491458 | 3.371412 | 34.08706 | 0.000202 | 0.006654 |
| SLC5A12 | 1.774426 | 1.156282 | 34.05952 | 0.000202 | 0.006663 |
| ADGRG2 | 2.079572 | 2.532618 | 34.02334 | 0.000203 | 0.00668 |
| PLEKHD1 | 1.765164 | 2.199422 | 33.99463 | 0.000204 | 0.006688 |
| AC096677.1 | 2.526822 | 0.1546 | 33.98348 | 0.000204 | 0.006688 |
| MICAL2 | 1.392422 | 4.430053 | 33.95962 | 0.000205 | 0.006695 |
| SERPINH1 | -1.27224 | 6.82199 | 33.88141 | 0.000207 | 0.006743 |
| RBFADN | -1.55175 | 1.696411 | 33.8581 | 0.000207 | 0.00675 |
| HS3ST4 | -1.20923 | 5.104569 | 33.83492 | 0.000208 | 0.006751 |
| AL592183.1 | 1.576864 | 1.772756 | 33.8282 | 0.000208 | 0.006751 |
| WLS | -1.50664 | 7.140645 | 33.73883 | 0.00021 | 0.006808 |
| PLSCR4 | -2.30441 | 2.130734 | 33.71201 | 0.00021 | 0.006818 |
| G6PD | 1.193629 | 5.436951 | 33.69404 | 0.000211 | 0.006821 |
| KCTD8 | 1.318521 | 2.912929 | 33.6463 | 0.000212 | 0.006845 |
| ZFP36L2 | -1.74096 | 6.694792 | 33.63501 | 0.000212 | 0.006845 |
| UBL4A | 1.056912 | 6.00584 | 33.599 | 0.000213 | 0.006862 |
| SLC2A1 | 1.449066 | 6.614626 | 33.57953 | 0.000214 | 0.006866 |
| FAM81B | -2.35047 | -0.2203 | 33.45065 | 0.000217 | 0.006956 |
| GLB1L3 | 1.75803 | 0.458514 | 33.04259 | 0.000219 | 0.006973 |
| ASCL1 | 2.890492 | 2.36802 | 33.37511 | 0.000219 | 0.006973 |
| GCHFR | 1.521815 | 0.870127 | 33.01006 | 0.000219 | 0.006973 |
| UBA1 | 1.031301 | 9.280553 | 33.34562 | 0.000219 | 0.006973 |
| PLXDC1 | 1.296444 | 3.730017 | 33.33347 | 0.00022 | 0.006973 |
| FAM163B | 1.768386 | 2.365766 | 33.33234 | 0.00022 | 0.006973 |
| VSTM2B | -1.49594 | 2.432347 | 33.32867 | 0.00022 | 0.006973 |
| MID1IP1 | 1.030714 | 5.574713 | 33.31856 | 0.00022 | 0.006973 |
| TRPM8 | 2.387976 | 1.337684 | 33.28597 | 0.000221 | 0.006988 |
| CDH22 | 1.291386 | 4.120977 | 33.25278 | 0.000222 | 0.006995 |
| H19 | -1.82323 | 8.517608 | 33.24298 | 0.000222 | 0.006995 |
| TSPY26P | 0.865519 | 5.191195 | 32.89302 | 0.000222 | 0.006995 |
| SDK2 | 1.035979 | 5.738834 | 33.17415 | 0.000224 | 0.007015 |
| COL11A2 | -1.2518 | 3.816739 | 33.15796 | 0.000224 | 0.007015 |
| WDR72 | 2.054959 | 1.461283 | 33.15676 | 0.000224 | 0.007015 |
| CACNA2D2 | 1.04304 | 6.168752 | 33.12766 | 0.000225 | 0.007015 |
| APCDD1L | 2.476044 | 0.583458 | 33.1271 | 0.000225 | 0.007015 |

| | | | | | |
|---|---|---|---|---|---|
| EN1 | 1.54068 | 6.07417 | 33.125 | 0.000225 | 0.007015 |
| DNM1P51 | -1.54813 | 0.872943 | 32.77579 | 0.000225 | 0.007015 |
| MMP23A | -1.9271 | 1.090396 | 33.01004 | 0.000228 | 0.007091 |
| HK2P1 | 2.670692 | 1.729938 | 32.9955 | 0.000228 | 0.007092 |
| RTP5 | 1.523327 | 3.855803 | 32.80187 | 0.000233 | 0.00724 |
| COL1A2 | 2.460043 | -0.21897 | 32.70099 | 0.000236 | 0.007313 |
| COL8A1 | 2.431167 | 3.028606 | 32.6549 | 0.000237 | 0.007337 |
| ELK1 | 1.22896 | 4.753689 | 32.64639 | 0.000238 | 0.007337 |
| NECAB2 | 0.945879 | 6.072586 | 32.61619 | 0.000238 | 0.007344 |
| HCN4 | 1.292909 | 4.791497 | 32.61253 | 0.000239 | 0.007344 |
| BMS1P8 | 1.196206 | 2.779161 | 32.25852 | 0.000239 | 0.00735 |
| BCL2L1 | 1.006786 | 6.463574 | 32.48359 | 0.000242 | 0.007419 |
| C5orf64 | 2.150402 | -0.1476 | 32.47592 | 0.000242 | 0.007419 |
| HBD | 4.470635 | -1.20261 | 32.47317 | 0.000242 | 0.007419 |
| TPX2 | 1.436896 | 2.899779 | 32.3676 | 0.000245 | 0.007499 |
| CD40 | 2.640014 | 0.282587 | 32.32765 | 0.000247 | 0.007522 |
| SLC16A4 | -1.65195 | 2.955318 | 32.2375 | 0.000249 | 0.00759 |
| DKC1 | 0.815159 | 5.975328 | 31.88959 | 0.00025 | 0.007597 |
| ITPKA | 2.360034 | 0.890374 | 32.18182 | 0.000251 | 0.007611 |
| ALK | 1.476072 | 4.504524 | 32.16485 | 0.000251 | 0.007611 |
| TNMD | 3.420837 | -1.29179 | 32.16308 | 0.000251 | 0.007611 |
| CNGB1 | 2.009575 | 1.658203 | 32.10233 | 0.000253 | 0.007654 |
| AP001178.3 | 1.624105 | 1.620242 | 32.08093 | 0.000254 | 0.007662 |
| HTR1B | 4.618265 | -0.30809 | 31.98823 | 0.000257 | 0.007734 |
| ADAM15 | 1.112685 | 4.186075 | 31.93539 | 0.000258 | 0.007771 |
| EFCAB13 | 1.612388 | 1.205904 | 31.86632 | 0.00026 | 0.007823 |
| NSMCE1 | 0.915565 | 4.679571 | 31.51903 | 0.000261 | 0.007835 |
| FOXD2-AS1 | 2.370287 | 1.167915 | 31.79417 | 0.000262 | 0.007866 |
| COL4A2 | 1.222379 | 5.356598 | 31.75774 | 0.000264 | 0.007888 |
| AC006206.2 | 2.026506 | -0.41227 | 31.39282 | 0.000265 | 0.007919 |
| XYLT1 | 0.962 | 4.240883 | 31.3379 | 0.000267 | 0.00794 |
| LINC02300 | 2.362673 | -0.70708 | 31.50951 | 0.000267 | 0.00794 |
| DCLK3 | 1.497003 | 2.069916 | 31.65307 | 0.000267 | 0.00794 |
| TRABD2B | -1.90646 | 5.430302 | 31.63205 | 0.000268 | 0.007948 |
| EDA | 1.153122 | 3.893588 | 31.47458 | 0.000273 | 0.008086 |
| CDH13 | -1.06734 | 4.863745 | 31.45339 | 0.000273 | 0.008095 |
| RPS6KA3 | 1.07084 | 7.553208 | 31.40804 | 0.000275 | 0.008127 |
| AL035425.3 | 3.153559 | -0.69694 | 31.34885 | 0.000277 | 0.008171 |
| NEK9 | -1.26884 | 6.655065 | 31.32199 | 0.000278 | 0.008171 |
| NXPH2 | 2.240288 | 0.079085 | 31.31916 | 0.000278 | 0.008171 |
| CER1 | 2.633015 | -0.96003 | 31.30461 | 0.000278 | 0.008171 |
| PDLIM1 | -1.51279 | 4.170707 | 31.30192 | 0.000278 | 0.008171 |
| GSTT2B | -1.39077 | 1.613503 | 30.90637 | 0.000281 | 0.008241 |
| AC112206.3 | 1.873293 | 0.457475 | 31.19254 | 0.000282 | 0.008255 |
| Z83843.1 | 1.133625 | 5.617155 | 31.17691 | 0.000282 | 0.008258 |
| ABCD1 | 1.508955 | 5.305737 | 31.13655 | 0.000284 | 0.008287 |
| OR2L2 | 2.563994 | -0.07982 | 31.10824 | 0.000285 | 0.008303 |

| | | | | | |
|---|---|---|---|---|---|
| MIR9-3HG | 1.211413 | 2.141346 | 30.75248 | 0.000286 | 0.008337 |
| SPIN3 | 1.036337 | 6.029653 | 31.03725 | 0.000287 | 0.008343 |
| TIGD2 | 1.071822 | 3.326063 | 30.7194 | 0.000287 | 0.008343 |
| PHF24 | 0.865359 | 6.067939 | 31.02095 | 0.000288 | 0.008343 |
| KLKB1 | -1.4494 | 1.67691 | 30.89667 | 0.000291 | 0.008414 |
| PAQR6 | -1.19392 | 3.748271 | 30.86703 | 0.000293 | 0.008475 |
| PLXNA3 | 0.973359 | 7.360684 | 30.84002 | 0.000294 | 0.00848 |
| THSD4-AS1 | 2.558058 | -0.88364 | 30.83837 | 0.000294 | 0.00848 |
| GJD2 | 3.436276 | -0.63565 | 30.81281 | 0.000295 | 0.008494 |
| CHRNB4 | 1.340861 | 3.997839 | 30.77857 | 0.000296 | 0.008503 |
| AL590814.1 | 3.669352 | -1.41142 | 30.77294 | 0.000296 | 0.008503 |
| CACHD1 | 0.825852 | 5.649076 | 30.46491 | 0.000297 | 0.008503 |
| MMP23B | -2.14355 | -0.19023 | 30.74473 | 0.000297 | 0.008516 |
| AGBL2 | -3.65039 | -0.11181 | 30.70668 | 0.000299 | 0.00854 |
| PGRMC1 | 0.843759 | 8.901396 | 30.49164 | 0.000299 | 0.00854 |
| AC239585.2 | 1.72942 | 0.610631 | 30.67497 | 0.0003 | 0.008553 |
| PCDHGC4 | 2.854985 | 0.519703 | 30.62063 | 0.000302 | 0.008591 |
| MED12L | 1.198147 | 2.753328 | 30.44448 | 0.000302 | 0.008591 |
| MYO3A | 2.394106 | 0.356739 | 30.57321 | 0.000304 | 0.008618 |
| SCDP1 | 1.901413 | 2.349385 | 30.55682 | 0.000304 | 0.008618 |
| MXRA5 | 1.949916 | 1.846878 | 30.55565 | 0.000304 | 0.008618 |
| BHLHB9 | 0.959449 | 5.347878 | 30.49605 | 0.000307 | 0.008668 |
| TNFSF10 | 3.538094 | 0.031321 | 30.44869 | 0.000308 | 0.008691 |
| PHYHD1 | -1.92995 | 0.270809 | 30.43538 | 0.000309 | 0.008691 |
| ADI1 | -1.05265 | 4.96481 | 30.42525 | 0.000309 | 0.008691 |
| IL1RAPL1 | 1.40998 | 1.232037 | 30.10769 | 0.00031 | 0.008691 |
| EGFL6 | 1.450724 | 3.503635 | 30.40408 | 0.00031 | 0.008691 |
| AC004836.1 | -2.14027 | 4.64911 | 30.39672 | 0.00031 | 0.008691 |
| MMGT1 | 0.82088 | 7.057789 | 30.34314 | 0.00031 | 0.008691 |
| GABRA3 | 1.087357 | 6.901895 | 30.37284 | 0.000311 | 0.008703 |
| AL513318.2 | 0.924753 | 4.533198 | 29.96154 | 0.000316 | 0.008814 |
| MYO18B | 2.352659 | -1.03362 | 29.89048 | 0.000318 | 0.00888 |
| CRNDE | -1.13571 | 4.516912 | 30.13407 | 0.00032 | 0.008925 |
| FAM201A | 1.362147 | 1.27099 | 29.81193 | 0.000322 | 0.008941 |
| COL6A4P2 | -1.09162 | 3.377022 | 29.94492 | 0.000322 | 0.008941 |
| B3GNT7 | -1.46078 | 3.041789 | 30.06909 | 0.000323 | 0.00896 |
| ICAM5 | 3.227926 | -0.03891 | 29.98241 | 0.000326 | 0.00904 |
| HRC | -1.37271 | 2.847679 | 29.97363 | 0.000327 | 0.00904 |
| SCN4A | 2.231636 | 0.838236 | 29.96354 | 0.000327 | 0.00904 |
| PNOC | 1.23173 | 2.504724 | 29.76197 | 0.000329 | 0.009082 |
| LINC01135 | -2.46837 | -0.61441 | 29.90121 | 0.00033 | 0.009082 |
| CD27-AS1 | -0.98227 | 4.011404 | 29.65881 | 0.00033 | 0.009082 |
| CACNA2D3 | 1.45288 | 3.668663 | 29.83348 | 0.000333 | 0.009138 |
| MR1 | -1.36193 | 2.193445 | 29.81365 | 0.000333 | 0.009148 |
| PDCD6IPP2 | -2.08334 | 1.438518 | 29.79935 | 0.000334 | 0.009152 |
| GPC6 | -1.39021 | 5.357346 | 29.76422 | 0.000335 | 0.009164 |
| SIM2 | -1.4309 | 4.315992 | 29.7585 | 0.000336 | 0.009164 |

| | | | | | |
|---|---|---|---|---|---|
| FRMPD4 | 2.079155 | 2.899669 | 29.75589 | 0.000336 | 0.009164 |
| SH3TC1 | 1.851568 | 0.193286 | 29.71691 | 0.000337 | 0.009197 |
| MYLK2 | 2.112341 | -0.29231 | 29.68962 | 0.000339 | 0.009216 |
| NECTIN3 | -1.5961 | 7.385955 | 29.658 | 0.00034 | 0.00924 |
| MRI1 | 1.433584 | 2.799576 | 29.64272 | 0.00034 | 0.009245 |
| MAL2 | 1.230967 | 5.0354 | 29.59952 | 0.000342 | 0.009282 |
| ZNF528 | 1.346296 | 3.764724 | 29.40728 | 0.000351 | 0.009484 |
| IQGAP3 | 2.44497 | -0.97163 | 29.25053 | 0.000351 | 0.009484 |
| WIF1 | 2.513327 | 0.964153 | 29.38084 | 0.000352 | 0.009502 |
| MYRF | 1.528808 | 2.572707 | 29.28434 | 0.000356 | 0.009605 |
| LRRIQ3 | 1.522103 | 0.849897 | 28.91112 | 0.00036 | 0.009708 |
| GALNT5 | -1.50822 | 0.817332 | 28.8849 | 0.000362 | 0.009725 |
| TRIM4 | -1.27967 | 3.84803 | 29.10976 | 0.000364 | 0.009768 |
| TOP2A | 1.811096 | 1.459865 | 29.10928 | 0.000364 | 0.009768 |
| ARMC3 | -1.86119 | 2.372836 | 29.08925 | 0.000365 | 0.009768 |
| ZNF83 | -1.05208 | 6.612091 | 29.0883 | 0.000365 | 0.009768 |
| AC097478.1 | 1.851837 | -0.42134 | 28.79917 | 0.000365 | 0.009768 |
| LRIG1 | -1.16634 | 6.08905 | 29.05397 | 0.000367 | 0.009785 |
| ZNF704 | 0.928392 | 6.410865 | 28.99936 | 0.000369 | 0.00984 |
| GABRA4 | 3.782475 | -0.74809 | 28.98402 | 0.00037 | 0.009845 |
| RAPH1 | 1.407366 | 2.497027 | 28.96681 | 0.000371 | 0.009845 |
| PCDHGB5 | 2.914984 | -0.39043 | 28.95186 | 0.000371 | 0.009845 |
| GSTT2 | -1.19389 | 3.541148 | 28.94957 | 0.000371 | 0.009845 |
| NOC2LP1 | 1.227178 | 2.967152 | 28.94366 | 0.000372 | 0.009845 |
| FOXQ1 | 3.971371 | -1.17545 | 28.91202 | 0.000373 | 0.009872 |
| HOXA5 | -1.26033 | 4.509714 | 28.82126 | 0.000378 | 0.009974 |
| AL121949.1 | 2.64913 | -1.40979 | 28.53179 | 0.000378 | 0.00998 |
| KCNA1 | -1.79977 | 0.743924 | 28.7569 | 0.000381 | 0.01003 |
| HIST1H3E | -1.902 | 2.745071 | 28.71955 | 0.000382 | 0.010058 |
| CDC42BPB | 1.040098 | 6.541885 | 28.71518 | 0.000383 | 0.010058 |
| GLIS3 | -1.3912 | 5.498528 | 28.70511 | 0.000383 | 0.010058 |
| AL157778.1 | 1.188577 | 2.511966 | 28.4879 | 0.000385 | 0.010079 |
| MASP1 | 0.993974 | 3.630656 | 28.3589 | 0.000387 | 0.010119 |
| ADAMTSL3 | 1.593693 | 1.555163 | 28.62236 | 0.000387 | 0.010119 |
| TBX20 | 4.335775 | -0.07868 | 28.61716 | 0.000388 | 0.010119 |
| FHIT | 0.971844 | 3.003486 | 28.32435 | 0.000389 | 0.010125 |
| KITLG | 1.050488 | 5.538412 | 28.59298 | 0.000389 | 0.010125 |
| CALHM5 | 1.518009 | 4.283919 | 28.58156 | 0.000389 | 0.010127 |
| SNU13 | 0.707413 | 7.852018 | 28.27561 | 0.000391 | 0.010151 |
| COL19A1 | 1.941527 | 0.001594 | 28.54278 | 0.000391 | 0.010151 |
| RPS20P14 | -1.10957 | 4.445652 | 28.5221 | 0.000392 | 0.010155 |
| MYO5B | 1.421457 | 2.213902 | 28.51754 | 0.000393 | 0.010155 |
| EIF4E3 | 1.200274 | 5.163979 | 28.51041 | 0.000393 | 0.010155 |
| CLCN3P1 | 1.988191 | -0.25891 | 28.47074 | 0.000394 | 0.010155 |
| DPP7 | -1.10499 | 5.50089 | 28.48829 | 0.000394 | 0.010155 |
| SLC9A7P1 | 1.708892 | 0.505279 | 28.48125 | 0.000394 | 0.010155 |
| FGF10 | 2.932851 | 0.411356 | 28.41307 | 0.000398 | 0.010222 |

| | | | | | |
|---|---|---|---|---|---|
| SMOC1 | -1.79075 | 8.802415 | 28.40993 | 0.000398 | 0.010222 |
| AC110285.1 | 2.222172 | 0.122934 | 28.39751 | 0.000399 | 0.010226 |
| ADCY3 | 1.229064 | 3.631617 | 28.3574 | 0.000401 | 0.010245 |
| H3F3B | -0.87507 | 8.453422 | 28.35446 | 0.000401 | 0.010245 |
| CD99L2 | 0.760779 | 7.862123 | 28.08643 | 0.000401 | 0.010245 |
| AC234775.3 | 1.906524 | 1.23561 | 28.31106 | 0.000403 | 0.010289 |
| SHROOM4 | 1.244795 | 3.11985 | 28.25372 | 0.000406 | 0.010352 |
| ACAA2 | -0.993 | 5.461231 | 28.22525 | 0.000408 | 0.010356 |
| TMEM176A | 1.695354 | 4.190465 | 28.21414 | 0.000408 | 0.010356 |
| LAMA2 | 1.416709 | 2.094253 | 28.21342 | 0.000408 | 0.010356 |
| SEPT6 | 1.029206 | 8.246128 | 28.21248 | 0.000408 | 0.010356 |
| SYN1 | 1.005941 | 8.123154 | 28.19582 | 0.000409 | 0.010366 |
| TSPYL5 | 2.623266 | 1.051951 | 28.15166 | 0.000411 | 0.010404 |
| ARF4 | -0.86267 | 8.618568 | 28.14789 | 0.000412 | 0.010404 |
| PPFIA4 | 1.193029 | 5.913757 | 28.13026 | 0.000413 | 0.010415 |
| NUPR1 | -1.479 | 3.208421 | 28.11685 | 0.000413 | 0.010417 |
| NR4A2 | -1.93683 | 2.271371 | 28.10999 | 0.000414 | 0.010417 |
| AC027279.1 | -1.88376 | 0.717826 | 28.0721 | 0.000416 | 0.010443 |
| AC021054.1 | -1.7599 | 1.888394 | 28.07179 | 0.000416 | 0.010443 |
| GPR27 | 0.85443 | 5.419872 | 28.00654 | 0.000419 | 0.010519 |
| LAMP2 | 0.86625 | 7.713639 | 27.97847 | 0.000421 | 0.010545 |
| WDR38 | -3.05703 | -0.13728 | 27.95491 | 0.000422 | 0.010564 |
| DDN | 1.603698 | 2.924534 | 27.89893 | 0.000425 | 0.010629 |
| PTPRN2 | 0.897671 | 7.349493 | 27.88283 | 0.000426 | 0.010634 |
| CALB2 | 0.981921 | 4.776389 | 27.86036 | 0.000427 | 0.010634 |
| AC017076.1 | 1.580512 | 1.087193 | 27.85772 | 0.000427 | 0.010634 |
| TRPV3 | 1.623235 | 0.135421 | 27.59364 | 0.000428 | 0.010634 |
| CU633906.1 | -2.2747 | -0.96156 | 27.58868 | 0.000428 | 0.010634 |
| HOXB2 | -1.57725 | 5.029144 | 27.82311 | 0.000429 | 0.010654 |
| ADRA1D | 1.703537 | 1.876777 | 27.8152 | 0.00043 | 0.010654 |
| AP003063.1 | 2.094712 | -0.90143 | 27.50874 | 0.000433 | 0.010694 |
| SAP30L | 0.73766 | 5.58604 | 27.50562 | 0.000433 | 0.010694 |
| RIPPLY3 | -2.11622 | 4.311753 | 27.75946 | 0.000433 | 0.010694 |
| SLC25A5 | 0.730786 | 7.126867 | 27.47491 | 0.000435 | 0.010711 |
| AL109614.1 | 2.811632 | -0.82081 | 27.72883 | 0.000435 | 0.010711 |
| ARMCX5-GPRASP2 | 0.950978 | 3.64668 | 27.45395 | 0.000436 | 0.010718 |
| DNAH2 | -1.68906 | 2.879736 | 27.70519 | 0.000436 | 0.010718 |
| AC120036.3 | 2.235318 | -0.90372 | 27.44155 | 0.000438 | 0.01074 |
| FAM3A | 1.009341 | 6.127047 | 27.67124 | 0.000438 | 0.01074 |
| CAPN9 | 4.206236 | -1.00289 | 27.61548 | 0.000441 | 0.010806 |
| VIM | -1.27446 | 10.77843 | 27.50927 | 0.000448 | 0.010945 |
| HDC | 1.817813 | 0.000507 | 27.42269 | 0.00045 | 0.010987 |
| FAM181B | 1.37987 | 2.235013 | 27.44088 | 0.000452 | 0.011018 |
| GSTM1 | 1.41499 | 1.015183 | 27.16909 | 0.000453 | 0.011031 |
| NPNT | 1.889112 | 4.341266 | 27.40332 | 0.000454 | 0.011047 |
| MARCH11 | 1.164829 | 2.459864 | 27.18493 | 0.000455 | 0.011054 |
| LINC00698 | 1.978998 | -0.65821 | 27.10417 | 0.000457 | 0.011088 |

| | | | | | |
|---|---|---|---|---|---|
| LINC01561 | 1.690225 | -0.47437 | 27.08791 | 0.000458 | 0.0111 |
| CASQ1 | 1.302608 | 1.087761 | 27.05123 | 0.00046 | 0.011141 |
| APTR | -0.90446 | 3.998692 | 27.03949 | 0.000461 | 0.011142 |
| RNASEH1-AS1 | 0.984592 | 2.786012 | 27.03259 | 0.000461 | 0.011142 |
| FMN1 | 1.085005 | 3.556513 | 27.27573 | 0.000462 | 0.011142 |
| UVRAG | 0.748292 | 5.238368 | 26.99332 | 0.000463 | 0.011176 |
| AL672207.1 | 1.417237 | 2.396507 | 27.21146 | 0.000466 | 0.011212 |
| WDR44 | 0.886303 | 6.083967 | 27.19718 | 0.000466 | 0.01122 |
| ZNF681 | 1.278695 | 3.009044 | 27.15662 | 0.000469 | 0.011267 |
| RPS6KA6 | 1.030912 | 5.970076 | 27.13126 | 0.000471 | 0.011275 |
| SUPT16HP1 | 1.399009 | 1.479827 | 27.1128 | 0.000471 | 0.011275 |
| WDR78 | -1.35272 | 4.362179 | 27.12044 | 0.000471 | 0.011275 |
| SPIN2B | 0.889392 | 4.703764 | 27.00327 | 0.000471 | 0.011275 |
| MECOM | -1.46058 | 1.177773 | 27.03166 | 0.000475 | 0.011348 |
| KLHL4 | 1.709409 | 3.575799 | 27.04722 | 0.000476 | 0.011348 |
| MARVELD1 | -1.18141 | 4.18144 | 27.04182 | 0.000476 | 0.011348 |
| GPRIN3 | 0.862236 | 6.885786 | 26.96102 | 0.000481 | 0.011458 |
| RAI14 | -1.1062 | 6.791132 | 26.92869 | 0.000483 | 0.011494 |
| DYNLT3P2 | 2.254097 | -0.99276 | 26.66511 | 0.000485 | 0.011512 |
| VWA3A | -2.41691 | 2.333476 | 26.88785 | 0.000486 | 0.011531 |
| ADGRG4 | 2.134894 | 2.096157 | 26.85339 | 0.000488 | 0.011559 |
| AIF1 | -5.24004 | -0.55191 | 26.85204 | 0.000488 | 0.011559 |
| LHX9 | 2.239546 | 2.129901 | 26.81973 | 0.000491 | 0.011592 |
| AC092111.1 | 1.88266 | 2.070821 | 26.81389 | 0.000491 | 0.011592 |
| SGCD | 1.759851 | 1.596933 | 26.80069 | 0.000492 | 0.011599 |
| SERPINE2 | 0.757246 | 5.972525 | 26.43441 | 0.0005 | 0.011777 |
| CCDC162P | -1.71738 | 4.88698 | 26.66774 | 0.000501 | 0.011777 |
| CDKN2B | 1.104583 | 3.013105 | 26.64622 | 0.000501 | 0.011777 |
| ANO4 | 1.44627 | 1.788415 | 26.65347 | 0.000502 | 0.011777 |
| NBDY | 0.656353 | 6.505695 | 26.38439 | 0.000503 | 0.011805 |
| GPRASP2 | 0.955205 | 7.212907 | 26.61935 | 0.000504 | 0.011805 |
| GLDCP1 | 1.445533 | 1.153942 | 26.53528 | 0.000507 | 0.011873 |
| NAAA | 1.225831 | 1.656574 | 26.31383 | 0.000508 | 0.011881 |
| LRRC34 | -1.21987 | 1.746547 | 26.21158 | 0.000516 | 0.012036 |
| IGFBPL1 | 1.288843 | 3.321297 | 26.40682 | 0.000519 | 0.012073 |
| BRINP2 | 1.213928 | 2.794 | 26.40619 | 0.000519 | 0.012073 |
| PRDM16 | 1.966773 | -0.28553 | 26.40346 | 0.000519 | 0.012073 |
| FAM9A | 3.874391 | -0.09922 | 26.38435 | 0.00052 | 0.01209 |
| ANGPTL1 | -2.19186 | 6.700464 | 26.35421 | 0.000522 | 0.012123 |
| SCN5A | 1.293122 | 1.871124 | 26.29685 | 0.000524 | 0.012123 |
| CMTM8 | -1.88575 | 6.272357 | 26.3331 | 0.000524 | 0.012123 |
| AC084357.3 | 3.837905 | -1.28767 | 26.33145 | 0.000524 | 0.012123 |
| AC004233.3 | 1.626062 | 1.957312 | 26.32182 | 0.000525 | 0.012125 |
| TBC1D2 | 1.166938 | 1.841041 | 26.0732 | 0.000526 | 0.01213 |
| THNSL2 | 1.792676 | 3.27146 | 26.29007 | 0.000527 | 0.01215 |
| ZNF747 | -0.94523 | 3.186368 | 26.03754 | 0.000528 | 0.012163 |
| SLC35E1P1 | 2.694205 | -1.20426 | 26.21948 | 0.000532 | 0.012238 |

| | | | | | |
|---|---|---|---|---|---|
| TCF4 | -1.4292 | 6.085301 | 26.2127 | 0.000533 | 0.012238 |
| DUSP23 | 1.288336 | 2.883115 | 26.18628 | 0.000535 | 0.012265 |
| AC009159.3 | -1.60119 | 0.296081 | 25.98271 | 0.000535 | 0.012265 |
| HOXC4 | 2.300918 | -0.4276 | 26.15102 | 0.000537 | 0.012301 |
| ARNTL | 0.840066 | 3.803531 | 25.89202 | 0.000539 | 0.012328 |
| BTBD19 | -1.42913 | 2.894246 | 26.1019 | 0.000541 | 0.012356 |
| KCNE4 | 2.342066 | 4.085923 | 26.09022 | 0.000542 | 0.012362 |
| GUCA1A | 1.574747 | -0.00538 | 25.79968 | 0.000546 | 0.012446 |
| PLPP4 | 1.492902 | 0.412966 | 25.79053 | 0.000547 | 0.012448 |
| LINC01139 | 2.755 | -0.22348 | 25.99669 | 0.000549 | 0.012481 |
| AC091078.1 | -1.43154 | 2.217722 | 25.98502 | 0.00055 | 0.012487 |
| CFAP43 | -1.82791 | 3.65116 | 25.95906 | 0.000552 | 0.012508 |
| ZFX | -0.86978 | 5.71403 | 25.95032 | 0.000552 | 0.012508 |
| MYO3B | 1.737202 | -0.01242 | 25.81672 | 0.000553 | 0.012508 |
| TAF13 | -0.85359 | 5.661483 | 25.93287 | 0.000554 | 0.012508 |
| HSPA8P7 | 1.456614 | 1.529891 | 25.93252 | 0.000554 | 0.012508 |
| HIST1H2BC | -2.7766 | -1.29221 | 25.91733 | 0.000555 | 0.012511 |
| DANT2 | 1.967098 | -0.81899 | 25.68312 | 0.000555 | 0.012511 |
| MMD2 | 3.001157 | 0.143141 | 25.88347 | 0.000557 | 0.012551 |
| TBL1Y | 1.799469 | -0.51428 | 25.6031 | 0.000561 | 0.012613 |
| SLC15A1 | -2.65469 | 1.531791 | 25.82038 | 0.000562 | 0.012613 |
| WASF3 | 0.758128 | 6.114855 | 25.68876 | 0.000562 | 0.012613 |
| RD3 | 2.03305 | 0.505668 | 25.81611 | 0.000563 | 0.012613 |
| AC008280.3 | -1.95378 | 0.925902 | 25.71411 | 0.000571 | 0.012779 |
| CCDC80 | -1.99549 | 5.974623 | 25.6854 | 0.000573 | 0.012816 |
| PKD1P5 | 1.833701 | 1.480466 | 25.66245 | 0.000575 | 0.012843 |
| PYGL | -1.31578 | 4.394137 | 25.61787 | 0.000578 | 0.012887 |
| RPH3A | 1.398937 | 2.153092 | 25.61559 | 0.000578 | 0.012887 |
| COL5A1 | -1.10898 | 5.173625 | 25.60104 | 0.00058 | 0.012887 |
| ELMO1 | -1.37771 | 6.893297 | 25.59764 | 0.00058 | 0.012887 |
| BNC2 | 1.244345 | 2.068859 | 25.58567 | 0.00058 | 0.012887 |
| PALM2 | 1.512743 | 3.531317 | 25.58699 | 0.000581 | 0.012887 |
| PHKA2 | 1.151254 | 4.768544 | 25.58348 | 0.000581 | 0.012887 |
| PRAME | -2.22111 | -0.71016 | 25.5673 | 0.000582 | 0.012894 |
| NEU4 | -1.94798 | -0.10707 | 25.5637 | 0.000583 | 0.012894 |
| AGBL4 | 1.377336 | 2.798816 | 25.54046 | 0.000584 | 0.012922 |
| AC138356.3 | -3.82719 | 0.154523 | 25.52015 | 0.000586 | 0.012945 |
| SNHG5 | -1.02162 | 4.166043 | 25.45587 | 0.000591 | 0.013036 |
| KLK4 | 2.32642 | -0.01801 | 25.45458 | 0.000592 | 0.013036 |
| GLDN | 1.41141 | 3.124539 | 25.42427 | 0.000594 | 0.013077 |
| SLC17A7 | 2.015921 | 2.879569 | 25.39056 | 0.000597 | 0.013125 |
| CYP27C1 | -1.01538 | 5.171361 | 25.35774 | 0.0006 | 0.013161 |
| PNMA3 | 1.148574 | 6.3249 | 25.35592 | 0.0006 | 0.013161 |
| PLEKHG4B | -0.9785 | 3.184376 | 25.10342 | 0.000602 | 0.0132 |
| KLF8 | 1.329359 | 2.21456 | 25.30949 | 0.000604 | 0.013213 |
| C10orf82 | 1.314444 | 2.181392 | 25.30504 | 0.000604 | 0.013213 |
| SSR4 | 0.854258 | 7.02869 | 25.28024 | 0.000606 | 0.013244 |

| | | | | | |
|---|---|---|---|---|---|
| ATP11A | 0.745565 | 5.425071 | 25.05035 | 0.000607 | 0.013244 |
| ADAMTSL1 | -1.5465 | 6.282896 | 25.26541 | 0.000607 | 0.013244 |
| FUNDC2 | 0.744754 | 6.427989 | 25.10102 | 0.000611 | 0.013301 |
| DNER | 0.997057 | 7.534455 | 25.2198 | 0.000611 | 0.013301 |
| CLK3P2 | 2.887435 | -0.91312 | 25.19586 | 0.000613 | 0.013326 |
| ATP6AP2 | 0.782164 | 8.550625 | 25.18205 | 0.000614 | 0.013326 |
| LPA | -1.71742 | 0.978725 | 25.12701 | 0.000619 | 0.013412 |
| LRBA | -1.19738 | 4.601282 | 25.12066 | 0.00062 | 0.013412 |
| GALC | 0.926767 | 5.421576 | 25.12025 | 0.00062 | 0.013412 |
| HSD17B10 | 0.927142 | 5.631215 | 25.1159 | 0.00062 | 0.013412 |
| STS | 0.968842 | 5.706733 | 25.10623 | 0.000621 | 0.013416 |
| ZNF418 | 1.241347 | 2.778611 | 25.06841 | 0.000625 | 0.013468 |
| CNTNAP5 | 1.456667 | 3.201689 | 25.05775 | 0.000626 | 0.013468 |
| CHSY3 | 1.372963 | 1.852516 | 25.05658 | 0.000626 | 0.013468 |
| KRT222 | -2.0166 | 2.14762 | 25.04379 | 0.000627 | 0.013469 |
| PDGFA | -1.01642 | 2.312395 | 24.81842 | 0.000627 | 0.013469 |
| SVIL-AS1 | 1.650695 | 3.705715 | 25.03386 | 0.000628 | 0.013469 |
| NKX2-2 | -1.85205 | 1.645722 | 25.01757 | 0.000629 | 0.013486 |
| ITGA5 | -1.92467 | 2.843644 | 24.98283 | 0.000632 | 0.013539 |
| RSPO4 | -1.21154 | 2.168008 | 24.95276 | 0.000635 | 0.013582 |
| PKD1P6 | 1.434046 | 1.221249 | 24.84395 | 0.000645 | 0.013772 |
| FRMD4B | -0.84821 | 4.074347 | 24.62404 | 0.000645 | 0.013772 |
| BCAP31P1 | 1.035395 | 3.863118 | 24.78322 | 0.00065 | 0.013871 |
| AC007663.2 | -2.57921 | -0.97838 | 24.77181 | 0.000652 | 0.013879 |
| CCDC120 | 1.106048 | 4.799553 | 24.73294 | 0.000655 | 0.013942 |
| ZNF266 | -1.05415 | 5.001445 | 24.72492 | 0.000656 | 0.013943 |
| HOTAIRM1 | -1.04399 | 2.862119 | 24.51194 | 0.00066 | 0.013985 |
| SHROOM2 | 0.867289 | 6.615252 | 24.67572 | 0.000661 | 0.013985 |
| BCAT1 | 0.852487 | 6.553291 | 24.6756 | 0.000661 | 0.013985 |
| C1orf115 | 1.536973 | 1.734954 | 24.67517 | 0.000661 | 0.013985 |
| WIPF1 | -0.74966 | 5.113484 | 24.43872 | 0.000663 | 0.014015 |
| CCM2L | 1.958555 | 1.102674 | 24.64345 | 0.000664 | 0.014019 |
| PPEF1 | 1.581542 | 1.2268 | 24.62477 | 0.000665 | 0.014043 |
| GPR156 | 1.955583 | 0.793065 | 24.58794 | 0.000669 | 0.014103 |
| NUAK2 | 1.195356 | 2.076078 | 24.48656 | 0.000671 | 0.014129 |
| HIST1H2BN | -1.30324 | 1.077822 | 24.33742 | 0.000673 | 0.014151 |
| PLA2G4C | 1.041197 | 3.298002 | 24.5347 | 0.000674 | 0.014155 |
| CNTN3 | 1.027263 | 4.156261 | 24.53174 | 0.000674 | 0.014155 |
| CSAD | -1.2713 | 4.424578 | 24.52627 | 0.000675 | 0.014155 |
| DHCR24 | 0.844093 | 7.531065 | 24.51949 | 0.000676 | 0.014155 |
| SAXO1 | -2.18129 | 1.376758 | 24.50053 | 0.000677 | 0.014179 |
| GJA1 | -0.8992 | 4.712118 | 24.45736 | 0.000682 | 0.014253 |
| UST | 1.031985 | 3.548266 | 24.43847 | 0.000683 | 0.014257 |
| PRDM6 | -2.35106 | 2.163015 | 24.43204 | 0.000684 | 0.014257 |
| IQGAP1 | -0.87971 | 6.006582 | 24.43194 | 0.000684 | 0.014257 |
| TLX2 | 2.650377 | -1.60047 | 24.21072 | 0.000685 | 0.014257 |
| ROBO3 | 1.105815 | 2.929923 | 24.41834 | 0.000685 | 0.014257 |

| | | | | | |
|---|---|---|---|---|---|
| BRCC3 | 0.883941 | 5.314525 | 24.40549 | 0.000687 | 0.014257 |
| SLAMF9 | 1.680943 | 0.304346 | 24.40229 | 0.000687 | 0.014257 |
| COL24A1 | 2.545712 | 3.009718 | 24.39354 | 0.000688 | 0.014257 |
| SMARCC2 | 0.923479 | 6.46931 | 24.38625 | 0.000689 | 0.014257 |
| VMA21 | 0.761622 | 7.49096 | 24.3848 | 0.000689 | 0.014257 |
| CRYBB2P1 | 0.832556 | 4.14304 | 24.16548 | 0.00069 | 0.014262 |
| HPR | -2.59114 | -0.25665 | 24.35589 | 0.000692 | 0.01427 |
| RAB26 | 1.32054 | 2.813444 | 24.34573 | 0.000693 | 0.01427 |
| ARRB1 | 1.183586 | 4.997351 | 24.34022 | 0.000693 | 0.01427 |
| TRIP6 | -1.24616 | 2.928233 | 24.33568 | 0.000694 | 0.01427 |
| PDGFRA | -4.34051 | 2.950574 | 24.33078 | 0.000694 | 0.01427 |
| ANKRD34B | 2.04868 | 1.31346 | 24.32347 | 0.000695 | 0.01427 |
| WWC1 | 0.919296 | 5.458206 | 24.32302 | 0.000695 | 0.01427 |
| PRSS16 | -1.21259 | 1.944863 | 24.21651 | 0.000696 | 0.014279 |
| GPR146 | 3.223648 | -1.43521 | 24.29735 | 0.000698 | 0.014294 |
| HP | -2.95933 | 0.704343 | 24.28321 | 0.000699 | 0.014295 |
| TSPAN2 | -1.17988 | 7.365321 | 24.28294 | 0.000699 | 0.014295 |
| CST1 | -2.09847 | 0.889281 | 24.27433 | 0.0007 | 0.014299 |
| PCDH1 | 1.512641 | 4.616015 | 24.26563 | 0.000701 | 0.014303 |
| PCDHGA1 | 1.900104 | -0.44765 | 24.20956 | 0.000704 | 0.014358 |
| SYT1 | 1.110494 | 8.522041 | 24.22339 | 0.000705 | 0.014358 |
| SYN3 | 1.622411 | 3.488173 | 24.21831 | 0.000706 | 0.014358 |
| VWA5B1 | 1.590387 | 0.150571 | 23.99629 | 0.000712 | 0.014463 |
| PNPLA3 | 1.200072 | 2.883272 | 24.15464 | 0.000712 | 0.014463 |
| HSFX1 | 1.460483 | 2.469901 | 24.137 | 0.000714 | 0.014486 |
| LAMA3 | 1.619678 | 1.162251 | 24.1233 | 0.000715 | 0.014501 |
| TCAF2 | -1.02877 | 2.791147 | 23.89564 | 0.000718 | 0.014535 |
| TSPAN11 | 1.086131 | 6.46772 | 24.08368 | 0.00072 | 0.014556 |
| CCDC85C | 0.965263 | 4.208535 | 24.06316 | 0.000722 | 0.014585 |
| AF274858.1 | 0.988107 | 2.908206 | 23.84843 | 0.000723 | 0.01459 |
| AL022344.2 | 2.267138 | -0.68669 | 24.00117 | 0.000728 | 0.014689 |
| DLGAP3 | 1.26056 | 4.73383 | 23.94849 | 0.000734 | 0.014788 |
| ECEL1 | 1.07501 | 6.149543 | 23.9334 | 0.000736 | 0.014806 |
| FASN | 1.210568 | 7.725995 | 23.88288 | 0.000741 | 0.014902 |
| XBP1 | -1.14199 | 7.778537 | 23.87613 | 0.000742 | 0.014902 |
| PTPMT1 | -1.14883 | 3.119998 | 23.86486 | 0.000743 | 0.014912 |
| CAV1 | 1.914688 | -0.14862 | 23.85084 | 0.000744 | 0.014928 |
| RPL10P9 | 0.658811 | 6.364162 | 23.63365 | 0.000746 | 0.014933 |
| AL356585.2 | 1.374985 | 1.035851 | 23.7379 | 0.000747 | 0.014933 |
| LGR4 | -0.70842 | 7.066722 | 23.7028 | 0.000747 | 0.014933 |
| CACNG3 | 2.266783 | -0.43768 | 23.82183 | 0.000748 | 0.014933 |
| RASD2 | 1.200798 | 3.795833 | 23.81245 | 0.000749 | 0.014939 |
| TBXA2R | 3.169095 | 0.836315 | 23.79348 | 0.000751 | 0.014966 |
| SPRY3 | 0.979924 | 3.42142 | 23.7125 | 0.000758 | 0.015087 |
| ISM2 | 1.35315 | 1.278009 | 23.65844 | 0.000766 | 0.015222 |
| AC079296.1 | 1.898682 | -0.12666 | 23.65159 | 0.000767 | 0.015222 |
| AC140481.3 | -3.10659 | 0.302171 | 23.65094 | 0.000767 | 0.015222 |

| | | | | | |
|---|---|---|---|---|---|
| FAM50A | 0.695501 | 7.125993 | 23.46493 | 0.000767 | 0.015222 |
| BEX1 | 0.699344 | 9.465016 | 23.41756 | 0.000771 | 0.015272 |
| COL27A1 | -1.39854 | 7.657703 | 23.59332 | 0.000773 | 0.015311 |
| SNCA | 0.985855 | 8.252316 | 23.56933 | 0.000776 | 0.015351 |
| SATL1 | 1.292766 | 2.464874 | 23.54311 | 0.000779 | 0.015378 |
| DNAH10 | 1.08844 | 2.462631 | 23.42679 | 0.00078 | 0.015378 |
| SCD | 0.781125 | 9.031346 | 23.53801 | 0.00078 | 0.015378 |
| NR5A1 | 2.929639 | -0.73277 | 23.49679 | 0.000784 | 0.015457 |
| CCDC40 | -1.17939 | 4.376969 | 23.45475 | 0.000789 | 0.015522 |
| SNX8 | 0.924146 | 3.898247 | 23.45114 | 0.00079 | 0.015522 |
| TCAF1 | -0.86406 | 6.539708 | 23.44909 | 0.00079 | 0.015522 |
| CDK6 | 1.378554 | 3.202109 | 23.43484 | 0.000792 | 0.01554 |
| STAG3L5P | 1.365632 | 1.000993 | 23.29506 | 0.000797 | 0.015624 |
| GDPD2 | 0.894448 | 3.299584 | 23.1748 | 0.000799 | 0.015659 |
| RNA5SP515 | 2.449654 | -1.54234 | 23.16259 | 0.000801 | 0.015673 |
| ADAM29 | 2.826205 | -0.94842 | 23.33317 | 0.000804 | 0.015717 |
| AC023024.2 | 1.808304 | 0.489131 | 23.28739 | 0.000809 | 0.01581 |
| PPARGC1B | 1.008217 | 2.480892 | 23.06193 | 0.000813 | 0.015864 |
| KCND3 | 0.882664 | 5.331145 | 23.23881 | 0.000815 | 0.015864 |
| PLCG2 | 1.519352 | 0.633684 | 23.2368 | 0.000816 | 0.015864 |
| YPEL1 | 0.975998 | 2.610451 | 23.04136 | 0.000816 | 0.015864 |
| TMEM108 | 0.766278 | 4.34519 | 23.03856 | 0.000816 | 0.015864 |
| XK | 1.171115 | 4.626996 | 23.21799 | 0.000818 | 0.015868 |
| HLA-B | 0.769132 | 5.823765 | 23.21019 | 0.000819 | 0.015868 |
| PPL | 0.866709 | 4.289438 | 23.17406 | 0.000819 | 0.015868 |
| PSAT1P3 | 1.444259 | -0.13424 | 23.0121 | 0.000819 | 0.015868 |
| RNF144A-AS1 | 0.982943 | 3.371255 | 23.19526 | 0.000821 | 0.015879 |
| HPCAL1 | 1.106567 | 6.987555 | 23.15846 | 0.000825 | 0.015951 |
| ARMCX7P | 0.698856 | 5.729933 | 22.92845 | 0.00083 | 0.01599 |
| CCDC144NL-AS1 | -1.05169 | 5.468847 | 23.1139 | 0.000831 | 0.01599 |
| MYOZ2 | 1.823303 | 0.476169 | 23.11281 | 0.000831 | 0.01599 |
| ACKR3 | 2.133593 | -0.0019 | 23.11194 | 0.000831 | 0.01599 |
| NFIA | -1.54304 | 5.850772 | 23.11108 | 0.000831 | 0.01599 |
| PPIC | -1.07365 | 3.278888 | 23.08514 | 0.000834 | 0.016037 |
| MYCBP | -1.40428 | 2.635284 | 23.06045 | 0.000837 | 0.016082 |
| PPP1R1C | 1.187031 | 3.50153 | 23.04651 | 0.000839 | 0.016101 |
| AC080038.1 | -1.88935 | -0.10602 | 22.99726 | 0.000845 | 0.016206 |
| PYCARD | -2.18507 | -0.81636 | 22.93964 | 0.000853 | 0.016328 |
| CDON | 0.956943 | 3.548665 | 22.92671 | 0.000854 | 0.016328 |
| NKRF | 0.73239 | 5.630198 | 22.81338 | 0.000855 | 0.016328 |
| MIR503HG | -1.82034 | 0.335083 | 22.91279 | 0.000856 | 0.016328 |
| SERPINA12 | -3.14561 | -0.29957 | 22.91165 | 0.000856 | 0.016328 |
| NMU | 2.376973 | 0.035288 | 22.91042 | 0.000857 | 0.016328 |
| AC006058.4 | 1.739774 | 1.189406 | 22.88898 | 0.000859 | 0.016366 |
| LINC01201 | 1.222289 | 1.974323 | 22.87609 | 0.000861 | 0.016378 |
| GC | -1.01207 | 3.870456 | 22.87072 | 0.000862 | 0.016378 |
| NGB | 1.665179 | 0.562667 | 22.8631 | 0.000863 | 0.016378 |

| | | | | | |
|---|---|---|---|---|---|
| NXPH1 | 1.801831 | 5.519603 | 22.85981 | 0.000863 | 0.016378 |
| PCSK6 | 1.400991 | 2.98415 | 22.7867 | 0.000873 | 0.016545 |
| DSE | -1.17978 | 4.670575 | 22.76671 | 0.000875 | 0.01658 |
| AC134026.1 | 2.828567 | 1.047145 | 22.7491 | 0.000878 | 0.016609 |
| TAB3 | 0.903696 | 7.002773 | 22.73065 | 0.00088 | 0.01664 |
| TIMM8A | 0.870929 | 3.753269 | 22.53628 | 0.000881 | 0.016642 |
| CHMP1B2P | 1.223834 | 1.651775 | 22.628 | 0.000886 | 0.016715 |
| SMC1B | -2.05666 | -0.73755 | 22.67278 | 0.000888 | 0.016742 |
| CCDC163 | 1.148478 | 1.469694 | 22.45272 | 0.000893 | 0.016812 |
| NTF3 | 1.745572 | 0.57513 | 22.62117 | 0.000895 | 0.016843 |
| NEURL1B | 0.998611 | 3.279824 | 22.57057 | 0.000902 | 0.016938 |
| AL008721.2 | -1.02347 | 2.493301 | 22.38185 | 0.000903 | 0.016938 |
| C1QTNF1 | 1.985301 | 1.833405 | 22.56652 | 0.000903 | 0.016938 |
| ITGB3BP | -1.01695 | 3.135211 | 22.5203 | 0.000909 | 0.017042 |
| TSHZ2 | -0.82003 | 5.132332 | 22.50178 | 0.000912 | 0.017075 |
| RBFOX3 | 1.164911 | 5.319789 | 22.4838 | 0.000914 | 0.017107 |
| DAPL1 | -1.15209 | 3.390543 | 22.45428 | 0.000918 | 0.017169 |
| AC110285.2 | -1.60741 | 1.482841 | 22.42985 | 0.000922 | 0.017218 |
| MIR4697HG | 0.751604 | 5.907816 | 22.40669 | 0.000925 | 0.017263 |
| ZNF300 | 0.895925 | 5.831476 | 22.39743 | 0.000926 | 0.017272 |
| LRRN2 | 1.062548 | 6.454917 | 22.38632 | 0.000928 | 0.017286 |
| LGSN | 1.928884 | 3.247288 | 22.37098 | 0.00093 | 0.017311 |
| GLDC | 0.811119 | 3.575315 | 22.1818 | 0.000931 | 0.017313 |
| SEZ6 | 1.164838 | 7.011547 | 22.32631 | 0.000937 | 0.017397 |
| AC106798.1 | 2.904224 | -0.61959 | 22.31778 | 0.000938 | 0.017397 |
| SYNGR4 | -2.11459 | 0.46611 | 22.30542 | 0.00094 | 0.017397 |
| AC138356.2 | 1.242605 | 2.11009 | 22.30408 | 0.00094 | 0.017397 |
| NDN | 0.908637 | 8.383902 | 22.29918 | 0.000941 | 0.017397 |
| LINC01703 | 1.720008 | -0.56859 | 22.11594 | 0.000941 | 0.017397 |
| SOX21 | 1.070486 | 4.019451 | 22.24119 | 0.000949 | 0.017529 |
| EVC | 1.067104 | 2.991372 | 22.23739 | 0.00095 | 0.017529 |
| AC092017.1 | -0.90703 | 2.82952 | 22.0425 | 0.000952 | 0.017551 |
| ARSJ | -1.48519 | 3.731974 | 22.21425 | 0.000953 | 0.01756 |
| CEP41 | 0.690963 | 5.904878 | 22.0117 | 0.000956 | 0.017608 |
| OR2L6P | 3.125421 | -1.06788 | 22.17631 | 0.000959 | 0.017629 |
| CPAMD8 | 1.040911 | 3.806732 | 22.17159 | 0.000959 | 0.017629 |
| TSPAN9 | 1.009364 | 5.083118 | 22.14484 | 0.000963 | 0.017686 |
| GABRE | 1.194477 | 5.217226 | 22.12855 | 0.000966 | 0.017704 |
| MAOB | 1.33368 | 7.891897 | 22.12699 | 0.000966 | 0.017704 |
| CLEC18C | 1.130268 | 2.314518 | 22.10169 | 0.00097 | 0.017757 |
| AL662844.4 | -1.17754 | 2.608018 | 22.0951 | 0.000971 | 0.01776 |
| EPHA10 | 1.174615 | 3.860206 | 22.08546 | 0.000972 | 0.017771 |
| AC106886.5 | -1.34619 | 0.622939 | 21.88154 | 0.000976 | 0.017824 |
| RPL10P16 | 0.878754 | 2.760678 | 21.87429 | 0.000977 | 0.017829 |
| SORCS2 | 1.197037 | 4.475828 | 22.04105 | 0.000979 | 0.017846 |
| FP236241.1 | -1.43473 | 1.641027 | 22.03009 | 0.000981 | 0.017861 |
| HIST1H1C | -2.34593 | 2.053091 | 21.99729 | 0.000986 | 0.017912 |

| | | | | | |
|---|---|---|---|---|---|
| EBF4 | 1.135039 | 4.105957 | 21.9947 | 0.000986 | 0.017912 |
| PQLC3 | -1.11207 | 3.790519 | 21.99448 | 0.000986 | 0.017912 |
| AL162171.1 | 3.655108 | -1.41334 | 21.97961 | 0.000988 | 0.017938 |
| PRRC2B | 0.994098 | 7.24016 | 21.95049 | 0.000993 | 0.017991 |
| TNKS2-AS1 | -1.65678 | 0.308818 | 21.94927 | 0.000993 | 0.017991 |
| MAOA | 0.67539 | 5.665278 | 21.76574 | 0.000994 | 0.017993 |
| LINC02016 | -2.12754 | -0.6896 | 21.9189 | 0.000998 | 0.018045 |
| AC016700.2 | 0.809311 | 5.15786 | 21.89198 | 0.001002 | 0.01809 |
| AC044787.1 | -1.16837 | 6.072811 | 21.89161 | 0.001002 | 0.01809 |
| PART1 | -2.06209 | 4.295694 | 21.87715 | 0.001004 | 0.018112 |
| SEMA5B | 0.855389 | 4.768474 | 21.87267 | 0.001005 | 0.018112 |
| FBXW4 | -0.9312 | 4.551222 | 21.86243 | 0.001007 | 0.018116 |
| TMEM52 | 1.408233 | 2.514886 | 21.85373 | 0.001008 | 0.018116 |
| MUM1L1 | 1.435697 | 2.590298 | 21.84068 | 0.00101 | 0.018116 |
| AP000704.1 | -0.969 | 2.352896 | 21.6633 | 0.00101 | 0.018116 |
| RPL39P3 | 0.874675 | 6.376384 | 21.83771 | 0.001011 | 0.018116 |
| CYP2D6 | -0.85897 | 3.327319 | 21.65758 | 0.001011 | 0.018116 |
| EVX2 | 3.560316 | -1.49172 | 21.83196 | 0.001011 | 0.018116 |
| HMGB3P6 | 1.040903 | 3.257896 | 21.80765 | 0.001015 | 0.018165 |
| RGS10 | 1.536361 | 2.241993 | 21.80378 | 0.001016 | 0.018165 |
| FAT4 | 1.163827 | 3.300662 | 21.79324 | 0.001018 | 0.01817 |
| MAP1LC3A | 0.927954 | 6.365306 | 21.78911 | 0.001018 | 0.01817 |
| NEO1 | 0.614795 | 7.674463 | 21.61004 | 0.001019 | 0.01817 |
| SDC3 | 0.87444 | 8.942018 | 21.77727 | 0.00102 | 0.018177 |
| DEPTOR | 1.284514 | 0.515426 | 21.58775 | 0.001023 | 0.018203 |
| HEPH | -0.75213 | 5.986207 | 21.75628 | 0.001024 | 0.018205 |
| RAI2 | 0.697109 | 4.682362 | 21.57066 | 0.001025 | 0.018216 |
| THEMIS2 | 1.418101 | 2.30846 | 21.74152 | 0.001026 | 0.018216 |
| PTCH1 | 0.841557 | 4.551266 | 21.72406 | 0.001029 | 0.01825 |
| DRC1 | -1.38851 | 2.720745 | 21.71443 | 0.00103 | 0.018262 |
| EDC4 | 1.323238 | 4.455176 | 21.6922 | 0.001034 | 0.0183 |
| CPA4 | 2.100397 | -0.48933 | 21.69036 | 0.001034 | 0.0183 |
| COL3A1 | -1.15359 | 5.827981 | 21.67532 | 0.001037 | 0.018328 |
| DUSP10 | 1.641229 | 3.846972 | 21.66728 | 0.001038 | 0.018335 |
| AL359263.1 | 1.766743 | 0.755715 | 21.65122 | 0.001041 | 0.018366 |
| HOXB4 | -1.45529 | 3.342033 | 21.60774 | 0.001048 | 0.018457 |
| SRL | 1.180955 | 1.153726 | 21.43144 | 0.001048 | 0.018457 |
| AC087783.2 | 2.023945 | 1.039367 | 21.60338 | 0.001049 | 0.018457 |
| PDHA1 | 0.654576 | 7.108996 | 21.40206 | 0.001053 | 0.01851 |
| PCDHGB1 | 1.915699 | -0.49541 | 21.57448 | 0.001053 | 0.01851 |
| GPRC5D-AS1 | -1.19136 | 2.674051 | 21.5686 | 0.001054 | 0.018511 |
| FUNDC2P1 | 0.75956 | 3.793412 | 21.35318 | 0.001062 | 0.018624 |
| AL353596.1 | 1.916479 | -1.11588 | 21.34018 | 0.001064 | 0.018645 |
| AL353588.1 | 1.388638 | 0.007581 | 21.33158 | 0.001065 | 0.018645 |
| KLF9 | -1.61571 | 3.877086 | 21.50147 | 0.001066 | 0.018645 |
| MSL3 | 0.642592 | 5.518418 | 21.32195 | 0.001067 | 0.018653 |
| ATG9A | 1.358803 | 4.253468 | 21.45696 | 0.001073 | 0.018726 |

| | | | | | |
|---|---|---|---|---|---|
| ADAMTS7P3 | 1.197023 | 1.198695 | 21.28439 | 0.001073 | 0.018726 |
| WWC3 | 0.800199 | 5.835969 | 21.45284 | 0.001074 | 0.018726 |
| AL139393.2 | -0.95877 | 2.801 | 21.25804 | 0.001078 | 0.018767 |
| FAM89A | 1.425966 | 2.176092 | 21.42393 | 0.001079 | 0.018767 |
| TCF15 | 2.251176 | -0.0188 | 21.42282 | 0.001079 | 0.018767 |
| TFF3 | -1.31808 | 5.738211 | 21.40412 | 0.001082 | 0.018807 |
| MNX1-AS1 | 1.884688 | -0.79609 | 21.28324 | 0.001084 | 0.01883 |
| GABRR1 | -1.10882 | 2.686438 | 21.37119 | 0.001088 | 0.018874 |
| RBM7 | -1.30184 | 5.539208 | 21.36243 | 0.001089 | 0.018884 |
| ZDHHC11B | -1.05312 | 5.257331 | 21.33292 | 0.001095 | 0.018954 |
| PI4KAP2 | -0.71754 | 5.258234 | 21.17265 | 0.001095 | 0.018954 |
| WASF4P | 1.369738 | 0.455634 | 21.14476 | 0.001098 | 0.01898 |
| AL590004.3 | -1.8757 | 0.271567 | 21.30496 | 0.001099 | 0.01898 |
| NDUFA1 | 0.737388 | 6.649941 | 21.30433 | 0.0011 | 0.01898 |
| TMEM97 | 0.651132 | 5.104421 | 21.11516 | 0.001103 | 0.019023 |
| PGK1P2 | 1.278393 | 4.198413 | 21.27944 | 0.001104 | 0.019023 |
| AC078909.2 | -1.78227 | 0.033821 | 21.26579 | 0.001106 | 0.01904 |
| IKBKG | 0.850739 | 4.455327 | 21.26349 | 0.001107 | 0.01904 |
| AC019131.2 | -1.11991 | 1.716867 | 21.07885 | 0.00111 | 0.019061 |
| ZDBF2 | 0.879382 | 6.967091 | 21.24537 | 0.00111 | 0.019061 |
| BMP1 | -1.14588 | 4.997609 | 21.241 | 0.001111 | 0.019061 |
| FAM43A | 1.271585 | 2.287137 | 21.23036 | 0.001113 | 0.019077 |
| HNF4A | -2.21461 | 0.419764 | 21.21458 | 0.001115 | 0.019109 |
| FAM156A | 1.139529 | 1.618061 | 21.03778 | 0.001117 | 0.01912 |
| LRRC75A | 0.800205 | 4.645074 | 21.17774 | 0.001118 | 0.01912 |
| ADAMTS7 | 1.220502 | 4.011858 | 21.18035 | 0.001122 | 0.019166 |
| CRIPT | -0.94554 | 4.526385 | 21.16991 | 0.001123 | 0.019182 |
| GK | 1.059467 | 4.32425 | 21.16391 | 0.001125 | 0.019185 |
| GALNT7 | -0.92486 | 6.09814 | 21.15138 | 0.001127 | 0.019197 |
| SNTG1 | -1.83563 | 4.108017 | 21.14232 | 0.001128 | 0.019197 |
| LTBP4 | 1.154369 | 4.506403 | 21.14227 | 0.001128 | 0.019197 |
| DPEP1 | 1.891378 | 0.255064 | 21.13921 | 0.001129 | 0.019197 |
| ARID3C | 1.850789 | 0.293794 | 21.11048 | 0.001134 | 0.01927 |
| IGSF8 | 0.793464 | 6.811183 | 21.09576 | 0.001137 | 0.019291 |
| TMBIM1 | -1.84596 | 0.272345 | 21.09202 | 0.001138 | 0.019291 |
| ME1 | -0.81311 | 6.56176 | 21.08827 | 0.001138 | 0.019291 |
| SYN2 | 0.813982 | 5.46717 | 21.06563 | 0.001142 | 0.019345 |
| AC016252.1 | -2.1012 | -0.9799 | 21.04716 | 0.001146 | 0.019346 |
| ASS1 | 0.895572 | 4.986687 | 21.04221 | 0.001147 | 0.019346 |
| ZHX2 | -1.00265 | 4.355255 | 21.04086 | 0.001147 | 0.019346 |
| CORIN | -1.98642 | 5.160857 | 21.04063 | 0.001147 | 0.019346 |
| FKBP5 | 0.953618 | 3.377587 | 21.03965 | 0.001147 | 0.019346 |
| PTGFRN | -0.63665 | 7.417865 | 20.86103 | 0.00115 | 0.019363 |
| HIST2H2AA4 | -1.92373 | 2.67859 | 21.01973 | 0.001151 | 0.019363 |
| TENT5C | -1.85204 | 2.556648 | 21.0189 | 0.001151 | 0.019363 |
| AC234783.1 | 1.628348 | -0.5169 | 20.84775 | 0.001152 | 0.019366 |
| NUTM2A | 1.103684 | 1.493088 | 20.8315 | 0.001155 | 0.019385 |

| | | | | | |
|---|---|---|---|---|---|
| HLA-DPB2 | 2.460561 | -0.96139 | 20.99192 | 0.001156 | 0.019385 |
| PTPN3 | 0.992734 | 3.857197 | 20.99155 | 0.001156 | 0.019385 |
| CXCL16 | 1.375348 | 2.06076 | 20.97727 | 0.001159 | 0.019393 |
| IGSF9B | 1.390848 | 3.103534 | 20.97586 | 0.001159 | 0.019393 |
| RBP4 | 0.742148 | 4.21962 | 20.80374 | 0.001161 | 0.019393 |
| SYNDIG1 | -0.87942 | 4.252541 | 20.96512 | 0.001161 | 0.019393 |
| SEMA3F | 0.895408 | 3.753279 | 20.95967 | 0.001162 | 0.019393 |
| LUC7L | -0.92475 | 6.857909 | 20.95886 | 0.001162 | 0.019393 |
| AC244107.1 | 1.242869 | 0.880372 | 20.78628 | 0.001164 | 0.019402 |
| NPFFR2 | 1.423153 | 1.674771 | 20.9294 | 0.001168 | 0.019448 |
| AC087203.3 | 1.883566 | 1.595489 | 20.92605 | 0.001168 | 0.019448 |
| AC138969.1 | -0.77493 | 6.385475 | 20.91842 | 0.00117 | 0.019456 |
| ZBED3 | -0.89154 | 4.504979 | 20.90386 | 0.001173 | 0.019472 |
| SLIT3 | 0.98975 | 6.025557 | 20.90123 | 0.001173 | 0.019472 |
| BAMBI | 1.121496 | 3.495963 | 20.89832 | 0.001174 | 0.019472 |
| HOXB7 | -1.23905 | 3.708659 | 20.88395 | 0.001176 | 0.019502 |
| NKAP | 0.837451 | 5.635907 | 20.83187 | 0.001186 | 0.019651 |
| SSTR2 | 0.771939 | 4.79414 | 20.75588 | 0.001188 | 0.019667 |
| CRYZ | -1.28374 | 5.370358 | 20.7894 | 0.001195 | 0.019755 |
| STRA6 | 1.856889 | 2.556279 | 20.7657 | 0.001199 | 0.019801 |
| FST | 1.080951 | 4.747355 | 20.76003 | 0.0012 | 0.019801 |
| GRIK3 | 0.880082 | 5.800421 | 20.75678 | 0.001201 | 0.019801 |
| CD248 | 1.043236 | 2.799339 | 20.75552 | 0.001201 | 0.019801 |
| HIST1H4H | -2.00914 | 2.361283 | 20.72422 | 0.001207 | 0.019886 |
| AC021683.1 | 2.123653 | -0.79417 | 20.70477 | 0.001211 | 0.019933 |
| NIFK-AS1 | -1.02343 | 3.041668 | 20.69899 | 0.001212 | 0.019936 |
| CXorf56 | 0.729909 | 5.225927 | 20.61907 | 0.001214 | 0.01994 |
| AC239800.1 | -1.56886 | -0.4252 | 20.52666 | 0.001215 | 0.01994 |
| AL080317.2 | -1.32938 | 0.753149 | 20.59002 | 0.001218 | 0.019983 |
| PIFO | -1.53175 | 3.091873 | 20.66357 | 0.001219 | 0.019987 |
| ZCCHC24 | -1.14374 | 2.984669 | 20.61769 | 0.001229 | 0.020121 |
| PPP2R5D | 0.955379 | 5.392659 | 20.58782 | 0.001235 | 0.020204 |
| PALMD | -1.44342 | 3.148992 | 20.57257 | 0.001238 | 0.020238 |
| MLLT3 | -0.73353 | 6.200191 | 20.5668 | 0.001239 | 0.020239 |
| TBC1D14 | 0.957219 | 5.289296 | 20.55434 | 0.001241 | 0.020239 |
| AL049871.1 | -1.4653 | 1.236305 | 20.55416 | 0.001241 | 0.020239 |
| AL590560.2 | 1.678918 | 0.519922 | 20.5526 | 0.001242 | 0.020239 |
| ZRSR2 | -0.95002 | 4.912474 | 20.54496 | 0.001243 | 0.020244 |
| HSD17B7P2 | -1.00435 | 2.121934 | 20.38111 | 0.001244 | 0.020244 |
| MYCL | 1.09387 | 1.913944 | 20.40838 | 0.001245 | 0.020244 |
| LINC00884 | 2.279218 | 0.200745 | 20.50822 | 0.001251 | 0.020323 |
| ARHGEF11 | 0.736961 | 6.245604 | 20.49987 | 0.001253 | 0.020334 |
| RPS10P7 | 2.072016 | -0.82254 | 20.49187 | 0.001254 | 0.020341 |
| BEST4 | -1.6049 | 0.834271 | 20.48837 | 0.001255 | 0.020341 |
| KHDC3L | -2.17437 | -0.92865 | 20.47997 | 0.001257 | 0.020353 |
| ENPP1 | -1.60749 | 3.681908 | 20.45919 | 0.001261 | 0.020395 |
| GPX7 | -1.12969 | 4.176344 | 20.45753 | 0.001261 | 0.020395 |

| | | | | | |
|---|---|---|---|---|---|
| TOLLIP-AS1 | -1.19407 | 0.898895 | 20.26477 | 0.001269 | 0.020484 |
| TRANK1 | 1.655934 | 1.452743 | 20.4215 | 0.001269 | 0.020484 |
| LINC00205 | 0.619278 | 6.205382 | 20.2393 | 0.001274 | 0.020552 |
| AC018638.6 | -1.60256 | 2.022311 | 20.35605 | 0.001283 | 0.020674 |
| THBD | -2.17459 | 2.209215 | 20.34709 | 0.001284 | 0.020688 |
| SYNPO2L | -1.79382 | -0.28388 | 20.34003 | 0.001286 | 0.020696 |
| GAS8 | 0.700974 | 4.051577 | 20.17652 | 0.001287 | 0.020703 |
| DDC | 1.779821 | 5.466633 | 20.30396 | 0.001294 | 0.020787 |
| CDYL2 | 0.956882 | 5.260862 | 20.2921 | 0.001296 | 0.020798 |
| SH3RF3 | 0.727072 | 4.53342 | 20.13473 | 0.001296 | 0.020798 |
| BDNF | -0.71818 | 6.668518 | 20.28657 | 0.001297 | 0.020798 |
| CHST1 | 0.930034 | 5.837201 | 20.26567 | 0.001302 | 0.020854 |
| TMEM132E | 1.091012 | 3.200545 | 20.23651 | 0.001308 | 0.020926 |
| LMOD1 | 1.547498 | 0.363901 | 20.23391 | 0.001309 | 0.020926 |
| HIST1H4E | -1.79989 | 0.611825 | 20.22997 | 0.00131 | 0.020926 |
| AC008060.1 | 2.295857 | 1.103224 | 20.22604 | 0.00131 | 0.020926 |
| AC104825.1 | -0.69965 | 4.78723 | 20.06163 | 0.001312 | 0.020941 |
| LINC02397 | 1.897938 | -1.13364 | 20.05235 | 0.001314 | 0.020948 |
| PDRG1 | 0.637643 | 5.410253 | 20.05035 | 0.001315 | 0.020948 |
| F8 | 0.935049 | 3.583922 | 20.19758 | 0.001317 | 0.020957 |
| AC093915.1 | -1.59337 | -0.63124 | 20.03856 | 0.001318 | 0.020957 |
| AC245177.1 | 1.994606 | 0.326825 | 20.17935 | 0.001321 | 0.02099 |
| GPD1 | -1.8086 | 5.430888 | 20.16341 | 0.001324 | 0.021029 |
| AGAP6 | -0.81675 | 4.651687 | 20.15331 | 0.001326 | 0.021039 |
| KDM2B | 0.722302 | 5.148218 | 20.06644 | 0.001327 | 0.021039 |
| AC106047.1 | -1.74663 | -0.94188 | 19.9831 | 0.00133 | 0.021072 |
| SLC7A4 | 1.331406 | 2.827736 | 20.12659 | 0.001332 | 0.021093 |
| AC092375.2 | 1.323399 | 0.612304 | 20.00386 | 0.001336 | 0.02114 |
| SCG5 | -0.98394 | 6.009201 | 20.06835 | 0.001345 | 0.021253 |
| FBLN7 | 1.184259 | 1.378052 | 19.98409 | 0.001346 | 0.021253 |
| RTN4RL1 | 1.124125 | 3.100993 | 20.06258 | 0.001346 | 0.021253 |
| RASSF9 | -1.32952 | 1.784274 | 20.03675 | 0.001352 | 0.021329 |
| FZD10 | 2.678032 | -0.93293 | 20.03096 | 0.001354 | 0.021333 |
| MMP24 | 1.49866 | 5.984718 | 20.02123 | 0.001356 | 0.021351 |
| LINC01138 | -0.77931 | 3.757755 | 19.85333 | 0.001359 | 0.02139 |
| HAGLR | 0.857488 | 3.337985 | 19.83694 | 0.001363 | 0.021433 |
| ECHDC3 | -1.89883 | 1.455043 | 19.96321 | 0.001369 | 0.021509 |
| AC139426.1 | 2.322919 | -1.31571 | 19.9557 | 0.001371 | 0.02152 |
| SLC4A11 | -1.51444 | 2.373733 | 19.95008 | 0.001372 | 0.021524 |
| ABCB4 | 1.463242 | 1.81596 | 19.92135 | 0.001379 | 0.021604 |
| SPHKAP | 2.760186 | 1.132725 | 19.9186 | 0.001379 | 0.021604 |
| GPD2 | -0.99301 | 4.560466 | 19.8909 | 0.001386 | 0.021688 |
| NFE2L3 | 0.766665 | 4.958152 | 19.87544 | 0.001389 | 0.021728 |
| TCP10L | 1.198708 | 0.874318 | 19.71696 | 0.001391 | 0.02174 |
| RGS22 | -2.0415 | -0.15623 | 19.85384 | 0.001394 | 0.021774 |
| SH2B3 | 0.816792 | 4.229354 | 19.82034 | 0.001402 | 0.02188 |
| PAPLN | -1.52078 | 1.468877 | 19.79739 | 0.001408 | 0.021948 |

| | | | | | |
|---|---|---|---|---|---|
| WFIKKN2 | 2.193747 | -0.17366 | 19.77016 | 0.001414 | 0.022012 |
| SYTL4 | 1.227185 | 1.393519 | 19.76893 | 0.001414 | 0.022012 |
| HIST1H2AC | -2.25958 | 4.604308 | 19.76657 | 0.001415 | 0.022012 |
| AC107398.3 | 1.238264 | 3.766726 | 19.74505 | 0.00142 | 0.022075 |
| SCGN | 2.650137 | -0.0617 | 19.71683 | 0.001427 | 0.022164 |
| TIMM17B | 1.060252 | 5.573163 | 19.68876 | 0.001434 | 0.022249 |
| KHDC1 | -1.03811 | 1.718311 | 19.53646 | 0.001435 | 0.022249 |
| HOXA2 | -1.28257 | 2.861504 | 19.66851 | 0.001439 | 0.022295 |
| HCCS | 0.658115 | 4.91807 | 19.5124 | 0.001441 | 0.022308 |
| ZNF529-AS1 | -0.99628 | 2.029678 | 19.49973 | 0.001444 | 0.022339 |
| PDE2A | 0.924211 | 3.99079 | 19.64238 | 0.001445 | 0.022343 |
| AC012640.4 | -1.81749 | 0.428467 | 19.63298 | 0.001447 | 0.022361 |
| CDH9 | 1.640861 | 2.116878 | 19.5828 | 0.00146 | 0.022502 |
| LGALS3 | 0.868283 | 2.353566 | 19.43534 | 0.00146 | 0.022502 |
| FIBIN | -1.34819 | 2.14142 | 19.58117 | 0.00146 | 0.022502 |
| AC007614.1 | 1.980023 | -0.51127 | 19.57824 | 0.001461 | 0.022502 |
| AMMECR1 | -1.34059 | 3.350044 | 19.56009 | 0.001465 | 0.022554 |
| CFAP77 | -2.15114 | 1.688653 | 19.53181 | 0.001472 | 0.022646 |
| IL17RD | 0.702092 | 6.260421 | 19.5257 | 0.001474 | 0.022652 |
| FAR2P2 | 1.502172 | 1.570133 | 19.51864 | 0.001476 | 0.022653 |
| ZNF189 | -0.69582 | 5.807298 | 19.51663 | 0.001476 | 0.022653 |
| PLPP5 | -0.98594 | 4.778731 | 19.49557 | 0.001481 | 0.022705 |
| RND1 | -0.79126 | 4.461885 | 19.49443 | 0.001482 | 0.022705 |
| TUBBP5 | 1.141485 | 2.057552 | 19.47782 | 0.001486 | 0.022752 |
| KCNH4 | 1.265721 | 1.538054 | 19.46379 | 0.001489 | 0.02279 |
| HSP90AB3P | 0.902223 | 6.286576 | 19.45852 | 0.001491 | 0.022793 |
| ANKRD18CP | -1.95852 | -0.89331 | 19.42155 | 0.0015 | 0.02292 |
| DNPEP | 0.813043 | 4.13004 | 19.40439 | 0.001503 | 0.022946 |
| ENPP7P11 | -1.88994 | 2.237431 | 19.39476 | 0.001507 | 0.022991 |
| LINC00461 | 1.251763 | 3.058847 | 19.38158 | 0.001511 | 0.023016 |
| GPX8 | -1.34872 | 5.520995 | 19.37976 | 0.001511 | 0.023016 |
| AC013652.1 | 1.690062 | 0.041158 | 19.36862 | 0.001514 | 0.023021 |
| CAT | 1.592823 | 4.24194 | 19.36537 | 0.001515 | 0.023021 |
| PCSK1 | 0.870813 | 9.277311 | 19.36526 | 0.001515 | 0.023021 |
| PHLPP1 | 0.747885 | 5.494397 | 19.34578 | 0.00152 | 0.023081 |
| CA10 | -1.70519 | 5.012412 | 19.32867 | 0.001524 | 0.023132 |
| NDUFB5 | -0.85961 | 6.453176 | 19.30667 | 0.00153 | 0.023189 |
| AC064875.1 | 1.185969 | 1.73529 | 19.30472 | 0.001531 | 0.023189 |
| HS3ST1 | 1.159563 | 2.4016 | 19.30113 | 0.001531 | 0.023189 |
| TRAPPC2 | 0.752647 | 4.554027 | 19.2074 | 0.001534 | 0.023213 |
| PINLYP | -1.59917 | 0.241531 | 19.27465 | 0.001538 | 0.023261 |
| NXPH3 | 1.613677 | 2.918239 | 19.26647 | 0.001541 | 0.02327 |
| VCX3A | 1.470006 | 0.196332 | 19.26368 | 0.001541 | 0.02327 |
| WWTR1 | -1.04179 | 6.6588 | 19.25572 | 0.001543 | 0.023285 |
| ZNF233 | 1.13269 | 1.59156 | 19.17948 | 0.001553 | 0.023401 |
| IGFBP7 | -1.57253 | 2.782728 | 19.21807 | 0.001553 | 0.023401 |
| TEX26-AS1 | 1.454067 | -0.47584 | 19.06908 | 0.001555 | 0.023409 |

| | | | | | |
|---|---|---|---|---|---|
| ADAM23 | 0.689915 | 6.608848 | 19.17523 | 0.001565 | 0.02354 |
| TGFBR2 | -1.32983 | 5.029821 | 19.16157 | 0.001569 | 0.023578 |
| CPB1 | -2.02063 | -0.83665 | 19.12611 | 0.001578 | 0.023691 |
| XIAPP1 | 1.354537 | 0.626662 | 19.12517 | 0.001579 | 0.023691 |
| CARTPT | -1.31148 | 7.184526 | 19.11627 | 0.001581 | 0.02371 |
| RPL10P8 | 1.226657 | 1.847268 | 19.10693 | 0.001583 | 0.023724 |
| GLP1R | 1.495018 | 2.024188 | 19.10442 | 0.001584 | 0.023724 |
| AL354984.1 | -1.54332 | -0.45312 | 18.93847 | 0.001591 | 0.023809 |
| KCNK12 | 1.322344 | 5.048576 | 19.05092 | 0.001599 | 0.02391 |
| AKNAD1 | 2.161349 | -0.68268 | 19.0405 | 0.001602 | 0.023935 |
| S1PR1 | 1.992344 | -0.71689 | 19.02488 | 0.001606 | 0.023982 |
| HK2 | 1.544253 | 6.7892 | 19.0095 | 0.00161 | 0.024025 |
| HSPA6 | 3.034321 | 1.164282 | 19.00617 | 0.001611 | 0.024025 |
| ZCWPW1 | -0.99842 | 2.504253 | 18.99115 | 0.001613 | 0.024036 |
| AL445253.1 | -1.89341 | 1.201066 | 18.98855 | 0.001616 | 0.024064 |
| DMRT2 | 1.707247 | 1.308603 | 18.97791 | 0.001619 | 0.024091 |
| ADAMTS6 | -0.80918 | 3.478777 | 18.82776 | 0.001622 | 0.024094 |
| TNFAIP3 | 0.887664 | 2.576208 | 18.82738 | 0.001622 | 0.024094 |
| OCM | -1.56473 | 0.651505 | 18.96448 | 0.001623 | 0.024094 |
| KLHL14 | 1.373524 | 2.953078 | 18.94999 | 0.001627 | 0.024135 |
| PHLDB2 | -1.26882 | 3.518825 | 18.94633 | 0.001628 | 0.024135 |
| RHBDL3 | 0.91149 | 4.846455 | 18.93058 | 0.001633 | 0.024184 |
| AJ239328.1 | 1.925129 | -1.11398 | 18.81193 | 0.001634 | 0.024193 |
| AC003986.2 | -1.99043 | -0.42512 | 18.90764 | 0.001639 | 0.024245 |
| CACNG5 | 2.120534 | 0.916049 | 18.89568 | 0.001643 | 0.024278 |
| ZNF677 | 0.869278 | 3.943031 | 18.88839 | 0.001645 | 0.02428 |
| AL033530.1 | 1.320637 | 2.410401 | 18.88546 | 0.001645 | 0.02428 |
| CDC25A | 0.833965 | 4.060198 | 18.88288 | 0.001646 | 0.02428 |
| NT5C | -0.71288 | 4.359928 | 18.72946 | 0.00165 | 0.024306 |
| RSPH1 | -1.85622 | 4.840202 | 18.86425 | 0.001652 | 0.024306 |
| PYURF | 0.682099 | 6.54712 | 18.86058 | 0.001653 | 0.024306 |
| CCNK | 1.013447 | 1.714478 | 18.71865 | 0.001654 | 0.024306 |
| ST8SIA4 | -1.11192 | 6.046692 | 18.85365 | 0.001655 | 0.024306 |
| FAAH2 | 1.01118 | 2.20134 | 18.77521 | 0.001655 | 0.024306 |
| KCNRG | -1.66872 | -0.36109 | 18.84786 | 0.001656 | 0.024306 |
| MCF2L2 | 0.790231 | 4.33599 | 18.82867 | 0.001662 | 0.02435 |
| NPR3 | 1.427221 | 1.702631 | 18.82831 | 0.001662 | 0.02435 |
| TRPC5 | 1.990342 | 0.478863 | 18.82424 | 0.001663 | 0.02435 |
| CPNE4 | 0.872324 | 3.172755 | 18.72485 | 0.001664 | 0.02435 |
| SLCO3A1 | 0.715236 | 4.752469 | 18.68016 | 0.001669 | 0.024401 |
| ADCYAP1R1 | 1.011421 | 6.524423 | 18.77609 | 0.001677 | 0.024476 |
| SPTB | 1.48029 | 3.140227 | 18.77514 | 0.001677 | 0.024476 |
| TRAPPC12-AS1 | 1.45519 | 0.432102 | 18.77251 | 0.001678 | 0.024476 |
| HSPB7 | -1.54949 | 2.381197 | 18.77094 | 0.001679 | 0.024476 |
| EMILIN3 | 2.001315 | 1.667168 | 18.75785 | 0.001682 | 0.024515 |
| FAM13A | 0.752937 | 5.18925 | 18.74687 | 0.001686 | 0.024545 |
| ANXA4 | -1.05406 | 3.661015 | 18.73985 | 0.001688 | 0.024552 |

| | | | | | |
|---|---|---|---|---|---|
| EFL1P1 | 1.282145 | 1.17217 | 18.73405 | 0.001689 | 0.024552 |
| MGAT5B | 1.044801 | 5.90014 | 18.73303 | 0.00169 | 0.024552 |
| APCDD1 | -0.6084 | 5.656935 | 18.59135 | 0.001691 | 0.024555 |
| NFATC1 | -1.27334 | 1.695106 | 18.71355 | 0.001696 | 0.024601 |
| AC138969.2 | 1.110858 | 4.89175 | 18.69745 | 0.0017 | 0.024653 |
| FAM86B2 | 0.981346 | 1.545688 | 18.55621 | 0.001702 | 0.024656 |
| ZNF192P1 | -1.12693 | 1.737291 | 18.66107 | 0.001711 | 0.024774 |
| POMK | 1.270395 | 1.880007 | 18.65732 | 0.001712 | 0.024774 |
| CYTL1 | 2.328007 | -1.17419 | 18.63105 | 0.00172 | 0.024871 |
| MAP3K15 | 1.679284 | 0.586925 | 18.62717 | 0.001721 | 0.024871 |
| NUTM2A-AS1 | 0.810871 | 2.88748 | 18.48741 | 0.001723 | 0.024871 |
| GBP3 | -1.52187 | 0.662856 | 18.58915 | 0.001733 | 0.025002 |
| ADCK2 | 0.83479 | 5.189011 | 18.58313 | 0.001735 | 0.025011 |
| BEND4 | 1.037732 | 4.355802 | 18.56735 | 0.00174 | 0.025063 |
| GCH1 | 1.281942 | 5.809641 | 18.55807 | 0.001743 | 0.025086 |
| DEAF1 | 0.784693 | 6.764677 | 18.55021 | 0.001745 | 0.025104 |
| IDSP1 | 1.618993 | 0.352138 | 18.54405 | 0.001747 | 0.025113 |
| CXorf51B | 2.294522 | -1.3333 | 18.54008 | 0.001748 | 0.025113 |
| TMEM63B | 0.84617 | 6.999463 | 18.53241 | 0.00175 | 0.025129 |
| DUSP27 | -2.97009 | 2.578272 | 18.52866 | 0.001752 | 0.025129 |
| AP000280.1 | -1.72117 | 3.674246 | 18.5204 | 0.001754 | 0.025145 |
| DERL2 | -0.64868 | 6.394901 | 18.49299 | 0.001755 | 0.025145 |
| AC005062.1 | 1.531471 | 2.301769 | 18.50184 | 0.00176 | 0.025195 |
| ZNF814 | -0.77273 | 4.668404 | 18.49043 | 0.001763 | 0.025228 |
| UNC5A | 1.107897 | 4.051262 | 18.48625 | 0.001765 | 0.025229 |
| PSMA3-AS1 | -1.15806 | 6.029918 | 18.4823 | 0.001766 | 0.025229 |
| AC140125.2 | -3.01566 | -1.31627 | 18.45184 | 0.001776 | 0.025347 |
| FAM13A-AS1 | 1.20117 | 1.25702 | 18.43675 | 0.00178 | 0.025398 |
| CAVIN1 | -1.00818 | 3.725736 | 18.42644 | 0.001784 | 0.025416 |
| HSDL2 | -0.88503 | 5.082118 | 18.42484 | 0.001784 | 0.025416 |
| AC145124.1 | 0.977298 | 3.218909 | 18.38612 | 0.001796 | 0.025573 |
| RORB | 1.19508 | 1.293712 | 18.37911 | 0.001799 | 0.025586 |
| CNNM2 | 0.804437 | 4.860202 | 18.37532 | 0.0018 | 0.025586 |
| VWA8 | 0.672185 | 4.461982 | 18.23312 | 0.001803 | 0.025595 |
| QRFPR | 1.834477 | -0.49601 | 18.36565 | 0.001803 | 0.025595 |
| FAM24B | 1.935803 | -1.11025 | 18.29806 | 0.001805 | 0.025604 |
| PCYT2 | 0.738709 | 4.769946 | 18.34782 | 0.001806 | 0.025604 |
| CATIP | -2.22147 | -0.54394 | 18.34836 | 0.001808 | 0.02562 |
| SPTBN2 | 0.968747 | 6.484045 | 18.31404 | 0.001819 | 0.025746 |
| CTAGE1 | 1.86752 | -0.80602 | 18.30973 | 0.001821 | 0.025746 |
| MPZ | 1.26267 | 0.874333 | 18.30903 | 0.001821 | 0.025746 |
| APBA2 | 0.853139 | 7.261389 | 18.29872 | 0.001824 | 0.025776 |
| SLC7A3 | 1.174228 | 3.132862 | 18.28705 | 0.001828 | 0.025806 |
| KCNH6 | 1.342247 | 0.40716 | 18.26652 | 0.001829 | 0.025806 |
| AL033397.2 | -2.08466 | -0.80022 | 18.2752 | 0.001832 | 0.025831 |
| LINC02506 | 1.596269 | 1.624789 | 18.27108 | 0.001833 | 0.025832 |
| ANO1 | 2.185762 | -0.93877 | 18.26501 | 0.001835 | 0.025842 |

| | | | | | |
|---|---|---|---|---|---|
| AC132192.2 | -1.12159 | 1.912035 | 18.25203 | 0.00184 | 0.025869 |
| NUDT4B | 0.872768 | 2.51446 | 18.12016 | 0.00184 | 0.025869 |
| PGK1 | 0.833424 | 9.03742 | 18.24671 | 0.001841 | 0.025873 |
| FDXACB1 | -0.95443 | 2.035995 | 18.10874 | 0.001844 | 0.025887 |
| MYL6 | -0.59356 | 9.372824 | 18.10198 | 0.001846 | 0.025901 |
| AL627230.2 | 1.259788 | 0.305075 | 18.09428 | 0.001848 | 0.02591 |
| WIPF3 | 0.671408 | 5.39704 | 18.16053 | 0.00185 | 0.02591 |
| MTCO1P40 | -1.02777 | 5.534059 | 18.21951 | 0.00185 | 0.02591 |
| JUNB | -1.21086 | 5.755435 | 18.20849 | 0.001854 | 0.025932 |
| MCAM | 0.821318 | 5.757219 | 18.20715 | 0.001854 | 0.025932 |
| ALS2CL | 0.883689 | 4.04214 | 18.19991 | 0.001857 | 0.025939 |
| VWA5B2 | 1.309517 | 3.206619 | 18.19781 | 0.001858 | 0.025939 |
| GLRA3 | 1.746395 | 2.37533 | 18.18642 | 0.001861 | 0.025974 |
| WASF2 | -0.96196 | 5.974807 | 18.17933 | 0.001864 | 0.025975 |
| CLDN9 | 1.621415 | 0.673295 | 18.17868 | 0.001864 | 0.025975 |
| DNAH3 | 2.289228 | -0.96042 | 18.17404 | 0.001865 | 0.025979 |
| ALG1L6P | 1.499963 | -0.54307 | 18.03233 | 0.001869 | 0.026014 |
| OPRK1 | 1.060836 | 3.110118 | 18.15892 | 0.00187 | 0.026014 |
| PIGCP1 | 1.057276 | 1.388948 | 18.01695 | 0.001874 | 0.026052 |
| S100A16 | -1.20286 | 0.697175 | 18.00684 | 0.001878 | 0.026067 |
| SERPINB7 | 1.373975 | 1.38438 | 18.13618 | 0.001878 | 0.026067 |
| PCDHAC2 | 1.385754 | 3.066988 | 18.11416 | 0.001886 | 0.026152 |
| CD36 | -0.81997 | 7.6958 | 18.09572 | 0.001892 | 0.026221 |
| C11orf71 | -0.94169 | 5.69831 | 18.08217 | 0.001896 | 0.026267 |
| MAP7D2 | 0.82022 | 6.462651 | 18.07069 | 0.0019 | 0.026281 |
| REPS2 | 1.025982 | 5.295021 | 18.06914 | 0.001901 | 0.026281 |
| ZSCAN1 | 0.924973 | 2.940416 | 18.06778 | 0.001901 | 0.026281 |
| CHRDL1 | -1.90797 | 9.464617 | 18.06235 | 0.001903 | 0.026289 |
| H2BFS | -1.60372 | 1.301333 | 18.0563 | 0.001905 | 0.0263 |
| DNMT3A | -0.82094 | 7.098907 | 18.05156 | 0.001907 | 0.026305 |
| AC007256.1 | 1.249113 | 0.554808 | 17.93612 | 0.00191 | 0.026326 |
| F13A1 | -2.87723 | 6.199464 | 18.02921 | 0.001914 | 0.026362 |
| PDE6B | -0.82929 | 2.677602 | 17.89956 | 0.001915 | 0.026362 |
| AC005821.1 | -1.70723 | -1.05612 | 17.8648 | 0.001927 | 0.026494 |
| YPEL2 | 0.861297 | 4.245178 | 17.99276 | 0.001927 | 0.026494 |
| SYTL2 | 1.039311 | 4.404346 | 17.9893 | 0.001928 | 0.026494 |
| IL1A | 1.013978 | 1.499248 | 17.83358 | 0.001938 | 0.02661 |
| FRG1BP | 1.038943 | 2.966436 | 17.94656 | 0.001943 | 0.026659 |
| ZNF441 | -1.30852 | 4.110487 | 17.94375 | 0.001944 | 0.026659 |
| AC004543.1 | 1.804962 | 0.084244 | 17.92066 | 0.001952 | 0.02674 |
| BRCC3P1 | 1.622009 | 0.519913 | 17.91949 | 0.001953 | 0.02674 |
| TICRR | 1.182277 | 0.617898 | 17.77734 | 0.001958 | 0.026794 |
| SLC25A5-AS1 | 0.885479 | 2.724767 | 17.76454 | 0.001963 | 0.026839 |
| TRHDE-AS1 | 1.063214 | 2.187827 | 17.86718 | 0.001971 | 0.026924 |
| C16orf82 | 1.38211 | 0.390014 | 17.86675 | 0.001971 | 0.026924 |
| ZNF93 | 0.908136 | 2.019559 | 17.72115 | 0.001978 | 0.026999 |
| AC245297.3 | -0.77247 | 3.094615 | 17.71278 | 0.001981 | 0.027013 |

| | | | | | |
|---|---|---|---|---|---|
| TDG | -0.72462 | 5.578664 | 17.83732 | 0.001982 | 0.027013 |
| AC027612.1 | 1.020931 | 2.093721 | 17.81627 | 0.001984 | 0.027026 |
| ZNF224 | -0.81141 | 4.553377 | 17.82519 | 0.001986 | 0.027037 |
| RNFT1 | -1.16721 | 3.507649 | 17.81061 | 0.001992 | 0.027081 |
| C4orf47 | -1.48688 | 3.358536 | 17.80892 | 0.001992 | 0.027081 |
| GABRA5 | 0.877257 | 5.742163 | 17.80134 | 0.001995 | 0.027092 |
| SERPINB1 | -0.74187 | 3.505106 | 17.67345 | 0.001996 | 0.027092 |
| ANKH | 0.600577 | 5.847117 | 17.66265 | 0.002 | 0.027127 |
| KCNQ1DN | 3.023993 | -0.95935 | 17.77988 | 0.002003 | 0.027151 |
| HHIPL1 | 1.049399 | 1.38638 | 17.64524 | 0.002006 | 0.027151 |
| VAT1L | 0.761269 | 9.331662 | 17.76916 | 0.002007 | 0.027151 |
| ID4 | 0.731552 | 7.375204 | 17.76697 | 0.002007 | 0.027151 |
| RTL8A | 0.685488 | 7.002498 | 17.76539 | 0.002008 | 0.027151 |
| PCDHA11 | 1.307154 | 1.218524 | 17.75686 | 0.002011 | 0.027158 |
| NUDT4P2 | 0.980138 | 2.943737 | 17.75673 | 0.002011 | 0.027158 |
| AC131097.3 | -1.79207 | -0.55727 | 17.7512 | 0.002013 | 0.027168 |
| FANCB | 1.272555 | 4.36E-06 | 17.59352 | 0.002025 | 0.027312 |
| MCM6 | 0.779825 | 3.084266 | 17.58478 | 0.002028 | 0.027338 |
| GRK3 | 0.650891 | 6.557143 | 17.7039 | 0.002031 | 0.027348 |
| SPRY1 | 1.09652 | 2.496495 | 17.67902 | 0.00204 | 0.027454 |
| FAM155B | 0.873853 | 5.45399 | 17.66842 | 0.002044 | 0.027489 |
| LRRIQ1 | -1.43622 | 3.163519 | 17.65342 | 0.002049 | 0.027533 |
| WTAP | -0.56815 | 6.942437 | 17.52831 | 0.00205 | 0.027533 |
| SOCS3 | -1.65333 | 2.93691 | 17.64425 | 0.002053 | 0.027556 |
| AP006284.1 | 1.098252 | 0.966707 | 17.50309 | 0.002059 | 0.027623 |
| EML5 | 1.413453 | 1.41845 | 17.62391 | 0.00206 | 0.027623 |
| AL022068.1 | -1.17858 | 1.361229 | 17.60569 | 0.002067 | 0.027697 |
| D2HGDH | 1.012384 | 3.167384 | 17.59442 | 0.002072 | 0.0277 |
| NFASC | 0.617391 | 7.555261 | 17.52405 | 0.002072 | 0.0277 |
| PSEN2 | 0.718745 | 5.303778 | 17.59133 | 0.002073 | 0.0277 |
| TLR2 | 1.778716 | -0.36384 | 17.59077 | 0.002073 | 0.0277 |
| BCAS4 | 0.591688 | 5.622735 | 17.46165 | 0.002075 | 0.027711 |
| PGAP1 | 0.963823 | 7.567962 | 17.56949 | 0.002081 | 0.02775 |
| NEB | 0.822936 | 2.932482 | 17.44519 | 0.002081 | 0.02775 |
| KCNJ12 | 1.683306 | 2.232644 | 17.56662 | 0.002082 | 0.02775 |
| ORMDL1 | -0.74329 | 6.446323 | 17.56202 | 0.002084 | 0.027753 |
| ADAMTS20 | 1.415308 | 0.269504 | 17.558 | 0.002085 | 0.027753 |
| AC026801.2 | -0.99045 | 1.676259 | 17.43237 | 0.002086 | 0.027753 |
| GLYCTK-AS1 | -1.0334 | 2.60615 | 17.54476 | 0.00209 | 0.027789 |
| EPHB3 | 0.896381 | 4.361361 | 17.52393 | 0.002098 | 0.027877 |
| AC207130.1 | 1.645601 | -0.11608 | 17.48497 | 0.002113 | 0.028059 |
| ZNRF2P2 | 1.689884 | -0.00252 | 17.4649 | 0.002121 | 0.028144 |
| ACACA | 0.584523 | 6.845589 | 17.32004 | 0.00213 | 0.028238 |
| XCL1 | 1.461322 | 0.943621 | 17.43973 | 0.002131 | 0.028238 |
| FAM66D | 1.381168 | 0.58388 | 17.4322 | 0.002134 | 0.028241 |
| PTPRF | 0.684724 | 7.301272 | 17.43218 | 0.002134 | 0.028241 |
| PHYKPL | -0.81669 | 4.089163 | 17.42822 | 0.002136 | 0.028243 |

| | | | | | |
|---|---|---|---|---|---|
| NETO2 | 0.579642 | 6.731545 | 17.29274 | 0.002141 | 0.028298 |
| STK32C | 0.730337 | 5.98758 | 17.40899 | 0.002143 | 0.028307 |
| PPFIA3 | 0.886879 | 6.100327 | 17.39529 | 0.002149 | 0.028343 |
| FER1L6 | 1.826428 | -1.17598 | 17.27375 | 0.002149 | 0.028343 |
| HEMK1 | -0.76523 | 4.610391 | 17.37126 | 0.002158 | 0.028449 |
| SDSL | -0.8126 | 3.270305 | 17.24715 | 0.00216 | 0.028452 |
| ZNF835 | 1.677619 | 2.301706 | 17.36089 | 0.002162 | 0.028467 |
| NPFFR1 | 2.190988 | -0.12215 | 17.33173 | 0.002174 | 0.028587 |
| FPGT-TNNI3K | 2.009278 | -0.96254 | 17.32675 | 0.002176 | 0.028587 |
| LINC02562 | 1.307169 | 0.44393 | 17.32114 | 0.002176 | 0.028587 |
| PRTFDC1 | -0.82811 | 4.192162 | 17.31961 | 0.002179 | 0.028587 |
| AC009102.2 | 1.726619 | -0.90135 | 17.24318 | 0.002182 | 0.028587 |
| AL137786.1 | -1.27468 | 1.653965 | 17.31046 | 0.002183 | 0.028587 |
| IGFBP4 | -0.8562 | 8.380312 | 17.30961 | 0.002183 | 0.028587 |
| GOLGA6L5P | -1.11369 | 1.563005 | 17.30952 | 0.002183 | 0.028587 |
| TBC1D2B | 0.799879 | 3.580861 | 17.23833 | 0.002185 | 0.028587 |
| PHGDH | 0.886314 | 4.955017 | 17.30074 | 0.002186 | 0.028587 |
| LMO1 | 1.016226 | 3.174198 | 17.29892 | 0.002187 | 0.028587 |
| NCAN | 0.729401 | 7.232657 | 17.29041 | 0.002191 | 0.028587 |
| CERS6 | 0.81907 | 6.195194 | 17.28992 | 0.002191 | 0.028587 |
| GPR143 | 0.922661 | 2.337373 | 17.16964 | 0.002191 | 0.028587 |
| FAM222A | 1.045564 | 5.055762 | 17.28435 | 0.002193 | 0.028597 |
| ZNF185 | 0.828527 | 4.97072 | 17.27538 | 0.002197 | 0.028626 |
| TIMMDC1 | -0.66988 | 6.207985 | 17.26658 | 0.0022 | 0.028655 |
| KCNE5 | 1.683562 | 0.366274 | 17.25237 | 0.002206 | 0.028712 |
| HIST1H2AE | -1.52421 | 0.465195 | 17.2404 | 0.002211 | 0.028757 |
| MPDU1 | -0.69999 | 4.886439 | 17.20037 | 0.002219 | 0.028841 |
| COMMD10 | -0.93949 | 3.95759 | 17.21451 | 0.002222 | 0.028859 |
| UPF3B | 0.913759 | 5.676555 | 17.1984 | 0.002228 | 0.028927 |
| NTAN1P2 | -1.07897 | 0.59212 | 17.06232 | 0.002235 | 0.029002 |
| AC006329.1 | 0.991022 | 2.18937 | 17.14463 | 0.002248 | 0.029134 |
| ACAT1 | -0.71096 | 6.378214 | 17.1499 | 0.002248 | 0.029134 |
| SEC23IP | -0.82307 | 6.155161 | 17.14011 | 0.002253 | 0.029168 |
| SMG6 | 1.01407 | 4.127512 | 17.07991 | 0.002278 | 0.029479 |
| HIST1H4I | -1.48204 | 2.334877 | 17.07427 | 0.00228 | 0.029491 |
| DISP3 | 0.968446 | 3.402418 | 17.05595 | 0.002288 | 0.029526 |
| NRIP3 | 1.227568 | 5.838405 | 17.05035 | 0.002291 | 0.029526 |
| TMEM123 | -1.33063 | 7.101898 | 17.04894 | 0.002291 | 0.029526 |
| ABCA2 | 0.857914 | 6.522438 | 17.04644 | 0.002292 | 0.029526 |
| NCR3LG1 | 1.293683 | 3.537545 | 17.04607 | 0.002292 | 0.029526 |
| LRRC37A3 | -0.69702 | 5.249394 | 17.04402 | 0.002293 | 0.029526 |
| NTSR1 | 1.302881 | 0.824015 | 17.04346 | 0.002294 | 0.029526 |
| AGAP14P | -0.90446 | 2.710965 | 17.00104 | 0.002295 | 0.029526 |
| ZNF667 | 0.732796 | 4.391232 | 17.00539 | 0.002298 | 0.029545 |
| ITPRIPL1 | 1.217214 | 2.120049 | 17.03082 | 0.002299 | 0.029545 |
| AC005224.3 | -2.01218 | -0.53552 | 17.0098 | 0.002308 | 0.029643 |
| AL049838.1 | -1.04532 | 5.382194 | 17.0004 | 0.002312 | 0.029676 |

| | | | | | |
|---|---|---|---|---|---|
| HIST2H2BE | -0.98859 | 5.212419 | 16.93635 | 0.00234 | 0.030016 |
| CELF3 | 0.908585 | 7.883331 | 16.92822 | 0.002344 | 0.03003 |
| POU3F1 | 1.678663 | 1.518088 | 16.92712 | 0.002344 | 0.03003 |
| MYOF | -1.38916 | 2.129663 | 16.9128 | 0.00235 | 0.030091 |
| ZNF561 | -0.6276 | 5.079995 | 16.79178 | 0.002353 | 0.030091 |
| PEAR1 | 1.409531 | -0.6597 | 16.79065 | 0.002353 | 0.030091 |
| ONECUT1 | 1.048066 | 2.143523 | 16.89392 | 0.002359 | 0.030143 |
| IRX5 | -1.19829 | 2.91438 | 16.88226 | 0.002364 | 0.03017 |
| CD247 | 1.335984 | -0.37046 | 16.76627 | 0.002364 | 0.03017 |
| AC008060.3 | 2.609352 | -1.6301 | 16.87911 | 0.002365 | 0.03017 |
| EPAS1 | 0.982996 | 2.152975 | 16.8491 | 0.002373 | 0.030235 |
| REEP3 | -1.33236 | 6.21184 | 16.8611 | 0.002373 | 0.030235 |
| MEIS1 | -0.79922 | 7.081841 | 16.85651 | 0.002375 | 0.030242 |
| MMP15 | 1.036901 | 6.543787 | 16.84292 | 0.002382 | 0.030301 |
| NDUFB11 | 0.675188 | 6.965682 | 16.83942 | 0.002383 | 0.030302 |
| ASPM | 1.245336 | 0.241046 | 16.7109 | 0.002389 | 0.030348 |
| BCL7A | 0.847693 | 5.513395 | 16.81914 | 0.002392 | 0.030348 |
| ENO1 | 0.821923 | 10.11455 | 16.8171 | 0.002393 | 0.030348 |
| CDK1 | 1.42811 | -0.22567 | 16.75072 | 0.002394 | 0.030348 |
| LINC01011 | -0.99113 | 2.757007 | 16.8146 | 0.002394 | 0.030348 |
| PTRHD1 | -0.74949 | 3.034511 | 16.69708 | 0.002396 | 0.030348 |
| NBPF15 | 0.693278 | 4.077451 | 16.67754 | 0.002405 | 0.030442 |
| CHRNA6 | 0.765068 | 3.050799 | 16.66868 | 0.002409 | 0.03046 |
| AC124303.2 | 1.966529 | 1.089864 | 16.78239 | 0.002409 | 0.03046 |
| TTLL3 | -0.98318 | 3.573027 | 16.77819 | 0.002411 | 0.030465 |
| LINC01915 | 1.454982 | 0.355567 | 16.74536 | 0.002426 | 0.030636 |
| CMTM6 | -1.28322 | 6.345258 | 16.73598 | 0.00243 | 0.030671 |
| SLC38A3 | -1.61913 | 3.954956 | 16.72264 | 0.002436 | 0.030711 |
| PTPRB | 1.044401 | 1.032067 | 16.60375 | 0.002439 | 0.030711 |
| CACNA1E | 0.690844 | 6.956413 | 16.71645 | 0.002439 | 0.030711 |
| CFAP70 | -1.77032 | 2.908625 | 16.7161 | 0.002439 | 0.030711 |
| AC006015.1 | 1.700047 | -0.3671 | 16.70956 | 0.002442 | 0.03073 |
| ERV3-1 | 0.742986 | 3.4312 | 16.57786 | 0.002451 | 0.030801 |
| AL139125.2 | -1.55376 | -0.72492 | 16.57763 | 0.002451 | 0.030801 |
| CDHR4 | -1.3729 | -0.1995 | 16.56729 | 0.002456 | 0.030841 |
| AQP10 | 2.604373 | -1.1124 | 16.67755 | 0.002457 | 0.030841 |
| ITGB5 | 1.075507 | 3.009326 | 16.65971 | 0.002465 | 0.03092 |
| FLNA | 0.834622 | 8.022775 | 16.65773 | 0.002466 | 0.03092 |
| CHRM4 | 1.771296 | 2.23608 | 16.65397 | 0.002468 | 0.030923 |
| POLI | -0.79047 | 5.685534 | 16.64729 | 0.002471 | 0.030943 |
| STX16-NPEPL1 | 1.76758 | 0.050706 | 16.63896 | 0.002475 | 0.030959 |
| RCN3 | -0.78494 | 4.381065 | 16.63821 | 0.002476 | 0.030959 |
| AC243591.1 | 2.263264 | -0.98182 | 16.62511 | 0.002482 | 0.031017 |
| DEC1 | -2.17765 | -0.55572 | 16.62182 | 0.002483 | 0.031017 |
| RGPD1 | 0.548203 | 5.979847 | 16.48363 | 0.002496 | 0.031137 |
| RPL9 | -0.74408 | 7.022759 | 16.59511 | 0.002496 | 0.031137 |
| NEK2 | 1.877735 | -0.79867 | 16.58062 | 0.002503 | 0.031204 |

| | | | | | |
|---|---|---|---|---|---|
| NR3C2 | 2.523147 | 0.033914 | 16.57224 | 0.002507 | 0.031235 |
| PLEKHM1P1 | 1.24139 | 1.59257 | 16.56157 | 0.002512 | 0.03128 |
| PNMA6A | 0.877331 | 5.614503 | 16.54431 | 0.00252 | 0.031364 |
| COA6-AS1 | -1.10161 | 0.718483 | 16.42932 | 0.002522 | 0.031365 |
| DPY19L2P2 | 0.761578 | 3.202518 | 16.40388 | 0.002534 | 0.0315 |
| AC104041.1 | 1.974197 | -0.50983 | 16.50932 | 0.002537 | 0.031516 |
| KLHDC8A | 0.76082 | 5.390659 | 16.50196 | 0.002541 | 0.031541 |
| COMMD7 | 0.53217 | 6.534233 | 16.38206 | 0.002545 | 0.03156 |
| PCDHA2 | 1.465966 | 1.416962 | 16.49248 | 0.002545 | 0.03156 |
| NTNG1 | -1.12568 | 3.718862 | 16.48925 | 0.002547 | 0.031561 |
| BCL11B | 1.394209 | -0.32988 | 16.36237 | 0.002555 | 0.031639 |
| GOLGA3 | -0.5891 | 7.130521 | 16.40374 | 0.002558 | 0.031656 |
| KCNMA1 | 0.574539 | 6.52 | 16.35261 | 0.002559 | 0.031656 |
| L3HYPDH | -1.2158 | 2.671591 | 16.46083 | 0.002561 | 0.031656 |
| NENF | 0.599487 | 5.78958 | 16.33598 | 0.002569 | 0.031742 |
| AC048380.2 | -1.45581 | 1.072718 | 16.42253 | 0.00258 | 0.031851 |
| SDHD | -0.71396 | 4.660693 | 16.40985 | 0.002586 | 0.031901 |
| SUPT3H | -0.70563 | 3.742989 | 16.29783 | 0.002587 | 0.031901 |
| AC009078.3 | -1.89062 | -1.11737 | 16.40178 | 0.00259 | 0.031911 |
| MFAP3 | 0.793651 | 5.443459 | 16.40011 | 0.002591 | 0.031911 |
| SEMA7A | 1.178744 | 5.45353 | 16.39466 | 0.002593 | 0.031925 |
| SAMHD1 | 0.593951 | 4.844911 | 16.27526 | 0.002598 | 0.031964 |
| CHRAC1 | -1.19294 | 3.368357 | 16.3709 | 0.002605 | 0.032032 |
| ZSCAN23 | 1.253442 | 0.67966 | 16.36437 | 0.002608 | 0.03204 |
| AC011287.1 | 1.707755 | -1.15994 | 16.25381 | 0.002609 | 0.03204 |
| CUL4B | 0.679731 | 7.44217 | 16.35835 | 0.002611 | 0.032052 |
| VTN | -1.57782 | 3.258012 | 16.33724 | 0.002622 | 0.032163 |
| HS6ST1P1 | 1.538625 | 2.864756 | 16.32981 | 0.002626 | 0.032187 |
| TFE3 | 0.669133 | 6.966331 | 16.32719 | 0.002627 | 0.032187 |
| ISLR | 1.228995 | 0.728224 | 16.30097 | 0.00264 | 0.03233 |
| IKBIP | -0.82285 | 4.52166 | 16.29656 | 0.002643 | 0.032338 |
| AL357055.3 | -1.26926 | 0.469399 | 16.29003 | 0.002646 | 0.03236 |
| ARHGAP35 | 0.731322 | 7.77821 | 16.28648 | 0.002648 | 0.032362 |
| AC015922.2 | 1.179852 | 1.298581 | 16.27824 | 0.002652 | 0.032395 |
| CXCL6 | -1.80625 | 0.361436 | 16.2627 | 0.00266 | 0.032472 |
| TMC6 | 1.722997 | 2.028462 | 16.24472 | 0.002669 | 0.032561 |
| COL6A3 | -1.69042 | 1.686689 | 16.2424 | 0.00267 | 0.032561 |
| BMP8A | 3.225862 | -0.81892 | 16.20546 | 0.00269 | 0.032771 |
| GABPB1-AS1 | -0.82161 | 5.038232 | 16.20305 | 0.002691 | 0.032771 |
| FAM19A1 | 1.687503 | 0.485286 | 16.19666 | 0.002694 | 0.032774 |
| ZCCHC9 | -0.75331 | 4.471387 | 16.1963 | 0.002694 | 0.032774 |
| AC019205.1 | -0.99952 | 2.312 | 16.18573 | 0.0027 | 0.032784 |
| GDE1 | -0.57236 | 6.687751 | 16.07656 | 0.002701 | 0.032784 |
| SNRNP25 | 0.721085 | 4.972483 | 16.18261 | 0.002702 | 0.032784 |
| PCDH18 | -0.66026 | 6.005965 | 16.17798 | 0.002704 | 0.032784 |
| BNIP3 | 0.708899 | 7.514698 | 16.17752 | 0.002704 | 0.032784 |
| MYCN | 0.754212 | 3.539626 | 16.06892 | 0.002705 | 0.032784 |

| | | | | | |
|---|---|---|---|---|---|
| ADAMTS18 | 0.99955 | 1.247954 | 16.04928 | 0.002715 | 0.032879 |
| TMX2 | -0.57919 | 6.133215 | 16.04784 | 0.002716 | 0.032879 |
| CLIC6 | 1.784405 | 0.377661 | 16.14832 | 0.00272 | 0.032879 |
| KIF14 | 1.537775 | -0.69283 | 16.05072 | 0.00272 | 0.032879 |
| AC048341.1 | -0.91448 | 2.953438 | 16.14622 | 0.002721 | 0.032879 |
| AC138932.1 | 1.348915 | 2.77068 | 16.13172 | 0.002728 | 0.032942 |
| ANKHD1-EIF4EBP3 | -0.8743 | 2.338231 | 16.02321 | 0.002729 | 0.032942 |
| GRIK4 | 0.663112 | 4.837986 | 16.05087 | 0.002732 | 0.032953 |
| ZNF674-AS1 | 0.723359 | 3.538597 | 16.00666 | 0.002738 | 0.03301 |
| AC103760.1 | -1.52264 | -0.5215 | 16.07614 | 0.002742 | 0.033038 |
| ZNF671 | 1.306882 | 2.408392 | 16.10219 | 0.002744 | 0.033044 |
| PDYN | 1.570982 | 0.033094 | 16.08969 | 0.002751 | 0.033105 |
| ZNF552 | -0.87057 | 2.007049 | 15.97892 | 0.002753 | 0.033105 |
| POP5 | -0.82397 | 4.266853 | 16.081 | 0.002755 | 0.033105 |
| VCY1B | 2.737013 | -1.17282 | 16.08052 | 0.002756 | 0.033105 |
| HLF | 0.845986 | 2.064544 | 15.96725 | 0.002759 | 0.03313 |
| MIR7-3HG | 1.088518 | 2.644256 | 16.06165 | 0.002766 | 0.033188 |
| CSPP1 | -0.84221 | 4.944052 | 16.0573 | 0.002768 | 0.033197 |
| MRPL19 | -0.73456 | 4.848645 | 16.04585 | 0.002774 | 0.033251 |
| CCDC65 | -0.84601 | 3.277544 | 16.03259 | 0.002781 | 0.03329 |
| PDK3 | 0.843844 | 3.265624 | 16.03239 | 0.002781 | 0.03329 |
| PPP3CB-AS1 | -1.52517 | 0.076081 | 16.02801 | 0.002784 | 0.03329 |
| SHPRH | 0.763085 | 3.621671 | 15.96406 | 0.002786 | 0.03329 |
| AC020891.2 | 1.906421 | -1.03658 | 16.02019 | 0.002788 | 0.03329 |
| CRHR2 | 1.458908 | 1.245093 | 16.01498 | 0.002791 | 0.03329 |
| EMP3 | -0.95925 | 4.989216 | 16.01337 | 0.002792 | 0.03329 |
| TRIM7 | 1.252981 | 1.263239 | 16.01042 | 0.002793 | 0.03329 |
| MROH7 | -2.13189 | -0.46162 | 16.01041 | 0.002793 | 0.03329 |
| AL354793.1 | 1.726202 | 0.464211 | 16.00997 | 0.002794 | 0.03329 |
| AC092969.1 | 1.211698 | 2.446939 | 15.9884 | 0.002805 | 0.033411 |
| KRT18P48 | 1.541245 | 0.034327 | 15.98342 | 0.002808 | 0.033424 |
| DIO3OS | 1.271391 | 2.330805 | 15.95588 | 0.002823 | 0.033584 |
| LHX1 | 1.514117 | 0.933476 | 15.95181 | 0.002826 | 0.033592 |
| MGST1 | -0.8879 | 6.386156 | 15.94736 | 0.002828 | 0.033601 |
| DLC1 | 0.679164 | 4.308726 | 15.83892 | 0.00283 | 0.033604 |
| GABRG2 | 0.710383 | 6.055094 | 15.92375 | 0.002841 | 0.033718 |
| NLRP1 | -1.25344 | 9.962205 | 15.92025 | 0.002843 | 0.033721 |
| KCNK3 | 1.117415 | 8.066769 | 15.90631 | 0.002851 | 0.033794 |
| LAMTOR5 | -0.61117 | 6.083178 | 15.88043 | 0.002861 | 0.033889 |
| ZNF512 | -0.58579 | 6.713755 | 15.82803 | 0.002862 | 0.033889 |
| UBE2A | 0.59216 | 7.559742 | 15.84219 | 0.002864 | 0.033889 |
| AC016727.1 | -1.33169 | 1.626466 | 15.87945 | 0.002866 | 0.03389 |
| CRYBB2 | 1.327538 | 0.35511 | 15.87479 | 0.002868 | 0.03389 |
| SYAP1 | -0.59136 | 7.450578 | 15.8428 | 0.002869 | 0.03389 |
| SRRM2-AS1 | -1.72803 | -0.32257 | 15.87119 | 0.00287 | 0.03389 |
| DLX1 | 1.896811 | -0.78738 | 15.86657 | 0.002873 | 0.033901 |
| ADAP2 | 0.916755 | 2.152931 | 15.76999 | 0.002878 | 0.033937 |

| | | | | | |
|---|---|---|---|---|---|
| DOC2B | 0.924751 | 7.09073 | 15.84267 | 0.002887 | 0.034021 |
| ZSCAN12P1 | -0.99981 | 1.127237 | 15.73467 | 0.002889 | 0.034029 |
| UBQLN4P1 | 1.084243 | 3.530359 | 15.81716 | 0.002901 | 0.034118 |
| KANK3 | 1.504695 | -0.09643 | 15.81398 | 0.002903 | 0.034118 |
| AL390728.6 | -1.45482 | 2.738808 | 15.81393 | 0.002903 | 0.034118 |
| MAD1L1 | 0.796398 | 4.447131 | 15.81357 | 0.002903 | 0.034118 |
| AL442067.2 | -2.19922 | -1.24699 | 15.78965 | 0.002917 | 0.034259 |
| PIK3IP1 | 0.924927 | 1.767976 | 15.68206 | 0.002919 | 0.034269 |
| SCARB1 | 0.746208 | 4.38552 | 15.76986 | 0.002928 | 0.034339 |
| SAPCD2 | 2.009759 | 1.625516 | 15.76704 | 0.00293 | 0.034339 |
| NBL1 | 1.170466 | 5.233397 | 15.76616 | 0.00293 | 0.034339 |
| NME3 | -1.45147 | 3.458688 | 15.75909 | 0.002934 | 0.034367 |
| FRRS1L | 0.707176 | 7.754484 | 15.75024 | 0.00294 | 0.034402 |
| SLC10A3 | 0.994056 | 4.241803 | 15.74813 | 0.002941 | 0.034402 |
| CAMKV | 0.770824 | 6.243511 | 15.7375 | 0.002947 | 0.03444 |
| AC107294.1 | -1.20712 | 0.063677 | 15.63407 | 0.002947 | 0.03444 |
| CDC14A | -1.51725 | 3.405264 | 15.73352 | 0.002949 | 0.034442 |
| MUC15 | -3.1716 | 1.217367 | 15.72353 | 0.002955 | 0.034457 |
| TMEM266 | 1.257621 | 2.470301 | 15.72178 | 0.002956 | 0.034457 |
| MTMR9LP | -0.90426 | 1.643616 | 15.61674 | 0.002958 | 0.034457 |
| INE1 | 1.46145 | -0.67821 | 15.61546 | 0.002958 | 0.034457 |
| DNAAF1 | -1.36996 | 4.116108 | 15.71691 | 0.002959 | 0.034457 |
| AL589987.1 | 1.502045 | 1.941318 | 15.68379 | 0.002978 | 0.034664 |
| AC233280.1 | 1.423575 | -0.4468 | 15.58663 | 0.002984 | 0.034706 |
| NPM1 | -0.78382 | 7.891159 | 15.65264 | 0.002997 | 0.034791 |
| TRAF3 | 0.61965 | 6.074337 | 15.64932 | 0.002999 | 0.034791 |
| PIM2 | 0.692712 | 3.935315 | 15.54509 | 0.003 | 0.034791 |
| AGER | -1.01934 | 2.5056 | 15.64625 | 0.003001 | 0.034791 |
| CCDC74A | 0.758827 | 4.198659 | 15.64529 | 0.003001 | 0.034791 |
| CFAP52 | -1.88116 | 2.186205 | 15.64431 | 0.003002 | 0.034791 |
| NUAK1 | 0.721505 | 6.06007 | 15.64225 | 0.003003 | 0.034791 |
| AC142472.1 | -1.13273 | 0.612462 | 15.53613 | 0.003006 | 0.034802 |
| KNSTRN | 0.604091 | 4.865923 | 15.5212 | 0.003015 | 0.034887 |
| EIF4A1P9 | 1.554445 | -0.57105 | 15.59125 | 0.003033 | 0.035085 |
| COL21A1 | -1.62877 | 2.460562 | 15.58453 | 0.003038 | 0.035112 |
| AC011043.1 | 1.259481 | 0.590707 | 15.57868 | 0.003041 | 0.03513 |
| PODXL | 0.653688 | 4.542829 | 15.47349 | 0.003044 | 0.03513 |
| ZNF829 | 1.193295 | 1.54604 | 15.5734 | 0.003044 | 0.03513 |
| SKAP2 | -0.54976 | 5.468431 | 15.4682 | 0.003047 | 0.03514 |
| CD74 | 0.769408 | 4.851003 | 15.56533 | 0.003049 | 0.035147 |
| C2orf72 | 0.549844 | 5.287021 | 15.46157 | 0.003051 | 0.035148 |
| PPCDC | -0.92214 | 1.624506 | 15.44475 | 0.003061 | 0.03524 |
| GPRASP1 | 0.794193 | 6.103163 | 15.54359 | 0.003062 | 0.03524 |
| LUC7L2 | -0.76751 | 5.73322 | 15.51811 | 0.003078 | 0.035399 |
| GALNT18 | 0.861863 | 4.489347 | 15.51107 | 0.003082 | 0.035429 |
| AMY1C | -1.77778 | 2.682959 | 15.49936 | 0.003089 | 0.035492 |
| TMEM176B | 1.117346 | 4.896007 | 15.49473 | 0.003092 | 0.035494 |

| | | | | | |
|---|---|---|---|---|---|
| GATA3-AS1 | 1.592143 | 0.537577 | 15.49338 | 0.003093 | 0.035494 |
| LRRC3 | 0.7547 | 5.531417 | 15.47538 | 0.003104 | 0.035561 |
| EBF3 | 0.92281 | 6.035369 | 15.47359 | 0.003105 | 0.035561 |
| RNF128 | 1.306456 | 1.928147 | 15.47323 | 0.003106 | 0.035561 |
| LINC01905 | 2.00837 | -1.51594 | 15.41598 | 0.003106 | 0.035561 |
| SFRP2 | 1.890018 | 5.170439 | 15.46841 | 0.003109 | 0.035572 |
| C1QTNF3 | -1.31998 | 4.690208 | 15.46457 | 0.003111 | 0.035579 |
| TUBA1B | 0.986967 | 7.580786 | 15.4585 | 0.003115 | 0.035598 |
| UCP2 | -1.14684 | 5.166788 | 15.45612 | 0.003116 | 0.035598 |
| BMPR1B | -1.38206 | 3.554308 | 15.45351 | 0.003118 | 0.035598 |
| MSTN | -1.26223 | 0.61781 | 15.44718 | 0.003122 | 0.035623 |
| RBPMS | 1.341801 | 1.137726 | 15.42693 | 0.003134 | 0.035747 |
| PCDHB17P | 1.082887 | 0.691451 | 15.32388 | 0.003137 | 0.035754 |
| NECTIN1 | 1.045977 | 6.727483 | 15.41871 | 0.00314 | 0.035757 |
| ZNF718 | 1.031922 | 2.34228 | 15.41722 | 0.00314 | 0.035757 |
| ZNF602P | 1.544094 | -0.57274 | 15.41147 | 0.003144 | 0.035766 |
| FAM228B | -0.86825 | 3.475383 | 15.41035 | 0.003145 | 0.035766 |
| AGAP1 | 0.80696 | 7.626589 | 15.40173 | 0.00315 | 0.035808 |
| TBX18 | -1.52413 | 3.960721 | 15.39815 | 0.003152 | 0.035814 |
| ARHGAP20 | 1.421256 | 2.58255 | 15.38872 | 0.003158 | 0.035862 |
| AC020916.1 | -0.94349 | 2.256262 | 15.371 | 0.00317 | 0.035969 |
| AC012507.2 | 1.375781 | -0.14576 | 15.36021 | 0.003172 | 0.035973 |
| GVQW2 | -1.49577 | 0.020798 | 15.35711 | 0.003178 | 0.03603 |
| S100B | -0.85335 | 3.165636 | 15.35323 | 0.003181 | 0.036036 |
| PRELP | 1.91233 | 0.997486 | 15.34936 | 0.003183 | 0.036036 |
| AC005332.2 | 1.21311 | 1.289904 | 15.34797 | 0.003184 | 0.036036 |
| AL035413.1 | 1.178683 | -0.06446 | 15.24228 | 0.003189 | 0.036068 |
| CCBE1 | 1.747595 | -0.89077 | 15.31638 | 0.003205 | 0.036189 |
| MID2 | 1.009454 | 4.267744 | 15.31635 | 0.003205 | 0.036189 |
| SCARNA7 | -2.71874 | -1.34006 | 15.31582 | 0.003205 | 0.036189 |
| KCNIP4 | 1.095089 | 3.265973 | 15.29429 | 0.003219 | 0.036314 |
| BMI1 | -0.65486 | 6.062496 | 15.29318 | 0.00322 | 0.036314 |
| GAPDHP38 | 0.922663 | 2.454563 | 15.27764 | 0.00323 | 0.036381 |
| LINC01592 | -1.64735 | -0.43964 | 15.27383 | 0.003232 | 0.036381 |
| SSPN | -1.45706 | 3.565007 | 15.27318 | 0.003232 | 0.036381 |
| AL691520.1 | 1.581144 | -0.92113 | 15.17604 | 0.003233 | 0.036381 |
| RESP18 | -1.56307 | -0.77194 | 15.19478 | 0.003249 | 0.036542 |
| AC009021.2 | 1.069078 | 1.153825 | 15.20378 | 0.003253 | 0.036542 |
| ZNF501 | 0.840324 | 2.471317 | 15.13401 | 0.00326 | 0.036542 |
| DDN-AS1 | 1.390501 | -0.38172 | 15.15558 | 0.00326 | 0.036542 |
| ANKRD18EP | 1.250216 | 1.944728 | 15.22891 | 0.003261 | 0.036542 |
| SDCBP2-AS1 | -0.94088 | 2.167279 | 15.22876 | 0.003262 | 0.036542 |
| HNRNPH1 | -0.83066 | 8.509241 | 15.22872 | 0.003262 | 0.036542 |
| ANOS1 | 1.463772 | 4.643163 | 15.22765 | 0.003262 | 0.036542 |
| GRK6 | 0.781813 | 5.392704 | 15.2256 | 0.003264 | 0.036542 |
| STARD4-AS1 | 0.962413 | 2.832239 | 15.22396 | 0.003265 | 0.036542 |
| FAM110C | -0.97879 | 1.540893 | 15.12701 | 0.003271 | 0.036556 |

| | | | | | |
|---|---|---|---|---|---|
| PCDHA12 | 1.327036 | 1.907407 | 15.21422 | 0.003271 | 0.036556 |
| TRABD2A | 1.139694 | 1.580707 | 15.21382 | 0.003271 | 0.036556 |
| GRM6 | 1.253424 | -0.11127 | 15.107 | 0.003278 | 0.036571 |
| SPINK5 | 1.567201 | 4.101727 | 15.20224 | 0.003279 | 0.036571 |
| FAIM | -1.1514 | 4.102725 | 15.2016 | 0.003279 | 0.036571 |
| RHOT1P1 | 1.001765 | 1.17879 | 15.1041 | 0.00328 | 0.036571 |
| BEX4 | 0.604819 | 8.621229 | 15.18267 | 0.003286 | 0.036616 |
| DNAJC30 | -0.76341 | 3.587516 | 15.17657 | 0.003289 | 0.036628 |
| ST8SIA1 | 0.877967 | 6.269237 | 15.1837 | 0.003291 | 0.036628 |
| LRRC46 | -1.22544 | 2.039748 | 15.18252 | 0.003292 | 0.036628 |
| AL133216.2 | -0.89296 | 1.834182 | 15.07272 | 0.003301 | 0.036692 |
| VAMP3 | -0.92782 | 6.170885 | 15.1685 | 0.003301 | 0.036692 |
| EPHA6 | 1.253022 | 0.04818 | 15.07692 | 0.003304 | 0.036703 |
| AC010654.1 | -1.52971 | -0.46665 | 15.14171 | 0.003319 | 0.036851 |
| AP001803.2 | 1.503721 | 1.51202 | 15.12973 | 0.003327 | 0.036909 |
| NDST4 | 1.127872 | 0.987862 | 15.12598 | 0.00333 | 0.036909 |
| ZNF714 | 0.952878 | 3.204434 | 15.12453 | 0.003331 | 0.036909 |
| IDH1-AS1 | -1.79867 | -1.3881 | 15.02714 | 0.003332 | 0.036909 |
| PCDHB15 | 0.797402 | 2.359178 | 15.01768 | 0.003338 | 0.03696 |
| AL133410.1 | -1.41386 | -0.59007 | 15.01334 | 0.003341 | 0.036961 |
| DNMT3B | 0.741332 | 4.537415 | 15.10518 | 0.003344 | 0.036961 |
| RALGAPA1P1 | 0.633656 | 5.611034 | 15.10493 | 0.003344 | 0.036961 |
| SCARA3 | 1.28495 | 3.222472 | 15.103 | 0.003345 | 0.036961 |
| TAF6 | 0.881088 | 4.696324 | 15.08757 | 0.003356 | 0.037041 |
| PRKG1 | -1.24516 | 4.523369 | 15.08701 | 0.003356 | 0.037041 |
| ADAMTS8 | 1.817198 | 1.826487 | 15.07683 | 0.003363 | 0.037083 |
| AC073349.2 | 1.107867 | 0.938794 | 15.05265 | 0.003364 | 0.037083 |
| SELENOF | -0.60453 | 7.11012 | 15.0735 | 0.003366 | 0.037083 |
| RPL10P6 | -0.66275 | 5.38794 | 15.06514 | 0.003371 | 0.037108 |
| KCNK6 | -1.72027 | -0.80694 | 15.06487 | 0.003371 | 0.037108 |
| TUBB2BP1 | 1.002618 | 7.960672 | 15.04383 | 0.003386 | 0.037247 |
| SLCO1C1 | 1.666919 | 3.265759 | 15.02692 | 0.003398 | 0.037351 |
| AC008759.2 | -2.18926 | -0.41299 | 15.02482 | 0.003399 | 0.037351 |
| FOXO1 | 0.884145 | 3.70063 | 15.00352 | 0.003414 | 0.037473 |
| AC120053.1 | -1.1241 | 3.149527 | 15.00345 | 0.003414 | 0.037473 |
| LYG1 | -0.97079 | 1.600515 | 14.91164 | 0.003425 | 0.037571 |
| SMIM24 | -1.3053 | 0.925114 | 14.97444 | 0.003434 | 0.037651 |
| KRCC1 | -1.1015 | 5.134691 | 14.97234 | 0.003436 | 0.037651 |
| ITPR3 | 1.452771 | 1.485327 | 14.96653 | 0.00344 | 0.037662 |
| IL13RA1 | 0.817246 | 5.346028 | 14.96557 | 0.00344 | 0.037662 |
| EMILIN1 | -0.80425 | 3.332794 | 14.96123 | 0.003443 | 0.037662 |
| OGN | -1.80627 | -0.69258 | 14.96044 | 0.003444 | 0.037662 |
| GOLGA8T | 1.27701 | 1.537166 | 14.95751 | 0.003446 | 0.037664 |
| FKBP7 | -0.73493 | 5.334489 | 14.94632 | 0.003454 | 0.03773 |
| AC016716.2 | -1.364 | -0.12375 | 14.93825 | 0.003459 | 0.037742 |
| ZSCAN5A | -0.76501 | 2.475469 | 14.84159 | 0.003461 | 0.037742 |
| RAB34 | -0.94334 | 4.112406 | 14.93508 | 0.003462 | 0.037742 |

| | | | | | |
|---|---|---|---|---|---|
| CLDN19 | 2.053423 | -0.33669 | 14.93427 | 0.003462 | 0.037742 |
| RALYL | -1.15684 | 4.178575 | 14.9303 | 0.003465 | 0.037742 |
| CASP8 | -0.88525 | 1.911141 | 14.83502 | 0.003466 | 0.037742 |
| DOP1B | 0.748248 | 5.254818 | 14.91477 | 0.003476 | 0.037833 |
| B3GLCT | -1.15078 | 5.081701 | 14.90974 | 0.00348 | 0.037837 |
| SYT7 | 1.025859 | 6.329841 | 14.909 | 0.00348 | 0.037837 |
| LINC00623 | -0.69524 | 4.731334 | 14.90518 | 0.003483 | 0.037846 |
| RNLS | 1.875483 | -0.45109 | 14.86834 | 0.003509 | 0.038113 |
| ALDH4A1 | -0.86395 | 4.185076 | 14.8527 | 0.003521 | 0.038215 |
| ZNF132 | -0.81936 | 2.421504 | 14.75461 | 0.003524 | 0.038232 |
| CWC25 | -0.63368 | 4.513891 | 14.74835 | 0.003529 | 0.038238 |
| SUCLG2 | -1.21849 | 2.86647 | 14.83986 | 0.00353 | 0.038238 |
| ZNF75D | 0.629166 | 4.736554 | 14.74617 | 0.00353 | 0.038238 |
| CLN5 | -0.87458 | 3.980166 | 14.83387 | 0.003534 | 0.038261 |
| GPLD1 | 0.925629 | 2.794459 | 14.80356 | 0.003556 | 0.038479 |
| TMEM233 | 1.529048 | -0.70117 | 14.77011 | 0.003569 | 0.038595 |
| OTOGL | 1.503342 | 1.089245 | 14.77621 | 0.003576 | 0.038655 |
| CTDSP1 | -0.98231 | 4.014936 | 14.77016 | 0.003581 | 0.038683 |
| AC093218.1 | -1.28156 | 0.90396 | 14.75992 | 0.003588 | 0.038744 |
| VEGFA | 0.642672 | 7.561574 | 14.75254 | 0.003594 | 0.038768 |
| TMEM230 | -0.68282 | 6.78778 | 14.7518 | 0.003594 | 0.038768 |
| NEURL1 | 1.018942 | 4.855816 | 14.74811 | 0.003597 | 0.038777 |
| RPL10P3 | 1.619716 | -0.9731 | 14.68338 | 0.003607 | 0.038869 |
| SHISAL1 | 0.813392 | 6.140889 | 14.72822 | 0.003612 | 0.038877 |
| FXYD6 | 0.712848 | 7.763556 | 14.7279 | 0.003612 | 0.038877 |
| AC066613.1 | 1.65318 | -0.49731 | 14.72059 | 0.003617 | 0.038916 |
| HAPLN3 | 1.536607 | 1.210676 | 14.71415 | 0.003622 | 0.038947 |
| LRRC32 | 1.239338 | 3.094235 | 14.70563 | 0.003629 | 0.038976 |
| SULT1A1 | -1.28255 | 0.246755 | 14.70542 | 0.003629 | 0.038976 |
| ZPLD1 | 2.03954 | 0.881377 | 14.6888 | 0.003641 | 0.039062 |
| NEK5 | -1.48047 | 2.276682 | 14.68874 | 0.003641 | 0.039062 |
| AC024896.1 | -1.13357 | 3.971827 | 14.68706 | 0.003643 | 0.039062 |
| TFPI2 | -1.10763 | 1.34448 | 14.68429 | 0.003645 | 0.039064 |
| UAP1L1 | 0.951261 | 4.134389 | 14.67687 | 0.00365 | 0.039104 |
| IL17RC | -1.11041 | 3.00587 | 14.671 | 0.003655 | 0.039131 |
| B3GNT9 | -1.11792 | 2.986636 | 14.6655 | 0.003659 | 0.039145 |
| ALPL | 1.983146 | 2.473022 | 14.66225 | 0.003661 | 0.039145 |
| KDM6A | 0.59003 | 5.1744 | 14.57034 | 0.003662 | 0.039145 |
| TCP11L2 | 0.735806 | 2.538116 | 14.55339 | 0.003675 | 0.039253 |
| TP53I11 | 0.84728 | 7.189004 | 14.64323 | 0.003676 | 0.039253 |
| MUC20P1 | 1.295283 | 0.780529 | 14.63694 | 0.00368 | 0.039284 |
| CYP51A1P2 | 0.808361 | 4.398071 | 14.63204 | 0.003684 | 0.03929 |
| C17orf51 | -0.60712 | 4.316346 | 14.54014 | 0.003685 | 0.03929 |
| ANKRD18A | 1.270846 | 0.918529 | 14.62847 | 0.003687 | 0.039291 |
| BEST1 | 0.912057 | 2.886161 | 14.62062 | 0.003693 | 0.039335 |
| HLA-DMB | 1.07918 | 0.755561 | 14.53195 | 0.003698 | 0.039366 |
| SNX24 | -0.67353 | 4.634773 | 14.61174 | 0.0037 | 0.039366 |

| | | | | | |
|---|---|---|---|---|---|
| KRT18 | 0.719767 | 3.646961 | 14.53438 | 0.003705 | 0.039405 |
| CMTM7 | -1.30204 | 4.685035 | 14.59833 | 0.00371 | 0.039435 |
| GPR158 | 0.651661 | 4.393831 | 14.49607 | 0.00372 | 0.039521 |
| ATP2B4 | 0.586108 | 8.138737 | 14.57267 | 0.003726 | 0.03956 |
| ACO2 | 0.887809 | 5.531791 | 14.56859 | 0.003733 | 0.039577 |
| AL049840.4 | -1.25939 | 1.750574 | 14.56451 | 0.003736 | 0.039577 |
| RGL3 | -1.30779 | 1.617668 | 14.56387 | 0.003737 | 0.039577 |
| CLTCL1 | 0.82703 | 3.715427 | 14.56039 | 0.003739 | 0.039577 |
| ARHGAP5 | 0.691457 | 6.204489 | 14.55913 | 0.00374 | 0.039577 |
| TRIM29 | 1.258588 | 1.463944 | 14.55849 | 0.003741 | 0.039577 |
| SNX25 | 0.536968 | 5.601261 | 14.46753 | 0.003741 | 0.039577 |
| SLC38A6 | -0.85568 | 3.79927 | 14.556 | 0.003743 | 0.039577 |
| LINC02381 | 1.245872 | 2.338956 | 14.5479 | 0.003749 | 0.039623 |
| CPXM1 | 0.589744 | 5.410808 | 14.48409 | 0.003751 | 0.039623 |
| SNN | 0.666424 | 7.913301 | 14.53861 | 0.003756 | 0.039649 |
| ZNF853 | 0.679889 | 6.82894 | 14.53726 | 0.003757 | 0.039649 |
| BNIP3P1 | 0.693716 | 6.260397 | 14.53066 | 0.003762 | 0.039683 |
| ERICH3 | 1.074708 | 3.225236 | 14.52505 | 0.003767 | 0.039708 |
| FUNDC1 | 0.546361 | 5.757707 | 14.43283 | 0.003769 | 0.039708 |
| AL450326.2 | -1.23613 | 1.029258 | 14.51603 | 0.003774 | 0.039743 |
| NPTN-IT1 | 1.420214 | 1.33562 | 14.50844 | 0.00378 | 0.039755 |
| SLF1 | -0.70983 | 4.423811 | 14.50815 | 0.00378 | 0.039755 |
| TCEANC | 0.815051 | 3.023672 | 14.50721 | 0.003781 | 0.039755 |
| WDFY4 | 2.025065 | 0.303151 | 14.50038 | 0.003786 | 0.039791 |
| IGF2BP2 | -0.78387 | 5.977226 | 14.49037 | 0.003794 | 0.039853 |
| NES | 0.814067 | 7.412596 | 14.4719 | 0.003809 | 0.039967 |
| SLC25A5P2 | 1.07447 | 1.939985 | 14.47173 | 0.003809 | 0.039967 |
| PRKRIP1 | -0.55199 | 5.574572 | 14.3669 | 0.003821 | 0.040068 |
| LONRF3 | 1.523836 | 1.158052 | 14.45479 | 0.003822 | 0.040068 |
| SPCS2 | -0.55007 | 6.742709 | 14.37838 | 0.003826 | 0.040085 |
| MIR3682 | -1.28296 | -0.51996 | 14.35525 | 0.003831 | 0.040114 |
| LYPD3 | 1.054909 | 1.210534 | 14.43864 | 0.003835 | 0.040141 |
| ALDOA | 1.55271 | 3.227382 | 14.43132 | 0.003841 | 0.04015 |
| CYS1 | -1.79945 | 0.371451 | 14.43088 | 0.003842 | 0.04015 |
| ANKRD20A10P | 1.402745 | -0.32894 | 14.43018 | 0.003842 | 0.04015 |
| SEC24C | 0.847543 | 4.543913 | 14.39605 | 0.00387 | 0.040417 |
| NEXN | -1.868 | 1.912115 | 14.38838 | 0.003876 | 0.040457 |
| DND1P1 | 2.488558 | -1.19464 | 14.38468 | 0.003879 | 0.040457 |
| UBB | 0.710178 | 6.934693 | 14.38087 | 0.003882 | 0.040457 |
| NUDT9 | -0.69874 | 5.962913 | 14.37949 | 0.003883 | 0.040457 |
| LINC01224 | 1.504996 | 1.829866 | 14.3769 | 0.003885 | 0.040457 |
| TRIM73 | -0.90307 | 2.475813 | 14.37676 | 0.003885 | 0.040457 |
| OLFML2B | -1.68794 | 4.14635 | 14.37281 | 0.003888 | 0.04047 |
| RNF144A | 0.656228 | 6.744866 | 14.36679 | 0.003893 | 0.0405 |
| ZNF660 | -0.87275 | 3.693493 | 14.34935 | 0.003908 | 0.040616 |
| CDK18 | 1.076388 | 3.570578 | 14.34829 | 0.003908 | 0.040616 |
| CCNQP2 | 1.763299 | -0.9584 | 14.33937 | 0.003916 | 0.040664 |

| | | | | | |
|---|---|---|---|---|---|
| MRAP2 | 0.873123 | 4.727898 | 14.33767 | 0.003917 | 0.040664 |
| PCDHB6 | 0.753567 | 2.962011 | 14.24748 | 0.003919 | 0.040664 |
| AL158151.4 | -1.20076 | 0.469508 | 14.31252 | 0.003938 | 0.040833 |
| AC084018.1 | 1.388289 | -0.37993 | 14.2989 | 0.003939 | 0.040833 |
| IQCA1 | 0.94659 | 2.13739 | 14.29889 | 0.003949 | 0.040912 |
| KCTD12 | -0.69606 | 7.729325 | 14.29675 | 0.003951 | 0.040912 |
| AC006466.1 | 1.403374 | -0.16648 | 14.29393 | 0.003953 | 0.040916 |
| SLC1A5 | 1.059048 | 4.460086 | 14.2798 | 0.003965 | 0.041017 |
| RAB11FIP5 | 0.949483 | 4.872434 | 14.27298 | 0.003971 | 0.041047 |
| RARA-AS1 | -0.9059 | 1.357846 | 14.18419 | 0.003972 | 0.041047 |
| RF00019 | -0.77139 | 2.908602 | 14.18015 | 0.003975 | 0.041061 |
| HOXB3 | -0.8534 | 5.004791 | 14.26292 | 0.003979 | 0.041062 |
| SLC25A34 | -0.90065 | 1.201498 | 14.17535 | 0.003979 | 0.041062 |
| ZFPM2-AS1 | 1.367089 | 1.780728 | 14.25798 | 0.003983 | 0.041068 |
| GPR183 | -1.81944 | 0.736165 | 14.25708 | 0.003984 | 0.041068 |
| ZNF18 | -0.80081 | 3.962877 | 14.25143 | 0.003989 | 0.041096 |
| AC010735.2 | -1.66341 | -0.99987 | 14.24709 | 0.003992 | 0.041113 |
| SLC25A15 | 0.858229 | 3.121394 | 14.2413 | 0.003997 | 0.041133 |
| PCDH17 | 0.847477 | 5.659733 | 14.23964 | 0.003999 | 0.041133 |
| AC079385.2 | -2.12615 | -0.46476 | 14.23759 | 0.004 | 0.041133 |
| TRIM6 | -1.36378 | 0.688194 | 14.23477 | 0.004003 | 0.041137 |
| AL512283.1 | 1.537453 | -1.09722 | 14.12878 | 0.004019 | 0.04128 |
| AL031658.2 | 1.002813 | 2.810714 | 14.21351 | 0.004021 | 0.04128 |
| ENPP6 | 1.358845 | -0.49224 | 14.12906 | 0.004025 | 0.041295 |
| LINC00632 | 0.818003 | 6.395679 | 14.20622 | 0.004027 | 0.041295 |
| MSX1 | -1.29717 | 0.635492 | 14.20469 | 0.004028 | 0.041295 |
| CHST15 | 1.267678 | 3.033014 | 14.20072 | 0.004032 | 0.041309 |
| AL592295.1 | 1.428941 | -0.3537 | 14.19606 | 0.004036 | 0.041329 |
| AC073869.3 | -1.19379 | 0.489694 | 14.17564 | 0.004053 | 0.041487 |
| TENM2 | 0.698979 | 5.517608 | 14.16722 | 0.00406 | 0.041525 |
| PTPRH | 0.792282 | 6.164657 | 14.16652 | 0.004061 | 0.041525 |
| CMTM1 | 1.302817 | 1.084931 | 14.15436 | 0.004071 | 0.041602 |
| C15orf39 | 1.034513 | 3.832279 | 14.15018 | 0.004075 | 0.041602 |
| AC000089.1 | -0.85992 | 5.061238 | 14.14788 | 0.004077 | 0.041602 |
| AZU1 | -1.09388 | 2.325844 | 14.14628 | 0.004078 | 0.041602 |
| FOXS1 | 2.957786 | -0.47874 | 14.14594 | 0.004079 | 0.041602 |
| ZNF710 | 1.006382 | 4.158153 | 14.13973 | 0.004084 | 0.041602 |
| LAPTM4B | 0.613529 | 7.661658 | 14.13915 | 0.004084 | 0.041602 |
| TPT1 | -0.81019 | 9.77357 | 14.1364 | 0.004087 | 0.041602 |
| AGPAT4-IT1 | -1.25339 | 0.100691 | 14.12915 | 0.004088 | 0.041602 |
| SLC1A4 | 0.646497 | 6.962749 | 14.13426 | 0.004089 | 0.041602 |
| LINC01006 | 0.703595 | 3.049948 | 14.03909 | 0.004097 | 0.041663 |
| NPM1P27 | -0.60942 | 6.556314 | 14.12029 | 0.004101 | 0.041684 |
| MFSD4A | 0.914207 | 2.781273 | 14.11697 | 0.004104 | 0.041692 |
| DOCK5 | -1.67544 | 2.335511 | 14.10656 | 0.004113 | 0.041763 |
| LGR5 | 1.436709 | -0.3061 | 14.10018 | 0.004118 | 0.041799 |
| ACSL1 | 0.554713 | 5.763978 | 14.00777 | 0.004125 | 0.041845 |

| | | | | | |
|---|---|---|---|---|---|
| PPP1R16B | 0.838034 | 4.786272 | 14.07696 | 0.004139 | 0.041964 |
| UPF3A | -0.75147 | 6.327548 | 14.05841 | 0.004155 | 0.042109 |
| CD63 | -0.5734 | 8.533997 | 14.02678 | 0.004158 | 0.042114 |
| LINC02615 | -1.55833 | 2.02626 | 14.0532 | 0.00416 | 0.042114 |
| AC107982.3 | -1.10194 | 1.188339 | 14.03948 | 0.004172 | 0.042216 |
| GPR157 | 1.017695 | 2.527376 | 14.03549 | 0.004175 | 0.042219 |
| NECAB1 | 0.713542 | 2.749864 | 13.94951 | 0.004176 | 0.042219 |
| GRM7 | 0.654077 | 4.432938 | 13.98025 | 0.004185 | 0.042291 |
| LINC01956 | 1.641707 | 0.187702 | 14.02087 | 0.004188 | 0.042299 |
| SELENOT | -0.61948 | 7.589625 | 14.01617 | 0.004192 | 0.042321 |
| DNAH12 | -1.64521 | 0.86336 | 14.01365 | 0.004195 | 0.042323 |
| HEXIM1 | -0.66778 | 5.752572 | 14.00651 | 0.004201 | 0.042366 |
| FOXJ1 | -1.79879 | 4.961686 | 13.99097 | 0.004215 | 0.042486 |
| SLC35A2 | 0.589771 | 5.320767 | 13.94088 | 0.004219 | 0.042508 |
| AC004987.2 | 0.63174 | 5.118267 | 13.98362 | 0.004222 | 0.04251 |
| IFT80 | -1.03965 | 5.367148 | 13.97649 | 0.004228 | 0.042554 |
| PDK1 | 1.391445 | 6.16823 | 13.97176 | 0.004232 | 0.042559 |
| AF067845.4 | 1.767238 | -0.54225 | 13.97094 | 0.004233 | 0.042559 |
| P2RY1 | 1.142067 | 3.041055 | 13.96679 | 0.004237 | 0.042559 |
| CCDC198 | -1.37677 | 1.661399 | 13.96678 | 0.004237 | 0.042559 |
| CLSTN2 | 0.571895 | 7.384034 | 13.95826 | 0.004244 | 0.042603 |
| ZFP36L1 | -0.94701 | 7.401988 | 13.95633 | 0.004246 | 0.042603 |
| WDR82P2 | 1.161821 | 2.028818 | 13.95505 | 0.004247 | 0.042603 |
| LINC02591 | -0.93136 | 2.637222 | 13.94564 | 0.004256 | 0.042666 |
| PDE7B | 1.279647 | 2.768663 | 13.941 | 0.00426 | 0.042666 |
| GCNA | -0.84938 | 2.762048 | 13.93937 | 0.004262 | 0.042666 |
| IMPDH1P5 | 1.431428 | 0.313381 | 13.9389 | 0.004262 | 0.042666 |
| TCEAL2 | -0.77505 | 6.477524 | 13.93637 | 0.004264 | 0.042668 |
| XPOTP1 | 0.867612 | 2.850855 | 13.93202 | 0.004268 | 0.042669 |
| AC073857.1 | -0.93636 | 1.578284 | 13.86414 | 0.004269 | 0.042669 |
| TAC1 | 1.57941 | 6.620352 | 13.92525 | 0.004274 | 0.042707 |
| LINC01828 | 1.772954 | 0.107569 | 13.91571 | 0.004283 | 0.042773 |
| ZNF77 | -0.72478 | 2.981065 | 13.82225 | 0.004292 | 0.042842 |
| SOGA1 | 0.611659 | 7.305078 | 13.89573 | 0.004302 | 0.042915 |
| GRIP2 | 0.643452 | 5.054853 | 13.88525 | 0.004311 | 0.04299 |
| AC118344.2 | -0.98569 | 0.49301 | 13.79851 | 0.004314 | 0.043 |
| AC112484.3 | -0.77954 | 1.94802 | 13.78309 | 0.004329 | 0.043123 |
| CLIC4P3 | 1.55991 | -0.14356 | 13.85785 | 0.004337 | 0.04318 |
| LRRN1 | 0.544459 | 5.861682 | 13.76207 | 0.004348 | 0.043277 |
| CSNK2B | -0.63177 | 5.702747 | 13.83925 | 0.004354 | 0.04331 |
| FOXM1 | 1.487403 | -0.14729 | 13.83674 | 0.004356 | 0.043313 |
| MFSD14A | 1.069841 | 1.221754 | 13.834 | 0.004359 | 0.043317 |
| HSPA12B | 1.513279 | 0.250804 | 13.81607 | 0.004376 | 0.043449 |
| AC005261.1 | -0.807 | 6.661853 | 13.81536 | 0.004376 | 0.043449 |
| XRCC1 | -0.72382 | 5.342576 | 13.80131 | 0.004389 | 0.043559 |
| ADCY7 | 1.363758 | 3.725888 | 13.79065 | 0.0044 | 0.043638 |
| LINC02526 | -1.58377 | -0.83718 | 13.78316 | 0.004407 | 0.043687 |

| | | | | | |
|---|---|---|---|---|---|
| AFG3L1P | -0.81006 | 4.298335 | 13.78001 | 0.00441 | 0.043695 |
| TDRKH | 0.608624 | 4.342108 | 13.68866 | 0.004418 | 0.043759 |
| GINS2 | 0.960364 | 1.123088 | 13.68439 | 0.004422 | 0.043779 |
| MPV17 | -0.64883 | 5.580005 | 13.76127 | 0.004427 | 0.043808 |
| TUBB8P12 | 1.124911 | -0.19716 | 13.67183 | 0.004434 | 0.043856 |
| COBL | 1.258248 | 2.895166 | 13.75174 | 0.004437 | 0.043856 |
| FSTL1 | -0.83023 | 9.283264 | 13.7471 | 0.004441 | 0.043861 |
| BPGM | -0.55224 | 6.336593 | 13.70411 | 0.004443 | 0.043861 |
| AC091849.1 | 1.036737 | 0.561925 | 13.65867 | 0.004447 | 0.043861 |
| DOCK6 | 0.687683 | 4.966607 | 13.73925 | 0.004449 | 0.043861 |
| USP44 | 1.076374 | 0.527887 | 13.6571 | 0.004449 | 0.043861 |
| CFAP44 | -1.07866 | 4.872353 | 13.73774 | 0.00445 | 0.043861 |
| AL080317.3 | -1.00243 | 1.575621 | 13.73412 | 0.004453 | 0.043873 |
| TFPT | -0.68637 | 3.876553 | 13.68846 | 0.004456 | 0.043873 |
| ETNK2 | 0.865043 | 6.108555 | 13.72983 | 0.004458 | 0.043873 |
| WNT7B | 2.547383 | -0.89708 | 13.72631 | 0.004461 | 0.043877 |
| OSBPL6 | -0.56348 | 6.336066 | 13.71768 | 0.004465 | 0.043877 |
| AL845472.1 | 0.929784 | 3.460706 | 13.72203 | 0.004465 | 0.043877 |
| AL450311.2 | 1.561606 | -0.93662 | 13.69747 | 0.004467 | 0.043877 |
| LINC00858 | 1.091393 | 0.20069 | 13.62978 | 0.004475 | 0.043941 |
| MRPS9 | -0.68897 | 5.203625 | 13.69835 | 0.004488 | 0.043977 |
| SIX4 | -0.93282 | 2.156388 | 13.69661 | 0.00449 | 0.043977 |
| MYL12A | -0.65303 | 5.692103 | 13.69409 | 0.004492 | 0.043977 |
| AL359924.1 | 1.739051 | -1.33735 | 13.64622 | 0.004493 | 0.043977 |
| CTCFL | -1.51266 | 2.631221 | 13.69261 | 0.004494 | 0.043977 |
| SNAP23 | -0.93801 | 2.827299 | 13.69195 | 0.004494 | 0.043977 |
| PRKAA1 | -0.71075 | 5.748652 | 13.69061 | 0.004495 | 0.043977 |
| RNF157 | 0.769101 | 7.326157 | 13.68996 | 0.004496 | 0.043977 |
| LARGE1 | 0.596491 | 5.764211 | 13.68653 | 0.004499 | 0.043977 |
| MTATP6P26 | -1.1866 | -0.1396 | 13.60408 | 0.0045 | 0.043977 |
| AKAP14 | 1.355056 | -0.22737 | 13.68214 | 0.004504 | 0.043988 |
| TMEM132A | 0.976222 | 6.887753 | 13.67695 | 0.004509 | 0.044002 |
| AMYP1 | -1.62551 | 2.098502 | 13.67626 | 0.004509 | 0.044002 |
| MAPK14 | -0.63111 | 6.288697 | 13.67265 | 0.004513 | 0.044012 |
| FGF7P8 | 1.523139 | 1.460172 | 13.67086 | 0.004515 | 0.044012 |
| TUBBP1 | 0.865393 | 5.896837 | 13.65446 | 0.004531 | 0.044147 |
| CCS | -0.71684 | 4.945405 | 13.64547 | 0.00454 | 0.044212 |
| ARG2 | 0.727637 | 4.741473 | 13.63972 | 0.004545 | 0.044246 |
| AC124303.1 | 1.883718 | -1.24524 | 13.63582 | 0.004549 | 0.044262 |
| AC005224.4 | 1.448423 | 0.291858 | 13.62759 | 0.004557 | 0.04432 |
| FUNDC2P2 | 1.014048 | 1.950489 | 13.62503 | 0.00456 | 0.044324 |
| ZNF396 | -0.7986 | 2.842074 | 13.60865 | 0.004566 | 0.044361 |
| NDUFAF1 | -0.55761 | 4.369542 | 13.52595 | 0.004578 | 0.044459 |
| VTCN1 | 1.313227 | 0.832844 | 13.60222 | 0.004582 | 0.04448 |
| ATP6V1G1 | -0.62838 | 7.652979 | 13.59186 | 0.004593 | 0.044559 |
| MEGF6 | -0.86917 | 5.120427 | 13.58574 | 0.004599 | 0.044597 |
| CAMK2A | 0.787161 | 4.625661 | 13.57981 | 0.004605 | 0.044625 |

| | | | | | |
|---|---|---|---|---|---|
| RBM18 | -0.6127 | 6.24487 | 13.57846 | 0.004606 | 0.044625 |
| CTSC | 1.104051 | 2.059446 | 13.55976 | 0.004625 | 0.044757 |
| RPL21 | -0.68456 | 6.657117 | 13.55899 | 0.004625 | 0.044757 |
| AC068491.1 | 1.016538 | 2.839694 | 13.55841 | 0.004626 | 0.044757 |
| LINC01801 | 1.664601 | 1.723439 | 13.54548 | 0.004639 | 0.044861 |
| PPM1J | 1.079665 | 2.151391 | 13.53823 | 0.004646 | 0.044911 |
| PI4KAP1 | -0.61623 | 5.453127 | 13.53519 | 0.004649 | 0.04492 |
| AC009509.4 | -1.66411 | -0.41206 | 13.52304 | 0.004662 | 0.044997 |
| SERPINF1 | -1.04345 | 5.75881 | 13.52296 | 0.004662 | 0.044997 |
| LRP3 | 1.154551 | 5.996624 | 13.5136 | 0.004671 | 0.045067 |
| MTMR8 | 1.133716 | 0.457126 | 13.49893 | 0.004675 | 0.045084 |
| PWAR1 | 1.311958 | 0.566745 | 13.50687 | 0.004678 | 0.045085 |
| IL31RA | 1.069789 | 2.400038 | 13.50509 | 0.00468 | 0.045085 |
| FTLP3 | 0.525386 | 6.547637 | 13.42353 | 0.004682 | 0.045085 |
| LINC01166 | 1.543175 | 0.650515 | 13.4995 | 0.004686 | 0.0451 |
| ZNF561-AS1 | -0.79713 | 3.200109 | 13.49432 | 0.004691 | 0.04513 |
| CKAP2L | 1.579841 | -0.44426 | 13.48934 | 0.004696 | 0.045157 |
| ANXA5 | -0.84282 | 7.61466 | 13.4793 | 0.004706 | 0.045235 |
| AC009244.1 | -0.83502 | 3.516665 | 13.46734 | 0.004719 | 0.045315 |
| THSD1 | -0.78532 | 1.78953 | 13.38759 | 0.004719 | 0.045315 |
| SLC47A2 | -1.13635 | 0.721453 | 13.45365 | 0.004733 | 0.045425 |
| PDCD6 | -0.56822 | 6.11156 | 13.43773 | 0.004749 | 0.045562 |
| COQ6 | 0.760539 | 2.018214 | 13.35391 | 0.004754 | 0.045589 |
| SLC24A2 | 0.732813 | 4.025866 | 13.42083 | 0.004767 | 0.045665 |
| FANCE | 0.814254 | 3.481238 | 13.4175 | 0.00477 | 0.045665 |
| STXBP5L | 1.049158 | 4.0788 | 13.41713 | 0.004771 | 0.045665 |
| AC009312.1 | 1.487836 | -1.05593 | 13.33794 | 0.004771 | 0.045665 |
| RPL36AL | -0.65537 | 6.585718 | 13.39639 | 0.004793 | 0.045838 |
| CD164 | -1.00884 | 7.457095 | 13.39511 | 0.004794 | 0.045838 |
| ART5 | 1.235735 | -0.51801 | 13.3125 | 0.004798 | 0.045852 |
| GPRIN1 | 0.884576 | 6.121298 | 13.38954 | 0.0048 | 0.045852 |
| HOXB-AS1 | -1.17595 | 1.107422 | 13.38301 | 0.004807 | 0.045888 |
| NSFP1 | 0.945467 | 4.240715 | 13.38171 | 0.004808 | 0.045888 |
| DENND1C | 1.56336 | 3.127714 | 13.37854 | 0.004811 | 0.045898 |
| SPRYD4 | -0.65279 | 3.610195 | 13.29302 | 0.004819 | 0.045928 |
| CYP51A1 | 1.254048 | 2.20073 | 13.37016 | 0.00482 | 0.045928 |
| AP000911.3 | 1.359627 | -0.80179 | 13.29085 | 0.004821 | 0.045928 |
| SIAH3 | 0.72175 | 2.926519 | 13.28846 | 0.004824 | 0.045931 |
| AMY2A | -1.93863 | 3.491366 | 13.36024 | 0.004831 | 0.045956 |
| AC244453.2 | -0.94591 | 0.659456 | 13.28184 | 0.004831 | 0.045956 |
| GNAS | 0.628115 | 11.52657 | 13.32295 | 0.004837 | 0.045994 |
| DGAT2 | 1.112441 | 3.499299 | 13.34959 | 0.004842 | 0.046017 |
| PALD1 | 1.055323 | 3.821294 | 13.34765 | 0.004844 | 0.046017 |
| APOC1 | -1.28667 | 0.895007 | 13.34133 | 0.004851 | 0.04606 |
| KIF6 | 0.817996 | 1.722751 | 13.26015 | 0.004854 | 0.04607 |
| C1QBP | -0.52532 | 6.748099 | 13.26457 | 0.004857 | 0.046072 |
| GRIN3B | 1.99216 | 1.033469 | 13.33379 | 0.004859 | 0.046072 |

| | | | | | |
|---|---|---|---|---|---|
| SPATA6 | -0.83222 | 4.577269 | 13.32962 | 0.004863 | 0.046093 |
| FNBP4 | -0.60587 | 6.193081 | 13.32627 | 0.004867 | 0.046105 |
| PCYT1B | 0.753497 | 4.446421 | 13.32158 | 0.004872 | 0.046132 |
| PTPRN | 0.927425 | 8.821018 | 13.30834 | 0.004886 | 0.046197 |
| ETFRF1 | -0.95616 | 5.241549 | 13.3057 | 0.004889 | 0.046197 |
| BAIAP2-DT | 0.558051 | 5.248375 | 13.22797 | 0.004889 | 0.046197 |
| FGFBP3 | 0.551078 | 7.111143 | 13.30563 | 0.004889 | 0.046197 |
| GALNT16 | 0.758292 | 5.312918 | 13.30388 | 0.004891 | 0.046197 |
| ABCG8 | 1.212585 | -0.24912 | 13.22288 | 0.004894 | 0.046197 |
| HEYL | 1.324268 | 1.105063 | 13.29935 | 0.004896 | 0.046197 |
| SYPL1 | -1.22583 | 4.583615 | 13.29806 | 0.004897 | 0.046197 |
| UBE2HP1 | 1.73404 | -0.29288 | 13.29559 | 0.0049 | 0.046197 |
| IRX3 | -1.09938 | 1.868024 | 13.29419 | 0.004901 | 0.046197 |
| ZBTB7C | 0.800957 | 2.85226 | 13.28359 | 0.004913 | 0.046283 |
| SQOR | -1.0523 | 0.515498 | 13.19901 | 0.004921 | 0.046316 |
| KLK6 | 1.46879 | -1.22401 | 13.1977 | 0.004922 | 0.046316 |
| AC018693.2 | 1.945835 | -0.92104 | 13.27414 | 0.004923 | 0.046316 |
| ELF1 | -0.83146 | 4.775939 | 13.27018 | 0.004927 | 0.046316 |
| VAMP5 | 1.189006 | 1.562706 | 13.26992 | 0.004928 | 0.046316 |
| AC122688.3 | 1.225085 | -0.35816 | 13.18639 | 0.004934 | 0.046359 |
| ARPC1A | -0.51725 | 5.445307 | 13.17986 | 0.004942 | 0.046405 |
| TTC8 | -0.75679 | 5.345816 | 13.24943 | 0.00495 | 0.046441 |
| AC074043.1 | 1.060795 | 0.996862 | 13.24834 | 0.004951 | 0.046441 |
| C2CD2L | 0.6481 | 5.205539 | 13.24502 | 0.004955 | 0.046441 |
| ZFYVE9 | 0.49497 | 6.192892 | 13.16586 | 0.004957 | 0.046441 |
| KCNQ5 | 1.073504 | 0.737187 | 13.23438 | 0.004958 | 0.046441 |
| AL645608.2 | 1.325823 | 0.668276 | 13.24111 | 0.004959 | 0.046441 |
| KANK4 | 1.036396 | 2.488423 | 13.23518 | 0.004965 | 0.046462 |
| MB21D2 | 0.679419 | 5.157002 | 13.23494 | 0.004966 | 0.046462 |
| SRP72P1 | 1.187726 | -0.22421 | 13.15054 | 0.004974 | 0.046517 |
| TGFA | 1.072894 | 0.505488 | 13.16181 | 0.004978 | 0.046534 |
| TRIM65 | -0.80063 | 3.537335 | 13.21544 | 0.004987 | 0.046598 |
| CREG1 | 0.516493 | 6.116381 | 13.1312 | 0.004995 | 0.046636 |
| AC034231.1 | 2.026673 | -0.78178 | 13.20765 | 0.004996 | 0.046636 |
| PRODH | 1.047002 | 0.624413 | 13.13618 | 0.004998 | 0.046636 |
| SOWAHA | 1.05801 | 1.969242 | 13.19994 | 0.005004 | 0.046673 |
| FOXO3B | 1.475478 | -0.00596 | 13.19241 | 0.005013 | 0.046729 |
| ZFAS1 | -0.57485 | 4.687793 | 13.11299 | 0.005016 | 0.046738 |
| ABCC8 | 1.051235 | 2.891706 | 13.18444 | 0.005021 | 0.046769 |
| MAP1B | 0.754875 | 12.01089 | 13.1789 | 0.005028 | 0.046787 |
| AF129075.1 | -1.22875 | -0.60476 | 13.10218 | 0.005028 | 0.046787 |
| WNT11 | -1.13424 | 1.99833 | 13.17364 | 0.005033 | 0.046801 |
| MSRB3 | -0.83998 | 3.232531 | 13.17317 | 0.005034 | 0.046801 |
| AC091544.4 | -1.14772 | 0.92563 | 13.17057 | 0.005037 | 0.046806 |
| WNT7A | 1.411031 | 2.580333 | 13.16363 | 0.005045 | 0.046857 |
| UTS2 | -1.33449 | 8.374823 | 13.15905 | 0.00505 | 0.046883 |
| C3orf38 | -0.71137 | 5.070146 | 13.1447 | 0.005066 | 0.047001 |

| | | | | | |
|---|---|---|---|---|---|
| GNG12-AS1 | -1.46209 | 0.397926 | 13.14357 | 0.005067 | 0.047001 |
| AC116562.4 | 1.571962 | -0.67479 | 13.1416 | 0.005069 | 0.047001 |
| ST8SIA6 | 1.003299 | 1.03479 | 13.09833 | 0.005074 | 0.047012 |
| FZD9 | 1.322276 | 3.612212 | 13.13647 | 0.005075 | 0.047012 |
| SNX3 | -0.62623 | 7.527714 | 13.13384 | 0.005078 | 0.047018 |
| SMAP1 | 0.935769 | 4.590821 | 13.13113 | 0.005081 | 0.047018 |
| CCDC8 | -1.00339 | 5.00806 | 13.12977 | 0.005083 | 0.047018 |
| WTIP | 0.7417 | 3.993213 | 13.12455 | 0.005089 | 0.047051 |
| FBLN1 | 1.032035 | 5.044992 | 13.12083 | 0.005093 | 0.047069 |
| LINC00863 | 0.83259 | 2.563236 | 13.11026 | 0.005105 | 0.047158 |
| PCDHA3 | 1.789257 | -0.30492 | 13.1059 | 0.00511 | 0.047183 |
| AL021807.1 | -1.65531 | -1.36233 | 13.027 | 0.005113 | 0.047195 |
| SLC26A8 | 1.594676 | -0.33406 | 13.09958 | 0.005117 | 0.047206 |
| ZNF41 | 0.501818 | 5.309431 | 13.01215 | 0.00513 | 0.04731 |
| AC005052.1 | 1.192659 | 2.192698 | 13.08303 | 0.005136 | 0.047338 |
| TEX26 | -1.11658 | 2.504839 | 13.07665 | 0.005143 | 0.047383 |
| NINL | -0.4922 | 5.769556 | 12.99056 | 0.005155 | 0.047475 |
| AP005229.2 | 2.107733 | -0.48792 | 13.05525 | 0.005167 | 0.047567 |
| ALCAM | -1.33607 | 10.26564 | 13.04872 | 0.005175 | 0.047578 |
| PODXL2 | 0.750586 | 8.081345 | 13.04863 | 0.005175 | 0.047578 |
| IPO11 | -1.17552 | 4.95503 | 13.04808 | 0.005176 | 0.047578 |
| HMGA1P1 | 0.897314 | 1.274525 | 12.97097 | 0.005178 | 0.047578 |
| SOX1-OT | 1.889019 | 0.37896 | 13.0432 | 0.005181 | 0.047578 |
| SLC43A2 | 0.794074 | 4.74273 | 13.04137 | 0.005183 | 0.047578 |
| DOCK2 | 1.164132 | 0.343542 | 13.04013 | 0.005185 | 0.047578 |
| ESPL1 | 1.933923 | -0.92768 | 13.0371 | 0.005188 | 0.047588 |
| FRMD4A | 0.663732 | 5.147234 | 13.03231 | 0.005194 | 0.047613 |
| AGAP2 | 0.957398 | 5.621346 | 13.02953 | 0.005197 | 0.047613 |
| AF287957.1 | -0.80765 | 1.910961 | 12.95373 | 0.005198 | 0.047613 |
| TSGA10 | -0.94166 | 2.727949 | 13.01643 | 0.005212 | 0.047723 |
| RTL8C | 0.666291 | 8.362224 | 12.99737 | 0.005235 | 0.047886 |
| MHENCR | -0.72324 | 2.816903 | 12.92237 | 0.005235 | 0.047886 |
| DNAJC1 | -0.80063 | 4.418052 | 12.99051 | 0.005243 | 0.047935 |
| CELP | -1.78271 | -0.99805 | 12.96677 | 0.00527 | 0.048169 |
| STIM2 | -0.62897 | 5.951075 | 12.94939 | 0.005291 | 0.048335 |
| HSPD1P6 | -1.08747 | 0.956079 | 12.94598 | 0.005295 | 0.04835 |
| C2orf70 | 1.841369 | 0.05609 | 12.94277 | 0.005299 | 0.048363 |
| TMEM145 | 0.73029 | 4.788719 | 12.93591 | 0.005307 | 0.048416 |
| ASAP2 | 0.6142 | 5.075713 | 12.93367 | 0.00531 | 0.048419 |
| AC092335.1 | 1.603696 | -0.83978 | 12.93116 | 0.005313 | 0.048422 |
| RBM6 | -0.71864 | 7.474568 | 12.92938 | 0.005315 | 0.048422 |
| NUTM2D | 1.146199 | 2.773758 | 12.92407 | 0.005321 | 0.048448 |
| ADIRF-AS1 | -1.15956 | -0.12015 | 12.84903 | 0.005322 | 0.048448 |
| AC018665.1 | 1.616126 | -1.13243 | 12.91173 | 0.005325 | 0.048455 |
| NIPA1 | 0.554922 | 7.27886 | 12.91556 | 0.005331 | 0.048472 |
| MTUS1 | -0.70653 | 5.017497 | 12.91305 | 0.005334 | 0.048472 |
| DBIL5P | 0.897749 | 1.376414 | 12.83897 | 0.005335 | 0.048472 |

| | | | | | |
|---|---|---|---|---|---|
| ZNF75A | -0.62451 | 5.189277 | 12.9107 | 0.005337 | 0.048474 |
| RAPGEF1 | 0.705607 | 6.992732 | 12.90867 | 0.00534 | 0.048475 |
| PEG10 | 0.759628 | 11.3961 | 12.90559 | 0.005343 | 0.048487 |
| VGLL3 | -1.08614 | 3.310995 | 12.89425 | 0.005357 | 0.048576 |
| BMP5 | 1.134946 | 3.830596 | 12.89285 | 0.005358 | 0.048576 |
| NOMO2 | 0.716441 | 4.440898 | 12.89143 | 0.00536 | 0.048576 |
| ADCY5 | 0.818776 | 6.147863 | 12.88066 | 0.005373 | 0.048672 |
| AC125257.1 | -0.60373 | 5.476251 | 12.8743 | 0.005381 | 0.04872 |
| PARVB | 0.947592 | 2.712146 | 12.87045 | 0.005386 | 0.048741 |
| CA5BP1 | 0.825898 | 3.254719 | 12.8594 | 0.005399 | 0.048814 |
| SPOCK2 | 0.628949 | 7.953625 | 12.85729 | 0.005401 | 0.048814 |
| GIT1 | 0.885404 | 7.678764 | 12.85525 | 0.005404 | 0.048814 |
| MYOM3 | 1.220196 | -0.03783 | 12.84701 | 0.005405 | 0.048814 |
| AC074019.1 | -1.59405 | -0.62646 | 12.8536 | 0.005406 | 0.048814 |
| TP53 | -0.83033 | 3.563817 | 12.85176 | 0.005408 | 0.048814 |
| CHORDC1 | -0.76778 | 6.510094 | 12.8486 | 0.005412 | 0.048814 |
| NUTM2B | 1.053673 | 1.721944 | 12.84811 | 0.005413 | 0.048814 |
| AC139100.2 | -1.05057 | 0.862806 | 12.84248 | 0.005419 | 0.048854 |
| MVK | 0.664131 | 4.374607 | 12.83889 | 0.005424 | 0.048872 |
| AL355472.3 | 1.435059 | -0.21428 | 12.83607 | 0.005427 | 0.048881 |
| PCP4L1 | 1.373194 | 2.370712 | 12.83214 | 0.005432 | 0.048903 |
| DHCR7 | 0.659658 | 6.077161 | 12.82665 | 0.005439 | 0.048942 |
| ADM | 1.470394 | 1.684132 | 12.80238 | 0.005469 | 0.049189 |
| TRIM74 | -1.45248 | -0.8546 | 12.75155 | 0.005485 | 0.049314 |
| ELF5 | -1.83178 | -0.54349 | 12.78712 | 0.005487 | 0.049315 |
| DACH2 | 0.888241 | 5.99634 | 12.78459 | 0.005491 | 0.049321 |
| SMG1P6 | 1.334353 | -0.29032 | 12.77699 | 0.0055 | 0.049384 |
| ADAR | 0.478739 | 7.967479 | 12.69857 | 0.005507 | 0.049428 |
| NFIA-AS2 | -1.7184 | 2.501827 | 12.76882 | 0.00551 | 0.049432 |
| BRI3BP | 0.571618 | 4.487367 | 12.69262 | 0.005515 | 0.049452 |
| RGS8 | 1.218943 | 4.331117 | 12.76043 | 0.005521 | 0.049482 |
| ACOT9 | 0.550932 | 4.679337 | 12.68108 | 0.005529 | 0.049517 |
| ARNT2 | 0.707874 | 7.351765 | 12.75302 | 0.00553 | 0.049517 |
| ST7-AS2 | -2.90803 | 0.405245 | 12.74877 | 0.005535 | 0.049517 |
| LINC02166 | -1.42902 | -0.42422 | 12.74857 | 0.005535 | 0.049517 |
| C22orf39 | -0.56022 | 5.312272 | 12.70664 | 0.005537 | 0.049517 |
| ZBTB33 | 0.621402 | 6.942054 | 12.73901 | 0.005547 | 0.049592 |
| NFAM1 | 1.208992 | 0.653854 | 12.73142 | 0.005557 | 0.049656 |
| SLC9A3R1 | -0.89973 | 5.15615 | 12.7295 | 0.005559 | 0.049656 |
| RPS6KA1 | 0.735963 | 4.213408 | 12.71666 | 0.005576 | 0.049775 |
| AC093909.6 | -0.6678 | 4.093128 | 12.71242 | 0.005581 | 0.049775 |
| PAF1 | -0.53081 | 6.060299 | 12.66965 | 0.005582 | 0.049775 |
| ZNF488 | 1.262918 | -0.11445 | 12.71123 | 0.005582 | 0.049775 |
| AMIGO1 | 0.57486 | 5.455033 | 12.69375 | 0.005605 | 0.049929 |
| SIMC1 | 0.490832 | 5.37919 | 12.62019 | 0.005607 | 0.049929 |
| PPM1L | 0.612842 | 6.43797 | 12.69183 | 0.005607 | 0.049929 |
| DEF6 | -0.89136 | 1.379021 | 12.61408 | 0.005615 | 0.049975 |

**Table. S3B: PD-associated genes.**

| Gene symbol | logFC | logCPM | F | PValue | FDR |
|---|---|---|---|---|---|
| CHCHD2 | -1.36634 | 6.379506 | 88.80828 | 3.73E-06 | 0.000715 |
| NR4A2 | -1.93683 | 2.271371 | 28.10999 | 0.000414 | 0.010417 |
| PODXL | 0.653688 | 4.542829 | 15.47349 | 0.003044 | 0.03513 |
| SNCA | 0.985855 | 8.252316 | 23.56933 | 0.000776 | 0.015351 |

https://pathcards.genecards.org/Card/parkinsons_disease_pathway?queryString=parkinson%20disease

**Table. S3C: SNCA-associated genes.**

| Gene symbol | logFC | logCPM | F | PValue | FDR |
|---|---|---|---|---|---|
| SNCA | 0.985855 | 8.252316 | 23.56933 | 0.000776 | 0.015351 |
| MAOB | 1.33368 | 7.891897 | 22.12699 | 0.000966 | 0.017704 |
| PPP2R5D | 0.955379 | 5.392659 | 20.58782 | 0.001235 | 0.020204 |
| KLK6 | 1.46879 | -1.22401 | 13.1977 | 0.004922 | 0.046316 |

https://pathcards.genecards.org/Card/alpha-synuclein_signaling?queryString=SNCA

**Table. S3D: Enrichment analysis report.**

| # | Processes | p-value | FDR | In Data |
|---|---|---|---|---|
| 1 | multicellular organism development | 6.138E-32 | 3.371E-28 | 681 |
| 2 | system development | 7.317E-32 | 3.371E-28 | 620 |
| 3 | anatomical structure development | 5.007E-30 | 1.538E-26 | 710 |
| 4 | developmental process | 2.744E-28 | 6.321E-25 | 747 |
| 5 | multicellular organismal process | 1.132E-22 | 2.086E-19 | 849 |
| 6 | nervous system development | 1.039E-21 | 1.596E-18 | 348 |
| 7 | animal organ development | 5.223E-20 | 6.832E-17 | 477 |
| 8 | regulation of localization | 5.932E-20 | 6.832E-17 | 386 |
| 9 | cell adhesion | 4.685E-18 | 4.536E-15 | 162 |
| 10 | biological adhesion | 4.923E-18 | 4.536E-15 | 164 |
| 11 | regulation of biological quality | 7.430E-17 | 6.223E-14 | 505 |
| 12 | cell-cell adhesion via plasma-membrane adhesion molecules | 1.001E-16 | 7.689E-14 | 68 |
| 13 | homophilic cell adhesion via plasma membrane adhesion molecules | 3.155E-16 | 2.236E-13 | 51 |
| 14 | regulation of signaling | 9.693E-16 | 6.303E-13 | 430 |
| 15 | regulation of cell communication | 1.026E-15 | 6.303E-13 | 427 |
| 16 | regulation of multicellular organismal process | 1.120E-15 | 6.452E-13 | 371 |
| 17 | negative regulation of biological process | 3.094E-14 | 1.600E-11 | 620 |
| 18 | cell-cell adhesion | 3.125E-14 | 1.600E-11 | 104 |
| 19 | negative regulation of cellular process | 1.283E-13 | 6.221E-11 | 571 |
| 20 | anatomical structure morphogenesis | 5.879E-13 | 2.708E-10 | 303 |
| 21 | regulation of transport | 1.365E-12 | 5.991E-10 | 261 |
| 22 | regulation of cellular process | 1.526E-12 | 6.393E-10 | 1109 |
| 23 | biological regulation | 2.398E-12 | 9.606E-10 | 1202 |
| 24 | regulation of biological process | 2.882E-12 | 1.106E-09 | 1152 |
| 25 | regulation of developmental process | 5.460E-12 | 2.012E-09 | 331 |
| 26 | tissue development | 6.048E-12 | 2.143E-09 | 264 |
| 27 | neurogenesis | 7.080E-12 | 2.416E-09 | 219 |
| 28 | regulation of ion transport | 7.982E-12 | 2.627E-09 | 208 |
| 29 | cell differentiation | 8.465E-12 | 2.690E-09 | 447 |
| 30 | cellular developmental process | 1.679E-11 | 5.156E-09 | 450 |
| 31 | skeletal system development | 2.104E-11 | 6.254E-09 | 94 |
| 32 | chemical synaptic transmission | 2.674E-11 | 7.465E-09 | 88 |
| 33 | anterograde trans-synaptic signaling | 2.674E-11 | 7.465E-09 | 88 |
| 34 | positive regulation of cellular process | 2.793E-11 | 7.569E-09 | 621 |
| 35 | cell-cell signaling | 2.919E-11 | 7.683E-09 | 173 |
| 36 | trans-synaptic signaling | 4.425E-11 | 1.133E-08 | 90 |
| 37 | generation of neurons | 5.171E-11 | 1.288E-08 | 201 |
| 38 | localization | 5.645E-11 | 1.369E-08 | 598 |
| 39 | positive regulation of biological process | 8.670E-11 | 2.048E-08 | 668 |
| 40 | cellular response to growth factor stimulus | 1.571E-10 | 3.620E-08 | 102 |
| 41 | cellular process | 2.041E-10 | 4.586E-08 | 1387 |
| 42 | growth | 2.114E-10 | 4.638E-08 | 93 |
| 43 | response to growth factor | 3.078E-10 | 6.595E-08 | 106 |

| | | | | |
|---|---|---|---|---|
| 44 | central nervous system development | 3.449E-10 | 7.222E-08 | 175 |
| 45 | synaptic signaling | 4.923E-10 | 1.008E-07 | 93 |
| 46 | regulation of secretion | 5.160E-10 | 1.034E-07 | 126 |
| 47 | animal organ morphogenesis | 5.294E-10 | 1.038E-07 | 159 |
| 48 | cell development | 6.009E-10 | 1.153E-07 | 242 |
| 49 | brain development | 6.887E-10 | 1.295E-07 | 143 |
| 50 | regulation of system process | 7.646E-10 | 1.409E-07 | 115 |
| 51 | response to endogenous stimulus | 7.912E-10 | 1.429E-07 | 240 |
| 52 | regulation of secretion by cell | 8.181E-10 | 1.450E-07 | 116 |
| 53 | developmental growth | 1.147E-09 | 1.995E-07 | 88 |
| 54 | regulation of cellular response to growth factor stimulus | 1.607E-09 | 2.742E-07 | 60 |
| 55 | head development | 1.867E-09 | 3.127E-07 | 147 |
| 56 | pattern specification process | 3.920E-09 | 6.450E-07 | 80 |
| 57 | regulation of signal transduction | 5.334E-09 | 8.623E-07 | 354 |
| 58 | regulation of hormone levels | 6.006E-09 | 9.542E-07 | 102 |
| 59 | positive regulation of developmental process | 6.867E-09 | 1.072E-06 | 194 |
| 60 | neuron differentiation | 8.755E-09 | 1.345E-06 | 163 |
| 61 | regulation of cell population proliferation | 9.611E-09 | 1.452E-06 | 236 |
| 62 | negative regulation of multicellular organismal process | 1.150E-08 | 1.709E-06 | 156 |
| 63 | extracellular matrix organization | 1.280E-08 | 1.872E-06 | 67 |
| 64 | regulation of cell differentiation | 1.300E-08 | 1.872E-06 | 223 |
| 65 | extracellular structure organization | 1.384E-08 | 1.962E-06 | 67 |
| 66 | response to organic substance | 1.859E-08 | 2.595E-06 | 399 |
| 67 | circulatory system process | 2.291E-08 | 3.150E-06 | 93 |
| 68 | regulation of membrane potential | 2.449E-08 | 3.318E-06 | 83 |
| 69 | cellular response to endogenous stimulus | 2.919E-08 | 3.897E-06 | 192 |
| 70 | ion transmembrane transport | 4.611E-08 | 6.070E-06 | 140 |
| 71 | regulation of cellular component movement | 5.067E-08 | 6.576E-06 | 149 |
| 72 | embryo development | 5.498E-08 | 7.035E-06 | 171 |
| 73 | embryonic morphogenesis | 5.869E-08 | 7.408E-06 | 102 |
| 74 | regulation of multicellular organismal development | 7.903E-08 | 9.840E-06 | 194 |
| 75 | negative regulation of developmental process | 8.173E-08 | 1.004E-05 | 138 |
| 76 | anatomical structure formation involved in morphogenesis | 8.894E-08 | 1.064E-05 | 138 |
| 77 | cellular component morphogenesis | 8.971E-08 | 1.064E-05 | 101 |
| 78 | regulation of hormone secretion | 9.007E-08 | 1.064E-05 | 64 |
| 79 | establishment of localization | 1.032E-07 | 1.204E-05 | 474 |
| 80 | transmembrane transport | 1.101E-07 | 1.268E-05 | 164 |
| 81 | regulation of transmembrane receptor protein serine/threonine kinase | 1.232E-07 | 1.399E-05 | 49 |
| 82 | cellular component organization | 1.245E-07 | 1.399E-05 | 570 |
| 83 | regulation of anion transport | 1.618E-07 | 1.796E-05 | 140 |
| 84 | positive regulation of ion transport | 1.707E-07 | 1.872E-05 | 117 |
| 85 | positive regulation of multicellular organismal process | 1.976E-07 | 2.142E-05 | 200 |
| 86 | skeletal system morphogenesis | 2.175E-07 | 2.330E-05 | 48 |
| 87 | cartilage development | 2.376E-07 | 2.516E-05 | 37 |
| 88 | regulation of response to stimulus | 2.449E-07 | 2.564E-05 | 454 |
| 89 | regionalization | 2.521E-07 | 2.610E-05 | 61 |
| 90 | embryonic organ development | 2.602E-07 | 2.664E-05 | 84 |

| | | | | |
|---|---|---|---|---|
| 91 | negative regulation of response to stimulus | 3.002E-07 | 3.040E-05 | 209 |
| 92 | embryonic skeletal system development | 3.048E-07 | 3.052E-05 | 33 |
| 93 | positive regulation of cell communication | 3.297E-07 | 3.267E-05 | 220 |
| 94 | locomotion | 3.772E-07 | 3.697E-05 | 179 |
| 95 | positive regulation of signaling | 3.826E-07 | 3.711E-05 | 220 |
| 96 | negative regulation of signaling | 4.100E-07 | 3.935E-05 | 183 |
| 97 | regulation of locomotion | 4.212E-07 | 4.001E-05 | 141 |
| 98 | tube development | 5.299E-07 | 4.982E-05 | 140 |
| 99 | negative regulation of cell communication | 5.764E-07 | 5.365E-05 | 182 |
| 100 | embryonic skeletal system morphogenesis | 6.591E-07 | 6.073E-05 | 27 |
| 101 | regulation of anatomical structure morphogenesis | 6.664E-07 | 6.079E-05 | 140 |
| 102 | transport | 6.846E-07 | 6.184E-05 | 456 |
| 103 | regulation of molecular function | 7.584E-07 | 6.784E-05 | 343 |
| 104 | inorganic ion transmembrane transport | 7.895E-07 | 6.994E-05 | 95 |
| 105 | axon guidance | 9.521E-07 | 8.340E-05 | 51 |
| 106 | regulation of neurotransmitter levels | 9.594E-07 | 8.340E-05 | 54 |
| 107 | neuropeptide signaling pathway | 9.903E-07 | 8.527E-05 | 27 |
| 108 | neuron projection guidance | 1.027E-06 | 8.766E-05 | 51 |
| 109 | positive regulation of cellular metabolic process | 1.038E-06 | 8.775E-05 | 376 |
| 110 | connective tissue development | 1.070E-06 | 8.962E-05 | 45 |
| 111 | movement of cell or subcellular component | 1.251E-06 | 1.025E-04 | 199 |
| 112 | positive regulation of cell differentiation | 1.266E-06 | 1.025E-04 | 133 |
| 113 | cation transport | 1.268E-06 | 1.025E-04 | 114 |
| 114 | regulation of peptide secretion | 1.268E-06 | 1.025E-04 | 66 |
| 115 | cellular response to transforming growth factor beta stimulus | 1.303E-06 | 1.044E-04 | 40 |
| 116 | axonogenesis | 1.381E-06 | 1.097E-04 | 67 |
| 117 | response to oxygen-containing compound | 1.433E-06 | 1.129E-04 | 252 |
| 118 | regulation of cell motility | 1.472E-06 | 1.150E-04 | 133 |
| 119 | regulation of phosphate metabolic process | 1.548E-06 | 1.198E-04 | 224 |
| 120 | regulation of phosphorus metabolic process | 1.591E-06 | 1.217E-04 | 224 |
| 121 | muscle contraction | 1.599E-06 | 1.217E-04 | 49 |
| 122 | secretion | 1.823E-06 | 1.377E-04 | 138 |
| 123 | gland development | 1.957E-06 | 1.461E-04 | 89 |
| 124 | response to transforming growth factor beta | 1.966E-06 | 1.461E-04 | 41 |
| 125 | developmental growth involved in morphogenesis | 2.011E-06 | 1.482E-04 | 32 |
| 126 | modulation of chemical synaptic transmission | 2.092E-06 | 1.530E-04 | 83 |
| 127 | regulation of cell migration | 2.137E-06 | 1.550E-04 | 126 |
| 128 | regulation of transcription by RNA polymerase II | 2.166E-06 | 1.559E-04 | 283 |
| 129 | export from cell | 2.311E-06 | 1.630E-04 | 129 |
| 130 | regulation of trans-synaptic signaling | 2.314E-06 | 1.630E-04 | 83 |
| 131 | cellular response to organic substance | 2.318E-06 | 1.630E-04 | 303 |
| 132 | urogenital system development | 2.414E-06 | 1.685E-04 | 65 |
| 133 | response to organic cyclic compound | 2.921E-06 | 2.024E-04 | 173 |
| 134 | muscle structure development | 3.045E-06 | 2.094E-04 | 85 |
| 135 | embryonic organ morphogenesis | 3.156E-06 | 2.154E-04 | 57 |
| 136 | tube morphogenesis | 3.405E-06 | 2.293E-04 | 106 |
| 137 | multicellular organismal signaling | 3.409E-06 | 2.293E-04 | 32 |

| | | | | |
|---|---|---|---|---|
| 138 | blood circulation | 3.443E-06 | 2.299E-04 | 77 |
| 139 | neuron development | 3.610E-06 | 2.393E-04 | 128 |
| 140 | response to lipid | 3.712E-06 | 2.443E-04 | 161 |
| 141 | chemical homeostasis | 3.813E-06 | 2.491E-04 | 159 |
| 142 | cellular component organization or biogenesis | 3.908E-06 | 2.536E-04 | 573 |
| 143 | regulation of cellular metabolic process | 3.937E-06 | 2.537E-04 | 609 |
| 144 | cell projection organization | 4.741E-06 | 3.034E-04 | 157 |
| 145 | renal system development | 4.775E-06 | 3.034E-04 | 58 |
| 146 | kidney epithelium development | 5.104E-06 | 3.220E-04 | 31 |
| 147 | regulation of ion transmembrane transport | 5.136E-06 | 3.220E-04 | 90 |
| 148 | sensory organ development | 5.275E-06 | 3.284E-04 | 101 |
| 149 | positive regulation of transport | 5.730E-06 | 3.544E-04 | 140 |
| 150 | adenylate cyclase-modulating G protein-coupled receptor signaling path | 5.938E-06 | 3.626E-04 | 42 |
| 151 | regulation of endocrine process | 5.942E-06 | 3.626E-04 | 17 |
| 152 | cell junction organization | 6.214E-06 | 3.767E-04 | 85 |
| 153 | metal ion transport | 6.286E-06 | 3.786E-04 | 89 |
| 154 | cell projection morphogenesis | 6.416E-06 | 3.839E-04 | 80 |
| 155 | regulation of heart contraction | 6.504E-06 | 3.866E-04 | 47 |
| 156 | muscle organ development | 6.545E-06 | 3.866E-04 | 56 |
| 157 | circulatory system development | 6.749E-06 | 3.961E-04 | 132 |
| 158 | neuron projection development | 6.807E-06 | 3.969E-04 | 106 |
| 159 | cell part morphogenesis | 7.406E-06 | 4.292E-04 | 83 |
| 160 | positive regulation of nitrogen compound metabolic process | 7.682E-06 | 4.424E-04 | 350 |
| 161 | secretion by cell | 7.884E-06 | 4.487E-04 | 121 |
| 162 | plasma membrane bounded cell projection morphogenesis | 7.889E-06 | 4.487E-04 | 79 |
| 163 | cell morphogenesis involved in neuron differentiation | 8.351E-06 | 4.710E-04 | 74 |
| 164 | epithelial tube morphogenesis | 8.430E-06 | 4.710E-04 | 59 |
| 165 | positive regulation of metabolic process | 8.458E-06 | 4.710E-04 | 411 |
| 166 | import into cell | 8.485E-06 | 4.710E-04 | 35 |
| 167 | reproductive structure development | 9.084E-06 | 5.012E-04 | 82 |
| 168 | regulation of transmembrane transport | 9.561E-06 | 5.244E-04 | 90 |
| 169 | cellular chemical homeostasis | 1.028E-05 | 5.603E-04 | 115 |
| 170 | regulation of nervous system development | 1.067E-05 | 5.782E-04 | 78 |
| 171 | signal release | 1.079E-05 | 5.816E-04 | 40 |
| 172 | kidney morphogenesis | 1.104E-05 | 5.896E-04 | 23 |
| 173 | regulation of cellular component organization | 1.107E-05 | 5.896E-04 | 275 |
| 174 | neuron projection morphogenesis | 1.119E-05 | 5.924E-04 | 78 |
| 175 | import across plasma membrane | 1.133E-05 | 5.964E-04 | 28 |
| 176 | sex differentiation | 1.143E-05 | 5.986E-04 | 60 |
| 177 | negative regulation of cell differentiation | 1.192E-05 | 6.206E-04 | 101 |
| 178 | cellular response to chemical stimulus | 1.248E-05 | 6.441E-04 | 354 |
| 179 | reproductive system development | 1.253E-05 | 6.441E-04 | 82 |
| 180 | forebrain development | 1.268E-05 | 6.441E-04 | 72 |
| 181 | morphogenesis of an epithelium | 1.268E-05 | 6.441E-04 | 76 |
| 182 | cellular response to BMP stimulus | 1.279E-05 | 6.441E-04 | 25 |
| 183 | response to BMP | 1.279E-05 | 6.441E-04 | 25 |
| 184 | synapse organization | 1.293E-05 | 6.473E-04 | 61 |

| | | | | |
|---|---|---|---|---|
| 185 | tissue morphogenesis | 1.349E-05 | 6.717E-04 | 90 |
| 186 | ion homeostasis | 1.383E-05 | 6.853E-04 | 111 |
| 187 | epithelium development | 1.458E-05 | 7.184E-04 | 158 |
| 188 | calcium ion homeostasis | 1.509E-05 | 7.396E-04 | 77 |
| 189 | ion transport | 1.521E-05 | 7.415E-04 | 300 |
| 190 | positive regulation of phosphate metabolic process | 1.582E-05 | 7.633E-04 | 153 |
| 191 | positive regulation of phosphorus metabolic process | 1.582E-05 | 7.633E-04 | 153 |
| 192 | post-embryonic body morphogenesis | 1.594E-05 | 7.650E-04 | 4 |
| 193 | neurotransmitter transport | 1.609E-05 | 7.680E-04 | 32 |
| 194 | renal tubule development | 1.664E-05 | 7.901E-04 | 22 |
| 195 | cell morphogenesis | 1.784E-05 | 8.428E-04 | 104 |
| 196 | regulation of neuron differentiation | 1.822E-05 | 8.521E-04 | 43 |
| 197 | cation transmembrane transport | 1.824E-05 | 8.521E-04 | 85 |
| 198 | negative regulation of transport | 1.831E-05 | 8.521E-04 | 79 |
| 199 | positive regulation of secretion by cell | 1.864E-05 | 8.633E-04 | 58 |
| 200 | negative regulation of transmembrane receptor protein serine/threonir | 1.886E-05 | 8.690E-04 | 27 |
| 201 | divalent inorganic cation homeostasis | 1.920E-05 | 8.800E-04 | 81 |
| 202 | plasma membrane bounded cell projection organization | 1.959E-05 | 8.937E-04 | 149 |
| 203 | male sex differentiation | 1.986E-05 | 8.994E-04 | 40 |
| 204 | regulation of neurogenesis | 1.999E-05 | 8.994E-04 | 65 |
| 205 | skeletal muscle organ development | 2.001E-05 | 8.994E-04 | 35 |
| 206 | inorganic anion transmembrane transport | 2.065E-05 | 9.160E-04 | 26 |
| 207 | regulation of protein secretion | 2.067E-05 | 9.160E-04 | 57 |
| 208 | G protein-coupled receptor signaling pathway, coupled to cyclic nucleot | 2.068E-05 | 9.160E-04 | 44 |
| 209 | signaling | 2.087E-05 | 9.200E-04 | 575 |
| 210 | axon extension | 2.140E-05 | 9.392E-04 | 15 |
| 211 | nephron epithelium development | 2.161E-05 | 9.436E-04 | 24 |
| 212 | regulation of primary metabolic process | 2.171E-05 | 9.437E-04 | 584 |
| 213 | mammary gland development | 2.207E-05 | 9.547E-04 | 33 |
| 214 | neuron projection extension | 2.235E-05 | 9.622E-04 | 20 |
| 215 | mesonephros development | 2.299E-05 | 9.854E-04 | 23 |
| 216 | inorganic ion homeostasis | 2.322E-05 | 9.906E-04 | 106 |
| 217 | muscle tissue development | 2.353E-05 | 9.992E-04 | 60 |
| 218 | positive regulation of transcription by RNA polymerase II | 2.495E-05 | 1.054E-03 | 162 |
| 219 | genetic imprinting | 2.620E-05 | 1.102E-03 | 11 |
| 220 | negative regulation of anion transport | 2.783E-05 | 1.165E-03 | 48 |
| 221 | regulation of peptide hormone secretion | 2.963E-05 | 1.235E-03 | 48 |
| 222 | neuron migration | 3.097E-05 | 1.285E-03 | 28 |
| 223 | female sex differentiation | 3.255E-05 | 1.345E-03 | 32 |
| 224 | adenylate cyclase-inhibiting G protein-coupled receptor signaling pathw | 3.325E-05 | 1.368E-03 | 20 |
| 225 | cation homeostasis | 3.478E-05 | 1.424E-03 | 104 |
| 226 | regulation of cytosolic calcium ion concentration | 3.606E-05 | 1.470E-03 | 59 |
| 227 | chloride transmembrane transport | 3.646E-05 | 1.470E-03 | 23 |
| 228 | embryonic cranial skeleton morphogenesis | 3.648E-05 | 1.470E-03 | 15 |
| 229 | nephron development | 3.653E-05 | 1.470E-03 | 29 |
| 230 | negative regulation of cell population proliferation | 3.707E-05 | 1.485E-03 | 107 |
| 231 | axon development | 3.765E-05 | 1.502E-03 | 70 |

| | | | | |
|---|---|---|---|---|
| 232 | nephron tubule development | 3.782E-05 | 1.502E-03 | 20 |
| 233 | muscle system process | 3.948E-05 | 1.561E-03 | 54 |
| 234 | regulation of nitrogen compound metabolic process | 4.013E-05 | 1.577E-03 | 564 |
| 235 | negative regulation of signal transduction | 4.022E-05 | 1.577E-03 | 159 |
| 236 | regulation of blood circulation | 4.138E-05 | 1.616E-03 | 53 |
| 237 | semaphorin-plexin signaling pathway | 4.316E-05 | 1.678E-03 | 11 |
| 238 | hindlimb morphogenesis | 5.335E-05 | 2.065E-03 | 14 |
| 239 | vascular transport | 5.804E-05 | 2.235E-03 | 18 |
| 240 | positive regulation of signal transduction | 5.858E-05 | 2.235E-03 | 188 |
| 241 | response to abiotic stimulus | 5.875E-05 | 2.235E-03 | 175 |
| 242 | metanephros development | 5.883E-05 | 2.235E-03 | 21 |
| 243 | positive regulation of secretion | 5.894E-05 | 2.235E-03 | 62 |
| 244 | negative regulation of secretion | 6.034E-05 | 2.279E-03 | 41 |
| 245 | cell morphogenesis involved in differentiation | 6.256E-05 | 2.353E-03 | 84 |
| 246 | skeletal muscle tissue development | 6.310E-05 | 2.363E-03 | 32 |
| 247 | negative regulation of response to wounding | 6.605E-05 | 2.454E-03 | 21 |
| 248 | pancreas development | 6.605E-05 | 2.454E-03 | 21 |
| 249 | regulation of protein phosphorylation | 6.716E-05 | 2.485E-03 | 178 |
| 250 | positive regulation of cell population proliferation | 7.120E-05 | 2.619E-03 | 137 |
| 251 | catechol-containing compound biosynthetic process | 7.163E-05 | 2.619E-03 | 10 |
| 252 | catecholamine biosynthetic process | 7.163E-05 | 2.619E-03 | 10 |
| 253 | semaphorin-plexin signaling pathway involved in neuron projection guic | 7.388E-05 | 2.691E-03 | 6 |
| 254 | striated muscle tissue development | 7.554E-05 | 2.740E-03 | 56 |
| 255 | regulation of excretion | 7.588E-05 | 2.742E-03 | 12 |
| 256 | response to hypoxia | 7.740E-05 | 2.786E-03 | 66 |
| 257 | regulation of phosphorylation | 7.805E-05 | 2.791E-03 | 194 |
| 258 | actin filament-based process | 7.814E-05 | 2.791E-03 | 82 |
| 259 | ear development | 7.947E-05 | 2.827E-03 | 46 |
| 260 | cell communication | 9.170E-05 | 3.250E-03 | 579 |
| 261 | kidney development | 9.388E-05 | 3.314E-03 | 52 |
| 262 | chloride transport | 9.553E-05 | 3.360E-03 | 23 |
| 263 | cellular homeostasis | 9.689E-05 | 3.394E-03 | 125 |
| 264 | response to increased oxygen levels | 9.990E-05 | 3.487E-03 | 16 |
| 265 | ureteric bud development | 1.034E-04 | 3.596E-03 | 21 |
| 266 | response to temperature stimulus | 1.078E-04 | 3.735E-03 | 41 |
| 267 | metal ion homeostasis | 1.084E-04 | 3.742E-03 | 93 |
| 268 | negative regulation of growth | 1.092E-04 | 3.756E-03 | 44 |
| 269 | regulation of catalytic activity | 1.112E-04 | 3.794E-03 | 261 |
| 270 | renal tubule morphogenesis | 1.112E-04 | 3.794E-03 | 18 |
| 271 | nephron tubule morphogenesis | 1.144E-04 | 3.890E-03 | 17 |
| 272 | chemotaxis | 1.161E-04 | 3.933E-03 | 77 |
| 273 | anion transmembrane transport | 1.170E-04 | 3.949E-03 | 61 |
| 274 | positive regulation of phosphorylation | 1.189E-04 | 3.999E-03 | 140 |
| 275 | negative regulation of ion transport | 1.246E-04 | 4.176E-03 | 60 |
| 276 | mesonephric tubule development | 1.283E-04 | 4.269E-03 | 21 |
| 277 | mesonephric epithelium development | 1.283E-04 | 4.269E-03 | 21 |
| 278 | cell motility | 1.302E-04 | 4.300E-03 | 141 |

| | | | | |
|---|---|---|---|---|
| 279 | localization of cell | 1.302E-04 | 4.300E-03 | 141 |
| 280 | heart development | 1.308E-04 | 4.305E-03 | 86 |
| 281 | phenol-containing compound biosynthetic process | 1.353E-04 | 4.435E-03 | 12 |
| 282 | reproduction | 1.389E-04 | 4.538E-03 | 207 |
| 283 | negative regulation of wound healing | 1.422E-04 | 4.631E-03 | 18 |
| 284 | nephron epithelium morphogenesis | 1.483E-04 | 4.810E-03 | 17 |
| 285 | response to corticosteroid | 1.582E-04 | 5.115E-03 | 52 |
| 286 | taxis | 1.593E-04 | 5.133E-03 | 77 |
| 287 | positive regulation of morphogenesis of an epithelium | 1.623E-04 | 5.212E-03 | 12 |
| 288 | positive regulation of macromolecule metabolic process | 1.657E-04 | 5.295E-03 | 373 |
| 289 | reproductive process | 1.661E-04 | 5.295E-03 | 206 |
| 290 | mesenchyme development | 1.677E-04 | 5.327E-03 | 39 |
| 291 | transport across blood-brain barrier | 1.683E-04 | 5.328E-03 | 17 |
| 292 | supramolecular fiber organization | 1.730E-04 | 5.460E-03 | 69 |
| 293 | plasma membrane organization | 1.757E-04 | 5.525E-03 | 21 |
| 294 | neurotransmitter secretion | 1.841E-04 | 5.735E-03 | 24 |
| 295 | signal release from synapse | 1.841E-04 | 5.735E-03 | 24 |
| 296 | regulation of growth | 1.843E-04 | 5.735E-03 | 97 |
| 297 | regulation of postsynaptic membrane potential | 1.849E-04 | 5.735E-03 | 26 |
| 298 | positive regulation of nucleic acid-templated transcription | 1.919E-04 | 5.933E-03 | 192 |
| 299 | negative regulation of renal sodium excretion | 1.930E-04 | 5.948E-03 | 6 |
| 300 | catecholamine metabolic process | 1.956E-04 | 5.966E-03 | 14 |
| 301 | catechol-containing compound metabolic process | 1.956E-04 | 5.966E-03 | 14 |
| 302 | response to hyperoxia | 1.956E-04 | 5.966E-03 | 14 |
| 303 | positive regulation of RNA biosynthetic process | 1.962E-04 | 5.966E-03 | 192 |
| 304 | negative regulation of developmental growth | 2.015E-04 | 6.108E-03 | 24 |
| 305 | male gonad development | 2.047E-04 | 6.185E-03 | 34 |
| 306 | regulation of neural precursor cell proliferation | 2.080E-04 | 6.264E-03 | 23 |
| 307 | regulation of transcription, DNA-templated | 2.148E-04 | 6.408E-03 | 342 |
| 308 | positive regulation of cold-induced thermogenesis | 2.149E-04 | 6.408E-03 | 20 |
| 309 | embryonic pattern specification | 2.154E-04 | 6.408E-03 | 17 |
| 310 | regulation of parathyroid hormone secretion | 2.156E-04 | 6.408E-03 | 4 |
| 311 | developmental process involved in reproduction | 2.209E-04 | 6.545E-03 | 145 |
| 312 | regulation of fat cell differentiation | 2.243E-04 | 6.624E-03 | 27 |
| 313 | regulation of neurotransmitter transport | 2.288E-04 | 6.736E-03 | 31 |
| 314 | regulation of nucleic acid-templated transcription | 2.322E-04 | 6.811E-03 | 346 |
| 315 | cellular calcium ion homeostasis | 2.331E-04 | 6.811E-03 | 70 |
| 316 | development of primary male sexual characteristics | 2.343E-04 | 6.811E-03 | 34 |
| 317 | positive regulation of peptide secretion | 2.343E-04 | 6.811E-03 | 34 |
| 318 | aminoglycan biosynthetic process | 2.385E-04 | 6.909E-03 | 20 |
| 319 | regulation of biosynthetic process | 2.459E-04 | 7.103E-03 | 414 |
| 320 | regulation of metabolic process | 2.481E-04 | 7.145E-03 | 658 |
| 321 | positive regulation of branching involved in ureteric bud morphogenesis | 2.494E-04 | 7.159E-03 | 8 |
| 322 | response to decreased oxygen levels | 2.518E-04 | 7.181E-03 | 66 |
| 323 | noradrenergic neuron development | 2.525E-04 | 7.181E-03 | 3 |
| 324 | intrinsic apoptotic signaling pathway in response to hypoxia | 2.525E-04 | 7.181E-03 | 3 |
| 325 | ossification | 2.581E-04 | 7.252E-03 | 46 |

| | | | | |
|---|---|---|---|---|
| 326 | glycosaminoglycan metabolic process | 2.582E-04 | 7.252E-03 | 26 |
| 327 | regulation of amine transport | 2.582E-04 | 7.252E-03 | 26 |
| 328 | cell growth | 2.582E-04 | 7.252E-03 | 26 |
| 329 | regulation of RNA biosynthetic process | 2.602E-04 | 7.286E-03 | 346 |
| 330 | peptidyl-tyrosine dephosphorylation | 2.622E-04 | 7.320E-03 | 19 |
| 331 | cardiac conduction | 2.643E-04 | 7.356E-03 | 20 |
| 332 | development of primary sexual characteristics | 2.683E-04 | 7.446E-03 | 49 |
| 333 | blood vessel development | 2.717E-04 | 7.504E-03 | 77 |
| 334 | cellular biogenic amine biosynthetic process | 2.730E-04 | 7.504E-03 | 12 |
| 335 | calcium-dependent cell-cell adhesion via plasma membrane cell adhesic | 2.730E-04 | 7.504E-03 | 12 |
| 336 | nephron morphogenesis | 2.737E-04 | 7.504E-03 | 17 |
| 337 | regulation of animal organ morphogenesis | 2.792E-04 | 7.633E-03 | 32 |
| 338 | response to wounding | 2.884E-04 | 7.862E-03 | 85 |
| 339 | glycosaminoglycan biosynthetic process | 2.916E-04 | 7.914E-03 | 19 |
| 340 | positive regulation of macrophage cytokine production | 2.923E-04 | 7.914E-03 | 6 |
| 341 | regulation of macrophage cytokine production | 2.929E-04 | 7.914E-03 | 7 |
| 342 | regulation of body fluid levels | 3.027E-04 | 8.156E-03 | 75 |
| 343 | response to hormone | 3.042E-04 | 8.172E-03 | 142 |
| 344 | regulation of protein localization | 3.134E-04 | 8.395E-03 | 122 |
| 345 | amine biosynthetic process | 3.218E-04 | 8.571E-03 | 12 |
| 346 | energy homeostasis | 3.218E-04 | 8.571E-03 | 12 |
| 347 | inorganic cation transmembrane transport | 3.246E-04 | 8.619E-03 | 73 |
| 348 | feeding behavior | 3.272E-04 | 8.664E-03 | 23 |
| 349 | response to glucocorticoid | 3.334E-04 | 8.784E-03 | 47 |
| 350 | sympathetic ganglion development | 3.365E-04 | 8.784E-03 | 5 |
| 351 | neuron projection extension involved in neuron projection guidance | 3.365E-04 | 8.784E-03 | 5 |
| 352 | positive regulation of corticosteroid hormone secretion | 3.365E-04 | 8.784E-03 | 5 |
| 353 | axon extension involved in axon guidance | 3.365E-04 | 8.784E-03 | 5 |
| 354 | negative regulation of hormone secretion | 3.400E-04 | 8.851E-03 | 22 |
| 355 | hormone transport | 3.502E-04 | 9.090E-03 | 21 |
| 356 | enzyme linked receptor protein signaling pathway | 3.522E-04 | 9.110E-03 | 94 |
| 357 | cell migration | 3.530E-04 | 9.110E-03 | 123 |
| 358 | negative regulation of cellular metabolic process | 3.683E-04 | 9.480E-03 | 286 |
| 359 | positive regulation of nervous system process | 3.971E-04 | 1.019E-02 | 14 |
| 360 | embryonic hindlimb morphogenesis | 4.046E-04 | 1.036E-02 | 11 |
| 361 | vascular process in circulatory system | 4.077E-04 | 1.041E-02 | 47 |
| 362 | regulation of branching involved in ureteric bud morphogenesis | 4.194E-04 | 1.067E-02 | 8 |
| 363 | cellular ion homeostasis | 4.355E-04 | 1.106E-02 | 92 |
| 364 | positive regulation of cellular component movement | 4.427E-04 | 1.121E-02 | 82 |
| 365 | positive regulation of locomotion | 4.451E-04 | 1.123E-02 | 81 |
| 366 | aminoglycan metabolic process | 4.506E-04 | 1.134E-02 | 27 |
| 367 | carboxylic acid transmembrane transport | 4.621E-04 | 1.160E-02 | 23 |
| 368 | cranial skeletal system development | 4.664E-04 | 1.168E-02 | 16 |
| 369 | cellular divalent inorganic cation homeostasis | 4.708E-04 | 1.176E-02 | 72 |
| 370 | synapse assembly | 4.743E-04 | 1.181E-02 | 24 |
| 371 | digestive system process | 4.824E-04 | 1.198E-02 | 17 |
| 372 | negative regulation of blood pressure | 4.959E-04 | 1.228E-02 | 15 |

| | | | | |
|---|---|---|---|---|
| 373 | regulation of cell development | 5.024E-04 | 1.235E-02 | 77 |
| 374 | developmental cell growth | 5.026E-04 | 1.235E-02 | 23 |
| 375 | regulation of heart rate | 5.026E-04 | 1.235E-02 | 23 |
| 376 | anterior/posterior pattern specification | 5.041E-04 | 1.235E-02 | 37 |
| 377 | positive regulation of transcription, DNA-templated | 5.054E-04 | 1.235E-02 | 183 |
| 378 | homeostatic process | 5.174E-04 | 1.259E-02 | 209 |
| 379 | chordate embryonic development | 5.189E-04 | 1.259E-02 | 104 |
| 380 | cell surface receptor signaling pathway | 5.192E-04 | 1.259E-02 | 266 |
| 381 | ectodermal placode morphogenesis | 5.342E-04 | 1.288E-02 | 8 |
| 382 | ectodermal placode formation | 5.342E-04 | 1.288E-02 | 8 |
| 383 | regulation of cellular biosynthetic process | 5.369E-04 | 1.292E-02 | 401 |
| 384 | myelination | 5.462E-04 | 1.309E-02 | 23 |
| 385 | semaphorin-plexin signaling pathway involved in axon guidance | 5.469E-04 | 1.309E-02 | 5 |
| 386 | regulation of response to wounding | 5.580E-04 | 1.326E-02 | 31 |
| 387 | digestive system development | 5.580E-04 | 1.326E-02 | 31 |
| 388 | regulation of reactive oxygen species metabolic process | 5.585E-04 | 1.326E-02 | 36 |
| 389 | regulation of intracellular signal transduction | 5.622E-04 | 1.332E-02 | 200 |
| 390 | vasculature development | 5.718E-04 | 1.351E-02 | 79 |
| 391 | gonad development | 5.743E-04 | 1.353E-02 | 47 |
| 392 | response to oxygen levels | 5.851E-04 | 1.373E-02 | 73 |
| 393 | positive regulation of protein secretion | 5.856E-04 | 1.373E-02 | 29 |
| 394 | positive regulation of cell motility | 5.916E-04 | 1.383E-02 | 79 |
| 395 | organic acid transmembrane transport | 5.930E-04 | 1.383E-02 | 23 |
| 396 | positive regulation of calcium ion import | 5.973E-04 | 1.390E-02 | 10 |
| 397 | heterophilic cell-cell adhesion via plasma membrane cell adhesion mole | 6.027E-04 | 1.397E-02 | 13 |
| 398 | inorganic anion transport | 6.036E-04 | 1.397E-02 | 27 |
| 399 | cell fate commitment | 6.056E-04 | 1.398E-02 | 39 |
| 400 | negative regulation of secretion by cell | 6.075E-04 | 1.399E-02 | 34 |
| 401 | cellular cation homeostasis | 6.105E-04 | 1.403E-02 | 90 |
| 402 | positive regulation of nervous system development | 6.276E-04 | 1.439E-02 | 50 |
| 403 | cellular response to organonitrogen compound | 6.347E-04 | 1.451E-02 | 104 |
| 404 | regulation of cell adhesion | 6.473E-04 | 1.476E-02 | 101 |
| 405 | response to organonitrogen compound | 6.523E-04 | 1.484E-02 | 161 |
| 406 | regulation of morphogenesis of a branching structure | 6.547E-04 | 1.484E-02 | 16 |
| 407 | potassium ion transmembrane transport | 6.557E-04 | 1.484E-02 | 25 |
| 408 | segmentation | 6.605E-04 | 1.492E-02 | 21 |
| 409 | negative regulation of transforming growth factor beta receptor signalir | 6.642E-04 | 1.496E-02 | 17 |
| 410 | appendage development | 6.712E-04 | 1.505E-02 | 33 |
| 411 | limb development | 6.712E-04 | 1.505E-02 | 33 |
| 412 | ectodermal placode development | 6.731E-04 | 1.505E-02 | 8 |
| 413 | regulation of cold-induced thermogenesis | 7.068E-04 | 1.577E-02 | 25 |
| 414 | regulation of insulin secretion | 7.112E-04 | 1.583E-02 | 38 |
| 415 | positive regulation of protein phosphorylation | 7.133E-04 | 1.584E-02 | 127 |
| 416 | cellular response to nitrogen compound | 7.205E-04 | 1.594E-02 | 111 |
| 417 | regulation of metal ion transport | 7.215E-04 | 1.594E-02 | 63 |
| 418 | positive regulation of positive chemotaxis | 7.238E-04 | 1.595E-02 | 9 |
| 419 | actin cytoskeleton organization | 7.300E-04 | 1.602E-02 | 69 |

| | | | | |
|---|---|---|---|---|
| 420 | negative regulation of ERK1 and ERK2 cascade | 7.304E-04 | 1.602E-02 | 16 |
| 421 | heart morphogenesis | 7.334E-04 | 1.605E-02 | 43 |
| 422 | regulation of axonogenesis | 7.445E-04 | 1.625E-02 | 30 |
| 423 | ensheathment of neurons | 7.551E-04 | 1.639E-02 | 23 |
| 424 | axon ensheathment | 7.551E-04 | 1.639E-02 | 23 |
| 425 | potassium ion transport | 7.562E-04 | 1.639E-02 | 26 |
| 426 | negative regulation of cellular response to growth factor stimulus | 7.830E-04 | 1.690E-02 | 19 |
| 427 | positive regulation of biosynthetic process | 7.833E-04 | 1.690E-02 | 225 |
| 428 | cell fate determination | 7.974E-04 | 1.717E-02 | 12 |
| 429 | embryo development ending in birth or egg hatching | 8.036E-04 | 1.726E-02 | 105 |
| 430 | regulation of cellular response to transforming growth factor beta stimu | 8.170E-04 | 1.751E-02 | 23 |
| 431 | positive regulation of nucleobase-containing compound metabolic proc | 8.309E-04 | 1.776E-02 | 212 |
| 432 | regulation of serotonin secretion | 8.404E-04 | 1.792E-02 | 6 |
| 433 | regulation of kidney development | 8.448E-04 | 1.798E-02 | 10 |
| 434 | positive regulation of cellular biosynthetic process | 8.579E-04 | 1.821E-02 | 221 |
| 435 | digestive tract development | 8.727E-04 | 1.847E-02 | 29 |
| 436 | regulation of renal sodium excretion | 8.759E-04 | 1.847E-02 | 9 |
| 437 | regulation of positive chemotaxis | 8.759E-04 | 1.847E-02 | 9 |
| 438 | positive regulation of fat cell differentiation | 8.890E-04 | 1.870E-02 | 15 |
| 439 | positive regulation of RNA metabolic process | 8.968E-04 | 1.882E-02 | 195 |
| 440 | anatomical structure arrangement | 9.039E-04 | 1.893E-02 | 7 |
| 441 | glutamine transport | 9.077E-04 | 1.896E-02 | 4 |
| 442 | positive regulation of aldosterone secretion | 9.622E-04 | 1.983E-02 | 3 |
| 443 | aminergic neurotransmitter loading into synaptic vesicle | 9.622E-04 | 1.983E-02 | 3 |
| 444 | positive regulation of mineralocorticoid secretion | 9.622E-04 | 1.983E-02 | 3 |
| 445 | GPI anchor release | 9.622E-04 | 1.983E-02 | 3 |
| 446 | serotonin biosynthetic process | 9.622E-04 | 1.983E-02 | 3 |
| 447 | positive regulation of serotonin secretion | 9.622E-04 | 1.983E-02 | 3 |
| 448 | regulation of BMP signaling pathway | 9.730E-04 | 2.001E-02 | 18 |
| 449 | mesenchymal cell differentiation | 9.825E-04 | 2.016E-02 | 27 |
| 450 | regulation of vascular endothelial growth factor receptor signaling path | 9.969E-04 | 2.041E-02 | 10 |
| 451 | positive regulation of cell migration | 1.006E-03 | 2.056E-02 | 75 |
| 452 | cellular biogenic amine metabolic process | 1.028E-03 | 2.095E-02 | 19 |
| 453 | positive regulation of intracellular signal transduction | 1.030E-03 | 2.095E-02 | 126 |
| 454 | regulation of steroid hormone secretion | 1.038E-03 | 2.106E-02 | 8 |
| 455 | regulation of neurotransmitter secretion | 1.074E-03 | 2.176E-02 | 26 |
| 456 | regulation of morphogenesis of an epithelium | 1.098E-03 | 2.218E-02 | 17 |
| 457 | cellular amine metabolic process | 1.123E-03 | 2.261E-02 | 19 |
| 458 | positive regulation of ion transmembrane transport | 1.126E-03 | 2.261E-02 | 40 |
| 459 | positive regulation of transmembrane transport | 1.126E-03 | 2.261E-02 | 40 |
| 460 | negative regulation of phosphate metabolic process | 1.153E-03 | 2.309E-02 | 78 |
| 461 | regulation of epithelial cell proliferation | 1.155E-03 | 2.309E-02 | 57 |
| 462 | regulation of MAPK cascade | 1.161E-03 | 2.315E-02 | 95 |
| 463 | digestion | 1.185E-03 | 2.353E-02 | 21 |
| 464 | sensory perception of pain | 1.185E-03 | 2.353E-02 | 21 |
| 465 | negative regulation of phosphorus metabolic process | 1.191E-03 | 2.359E-02 | 78 |
| 466 | ventricular cardiac muscle cell membrane repolarization | 1.243E-03 | 2.441E-02 | 5 |

| | | | | |
|---|---|---|---|---|
| 467 | hair follicle placode formation | 1.243E-03 | 2.441E-02 | 5 |
| 468 | membrane repolarization during ventricular cardiac muscle cell action p | 1.243E-03 | 2.441E-02 | 5 |
| 469 | cranial nerve structural organization | 1.243E-03 | 2.441E-02 | 5 |
| 470 | negative regulation of transcription by RNA polymerase II | 1.249E-03 | 2.449E-02 | 116 |
| 471 | wound healing | 1.261E-03 | 2.468E-02 | 64 |
| 472 | aortic valve morphogenesis | 1.272E-03 | 2.483E-02 | 8 |
| 473 | response to carbohydrate | 1.329E-03 | 2.589E-02 | 42 |
| 474 | behavior | 1.398E-03 | 2.717E-02 | 94 |
| 475 | development of primary female sexual characteristics | 1.407E-03 | 2.723E-02 | 26 |
| 476 | sulfur compound biosynthetic process | 1.407E-03 | 2.723E-02 | 26 |
| 477 | positive regulation of anion transport | 1.410E-03 | 2.724E-02 | 77 |
| 478 | response to monosaccharide | 1.416E-03 | 2.729E-02 | 37 |
| 479 | response to heat | 1.442E-03 | 2.774E-02 | 25 |
| 480 | regulation of protein modification process | 1.448E-03 | 2.775E-02 | 207 |
| 481 | inorganic ion import across plasma membrane | 1.458E-03 | 2.775E-02 | 17 |
| 482 | inorganic cation import across plasma membrane | 1.458E-03 | 2.775E-02 | 17 |
| 483 | response to drug | 1.458E-03 | 2.775E-02 | 84 |
| 484 | regulation of pathway-restricted SMAD protein phosphorylation | 1.461E-03 | 2.775E-02 | 13 |
| 485 | gamma-aminobutyric acid signaling pathway | 1.461E-03 | 2.775E-02 | 13 |
| 486 | cellular metal ion homeostasis | 1.493E-03 | 2.831E-02 | 80 |
| 487 | regulation of transforming growth factor beta receptor signaling pathwa | 1.502E-03 | 2.837E-02 | 22 |
| 488 | vesicle-mediated transport in synapse | 1.503E-03 | 2.837E-02 | 26 |
| 489 | type B pancreatic cell development | 1.507E-03 | 2.840E-02 | 6 |
| 490 | regulation of GTPase activity | 1.511E-03 | 2.841E-02 | 63 |
| 491 | positive regulation of cell-substrate adhesion | 1.543E-03 | 2.883E-02 | 25 |
| 492 | response to parathyroid hormone | 1.546E-03 | 2.883E-02 | 8 |
| 493 | neurotransmitter loading into synaptic vesicle | 1.552E-03 | 2.883E-02 | 4 |
| 494 | positive regulation of ER-associated ubiquitin-dependent protein catabo | 1.552E-03 | 2.883E-02 | 4 |
| 495 | positive regulation of alkaline phosphatase activity | 1.552E-03 | 2.883E-02 | 4 |
| 496 | commitment of neuronal cell to specific neuron type in forebrain | 1.552E-03 | 2.883E-02 | 4 |
| 497 | limb morphogenesis | 1.594E-03 | 2.950E-02 | 28 |
| 498 | appendage morphogenesis | 1.594E-03 | 2.950E-02 | 28 |
| 499 | stem cell differentiation | 1.627E-03 | 3.004E-02 | 29 |
| 500 | regulation of peptide transport | 1.639E-03 | 3.021E-02 | 86 |
| 501 | morphogenesis of a branching epithelium | 1.651E-03 | 3.036E-02 | 30 |
| 502 | mucopolysaccharide metabolic process | 1.717E-03 | 3.152E-02 | 19 |
| 503 | response to chemical | 1.720E-03 | 3.152E-02 | 496 |
| 504 | bone growth | 1.764E-03 | 3.218E-02 | 9 |
| 505 | semi-lunar valve development | 1.764E-03 | 3.218E-02 | 9 |
| 506 | negative regulation of voltage-gated potassium channel activity | 1.767E-03 | 3.218E-02 | 5 |
| 507 | sympathetic nervous system development | 1.816E-03 | 3.293E-02 | 7 |
| 508 | regulation of catecholamine secretion | 1.816E-03 | 3.293E-02 | 16 |
| 509 | negative regulation of neuron differentiation | 1.820E-03 | 3.295E-02 | 18 |
| 510 | ureteric bud morphogenesis | 1.840E-03 | 3.324E-02 | 13 |
| 511 | transmission of nerve impulse | 1.858E-03 | 3.351E-02 | 15 |
| 512 | positive regulation of cytosolic calcium ion concentration | 1.897E-03 | 3.415E-02 | 46 |
| 513 | regulation of alcohol biosynthetic process | 1.914E-03 | 3.438E-02 | 17 |

| | | | | |
|---|---|---|---|---|
| 514 | positive regulation of neurogenesis | 1.921E-03 | 3.443E-02 | 42 |
| 515 | positive regulation of heart contraction | 1.981E-03 | 3.545E-02 | 12 |
| 516 | organic hydroxy compound biosynthetic process | 1.993E-03 | 3.559E-02 | 27 |
| 517 | regulation of wound healing | 2.009E-03 | 3.575E-02 | 25 |
| 518 | transforming growth factor beta receptor signaling pathway | 2.012E-03 | 3.575E-02 | 22 |
| 519 | negative regulation of molecular function | 2.014E-03 | 3.575E-02 | 140 |
| 520 | divalent inorganic cation transport | 2.051E-03 | 3.632E-02 | 43 |
| 521 | regulation of cell growth | 2.054E-03 | 3.632E-02 | 64 |
| 522 | mesonephric tubule morphogenesis | 2.057E-03 | 3.632E-02 | 13 |
| 523 | inner ear development | 2.075E-03 | 3.656E-02 | 38 |
| 524 | lipid translocation | 2.087E-03 | 3.663E-02 | 11 |
| 525 | autonomic nervous system development | 2.087E-03 | 3.663E-02 | 11 |
| 526 | glial cell development | 2.120E-03 | 3.713E-02 | 23 |
| 527 | regulation of hydrolase activity | 2.188E-03 | 3.826E-02 | 150 |
| 528 | pharyngeal system development | 2.232E-03 | 3.888E-02 | 8 |
| 529 | male genitalia development | 2.232E-03 | 3.888E-02 | 8 |
| 530 | ganglion development | 2.240E-03 | 3.895E-02 | 7 |
| 531 | anaphase | 2.292E-03 | 3.933E-02 | 3 |
| 532 | lateral line system development | 2.292E-03 | 3.933E-02 | 3 |
| 533 | ceramide translocation | 2.292E-03 | 3.933E-02 | 3 |
| 534 | indolalkylamine biosynthetic process | 2.292E-03 | 3.933E-02 | 3 |
| 535 | His-Purkinje system development | 2.292E-03 | 3.933E-02 | 3 |
| 536 | specification of loop of Henle identity | 2.292E-03 | 3.933E-02 | 3 |
| 537 | negative regulation of female gonad development | 2.292E-03 | 3.933E-02 | 3 |
| 538 | endochondral bone morphogenesis | 2.300E-03 | 3.939E-02 | 14 |
| 539 | negative regulation of nervous system development | 2.344E-03 | 4.006E-02 | 26 |
| 540 | negative regulation of metabolic process | 2.364E-03 | 4.034E-02 | 326 |
| 541 | glial cell differentiation | 2.413E-03 | 4.073E-02 | 34 |
| 542 | response to nitrogen compound | 2.419E-03 | 4.073E-02 | 166 |
| 543 | positive regulation of heart rate | 2.422E-03 | 4.073E-02 | 9 |
| 544 | cellular response to oxygen-containing compound | 2.423E-03 | 4.073E-02 | 160 |
| 545 | pinocytosis | 2.438E-03 | 4.073E-02 | 5 |
| 546 | regulation of heart rate by cardiac conduction | 2.449E-03 | 4.073E-02 | 10 |
| 547 | renal system pattern specification | 2.458E-03 | 4.073E-02 | 4 |
| 548 | regulation of atrial cardiac muscle cell membrane repolarization | 2.458E-03 | 4.073E-02 | 4 |
| 549 | ureter morphogenesis | 2.458E-03 | 4.073E-02 | 4 |
| 550 | hematopoietic stem cell migration | 2.458E-03 | 4.073E-02 | 4 |
| 551 | vascular wound healing | 2.458E-03 | 4.073E-02 | 4 |
| 552 | regulation of mineralocorticoid secretion | 2.458E-03 | 4.073E-02 | 4 |
| 553 | pattern specification involved in kidney development | 2.458E-03 | 4.073E-02 | 4 |
| 554 | insulin secretion involved in cellular response to glucose stimulus | 2.458E-03 | 4.073E-02 | 4 |
| 555 | cellular response to UV-C | 2.458E-03 | 4.073E-02 | 4 |
| 556 | regulation of aldosterone secretion | 2.458E-03 | 4.073E-02 | 4 |
| 557 | regulation of RNA metabolic process | 2.478E-03 | 4.099E-02 | 359 |
| 558 | positive regulation of macromolecule biosynthetic process | 2.533E-03 | 4.182E-02 | 205 |
| 559 | regulation of synapse organization | 2.552E-03 | 4.206E-02 | 37 |
| 560 | amine metabolic process | 2.559E-03 | 4.210E-02 | 19 |

| | | | | |
|---|---|---|---|---|
| 561 | cytoskeleton organization | 2.580E-03 | 4.237E-02 | 129 |
| 562 | divalent metal ion transport | 2.615E-03 | 4.283E-02 | 42 |
| 563 | positive regulation of osteoblast differentiation | 2.617E-03 | 4.283E-02 | 16 |
| 564 | hindbrain development | 2.667E-03 | 4.358E-02 | 34 |
| 565 | regeneration | 2.682E-03 | 4.374E-02 | 41 |
| 566 | regulation of blood pressure | 2.722E-03 | 4.431E-02 | 36 |
| 567 | regulation of osteoblast differentiation | 2.757E-03 | 4.480E-02 | 25 |
| 568 | roof of mouth development | 2.763E-03 | 4.482E-02 | 19 |
| 569 | lipid transport | 2.794E-03 | 4.523E-02 | 48 |
| 570 | collagen metabolic process | 2.809E-03 | 4.523E-02 | 12 |
| 571 | regulation of extrinsic apoptotic signaling pathway via death domain rec | 2.809E-03 | 4.523E-02 | 12 |
| 572 | sensory perception of temperature stimulus | 2.816E-03 | 4.523E-02 | 9 |
| 573 | negative regulation of platelet activation | 2.816E-03 | 4.523E-02 | 9 |
| 574 | blood vessel morphogenesis | 2.817E-03 | 4.523E-02 | 62 |
| 575 | regulation of calcium ion import | 2.842E-03 | 4.551E-02 | 13 |
| 576 | system process | 2.846E-03 | 4.551E-02 | 262 |
| 577 | regulation of nucleobase-containing compound metabolic process | 2.850E-03 | 4.551E-02 | 382 |
| 578 | regulation of G protein-coupled receptor signaling pathway | 2.887E-03 | 4.602E-02 | 29 |
| 579 | negative regulation of peptide secretion | 2.912E-03 | 4.634E-02 | 21 |
| 580 | body fluid secretion | 2.938E-03 | 4.667E-02 | 17 |
| 581 | cellular response to glucocorticoid stimulus | 2.958E-03 | 4.686E-02 | 20 |
| 582 | formation of primary germ layer | 2.960E-03 | 4.686E-02 | 23 |
| 583 | regulation of anatomical structure size | 3.001E-03 | 4.743E-02 | 75 |
| 584 | organic hydroxy compound transport | 3.020E-03 | 4.765E-02 | 27 |
| 585 | response to retinoic acid | 3.041E-03 | 4.785E-02 | 28 |
| 586 | morphogenesis of a branching structure | 3.047E-03 | 4.785E-02 | 30 |
| 587 | synaptic vesicle cycle | 3.048E-03 | 4.785E-02 | 24 |
| 588 | regulation of cellular protein metabolic process | 3.076E-03 | 4.815E-02 | 276 |
| 589 | carbohydrate derivative metabolic process | 3.082E-03 | 4.815E-02 | 108 |
| 590 | positive regulation of gene expression | 3.083E-03 | 4.815E-02 | 254 |
| 591 | sensory system development | 3.110E-03 | 4.849E-02 | 62 |
| 592 | response to catecholamine | 3.122E-03 | 4.851E-02 | 21 |
| 593 | endochondral bone growth | 3.138E-03 | 4.851E-02 | 8 |
| 594 | aortic valve development | 3.138E-03 | 4.851E-02 | 8 |
| 595 | negative regulation of stress fiber assembly | 3.138E-03 | 4.851E-02 | 8 |
| 596 | synaptic membrane adhesion | 3.138E-03 | 4.851E-02 | 8 |
| 597 | skeletal muscle cell differentiation | 3.152E-03 | 4.864E-02 | 13 |
| 598 | branching morphogenesis of an epithelial tube | 3.166E-03 | 4.876E-02 | 26 |
| 599 | regulation of corticosteroid hormone secretion | 3.175E-03 | 4.876E-02 | 6 |
| 600 | ureter development | 3.175E-03 | 4.876E-02 | 6 |
| 601 | proximal/distal pattern formation | 3.260E-03 | 4.998E-02 | 9 |

**Table. S3D: Genes associated to dopamine neurotransmiter, dopamine metabolism and tyrosine metabolism. to dopamine metabolism and dopamine neurotransmitter.**

**Genes asociated to dopamine neurotransmiter**

| **Gene symbol** | **logFC** | **logCPM** | **F** | **PValue** | **FDR** |
|---|---|---|---|---|---|
| SLC18A2 | 3.321849 | 2.117558 | 71.60916 | 1.01E-05 | 0.001176 |
| CASK | 1.166581 | 8.312961 | 54.60319 | 3.13E-05 | 0.002264 |
| SYN1 | 1.005941 | 8.123154 | 28.19582 | 0.000409 | 0.010366 |
| PPFIA4 | 1.193029 | 5.913757 | 28.13026 | 0.000413 | 0.010415 |
| SYT1 | 1.110494 | 8.522041 | 24.22339 | 0.000705 | 0.014358 |
| SYN3 | 1.622411 | 3.488173 | 24.21831 | 0.000706 | 0.014358 |
| SYN2 | 0.813982 | 5.46717 | 21.06563 | 0.001142 | 0.019345 |
| PPFIA3 | 0.886879 | 6.100327 | 17.39529 | 0.002149 | 0.028343 |

https://pathcards.genecards.org/Card/neurotransmitter_release_cycle?queryString=dopamine%20

**Genes associated to dopamine metabolism**

| **Gene symbol** | **logFC** | **logCPM** | **F** | **PValue** | **FDR** |
|---|---|---|---|---|---|
| COMT | -1.53814 | 5.126131 | 38.89146 | 0.000122 | 0.004861 |
| MAOB | 1.33368 | 7.891897 | 22.12699 | 0.000966 | 0.017704 |
| MAOA | 0.67539 | 5.665278 | 21.76574 | 0.000994 | 0.017993 |
| DDC | 1.779821 | 5.466633 | 20.30396 | 0.001294 | 0.020787 |

https://pathcards.genecards.org/Card/dopamine_metabolism?queryString=dopamine%20

**Genes associated to tyrosine metabolism**

| Gene symbol | logFC | logCPM | F | PValue | FDR |
|---|---|---|---|---|---|
| FAHD1 | -0.90894 | 5.481232 | 41.54503 | 8.96E-05 | 0.004059 |
| COMT | -1.53814 | 5.126131 | 38.89146 | 0.000122 | 0.004861 |
| MAOB | 1.33368 | 7.891897 | 22.12699 | 0.000966 | 0.017704 |
| MAOA | 0.67539 | 5.665278 | 21.76574 | 0.000994 | 0.017993 |
| DDC | 1.779821 | 5.466633 | 20.30396 | 0.001294 | 0.020787 |
| ALDH1A3 | -1.03574 | -0.92068 | 6.049635 | 0.034908 | 0.138598 |
| ALDH1B1 | 0.363203 | 5.062321 | 5.587796 | 0.040721 | 0.151322 |
| AOX1 | 0.783245 | -0.61863 | 5.330045 | 0.044669 | 0.159441 |

**Table. S3F: neurotransmitter release.**

| Gene symbol | logFC | logCPM | F | PValue | FDR |
|---|---|---|---|---|---|
| SLC18A2 | 3.321849 | 2.117558 | 71.60916 | 1.01E-05 | 0.001176 |
| GAD1 | 1.596769 | 6.242948 | 65.96434 | 1.43E-05 | 0.001462 |
| CASK | 1.166581 | 8.312961 | 54.60319 | 3.13E-05 | 0.002264 |
| SLC5A7 | 0.908425 | 6.258985 | 44.22808 | 6.98E-05 | 0.003546 |
| SYN1 | 1.005941 | 8.123154 | 28.19582 | 0.000409 | 0.010366 |
| PPFIA4 | 1.193029 | 5.913757 | 28.13026 | 0.000413 | 0.010415 |
| NAAA | 1.225831 | 1.656574 | 26.31383 | 0.000508 | 0.011881 |
| SLC17A7 | 2.015921 | 2.879569 | 25.39056 | 0.000597 | 0.013125 |
| SYT1 | 1.110494 | 8.522041 | 24.22339 | 0.000705 | 0.014358 |
| SYN3 | 1.622411 | 3.488173 | 24.21831 | 0.000706 | 0.014358 |
| MAOA | 0.67539 | 5.665278 | 21.76574 | 0.000994 | 0.017993 |
| SYN2 | 0.813982 | 5.46717 | 21.06563 | 0.001142 | 0.019345 |
| PPFIA3 | 0.886879 | 6.100327 | 17.39529 | 0.002149 | 0.028343 |

https://pathcards.genecards.org/Card/neurotransmitter_release_cycle?queryString=dopamine%20

**Table S3:** Results from transcriptomic data analysis. (S3A) List of genes significantly deregulated (FDR < 0.05) in 3xSNCA MOs with logFC value. (S3B) List of significantly deregulated genes in 3xSNCA MOs associated to PD. (S3C) List of significantly deregulated genes in 3xSNCA MOs associated to *SNCA*. (S3D) List of significantly deregulated GO processes in 3xSNCA MOs. (S3E) List of significantly deregulated genes in 3xSNCA MOs associated to dopamine neurotransmitter, dopamine metabolism and tyrosine metabolism. (S3F) List of significantly deregulated genes in 3xSNCA MOs associated to neurotransmitter release. Information for gene-pathway association was obtained from pathcards (https://pathcards.genecards.org).

**Table S4: Antibodies used in this study.**

| Antibodies | | | | |
|---|---|---|---|---|
| **Protein** | **Dilution** | **Company** | **Catalog number** | **Technique** |
| SOX2 | 1:200 | R&D systems | AF2018 | Immunostaining |
| OCT4 | 1:400 | Abcam | ab19857 | Immunostaining |
| NANOG | 1:100 | Millipore | AB5731 | Immunostaining |
| SEEA4 | 1:25 | Millipore | MAB4304 | Immunostaining |
| TRA-1-60 | 1:25 | Millipore | MAB4360 | Immunostaining |
| TRA-1-81 | 1:25 | Millipore | MAB4360 | Immunostaining |
| FOXA2 | 1:250 | Santa Cruz | sc-6554 | Immunostaining |
| PAX6 | 1:300 | Biolegend | 901301 | Immunostaining |
| NESTIN | 1:100 | Millipore | MAB5326 | Immunostaining |
| SOX1 | 1:100 | R&D systems | AF3369 | Immunostaining |
| LMX1A | 1:100 | Sigma Aldrich | HPA030088 | Immunostaining |
| EN1 | 1:100 | Santa Cruz | sc-46101 | Immunostaining |
| DOPAMINE | 1:100 | ImmuSmol | IS1005 | Immunostaining |
| DDC | 1:100 | Thermo Scientific | PA5-25450 | Immunostaining |
| GIRK2 | 1:100 | Abcam | ab65096 | Immunostaining |
| Calbindin | 1:100 | Immunostar | 24427 | Immunostaining |
| TH | 1:1000 | Millipore | MAB5280 | WB |
| TH | 1:1000 | Abcam | ab112 | Immunostaining |
| TH | 1:600 | Abcam | ab76442 | Immunostaining |
| TUJ1 | 1:1000 | OptimAB Eurogentec | PRB-435P-050 | Immunostaining |
| TUJ1 | 1:1000 / 1:20000 | BioLegend | 801201 | Immuno / WB |
| MAP2 | 1:1000 | Abcam | ab5392 | Immunostaining |
| α-syn | 1:1000 | NOVUS Biologicals | NBP1-05194 | Immunostaining |
| α-syn | 1:1000 | Santa Cruz | sc-7011-R | WB / DotBlot |
| α-syn pS129 | 1:5000 | Cell Signaling | 23706S | Immunostaining |
| α-syn conformation | 1:100000 | abcam | ab209538 | Immunostaining |
| Synapthophysin | 1:50 | Abcam | ab8049 | Immunostaining |
| PSD-95 | 1:250 | Invitrogen | 51-6900 | Immunostaining |
| GFAP | 1:1000 | Millipore | AB5541 | Immunostaining |
| Beta-actin | 1:20000 | Cell signaling technology | 3700 | WB |

**Table S4:** List of antibodies used in this manuscript including information from the company, catalog number, optimal concentration and technique in which the antibody was used.
